# Host transcriptional responses and SARS-CoV-2 isolates from the nasopharyngeal samples of Bangladeshi COVID-19 patients

**DOI:** 10.1101/2020.07.23.218198

**Authors:** Abul Bashar Mir Md. Khademul Islam, Md. Abdullah-Al-Kamran Khan, Rasel Ahmed, Md. Sabbir Hossain, Shah Md. Tamim Kabir, Md. Shahidul Islam, A.M.A.M. Zonaed Siddiki

## Abstract

As the COVID-19 pandemic progresses, fatality and cases of new infections are also increasing at an alarming rate. SARS-CoV-2 follows a highly variable course and it is becoming more evident that individual’s immune system has a decisive influence on the progression of the disease. However, the detailed underlying molecular mechanisms of the SARS-CoV-2 mediate disease pathogenesis are largely unknown. Only a few host transcriptional responses in COVID-19 have been reported so far from the Western world, but no such data has been generated from the South-Asian region yet to correlate the conjectured lower fatality around this part of the globe. In this context, we aimed to perform the transcriptomic profiling of the COVID-19 patients from Bangladesh along with the reporting of the SARS-CoV-2 isolates from these patients. Moreover, we performed a comparative analysis to demonstrate how differently the various SARS-CoV-2 infection systems are responding to the viral pathogen. We detected a unique missense mutation at 10329 position of ORF1ab gene, annotated to 3C like proteinase, which is found in 75% of our analyzed isolates; but is very rare globally. Upon the functional enrichment analyses of differentially modulated genes, we detected a similar host induced response reported earlier; this response was mainly mediated by the innate immune system, interferon stimulation, and upregulated cytokine expression etc. in the Bangladeshi patients. Surprisingly, we did not perceive the induction of apoptotic signaling, phagosome formation, antigen presentation and production, hypoxia response within these nasopharyngeal samples. Furthermore, while comparing with the other SARS-CoV-2 infection systems, we spotted that lung cells trigger the more versatile immune and cytokine signaling which was several folds higher compared to our reported nasopharyngeal samples. We also observed that lung cells did not express *ACE2* in a very high amount as suspected, however, the nasopharyngeal cells are found overexpressing *ACE2*. But the amount of *DPP4* expression within the nasal samples was significantly lower compared to the other cell types. Surprisingly, we observed that lung cells express a very high amount of integrins compared to the nasopharyngeal samples, which might suggest the putative reasons for an increased amount of viral infections in the lungs. From the network analysis, we got clues on the probable viral modulation for the overexpression of these integrins. Our data will provide valuable insights in developing potential studies to elucidate the roles of ethnicity effect on the viral pathogenesis, and incorporation of further data will enrich the search of an effective therapeutics.

## Introduction

Since the declaration of COVID-19 pandemic on 11 March, this Severe Acute Respiratory Syndrome Coronavirus 2 (SARS-CoV-2) mediated infection has spread ~213 countries and territories [1]. Approximately, 15 million individuals across the globe have fallen victim to this virus and the number is constantly increasing at an alarming rate, as of the writing of this manuscript [1]. Though the initial fatality percentage was as low as 3.5%, currently this value lies around ~6.66% [1] and it might be increased because of the withdrawal of earlier preventing measures taken throughout the world. Coronaviruses are not new to human civilization, as these viruses caused several earlier outbreaks during the past two decades. However, none of the earlier outbreaks spread as widely as the current ongoing pandemic. As the pandemic progresses, more researches on the molecular pathobiology of the COVID-19 are being rapidly carried out to search for effective therapeutic intervention.

Coronaviruses possess single-stranded RNA (positive sense) genomes lengthening approximately 30Kb [2]. Amongst the coronaviruses, SARS-CoV-2 is a member of the betacoronaviruses having a ~29.9Kb genome which contains 11 functional genes [3]. Though SARS-CoV-2 shows similar clinical characteristics as Severe Acute Respiratory Syndrome Coronavirus (SARS-CoV) and Middle East Respiratory Syndrome-related Coronavirus (MERS-CoV) viruses, it has only ~79% and ~50% genome sequence similarities with these viruses, respectively; whereas, the genome sequence of SARS-CoV-2 is ~90% identical to that of bat derived SARS-like coronavirus [4]. Moreover, several key genomic variances between SARS-CoV-2 and SARS-CoV such as- 380 different amino acid substitutions, ORF8a deletion, ORF8b elongation, and ORF3b truncation were also reported [2].

The clinical characteristics of the COVID-19 range from mild fever to severe lung injury [5]. Some of the commonly observed mild COVID-19 symptoms are- fever, cough, and fatigue; however, complications such as- myalgia, shortness of breath, headache, diarrhea, and sore throat were also reported [6]. Furthermore, severely affected patients had exhibited respiratory complications like- moderate to severe pneumonia, acute respiratory distress syndrome (ARDS), sepsis, acute lung injury (ALI), and multiple organ dysfunction (MOD) [7]. Primarily, the lungs of the COVID-19 patients are affected [8]; however, failures of other functional systems, namely- cardiovascular system, nervous system etc. were also reported [9, 10].

Several features of the SARS-CoV-2 infection made it more complicated for effective clinical management. From the earlier studies, the incubation period of SARS-CoV-2 was reported to be around 4-5 days, however, some recent studies suggested a prolonged incubation period of 8-27 days [11]. Additionally, several cases of viral latency within the host [12], and the recurrent presence of SARS-CoV-2 in clinically recovered patients were also recorded [13, 14]. However, the detailed molecular mechanism behind these phenomena is still elusive.

In COVID-19, an increased level infection associated pro-inflammatory cytokines were recorded [15], which thereby supports the term “Cytokine storm”, that was frequently used to describe the SARS-CoV and MERS-CoV disease pathobiology [16]. This phenomenon causes the hyperactivation and recruitment of the inflammatory cells within the lungs and results in the acute lung injury of the infected patients [17]. However, this illustrates one putative molecular mechanism of COVID-19, there are many other immune regulators and host genetic/epigenetic factors which can also play significant contribution towards the disease manifestation [18, 19]. This multifaceted regulation was also reported previously for other different coronavirus infections [20]. Host-pathogen interactions in different coronavirus infections can function as a double-edged sword, as these could be beneficial not only to the hosts but also the viruses [20]. Similar host-virus tug-of-war can also occur in COVID-19 which might be contributing towards the overcomplicated disease outcomes [21].

Collectively, more than 1.7 million (almost 9% of the total infections around the globe) people have been diagnosed with COVID-19 in the South-Asian region and the number is still increasing devastatingly [1]. Recently, it has been speculated that South-Asian people might be possessing a genomic region acting as the risk factor for COVID-19 [22]. Moreover, another study suggested some genomic variations in several Indian SARS-CoV-2 isolates that might be involved in the COVID-19 pathogenesis in Indian patients [23]. However, any data suggesting the COVID-19 patients’ transcriptomic responses from this part of the globe are yet to be reported.

Several previously conducted studies reported the host transcriptional responses in SARS-CoV-2 infections using patient samples, animal models, and cell lines to explain the pathobiology of COVID-19 [24–26]. However, a detailed comparison of the host transcriptional responses between these different infection models as well as the different sites of the respiratory system is still lacking; but it might provide valuable insights on the COVID-19 pathogenesis and disease severity. In this present study, we sought to discuss the host transcriptional responses observed in COVID-19 patients in Bangladesh. Additionally, we reported the genome variations observed in the four SARS-CoV-2 isolates obtained from these patients. Finally, we illuminated the differences in host transcriptional responses in different COVID-19 infection models and further pursued to discover the putative effects of these altered responses.

## Materials and Methods

### Sample collection and virus detection by Real-time reverse transcription-quantitative PCR (RT-qPCR)

The nasopharyngeal swab samples were collected from patients suspicious of COVID-19 and placed in sample collection vial containing normal saline. Collected samples were preserved at −20°C until further use for RNA extraction and RT-qPCR assay. The RT-qPCR was performed for ORF1ab and N genes of SARS-CoV-2 using Novel Coronavirus (2019-nCoV) Nucleic Acid Diagnostic Kit (PCR-Fluorescence Probing) of Sansure Biotech Inc. according to the manufacturer’s instructions. RNA was extracted from a 20 μL swab sample through lysis with sample release reagent provided by the kit and then directly used for RT-qPCR. Thermal cycling was performed at 50=□°C for 30 min for reverse transcription, followed by 95=□°C for 1 min and then 45 cycles of 95=□°C for 15 s, 60=□°C for 30 s on an Analytik-Jena qTOWER instrument (Analytik Jena, Germany).

### RNA sequencing

Total RNA was extracted from nasopharyngeal swab samples (labeled as S2, S3, S4, S9) collected from SARS-COV-2 infected COVID-19 patients using TRIzol (Invitrogen) reagent following the manufacturer’s protocol. RNA-seq libraries were prepared from total RNA using TruSeq Stranded Total RNA Library Prep kit (Illumina) according to the manufacturer’s instructions where the first-strand cDNA was synthesized using SuperScript II Reverse Transcriptase (Thermo Fisher) and random primers. Paired-end (150 bp reads) sequencing of the RNA library was performed on the Illumina NextSeq 500 platform.

### Data processing and identification of the viral agent

Firstly, the sequencing reads were adapter and quality trimmed using the Trimmomatic program [27]. The remaining reads were mapped against the SARS-CoV-2 reference sequence (NC_045512.2) using Bowtie 2 [28]. Then the mapped reads were assembled de novo using Megahit (v.1.1.3) [29].

### Mapping of the RNA-seq reads onto SARS-CoV-2 reference genome

We mapped the normalized (by count per million mapped reads-CPM) RNA-seq reads onto the SARS-CoV-2 genome track of the UCSC genome browser [30] using the “bamCoverage” feature of deepTools2 suite [31].

### Identification of SARS-CoV-2 genome variations and variation annotation

We identified the variations within our sequenced SARS-CoV-2 genome using the “Variation Identification” (https://bigd.big.ac.cn/ncov/online/tool/variation) tool of “2019 Novel Coronavirus Resource (2019nCoVR)” portal of China National Center for Bioinformation [32]. We then annotated the variations of the isolated SARS-CoV-2 isolates using the “Variation Annotation” (https://bigd.big.ac.cn/ncov/online/tool/annotation) tool from the same portal [32]. We also gathered the global frequency of every identified variation using this same information portal [32]. Different representations showing the information regarding the variations were produced using the Microsoft Excel program [33]. The impacts of the variations were further characterized utilizing the Ensembl Variant Effect Predictor (VEP) tool [34].

### Analysis of RNA-seq expression data

We analyzed both our RNA-seq and some publicly available RNA-seq data for COVID-19 host transcriptional profile analysis. Publicly available Illumina sequenced RNA-seq raw FastQ reads were extracted from the GEO database (accessions of the data used can be found in supplementary file 1) [35]. We have checked the raw sequence quality using FastQC program (v0.11.9) [36] and found that the "Per base sequence quality", and "Per sequence quality scores" were high over the threshold for all sequences (Supplementary file 2). The mapping of reads was done with TopHat (tophat v2.1.1 with Bowtie v2.4.1) [37]. Short reads were uniquely aligned allowing at best two mismatches to the human reference genome from (GRCh38) as downloaded from the UCSC database [38]. Sequence matched exactly more than one place with equally quality were discarded to avoid bias [39]. The reads that were not mapped to the genome were utilized to map against the transcriptome (junctions mapping). Ensembl gene model [40] (version 99, as extracted from UCSC) was used for this process. After mapping, we used the SubRead package featureCount (v2.21) [41] to calculate absolute read abundance (read count, rc) for each transcript/gene associated to the Ensembl genes. For differential expression (DE) analysis we used DESeq2 (v1.26.0) with R (v3.6.2; 2019-07-05) [42] that uses a model based on the negative binomial distribution. To avoid false positive, we considered only those transcripts where at least 10 reads are annotated in at least one of the samples used in this study and also applied a minimum Log2 fold change of 0.5 for to be differentially apart from adjusted p-value cut-off of ≤ 0.05 by FDR. To assess the fidelity of the RNA-seq data used in this study and normalization method applied here, we checked the normalized Log2 expression data quality using R/Bioconductor package “arrayQualityMetrics (v3.44.0)” [43]. From these analyses, no outlier was detected in our data by “Distance between arrays”, “Boxplots”, and “MA plots” methods and replicate samples are clustered together (data not shown). We also performed a multifactorial differential gene expression analysis using the edgeR tool [44] following the experimental design-(Sample A/control for sample A)/(Sample B/control for sample B).

### Construction of phylogenetic tree

We constructed a Neighbour-Joining phylogenetic tree with all available 145 SARS-CoV-2 genomes of Bangladeshi isolates (retrieved on 6^th^ May from GISAID [45]). Firstly, the genome sequences were aligned using MAFFT [46] tool using the auto-configuration. Then we used MEGA X [47] for constructing the phylogenetic tree utilizing 500 bootstrapping with substitution model/method: maximum composite likelihood, uniform rates of variation among sites, the partial deletion of gaps/missing data and site coverage cutoff 95%.

### Functional enrichment analysis

We utilized Gitools (v1.8.4) for enrichment analysis and heatmap generation [48]. We have utilized the Gene Ontology Biological Processes (GOBP) [49], Bioplanet pathways [50], KEGG pathway [51], and Reactome pathway [52] modules for the overrepresentation analysis. Resulting p-values were adjusted for multiple testing using the Benjamin and Hochberg’s method of False Discovery Rate (FDR) [53].

### Retrieval of the host proteins that interact with SARS-CoV-2

We have obtained the list of human proteins that form high confidence interactions with SARS-CoV-2 proteins from conducted previously study [21] and processed their provided protein names into the associated HGNC official gene symbols.

### Construction of biological networks

Construction, visualization, and analysis of biological networks with differentially expressed genes, their associated transcription factors, and interacting viral proteins were executed in the Cytoscape software (v3.8.0) [54]. We used the STRING [55] database to extract the highest confidences (0.9) edges only for the protein-protein interactions to reduce any false positive connection.

## Results

### Our sequenced SARS-CoV-2 isolates showed a divergent variation pattern compared to the other Bangladeshi isolates

We sought to find out the genome variations within the four SARS-CoV-2 isolates we sequenced and pursued the deviation of these genomes compared to the other isolates in Bangladesh. To accomplish these goals, we first identified and annotated the genome variations observed within our sequenced isolates. Then we produced informative statistics from these observed variations and compared the prevalence of those with the other isolates of Bangladesh and the rest of the world.

We mapped the RNA-seq reads of each of the samples and checked their distribution athwart the entire reference genome of SARS-CoV-2 (Figure 1A). High coverages and read evidence were observed for all the isolates across the whole genome of the SARS-CoV-2 (Figure 1A). This suggests that the sequenced genomes of these isolates are of high coverage and no such region is observed without the mapped reads.

**Figure 1:**
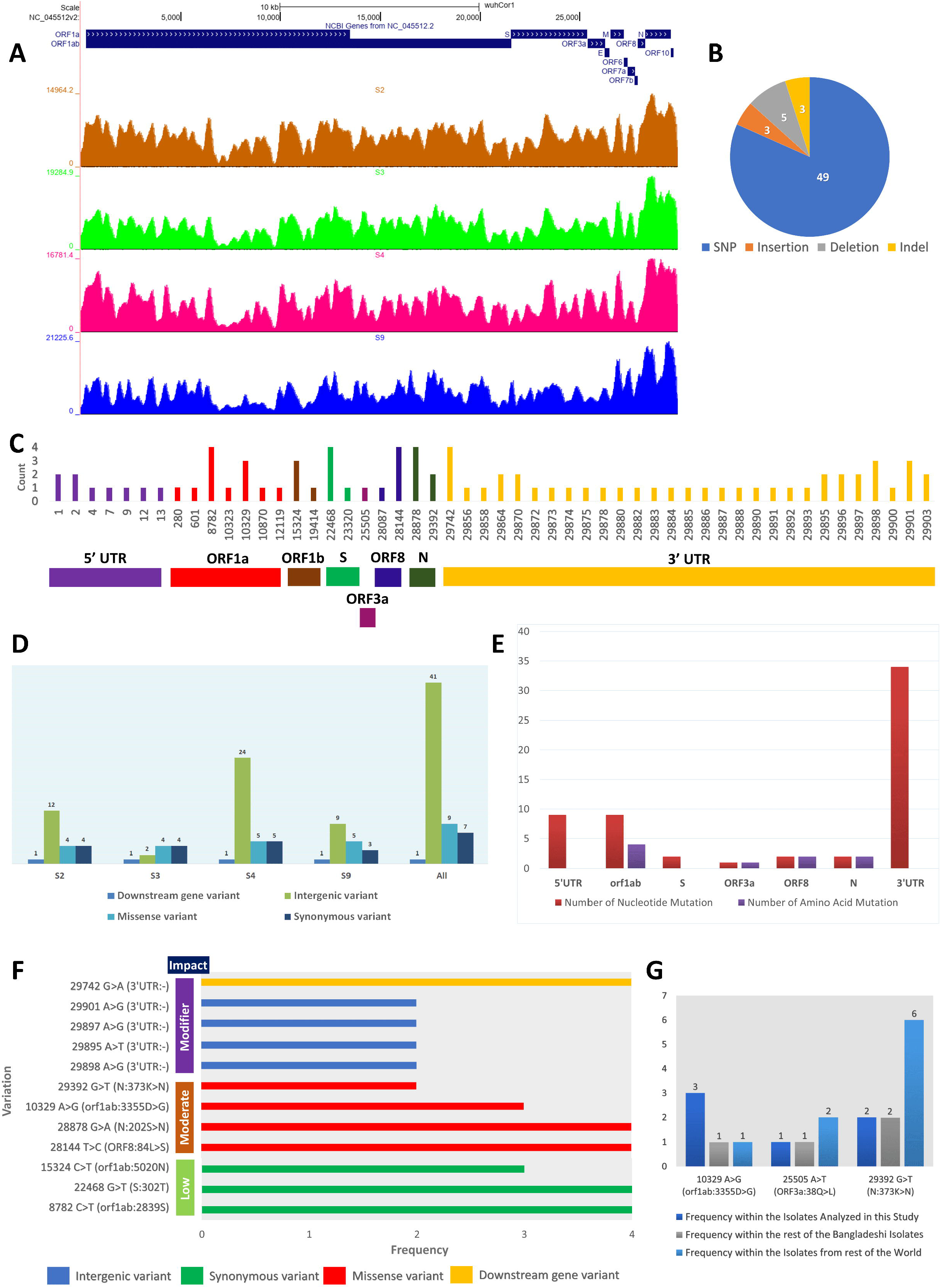
Genomic information of the sequenced SARS-CoV-2 isolates. **A.** Genome coverage normalized density map for the four sequenced SARS-CoV-2 isolates. **B.** Pie-chart illustrating the different types of variations found within these four isolates. **C.** Genome location-wise representation of the mutations and their associated frequency. **D.** Isolate-wise variation information. **E.** Gene-wise amount and type of mutations. **F.** Annotated impacts of the different mutations (only those are shown which have frequencies more than 1). **G.** Frequencies of selected unique mutations observed in these isolates.

We detected sixty different types of variations within these four analyzed SARS-CoV-2 isolates (Table 1). All the four different types of sequence variations were spotted in these isolates, however, single nucleotide polymorphisms (SNPs) were most prominent (Figure 1B). Among these variations, twelve variations were found in more than one isolate, whereas rest forty-eight variation occurred in only one isolate (Table 1, Figure 1C). Among the isolates, isolate S3 contained the lowest number of variations, whereas isolate S4 has the highest number of variations (Figure 1D). Most of these variations are intergenic variants that occurred either in the 5’-UTR or 3’-UTR regions (Figure 1C-D); whereas, globally most variations have occurred in the ORF1ab gene. Surprisingly, out of all these variations, we found only one downstream gene variation on the 3’-UTRs of all the four isolates; this variation can potentially impact the regulation of the ORF10 gene (Figure 1D). Most of the nucleic acid mutations were located on the 3’-UTR of the isolates, whereas the ORF1ab gene contained most of the amino acid mutations (Figure 1E).

**Table 1:**
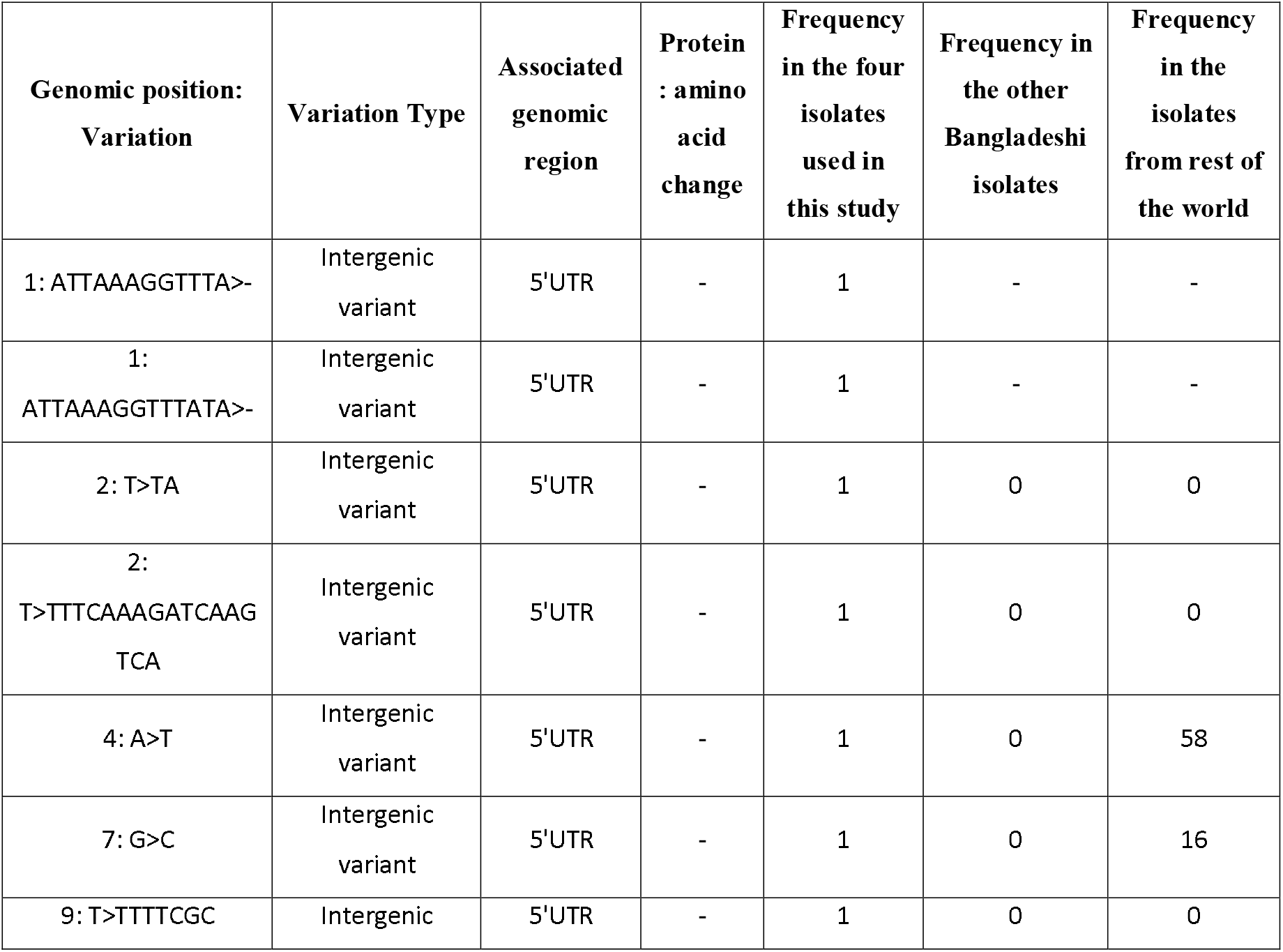
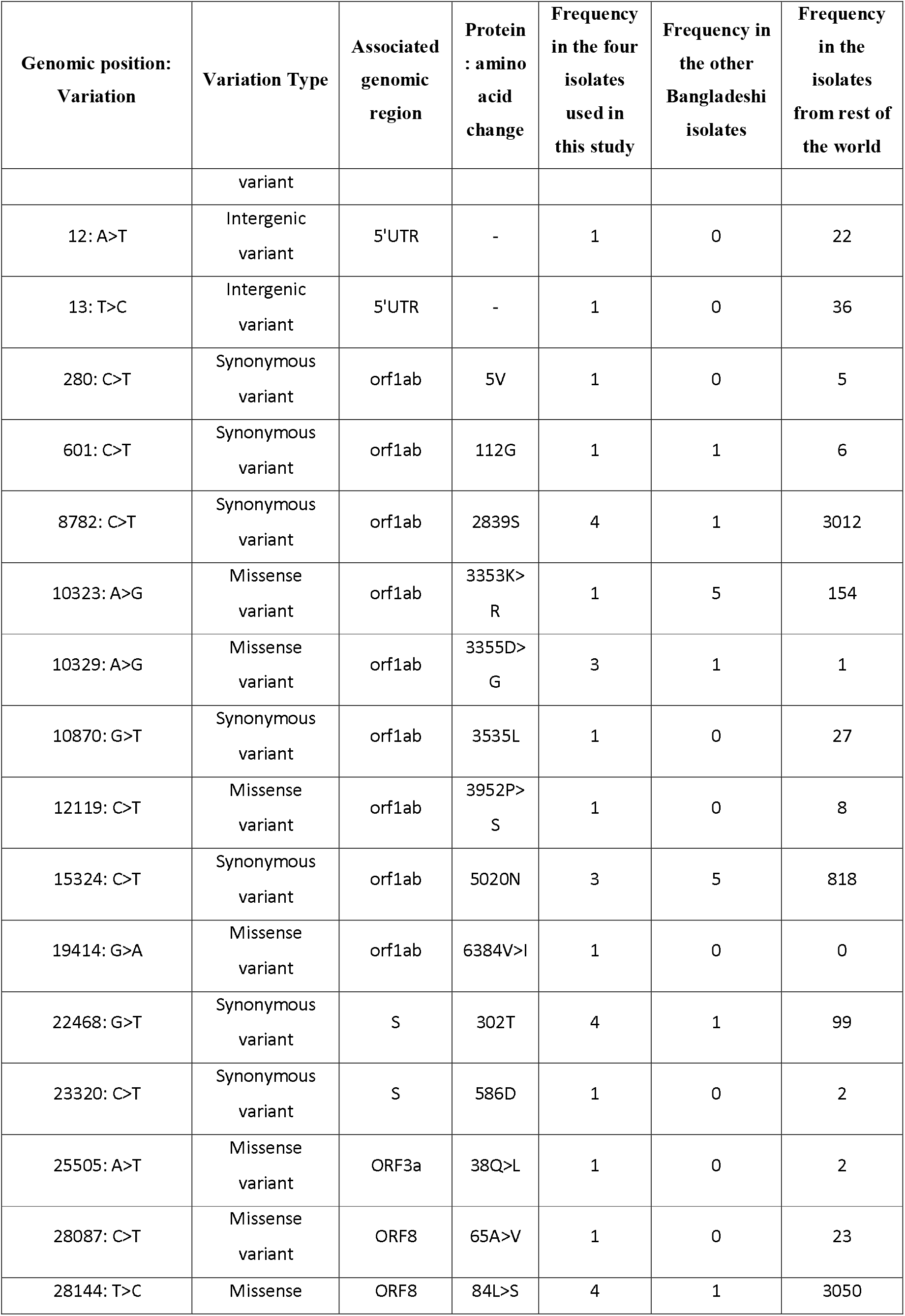
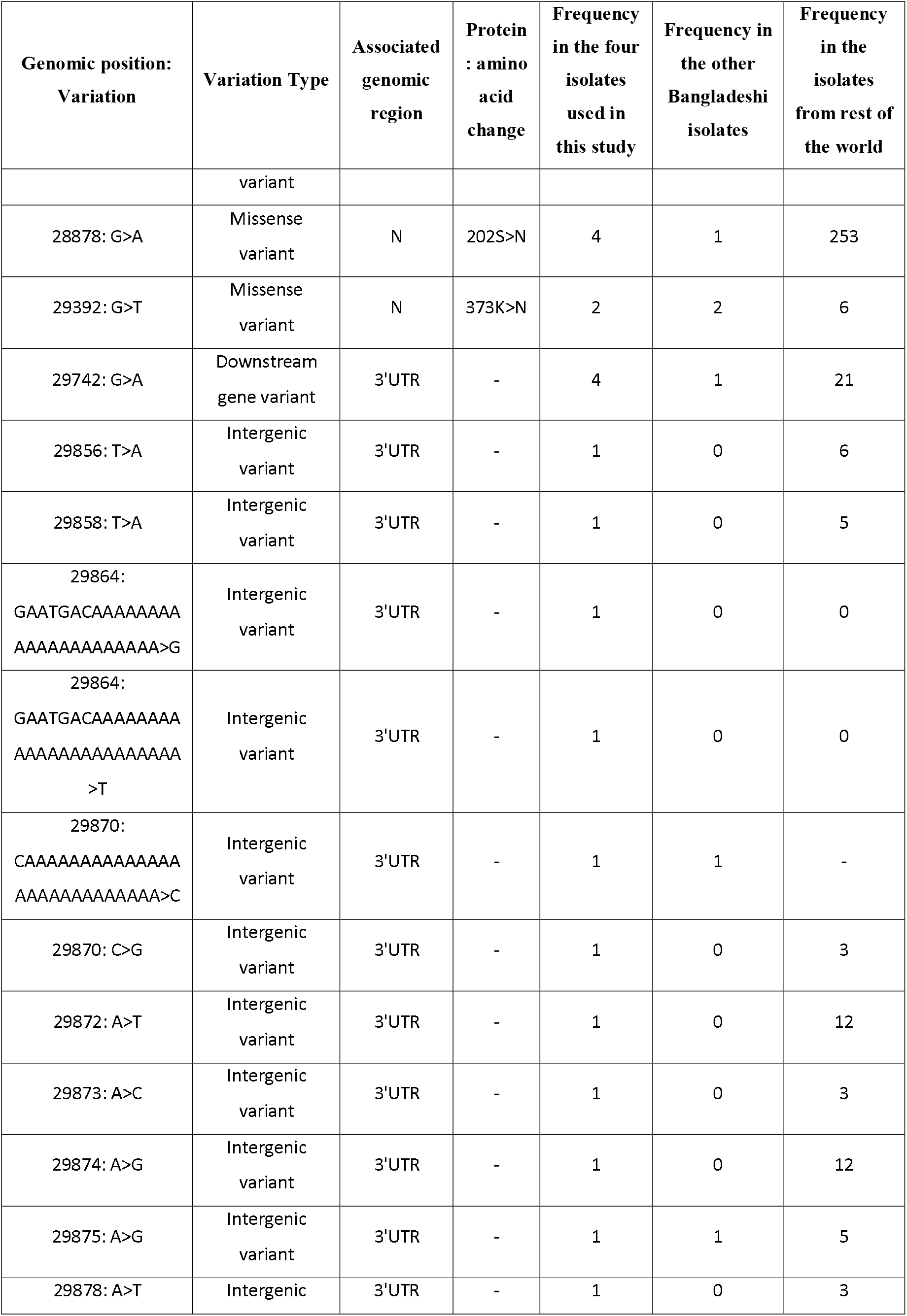
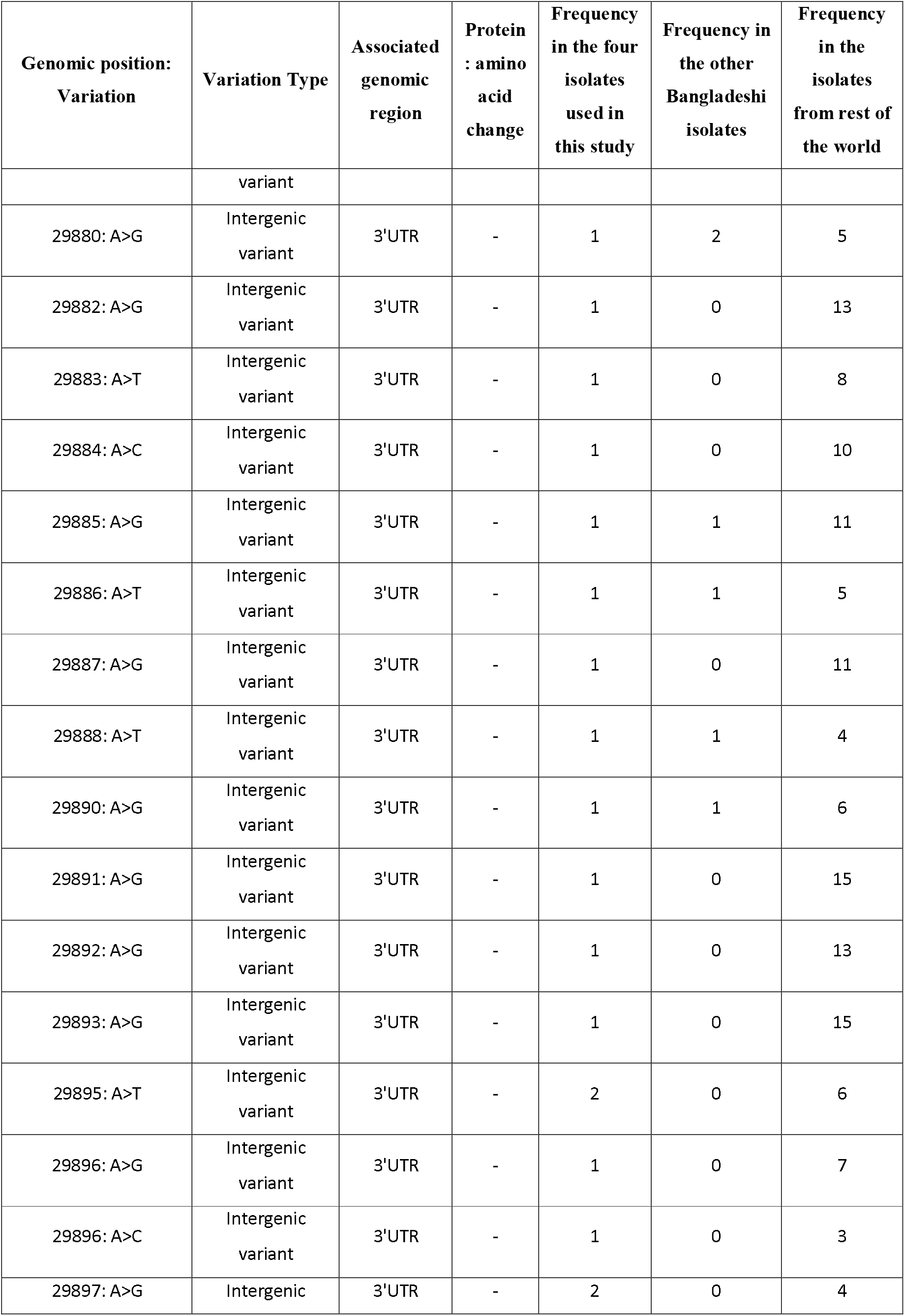
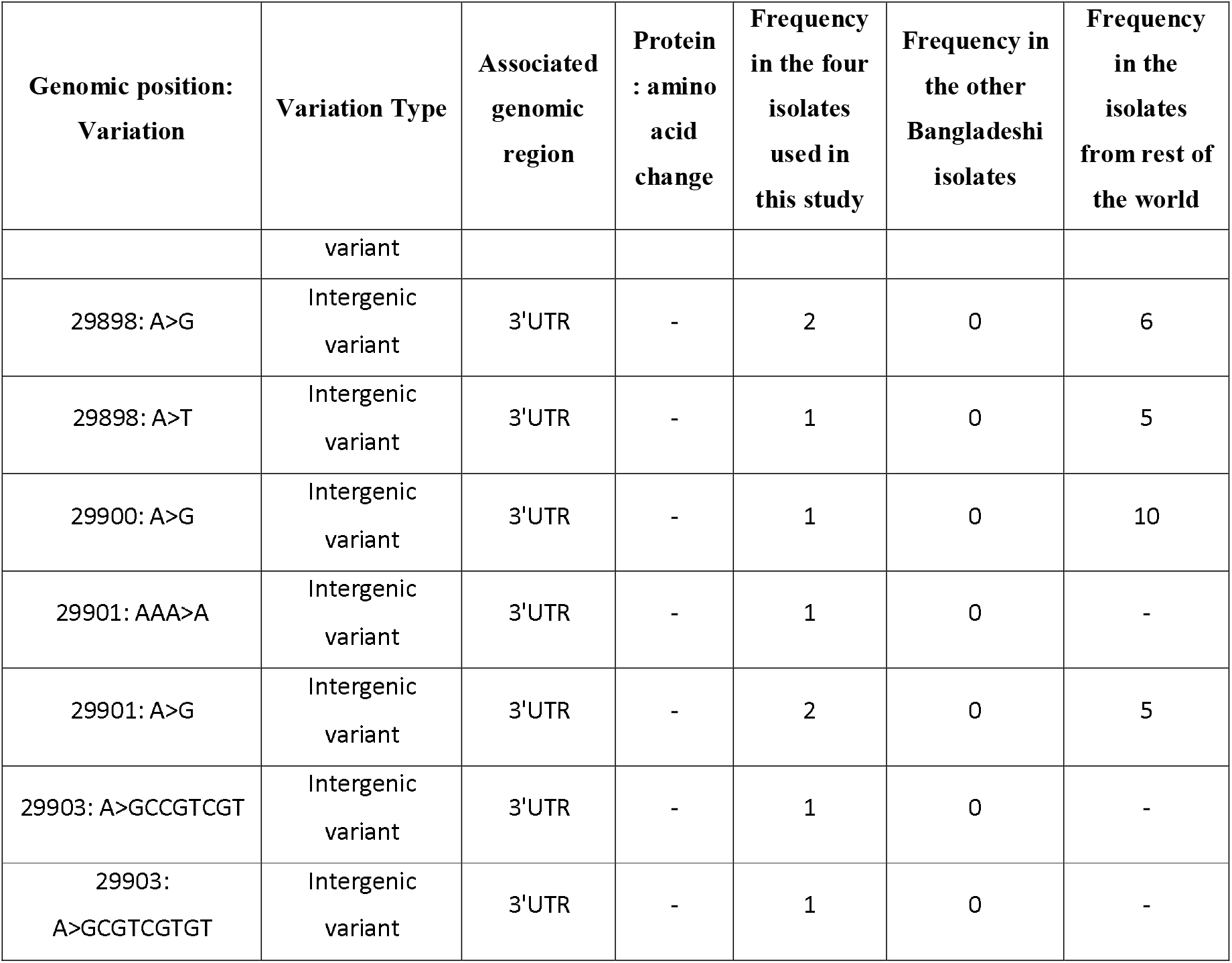
Observed variations within the four Bangladeshi SARS-CoV-2 isolates reported in this study.

No highly severe mutation was identified amongst these variations, but we found nine moderately impacting, seven low impacting, and forty-seven modifier variations within these isolates (Figure 1F, Supplementary file 3). As of 8^th^ July, thirty-eight out of the sixty variations within our sequenced isolates were completely absent in all other Bangladeshi SARS-CoV-2 isolates (Table 1). Strikingly, we observed that variation 10329: A>G is present within three of our sequenced isolates, only one other Bangladeshi and one other USA isolate contain this variation (Figure 1G). This variation is located within the 3C-like protease of SARS-CoV-2. Previously, the potential implication of the mutations of this protein was reported to alter its overall structure and functionality [56–58] in SARS-CoV. Also, few of our reported variations like 25505: A>T and 29392: G>T are not highly prevalent globally (Figure 1G).

Exploring the Nextstrain portal [59], we noticed that our analyzed SARS-CoV-2 sequences are closely placed to the Saudi-Arabian isolates (Supplementary Figure 1); although, most of the other Bangladeshi isolates were placed in the major European clusters (data not shown). Furthermore, these isolates analyzed in this study is distinctly placed in our constructed Neighbor-Joining phylogenetic tree (Figure 2), this also supports the differences between these isolates and other Bangladeshi SARS-CoV-2 isolates which might have been originated from the European nations.

**Figure 2:**
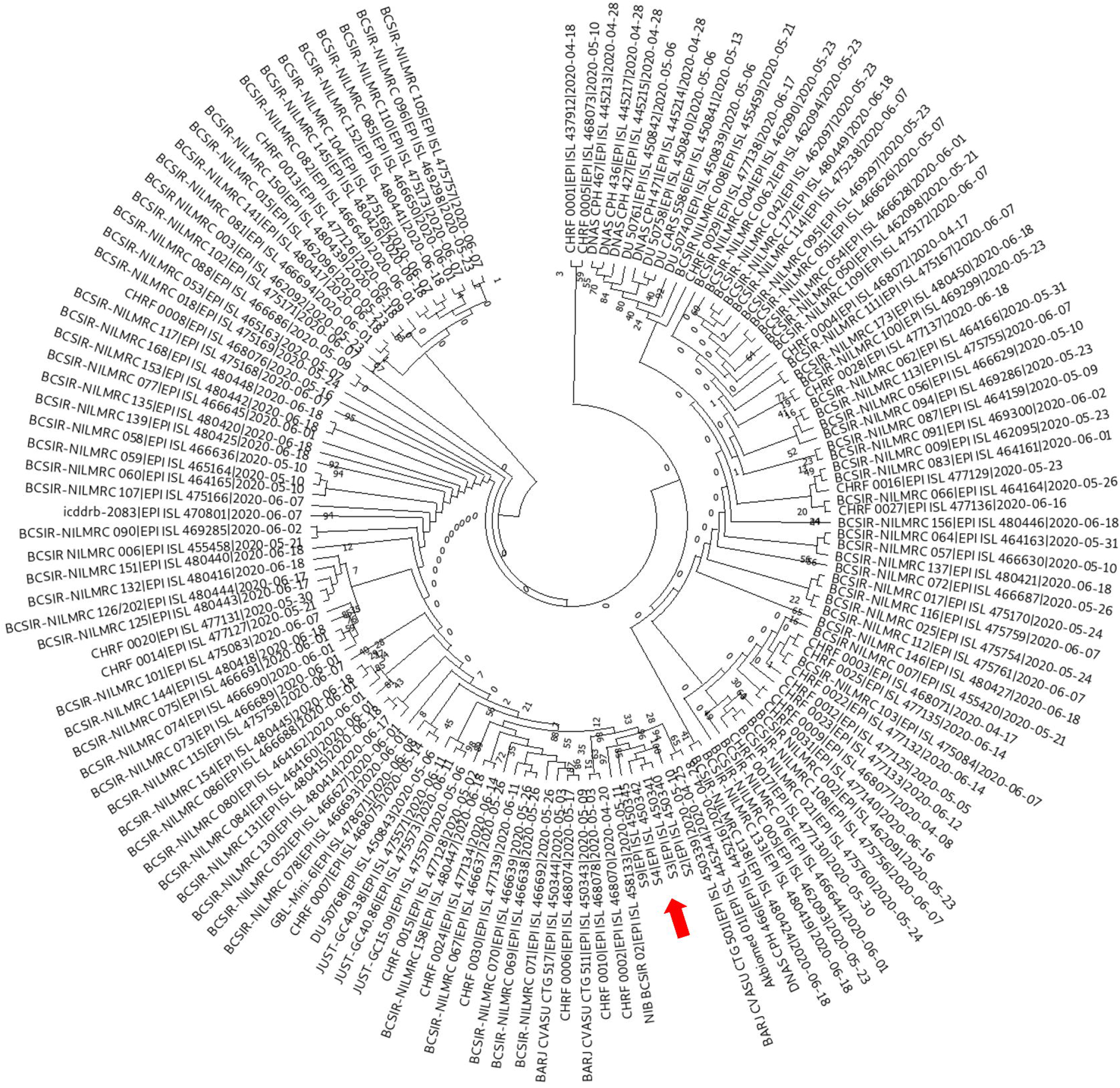
Phylogenetic tree of Bangladeshi SARS-CoV-2 isolates. Neighbor-joining tree using MEGA tools. Isolates reported in this study are indicated with a red arrow. The evolutionary history was inferred using the Neighbor-Joining method. The optimal tree with the sum of branch length = 0.01403419 is shown. The percentage of replicate trees in which the associated taxa clustered together in the bootstrap test (500 replicates) are shown next to the branches. The evolutionary distances were computed using the Maximum Composite Likelihood method and are in the units of the number of base substitutions per site. This analysis involved 145 nucleotide sequences. Codon positions included were 1st+2nd+3rd+Noncoding. All positions with less than 95% site coverage were eliminated, i.e., fewer than 5% alignment gaps, missing data, and ambiguous bases were allowed at any position (partial deletion option). There was a total of 29827 positions in the final dataset. Values represent bootstrap numbers (%).

### Stimulated antiviral immune responses are detected in the nasopharyngeal samples of Bangladeshi COVID-19 patients

Though initial researches suggested the potential implication of viral variations on the COVID-19 disease severity, one recent study indicated otherwise; Several host factors such as- abnormal immune responses, cytokine signaling etc. might be influencing the overall disease outcomes more prominently compared to the viral mutations [60]. Moreover, several data surmised that ethnicity might be a pivotal risk factor of being susceptible to COVID-19 [61].

In this context, we explored the transcriptome data obtained from the nasopharyngeal samples from Bangladeshi COVID-19 patients to find out how these patients were responding against the invading SARS-CoV-2. We compared the RNA-seq data of these patients with some random normal individuals’ nasopharyngeal RNA-seq data to find out the differentially expressed genes within our analyzed samples. We observed a roughly constant standard deviation for the normalized reads suggesting a lesser amount of variation occurred during the normalization (Figure 3A). Furthermore, we performed sample clustering to assess the quality of our generated normalized RNA-seq data. No anomalies were observed in the sample to sample distance matrix (Figure 3B) and principal component analysis (PCA) (Figure 3C) while comparing our samples with the used healthy individuals’ data. Moreover, the larger differences observed in the PCA plot (Figure 3C) and clustered heatmap of the count matrix with the top 50 significant genes (Figure 3D) suggest a significant transcriptomic response difference between our infected patients’ data and the normal individuals’ data. Likewise, the sample to sample distance plot suggested the similarities of samples of similar nature; the infected and healthy samples were clustered into two distinct groups (Figure 3B).

**Figure 3:**
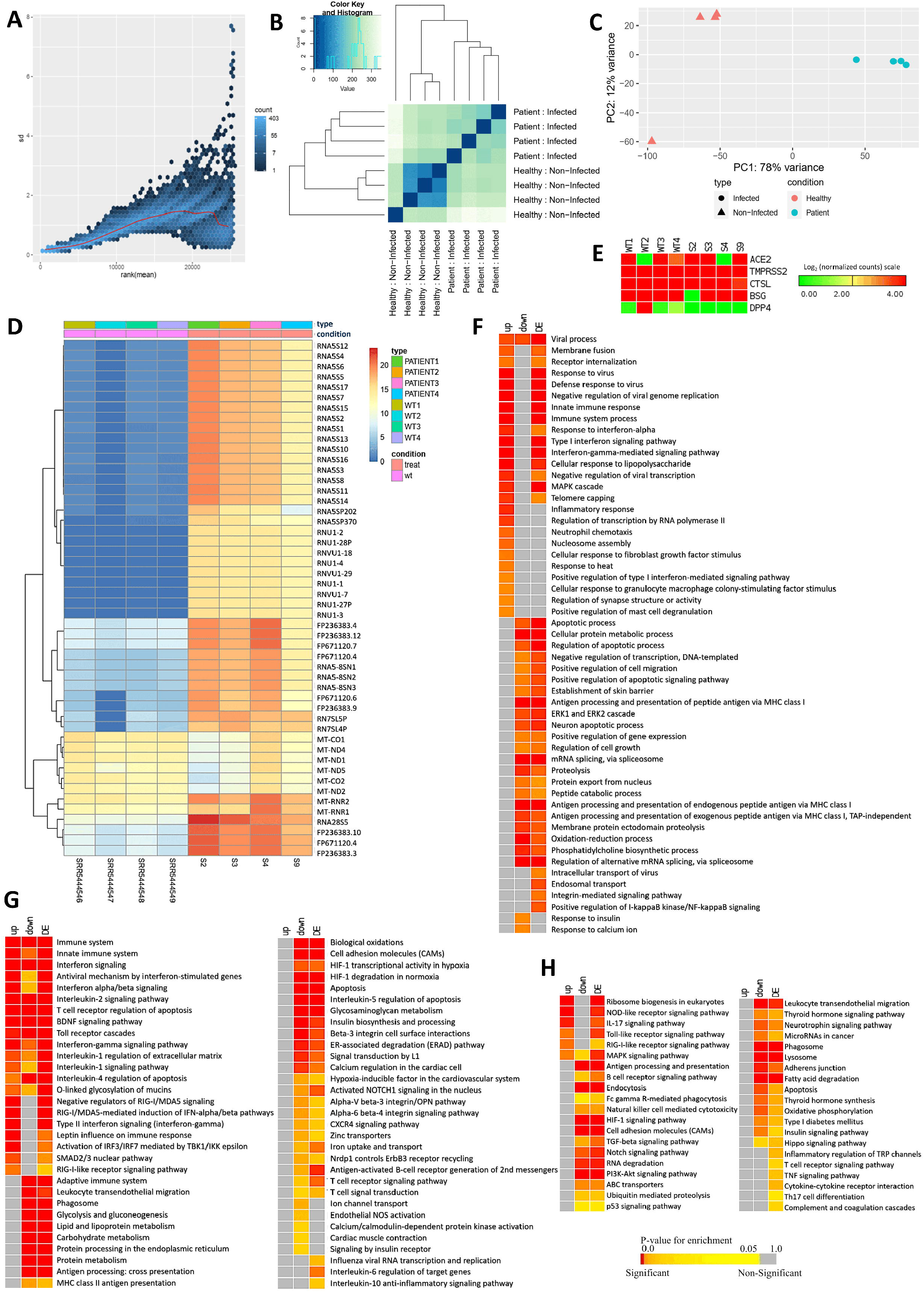
**A.** Variance plot. This plots the standard deviation of the transformed data, across samples, against the mean, using the variance stabilizing transformation. The vertical axis in the plots is the square root of the variance over all samples. **B.** Sample to sample distance plot. A heatmap of distance matrix providing an overview of similarities and dissimilarities between samples. Clustering is based on the distances between the rows/columns of the distance matrix. **C.** Principal component analysis plot. Samples are in the 2D plane spanned by their first two principal components. **D.** Clustered heatmap of the log_2_ converted normalized count matrix RNA-seq reads, top 50 genes, of nasopharyngeal samples. **E.** Normalized Log_2_ read counts of the genes encoding SARS-CoV-2 receptor and entry associated proteins. Enrichment analysis and comparison between deregulated genes and the genes of some selected processes in SARS-CoV-2 infected nasopharyngeal samples and SARS-CoV-2 infected lung biopsy samples using **F.** GOBP module, **G.** KEGG pathway, **H.** Bioplanet pathway module. Selected significant terms are represented in heatmaps. Significance of enrichment in terms of the adjusted p-value (< 0.05) is represented in color-coded P-value scale for all heatmaps; Color towards red indicates higher significance and color towards yellow indicates less significance, while grey means non-significant. Normalized Log_2_ converted read counts are considered as the expression values of the genes and represented in a color-coded scale; Color towards red indicating higher expression, while color towards green indicating little to no expression.

Sungnak et al. described the significance of several viral entry associated host proteins in SARS-CoV-2 pathogenesis, namely-ACE2, TMPRSS2, BSG, CTSL, DPP4 [62]. We also investigated the expression of the associated transcripts of these proteins within our patients’ samples. We spotted that both the healthy and infected samples have expressed these genes except the *DPP4* gene (Figure 3E).

We identified 1,614 differentially expressed genes within our reported four SARS-CoV-2 infected nasopharyngeal samples; among these differentially expressed genes, 558 genes were upregulated, and 1056 genes were downregulated (Supplementary file 4). Then we sought to discover in which biological functions/pathways these deregulated genes might be involved. To achieve this, we performed functional enrichment analyses with the observed deregulated genes using different ontology and pathway modules.

Several GOBP terms related to antiviral immune responses such as- viral process, defense response to virus, innate immune response, inflammatory response, negative regulation of viral transcription, negative regulation of viral genome replication etc. were observed enriched for the upregulated genes (Figure 3F, Supplementary Figure 2). Surprisingly, several other important antiviral defense related functions such as- apoptosis, antigen processing, and presentation etc. were found enriched for downregulated genes (Figure 3F). Similarly, this pattern was also observed for the functional enrichment using KEGG and Bioplanet pathways modules. Upregulated genes are observed involved in signaling pathways such as- innate immune system, antiviral mechanism by interferon-stimulated genes, interleukin-2 signaling, interferon-gamma signaling, interferon alpha-beta signaling, antiviral mechanism by interferon stimulated genes, IL-17 signaling pathway, Toll-like receptor signaling pathway, RIG-I like receptor signaling pathway, and MAPK signaling pathway, etc. (Figure 3G-H, Supplementary Figure 2). Strikingly, several important antiviral signaling pathways such as- antigen processing and presentation, apoptosis, HIF-1 signaling pathway, Natural killer cell mediated cytotoxicity, phagosome, PI3K-Akt signaling pathway, Interleukin-6 regulation of target genes, and Interleukin-10 inflammatory signaling pathway were enriched for the downregulated genes (Figure 3G-H). This unusual observation made us curious to search for a similar pattern of deregulated host responses in several other COVID-19 disease models.

### Host responses observed in nasopharyngeal samples are significantly different compared to the other SARS-CoV-2 infections models

We sought to compare the host responses of our analyzed samples with several other different SARS-CoV-2 infection models (two different lung biopsy samples from COVID-19 patients and two different cell lines). We performed functional enrichment analyses using differentially expressed genes from four other SARS-CoV-2 infection systems and compared the enriched terms of our samples with these four other samples. Moreover, how the host responds differently in different tissue types were also evaluated. To achieve these goals, we identified the differentially expressed genes across these different samples and systematically compared the enrichment results of those deregulated genes.

Using the similar parameterization of the differential gene expression analyses, we identified 6714 genes in lung cells (GSE147507), 232 genes in lung cells (GSE150316), 143 genes in NHBE cells (GSE147507), and 5637 genes in Calu-3 cells (GSE148729) as differentially expressed compared to their respective healthy controls (Supplementary file 5). Significant proportions of the deregulated genes detected in our nasopharyngeal samples are also found deregulated in lung (GSE147507) and Calu-3 cells (GSE148729) samples (Figure 4A), while a small number of our samples’ deregulated genes were also observed deregulated in rest of the two samples used (Figure 4A).

**Figure 4:**
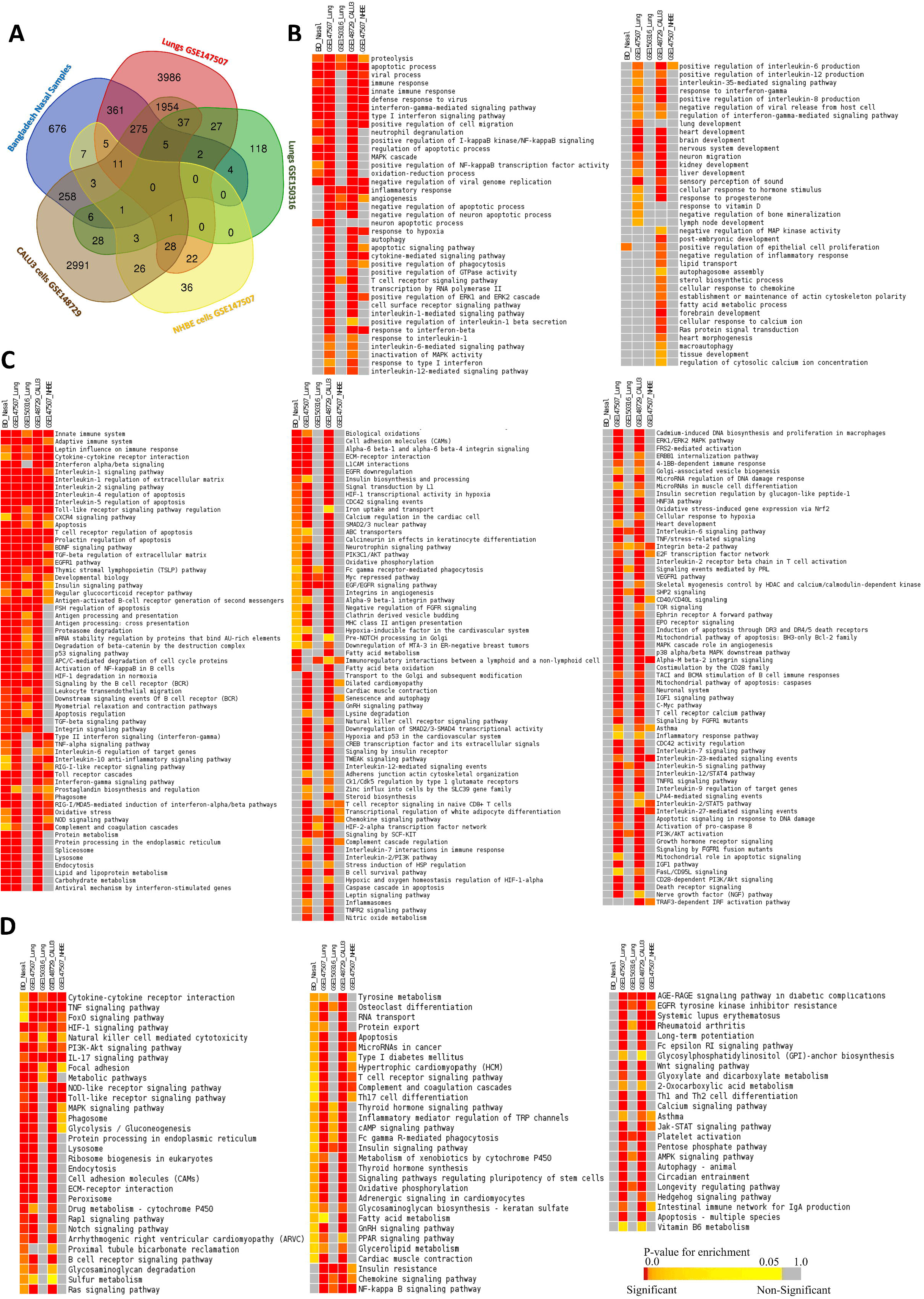
Comparison of the gene expression patterns in different SARS-CoV-2 infection models. **A.** Venn-diagram showing the observed deregulated genes (with their respective control) in the different cell types. Enrichment analysis and comparison between deregulated genes in different SARS-CoV-2 infection models using **B.** GOBP module, **C.** Bioplanet pathway module, **D.** KEGG pathway module. Selected significant terms are represented in heatmaps. Color scale/schemes are similar to Figure 3.

Enrichment analysis using these deregulated genes suggested the host response differences among the different infection systems used (Figure 4B-D). Upon the analysis, only a few GOBP terms were found enriched for both our samples, lung (GSE147507), and Calu-3 cells (GSE148729) samples, such as- viral process, immune response, innate immune response, defense response to virus, interferon signaling etc. (Figure 4B). However, genes in many important antiviral immune response related functions were not significantly enriched for our samples but were enriched for the lung (GSE147507), and Calu-3 cells (GSE148729) samples; these processes are-autophagy, apoptotic signaling pathway, interleukin-6 mediated signaling pathway, interleukin-12 mediated signaling pathway, cytokine-mediated signaling pathway, inflammatory response etc. (Figure 4B). Moreover, processes like- response to hypoxia, response to vitamin-D, and lung development were also not enriched for the deregulated genes of our nasal samples (Figure 4B).

We noticed several commonly enriched important immune signaling pathways for most of the samples used for the comparison (Figure 4C-D), such as adaptive immune system, innate immune system, interferon signaling, apoptosis, Toll-like receptor signaling pathway regulation, antigen processing and presentation, integrin signaling pathway, RIG-I like receptor signaling pathway, phagosomes etc. (Figure 4C-D, Supplementary Figure 3); however, pathways like JAK-STAT signaling pathway, Natural killer cell mediated cytotoxicity, NF-κB signaling pathway, asthma, PI3K-Akt pathway, cellular response to hypoxia, inflammasomes, and inflammatory response pathway etc. (Figure 4C-D, Supplementary Figure 3) were not enriched for the deregulated genes of our nasopharyngeal samples. These results suggest that host responses observed in nasopharyngeal samples have a different host response compared to the other infection systems. Therefore, to unveil the mystery behind this observation, we further analyzed these data to compare the gene expression patterns in different specific functionalities.

### Significant gene expression differences were spotted between the nasopharyngeal samples and lung biopsy samples

We compared the normalized read counts of each sample without integrating the respective controls to shed insights on the differences in gene expression patterns between the individual samples and tissues. A constant standard deviation was observed for the normalized read counts of the infected samples (Figure 5A) indicating the acceptability of the normalized reads for analysis. From the sample to sample distance clustering, principal component analysis, and clustered heatmap of the count matrix with top 50 genes, we observed that gene expression profiles of our nasopharyngeal samples are more relevant to that of lung samples; whereas, high level of variance was observed between the gene expression counts of the cell lines and primary nasopharyngeal samples (Figure 5B-D). Furthermore, we had a similar observation from the clustered normalized read counts of the samples based on Pearson’s correlation distance with all genes that vary across samples (Figure 5E). We then narrowed down our searches to the sample level gene expression profiles of several COVID-19 related important biological functions within these samples (Figure 6), to understand the gene expression similarities and dissimilarities among these infections systems, specially comparing nasal and lung tissues.

**Figure 5:**
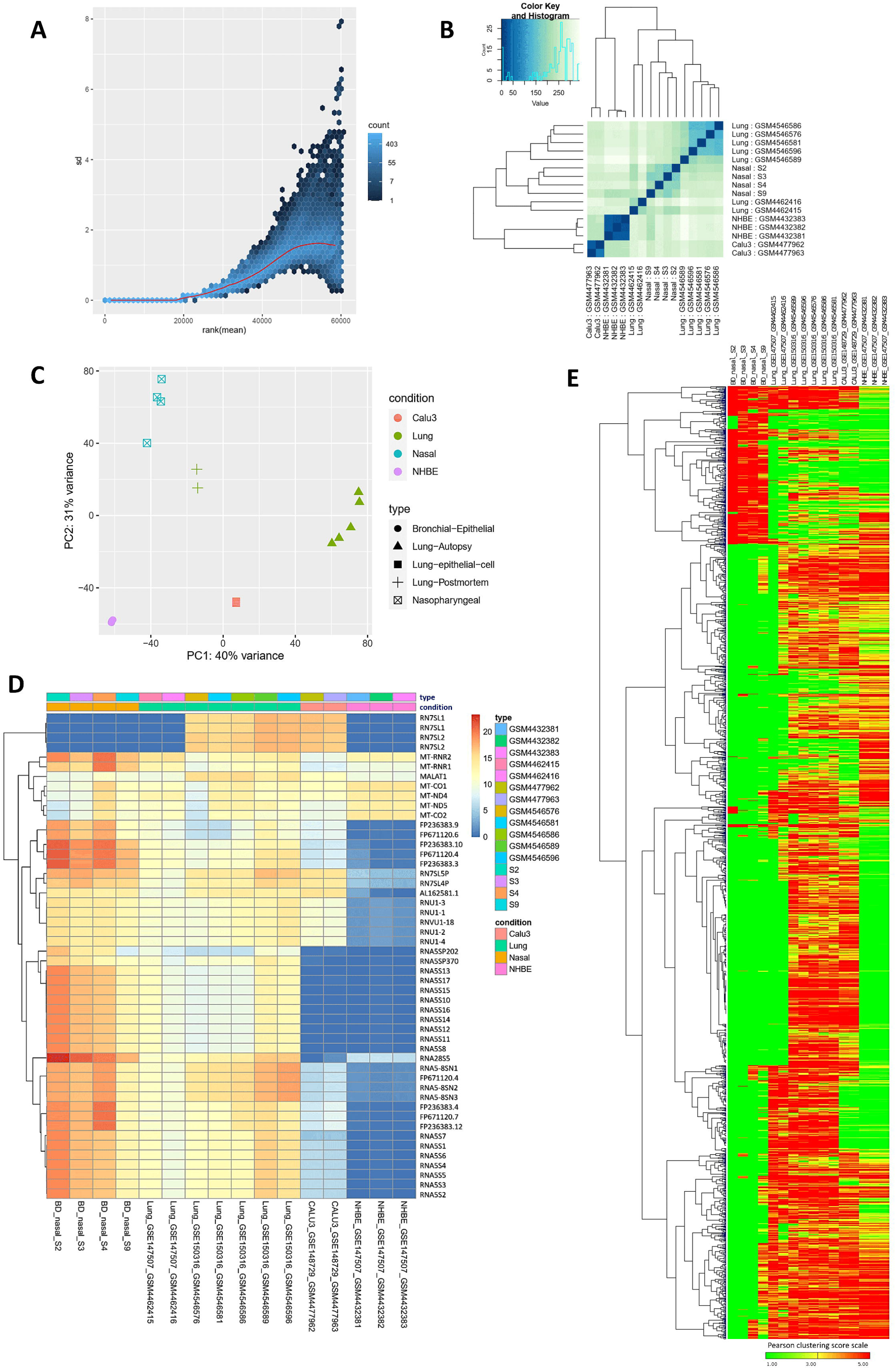
**A.** Variance plot, **B.** Sample to sample distance plot, **C.** Principal component analysis plot, **D.** Clustered heatmap of the count matrix of the normalized RNA-seq reads of different SARS-CoV-2 infection samples using to 50 genes. **E.** Gene expression heatmap showing global gene expression profiles in the individual infected samples of the various infection system. Heatmap is clustered based on Pearson’s distance with genes that vary across the sample, leaving out genes that do not vary significantly.

**Figure 6:**
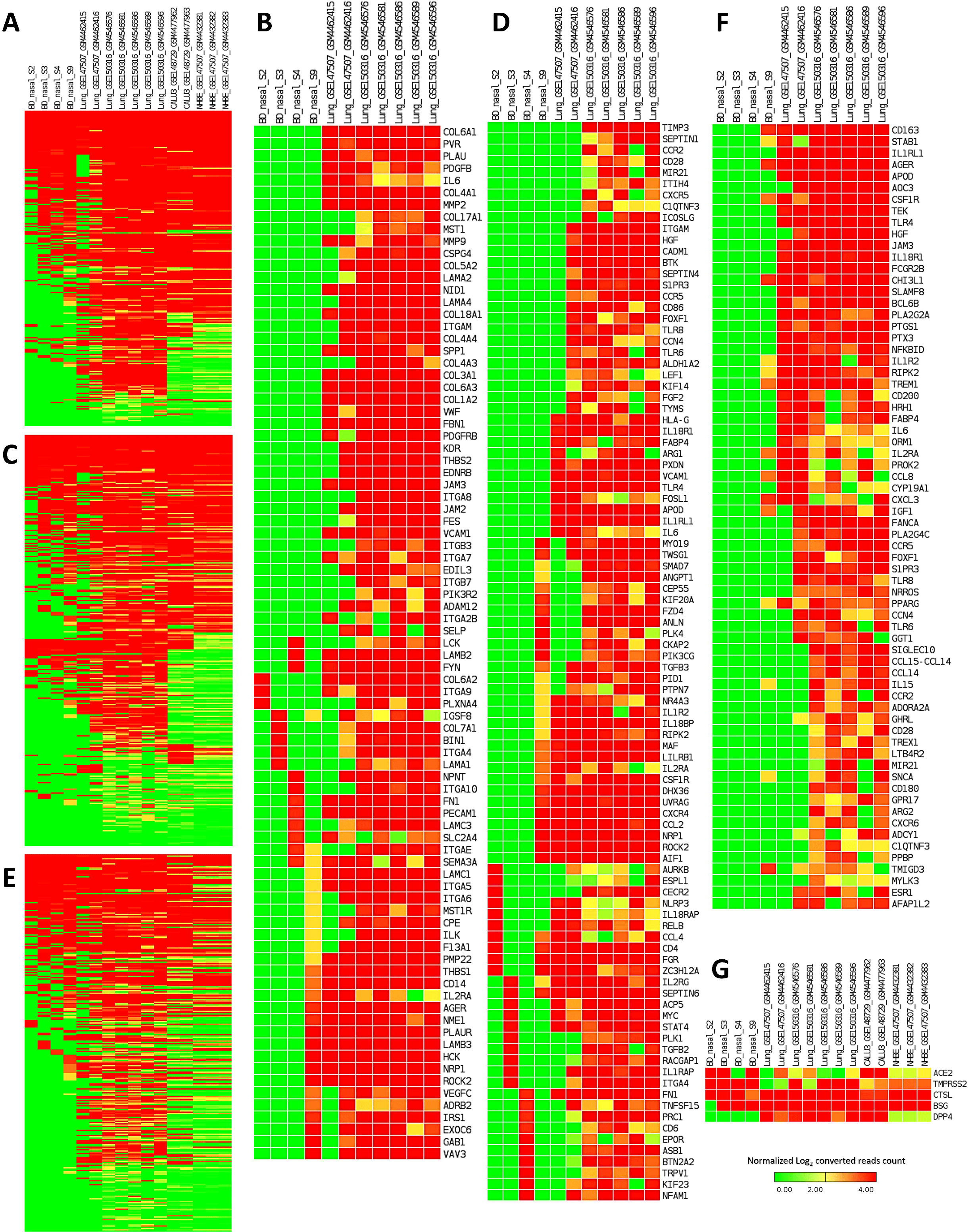
Heatmaps representing the sample level absolute expression of Integrin related genes. **A.** across the different SARS-CoV-2 infection models, **B.** in only nasopharyngeal samples and lung samples; Cytokine signaling related genes **C.** across the different SARS-CoV-2 infection models, **D.** in only nasopharyngeal samples and lung samples; and Inflammation related genes **E.** across the different SARS-CoV-2 infection models, **F.** in only nasopharyngeal samples and lung samples; **G.** Expression profiles of genes encoding SARS-CoV-2 receptor and entry associated proteins. Normalized (DESeq2) Log_2_ converted read counts are considered as the expression values of the genes and represented in a color-coded scale; Color towards red indicating higher expression, while color towards green indicating little to no expression.

### Genes related to integrins and integrin signaling pathway are highly expressed in lung samples compared to the nasopharyngeal samples

Several previous reports suggested an important aspect of integrins in SARS-CoV-2 pathogenesis, therefore, we sought to find out the expression profiles of integrin related genes in different COVID-19 infection models at sample level [63, 64]. RGD motif of the spike protein of SARS-CoV-2 can bind the integrins and this motif is placed near to the ACE2-receptor binding motif [63]. Moreover, evidence of integrin domain binding was also reported for SARS-CoV [65]. Therefore, we sought to discover the expression profiles of the integrin related genes in different SARS-CoV-2 infection models. To accomplish this, we filtered out the integrin and integrin signaling related genes (Supplementary file 6) within the terms of the GOBP, KEGG pathway, and Bioplanet pathway modules that we used for enrichment analysis. Intriguingly, we observed that the genes related to integrins and integrin signaling were highly expressed in analyzed lung samples, and the lowest number of these genes were expressed in the nasopharyngeal samples (Figure 6A-B, Supplementary Figure 4A). Based on these observations, we can assume that overexpression of integrins and integrin signaling related genes in the lungs might provide the virus a competitive edge in invading the lung cells more efficiently compared to the cells of the nasopharynx and respiratory tracts.

### Cytokine and inflammatory signaling genes are overexpressed in lung samples

Aberrant cytokine stimulation and inflammatory responses are thought to be the major contributor to pathogenic lung damages in severely affected COVID-19 patients [66, 67]. We wanted to find out whether the genes related to cytokine signaling and inflammation have differential expression profiles in lung cells compared to the other infection systems. We extracted and compared the gene expression values of the genes related to these two terms (Supplementary file 6). We are not surprised to observe that the genes of these two major contributing events of COVID-19 lung pathobiology are significantly overexpressed in lung samples compared to the rest of the SARS-CoV-2 infected cell types (Figure 6C-F, Supplementary Figure 4B-C). Particularly, the analyzed nasopharyngeal samples have very low expression values for the cytokine and inflammatory signaling genes (Figure 6C-F). Therefore, these observations are fueling the preexisting supposition of the roles of enhanced cytokine, and inflammatory signaling for worsening the disease condition in patients with SARS-CoV-2 infected lungs.

### A differential gene expression profile was detected for the SARS-CoV-2 entry receptors/associated proteins in different infection models

Expression of receptor protein ACE2 and entry associated proteins such as- TMPRSS2, BSG, CTSL, DPP4 on the cell surface of the host is essential for the invasion of SARS-CoV-2 [62]. Moreover, ACE2 overexpression is thought to increase the infection potentiality of SARS-CoV-2 [68]. Furthermore, Kuba et al. demonstrated the potential role of ACE2 in SARS-CoV induced lung injury [69]. So, we ventured to check the gene expression levels of ACE2 and the other entry associated proteins in the different SARS-CoV-2 infected cells. Surprisingly, we observed that the *ACE2* gene is not expressed in high levels in lung samples as speculated (Figure 6G). However, gene expression levels of the other entry associated proteins were higher in lung samples (Figure 6G). Nonetheless, in few of the lung samples, the *TMPRSS2* gene was not expressed in higher amounts (Figure 6G). Interestingly, we have not detected any expression of *DPP4* gene within the Bangladeshi nasopharyngeal samples (Figure 6G).

### Inflammatory immune responses were several folds higher in lungs than the nasopharynx of COVID-19 patients

From our previous observations, it was evident that COVID-19 patient’s lung responds to the viral infection differently compared to the epithelial cells of nasopharynx. We then sought to figure out the specific genes and biological functions/signaling pathways which have this differential pattern. We achieved this by designing a multifactorial differential gene expression analysis using a generalized linear model (GLM) [44]; in which we compared the fold changes of every differentially expressed gene in nasopharyngeal and lung (GSE150316) samples, to discover how many folds lung is alternatively expressing the genes than nasopharynx in COVID-19.

Firstly, we analyzed the suitability of the data for this design and observed no irregularities between the data used (Figure 7A-D). Moreover, upon this multifactorial differential gene expression analysis, we observed an acceptable common biological coefficient of variation; this variation decreases significantly as the expression values increases (Figure 7E). From the MA plot, we observed a very high amount of the significantly (p-value < 0.05) several fold upregulated and downregulated genes in lungs compared to nasopharyngeal samples (Figure 7F). We detected 807 upregulated and 298 downregulated genes in lungs compared to the nasopharyngeal samples (Supplementary file 7). Interestingly, we noticed the highly upregulated integrin and integrin signaling genes in lungs compared to the nasal samples (Figure 7G) which are consistent with our previous observations. Modulatory roles of integrins are well established in acute lung damages [70]. Similarly, aberrant expression of genes involved in integrin signaling can also provoke acute lung injuries, namely-*ADAM15* [71], *SDC1* [72], *CD14* [73], *CD47* [74], *CD9* [75], *HMGB1* [76], *ITA6* [77], and *ITAV* [78] etc. Therefore, SARS-CoV-2 infection induced deregulation of these genes might be contributing towards the worsening of the normal pathobiology and functionality of lungs in COVID-19.

**Figure 7:**
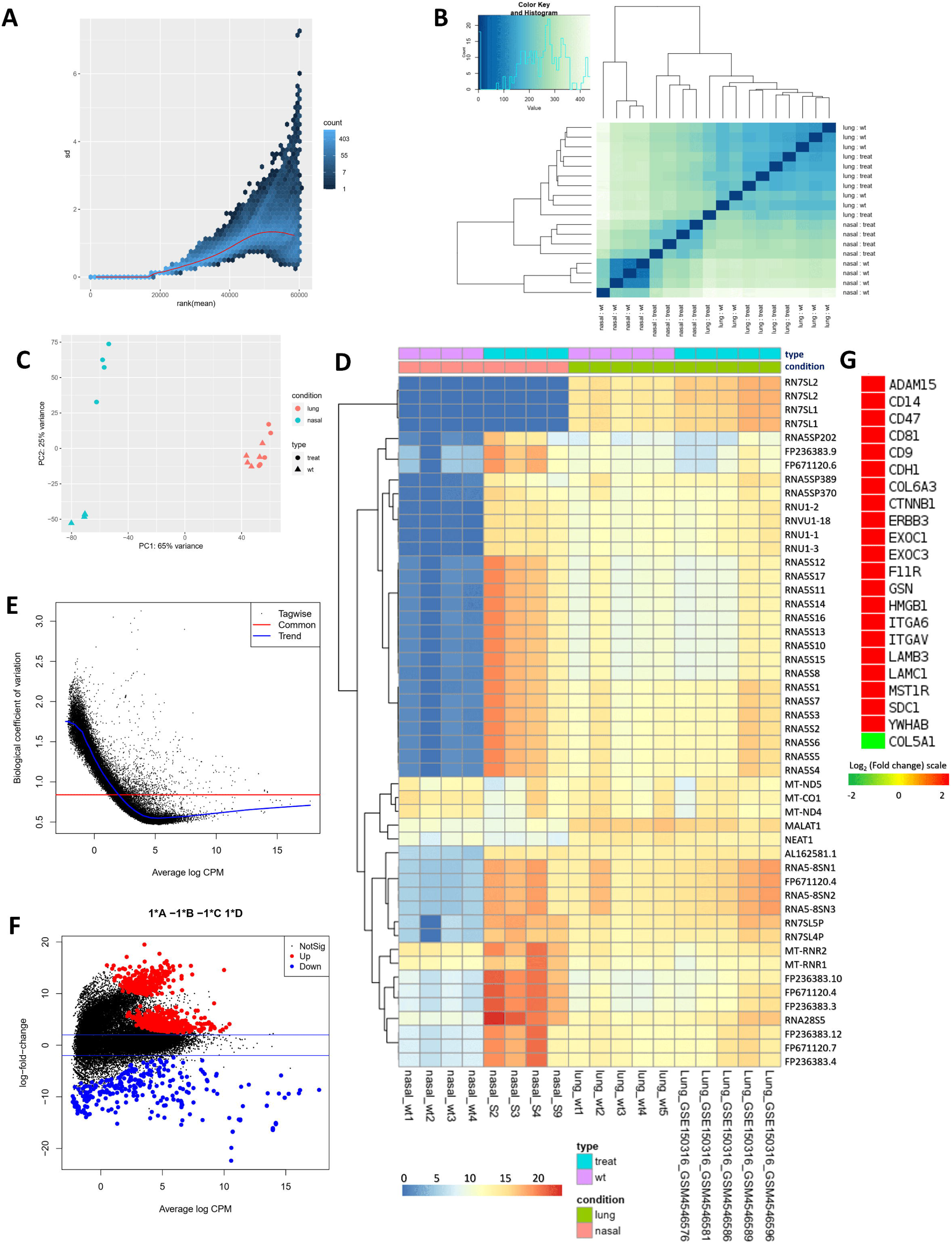
**A.** Variance plot, **B.** Sample to sample distance plot, **C.** Principal component analysis plot, **D.** Clustered heatmap of the count matrix of the normalized RNA-seq reads (top 50 genes) of the SARS-CoV-2 infected nasopharyngeal and lung samples. **E.** Common dispersion plot or the biological coefficient of variation plot. Here we are estimating the dispersion. The square root of the common dispersion gives the coefficient of variation of biological variation. Here the coefficient of biological variation is around 0.8. **F.** MA plot. Plot log-fold change against log-counts per million, with DE genes are highlighted. The blue lines indicate 2-fold changes. Red and blue points indicate genes with P-value less than 0.05. **G.** Expression profiles of genes encoding Integrins. Log_2_ (fold change) values are considered as the expression values of the genes and represented in a color-coded scale; Color towards red indicating higher expression, while color towards green indicating little to no expression.

We then performed functional enrichment analysis to hunt down the signaling pathways which are differentially expressed in lungs compared to the nasopharyngeal cells. These enrichment analyses revealed that biological functions such as viral process, antigen processing, and presentation etc. are highly upregulated, function such as- regulation of gene silencing by miRNA is found downregulated in lungs compared to the nasopharyngeal cells (Figure 8A). Furthermore, pathways that provide antiviral immunity such as apoptosis, phagosome, antigen processing and presentation, adaptive immune system, innate immune system, interferon signaling, different interleukin signaling, cytokine signaling in immune system etc. were highly upregulated in lungs compared to the nasopharyngeal samples (Figure 8B-D). Despite having the antiviral protective roles, hyperactivity from these pathways can significantly worsen the COVID-19 patient’s overall lung functionality which can be further complicated with progressive and permanent lung damage.

**Figure 8:**
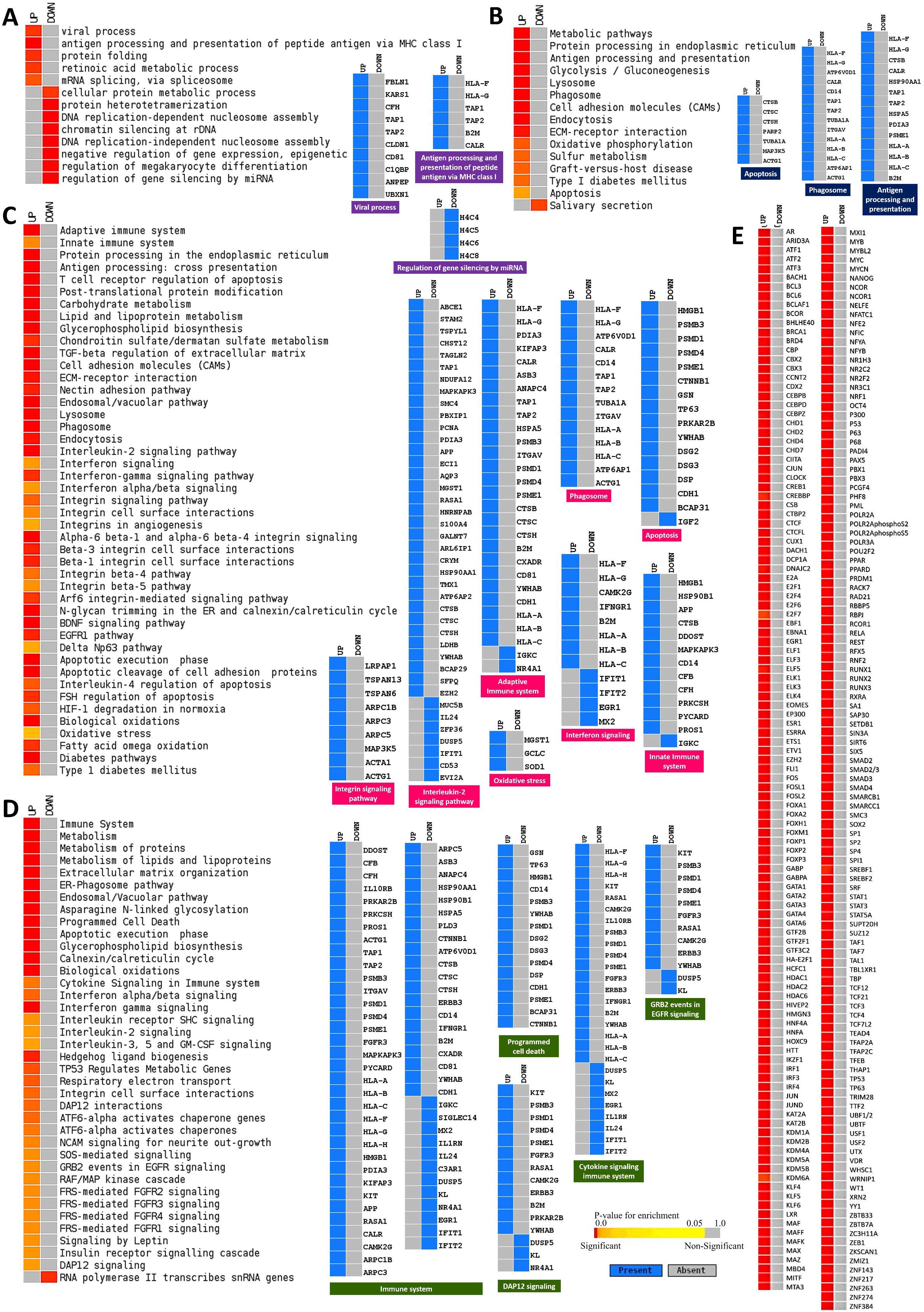
Enrichment analysis and comparison between deregulated genes and the genes of some selected processes in SARS-CoV-2 infected nasopharyngeal samples versus SARS-CoV-2 infected lung biopsy samples using. **A.** GOBP module, **B.** KEGG pathway, **C.** Bioplanet pathway module, **D.** Reactome pathway module. Selected significant terms are represented in heatmaps. Color schemes are similar to Figure 3. For individual processes, blue means presence (significantly differentially expressed gene) while grey means absence (not significantly differentially expressed genes for this module for this experimental condition).

Previously, it was reported that transcription factors can contribute to many inflammatory lung diseases [79, 80] which have similar lung characteristics observed in COVID-19. In this context, we identified the highly expressed transcription factors in lungs by comparing their respective expression values in nasopharyngeal samples (Figure 8E). Among these, transcription factors such as- CBP [79], CEBP [81], NFAT [79], ATF3 [82], GATA6 [83], HDAC2 [84], TCF12 [85] etc. have significant roles in lung’s overall functionality, acute lung injury and antiviral response mechanism in lungs.

### SARS-CoV-2 integrates its proteins in regulating the host antiviral immune responses

As we have observed the differential host responses in COVID-19 nasopharyngeal samples, then we sought to interconnect the virus-host interplay in those host responses. We first analyzed how many of the virus interacting host proteins’ genes reported by Gordon et al. [21] are differentially expressed in our reported nasopharyngeal samples. Only 51 genes of those proteins are found deregulated in our nasopharyngeal samples (Figure 9A). We then constructed a network interlinking the virus-host protein-protein interaction data from Gordon et al. [21] along with the deregulated genes from the nasopharyngeal samples (Figure 9B). Strikingly, we observed that most of the immune signaling related downregulated genes are directly or indirectly connected to the viral proteins (Figure 9B); this suggests the probable roles of the virus in the differential host responses in the COVID-19 affected patients.

**Figure 9:**
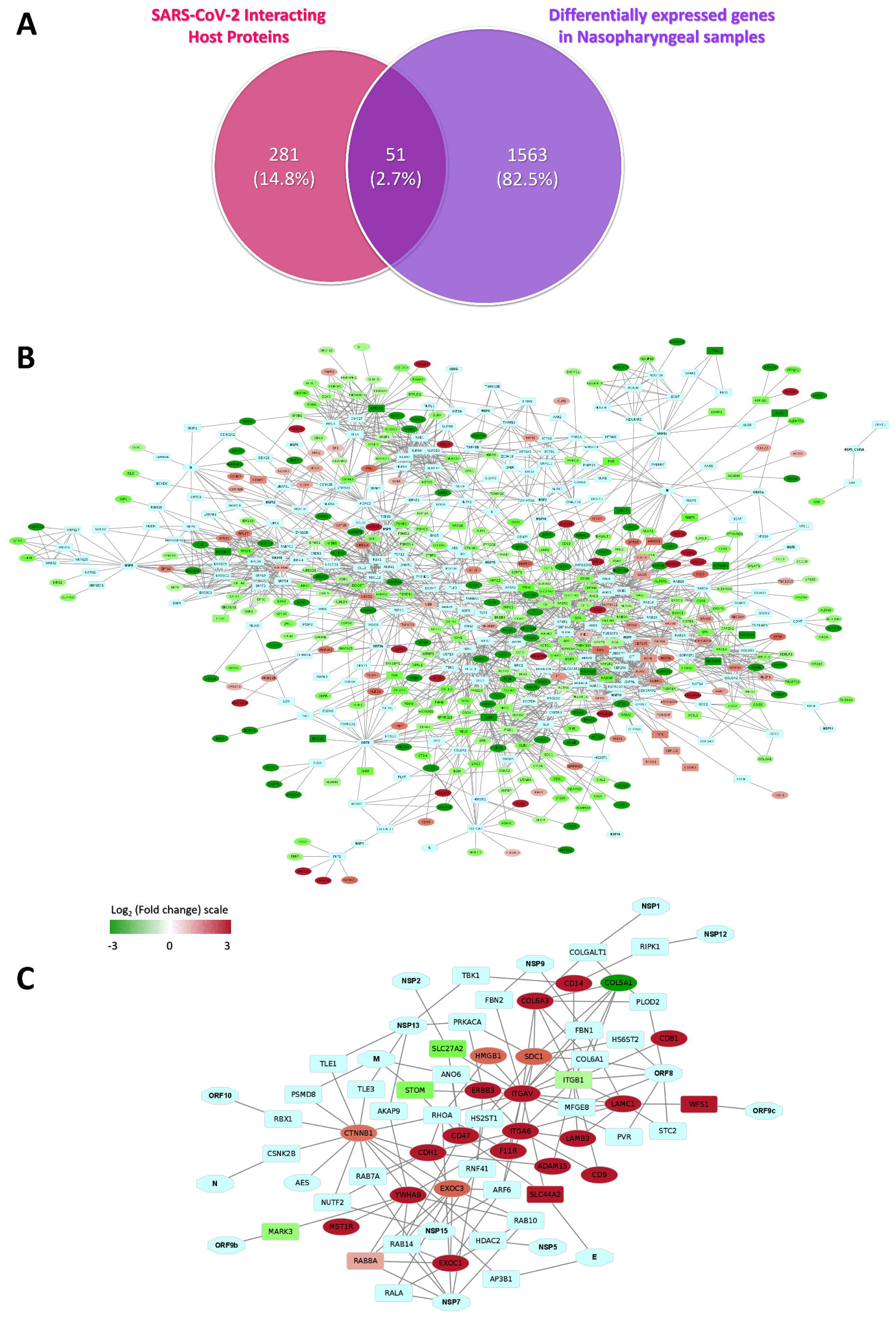
**A.** Venn diagram showing the commonly deregulated genes between deregulated genes in our nasopharyngeal samples and Gordon et al. reported viral protein-interacting high confidence host proteins. Network representing the interactions between genes in **B.** Deregulated genes in Bangladeshi nasopharyngeal samples along with SARS-CoV-2 proteins and Gordon et al. described viral interacting host proteins, and **C.** Differentially expressed Integrin related genes in lungs compared to the nasal samples along with SARS-CoV-2 proteins and Gordon et al. described viral interacting host proteins. Hexagon, ellipse, rounded rectangle represents viral proteins, process-related genes, and proteins that interact with viral proteins, respectively. Expression values of the genes and represented in a color-coded scale. Color towards red indicating higher expression, while color towards green indicating little to no expression.

Furthermore, we sought to establish the links between the viral proteins with integrin signaling associated genes by constructing a functional network with the viral-host protein-protein interaction data with the highly upregulated genes observed in lungs (from the comparison analysis between the lung and nasopharyngeal samples) (Figure 9C). From this constructed network, we observed that viral proteins such as ORF10, N, ORF9b, NSP7, NSP15, NSP5, M, NSP13, NSP2, NSP9, ORF8, ORF9c, NSP12, and NSP1 can directly or indirectly interact with the differentially expressed genes in lungs (Figure 9C), suggesting the putative mechanism behind the deregulated integrin signaling to promote the viral invasion in lungs.

## Discussion

For a better understanding of the host-virus interaction in the SARS-CoV-2 pathogenesis, transcriptional responses of hosts play an enormous role. Moreover, population-specific disease induced host transcriptome might also correlate the susceptibility from COVID-19 with an individual’s ethnicity [86]. In this context, we aimed to discover the representative host transcriptome response upon SARS-CoV-2 infection by performing and analyzing total RNA-seq from the nasopharyngeal samples of four COVID-19 positive Bangladeshi individuals. Also, we analyzed the genomic variations of the SARS-CoV-2 isolates from these patients to gain more insights into their probable origin. Moreover, we compared the transcriptome from different SARS-CoV-2 infection models, particularly, we compared the differential gene expression of the lung biopsy samples with the nasopharyngeal samples of ours to illustrate the possible molecular mechanisms behind the lung damages in severe COVID-19 patients.

Analyzing the variations within the sequenced isolates, we observed a very low frequency of missense amino acid variations within the proteins of SARS-CoV-2, particularly in the spike and ORF3a gene regions. Many concepts are there correlating the probable roles of variations with the COVID-19 disease severity [87, 88]. We did not observe any such variations within the spike region of our reported isolates; however, we recorded an unusual amount of 3’-UTR and 5’-UTR variations within these four isolates. Surprisingly, we spotted a very rare A>G mutation at position 10329 of ORF1ab gene in three of our four isolates; we also got another rare A>T variation at position 25505 of ORF3a gene in one of our isolates. Both of these variations are extremely rare amongst the other worldwide isolates (Table 1).

Phylogenetic analyses from Nextstrain portal suggested that our reported isolates are closely related to the Middle East Asian SARS-CoV-2 isolates; which is also very evident from our constructed neighbor-joining phylogenetic tree of 145 Bangladeshi isolates as our reported isolates are distinctly placed within the tree. This supports the idea of probable source of origin of these isolates from Middle-East Asia, while the rest of the Bangladeshi isolates are more related to the European isolates. As our isolates were obtained from the Chittagong area of Bangladesh, from where a large number of people recently immigrated to Middle-East (particularly Saudi Arabia) for work [89]; those immigrant people returning from the Middle-East during this pandemic might have brought these isolates into Bangladesh.

Previously, host transcriptional responses reported by Blanco-melo et al. [24] and Butler et al. [25] suggested a potential increase in the host antiviral immune responses such as- interferon signaling, interferon stimulated gene signaling, chemokine signaling, cytokine signaling etc.; however, Blanco-melo et al [90] also reported the presence of low IFN-I and IFN-III in COVID-19 patient’s lung cells. We observed similar host immune responses, interferon, and cytokine signaling in Bangladeshi COVID-19 patients too. Moreover, we also observed a stimulated innate immune response in our patients which was also reported for other COVID-19 patients [91].

Astoundingly, important signaling pathways those elicit antiviral immune responses such as- apoptosis [20], phagosome formation [92], antigen processing and presentation [93], Natural killer cell mediated cytotoxicity [94], and Toll-like receptor signaling [95] etc. were found downregulated in Bangladeshi COVID-19 patients. Also, pathways such as- HIF-1 response [96], PI3K-Akt signaling [97], and IL-17 signaling [98] etc. are also found deregulated which could assist the COVID-19 patients suffering from hypoxia, lung injury, and inflammation of the respiratory tract.

While we were comparing the nasopharyngeal cell’s transcriptional responses with other SARS-CoV-2 infection models, we observed that lung cells elicited the immense cytokine and inflammatory responses against the invading viral pathogen. These overstimulated responses sometimes can do irreversible damages to the lungs [99]. This might shed insights into the COVID-19 disease severity when the viral infection progresses into the lungs.

Though an increased amount of ACE2 will facilitate the invasion of SARS-CoV-2, nonetheless, we observed a significant downregulation of ACE2 in lung cells. This phenomenon could backup the concept of ACE2 downregulation by SARS-CoV-2 itself after using it [100], thus reducing the organ protective roles of ACE2 [101] and resulting in progressive lung damages.

Integrins were reported important for the entry of SARS-CoV into the host cells [65], so it was speculated similar phenomenon might also be present in SARS-CoV-2. This idea is further intensified after the study by Sigrist et al. [63], who suggested the presence of an integrin-binding RGD motif in the spike of SARS-CoV-2. Surprisingly, upon the gene expression comparison between the different SARS-CoV-2 infected cells, we observed several folds upregulated expressions of genes related to the integrin signaling pathway in lung cells. This observation could support the idea of increased viral infections in lungs might be happening due to the overexpression of these probable attachment proteins. Also, the network analysis suggests a probable mechanism of upregulation of these proteins by the virus itself by the putative interactions through its proteins. Though more targeted studies should be undertaken for conclusive evidence supporting this phenomenon.

## Conclusion

In this study, we present the very first report of the host transcriptional response data from COVID-19 patients of the South-Asian region along with the SARS-CoV-2 isolates obtained from these patients. This data might provide newer insights into the host susceptibility from the perspective of ethnicity as well as will provide newer aspects of host responses against the virus in the different parts of the respiratory tract. However, a limited number of patient data is used here, but subsequent incorporation of more patient data from other parts of the world will significantly increase the understanding of this complex host-virus response in COVID-19, which will help in designing therapeutic interventions as well as in current clinical management of the patients.

## Acknowledgement

We would like to thank all the authors who have kindly deposited and shared genome data on GISAID (https://www.gisaid.org/). We would also want to convey our gratitude to the developers maintaining the NextStrain novel coronavirus data portal (https://nextstrain.org/).

## Competing Interests

The authors declare no competing interests.

## Author Contributions

ABMMKI designed the workflow and conceived the project. AMAMZS performed sample collection and detection. RA, MSH, SMTK, and MSI performed the RNA sequencing and viral genome assembly. ABMMKI and MAAKK performed the RNA-seq data analysis, comparative genomics, and other bioinformatic analyses. MAAKK and ABMMKI wrote the manuscript. All authors read and approved the final manuscript. ABMMKI and MAAKK contributed equally to this work.

## Funding

This project was not associated with any internal or external source of funding.

## Data Availability

SARS-CoV-2 genomes are deposited at GISAID initiatives portal (https://www.gisaid.org/) with accessions-EPI_ISL_450340, EPI_ISL_450341, EPI_ISL_450342, EPI_ISL_450345. Raw RNA-seq data are deposited at NCBI-GEO (https://www.ncbi.nlm.nih.gov/geo/) under the accession-GSM4667504, GSM4667505, GSM4667506, GSM4667507. Additionally, publicly available data were utilized (Supplementary file 1). Analyses generated data are deposited as supplementary files.

## Ethics statements

Patients’ written-consents were taken before the collection of the samples.

## Supplementary Figure Legends

**Supplementary Figure 1:** Snapshot of Nextstrain data portal showing the phylogenetic relationship of two SARS-CoV-2 isolates used in this study. Isolates of this study are indicated using a red arrow.

**Supplementary Figure 2:** Deregulated genes of selected terms from Figure 3 in different SARS-CoV-2 infection systems. Genes of selected significant terms are represented here. For individual processes, blue means presence (differentially expressed gene of the module term) while grey means absence (not differentially expressed in the experimental condition in that module term). Processes in the green, blue, red color background represent KEGG, Bioplanet, GOBP enriched terms, respectively.

**Supplementary Figure 3:** Deregulated genes of selected terms from Figure 3 in different SARS-CoV-2 infection systems. For individual processes, blue means presence (differentially expressed gene of the module term) while grey means absence (not differentially expressed in the experimental condition in that module term). Processes in the green, blue, red color background represent KEGG, Bioplanet, GOBP enriched terms, respectively.

**Supplementary Figure 4:** Expanded view of the heatmaps A, B, C of Figure 6.

## List of supplementary files

**Supplementary file 1:** Sources of the data used in this study.

**Supplementary file 2:** Per base sequence quality reports of the generated RNA-seq reads of the four COVID-19 infected nasal samples used in this study.

**Supplementary file 3:** Isolate-wise variation information of the four SARS-CoV-2 isolates used in this study.

**Supplementary file 4:** Differentially expressed genes found in the four Nasal samples of Bangladeshi COVID-19 patient.

**Supplementary file 5:** Differentially expressed genes in different SARS-CoV-2 infected cell types.

**Supplementary file 6:** Genes and associated terms used for filtering the expression values used in Figure 6.

**Supplementary file 7:** Differentially expressed genes in SARS-CoV-2 infected Lungs compared to the Bangladeshi Nasal samples used in this study.

## Notes

### Competing Interest Statement

The authors have declared no competing interest.

