## Supplementary file 1 for "Host transcriptional responses and SARS-CoV-2 isolates from the nasopharyngeal samples of Bangladeshi COVID-19 patients"

**Supplementary file 1: Sources of the data used in this study.**

### RNA-seq data generated from Nasopharyngeal samples of Bangladeshi COVID-19 patients

| Sample ID | Specimen | Virus: GISAID Accession | Human RNA-seq: GEO accession | Total Read No. (R1) | Total Read No. (R2) | Mapped Read No. (Human GRCh38) | Sequencing technology | Virus Assembly method | Lineage | Clade |
|---|---|---|---|---|---|---|---|---|---|---|
| **S2** | Nasopharyngeal swab | **EPI_ISL_450340** | GSM4667504 | 10144042 | 10144042 | 8354586 | Illumina NextSeq500 | MEGAHIT v1.1.3 | A | S |
| **S3** | Nasopharyngeal swab | **EPI_ISL_450341** | GSM4667505 | 9276172 | 9276172 | 6542386 | Illumina NextSeq500 | MEGAHIT v1.1.3 | A | S |
| **S4** | Nasopharyngeal swab | **EPI_ISL_450342** | GSM4667506 | 9029322 | 9029322 | 4788596 | Illumina NextSeq500 | MEGAHIT v1.1.3 | A | S |
| **S9** | Nasopharyngeal swab | **EPI_ISL_450345** | GSM4667507 | 8234222 | 8234222 | 4170828 | Illumina NextSeq500 | MEGAHIT v1.1.3 | A | S |
| | | | GSE ID | | | | | | | |
| | | | **GSE154244** | | | | | | | |

### Publicly available data used in this study

| Nasal Control (WT) Healthy Data | | | | | | | |
|---|---|---|---|---|---|---|---|
| **GEO Accession** | **SAMPLE-1 (WT)** | **SAMPLE-2 (WT)** | **SAMPLE-3 (WT)** | **SAMPLE-4 (WT)** | | | |
| **GSE97668** | GSM2574998 | GSM2574999 | GSM2575000 | GSM2575001 | disease status: Non-asthmatic | age: Adult. No Treatment | cell type: Nasal epithelial |
| Citation: Heymann PW, Nguyen HT, Steinke JW, Turner RB et al. Rhinovirus infection results in stronger and more persistent genomic dysregulation: Evidence for altered innate immune response in asthmatics at baseline, early in infection, and during convalescence. PLoS One 2017;12(5):e0178096. PMID: 28552993 | | | | | | | |

**Infected Cell Data**

| GEO Accession | | | GEO: GSM (Sample ID) | |
|---|---|---|---|---|
| **GSE147507** | ung_GSE147507_GSM44624 | Lung | GSM4462415 | Lung-Postmortem |
| | Lung_GSE147507_GSM44624 | Lung | GSM4462416 | Lung-Postmortem |
| **GSE150316** | ung_GSE150316_GSM45465 | Lung | GSM4546576 | Lung-Autopsy |
| | Lung_GSE150316_GSM45465 | Lung | GSM4546581 | Lung-Autopsy |
| | Lung_GSE150316_GSM45465 | Lung | GSM4546586 | Lung-Autopsy |
| | Lung_GSE150316_GSM45465 | Lung | GSM4546589 | Lung-Autopsy |
| | Lung_GSE150316_GSM45465 | Lung | GSM4546596 | Lung-Autopsy |
| **GSE148729** | LU3_GSE148729_GSM4477 | Calu3 | GSM4477962 | Lung-epithelial-cell |
| | CALU3_GSE148729_GSM4477 | Calu3 | GSM4477963 | Lung-epithelial-cell |
| **GSE147507** | IBE_GSE147507_GSM4432 | NHBE | GSM4432381 | Bronchial-Epithelial |
| | NHBE_GSE147507_GSM4432 | NHBE | GSM4432382 | Bronchial-Epithelial |
| | NHBE_GSE147507_GSM4432 | NHBE | GSM4432383 | Bronchial-Epithelial |
