## Supplementary file 2 for "Host transcriptional responses and SARS-CoV-2 isolates from the nasopharyngeal samples of Bangladeshi COVID-19 patients"

**Supplementary file 2: Per base sequence quality reports of the generated RNA-seq reads of the four COVID-19 infected nasal samples used in this study.**

**A. Sample S2**

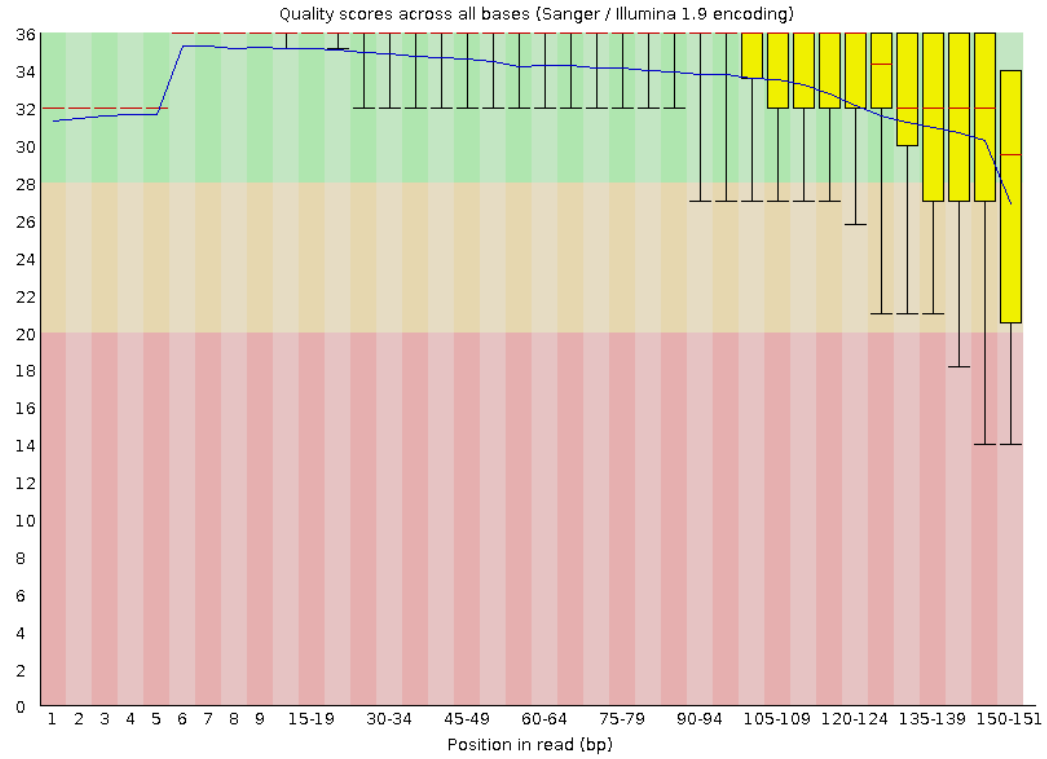


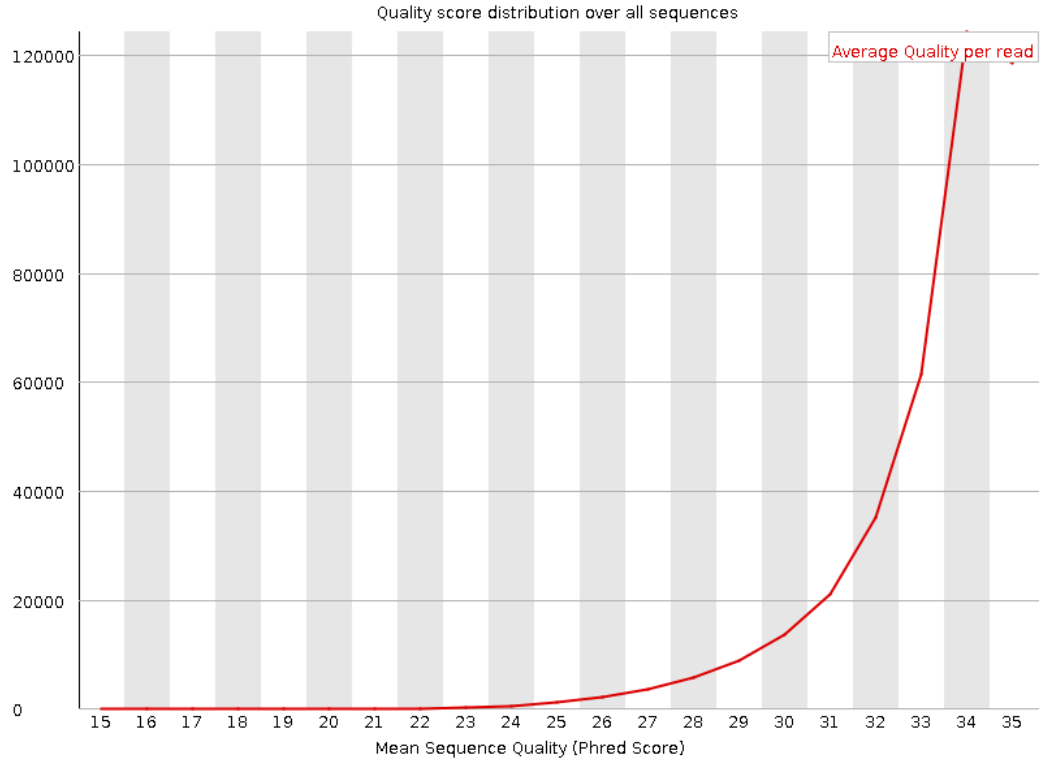

### B. Sample S3

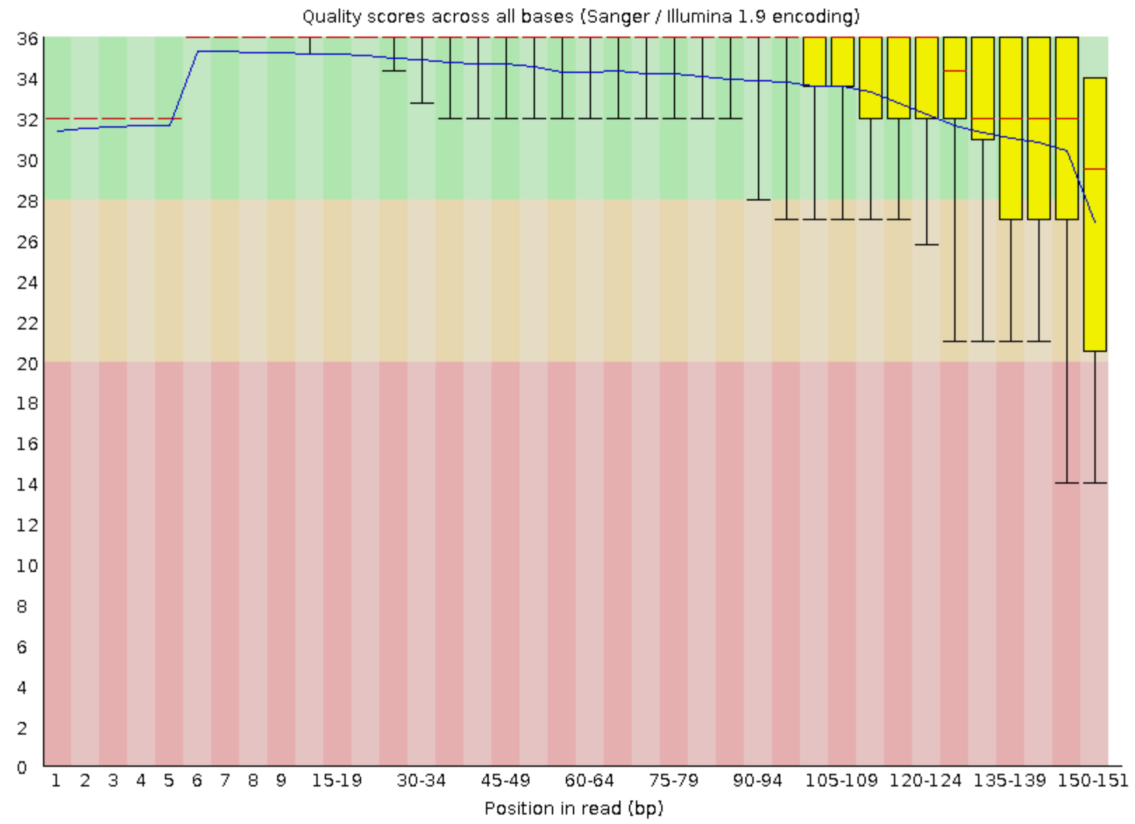

Quality scores across all bases (Sanger / Illumina 1.9 encoding)
36
34
32
30
28
26
24
22
20
18
16
14
12
10
8
6
4
2
0
1 2 3 4 5 6 7 8 9 15-19 30-34 45-49 60-64 75-79 90-94 105-109 120-124 135-139 150-151
Position in read (bp)


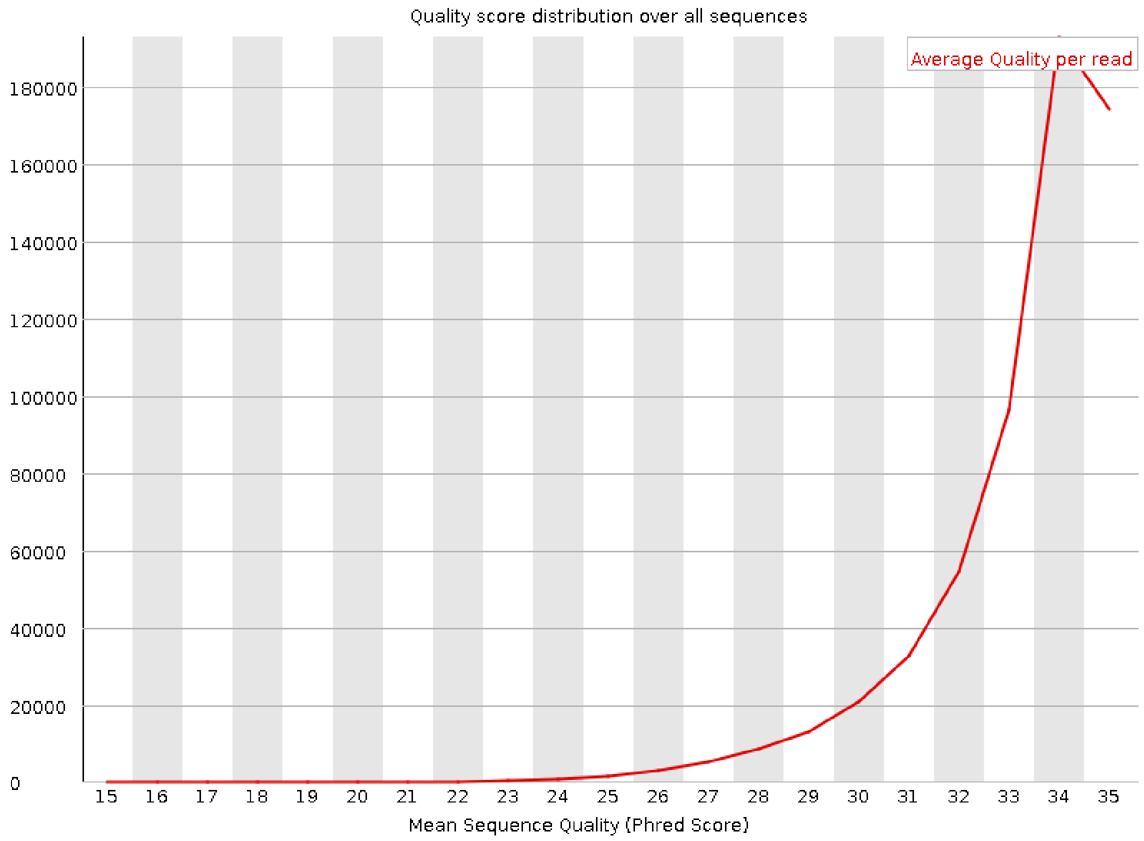

Quality score distribution over all sequences
Average Quality per read
180000
160000
140000
120000
100000
80000
60000
40000
20000
0
15 16 17 18 19 20 21 22 23 24 25 26 27 28 29 30 31 32 33 34 35
Mean Sequence Quality (Phred Score)

### C. Sample S4

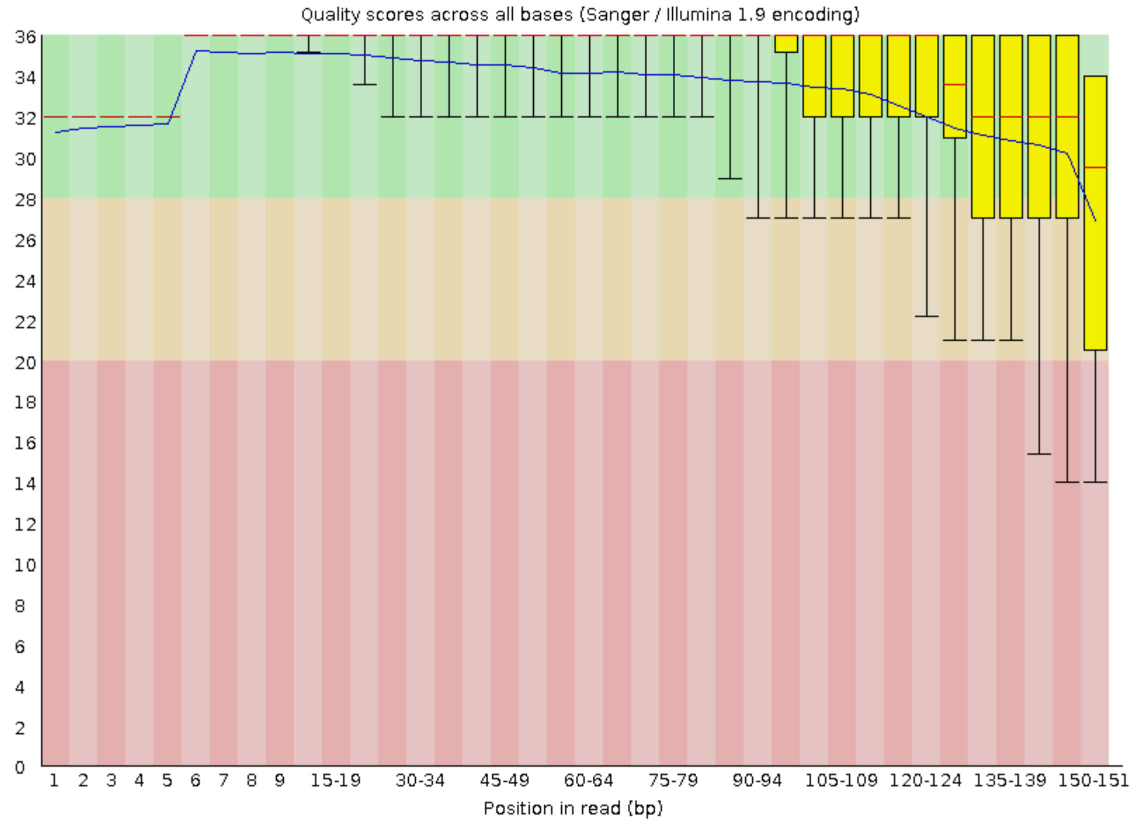

Quality scores across all bases (Sanger / Illumina 1.9 encoding)
36
34
32
30
28
26
24
22
20
18
16
14
12
10
8
6
4
2
0
1 2 3 4 5 6 7 8 9 15-19 30-34 45-49 60-64 75-79 90-94 105-109 120-124 135-139 150-151
Position in read (bp)


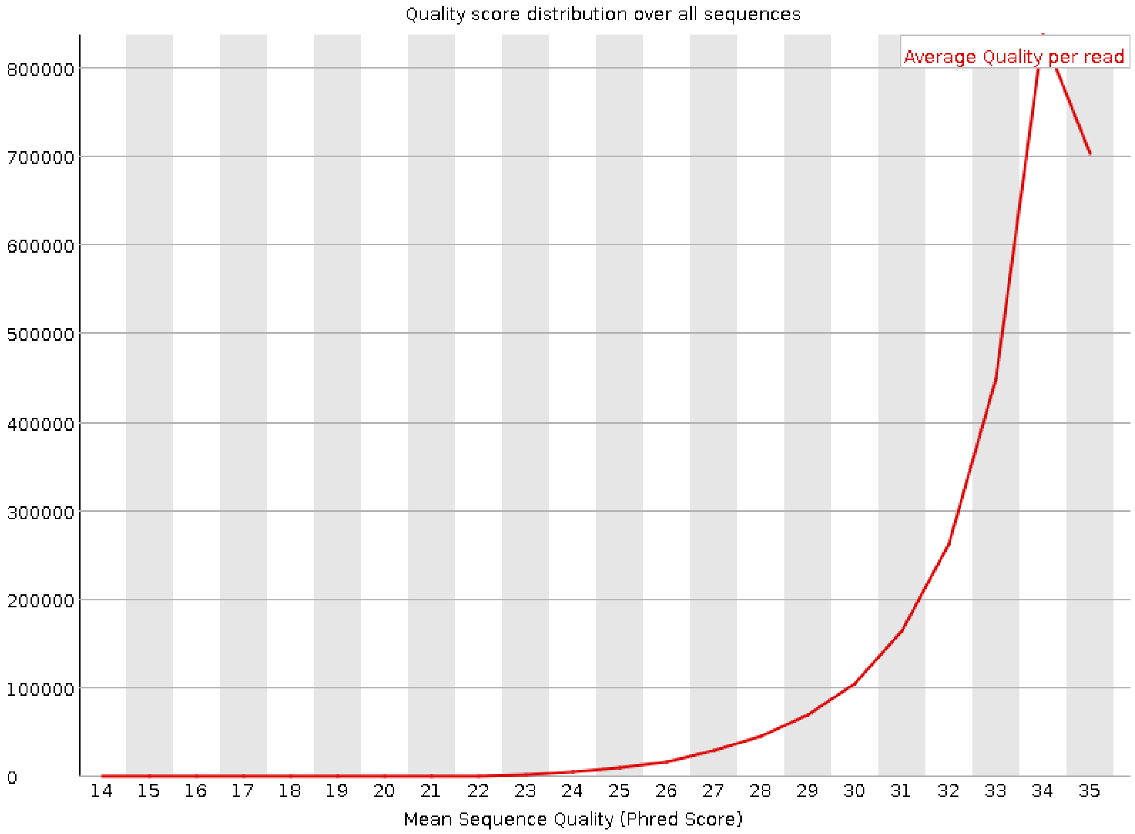

Quality score distribution over all sequences
Average Quality per read
800000
700000
600000
500000
400000
300000
200000
100000
0
14 15 16 17 18 19 20 21 22 23 24 25 26 27 28 29 30 31 32 33 34 35
Mean Sequence Quality (Phred Score)

### D. Sample S9

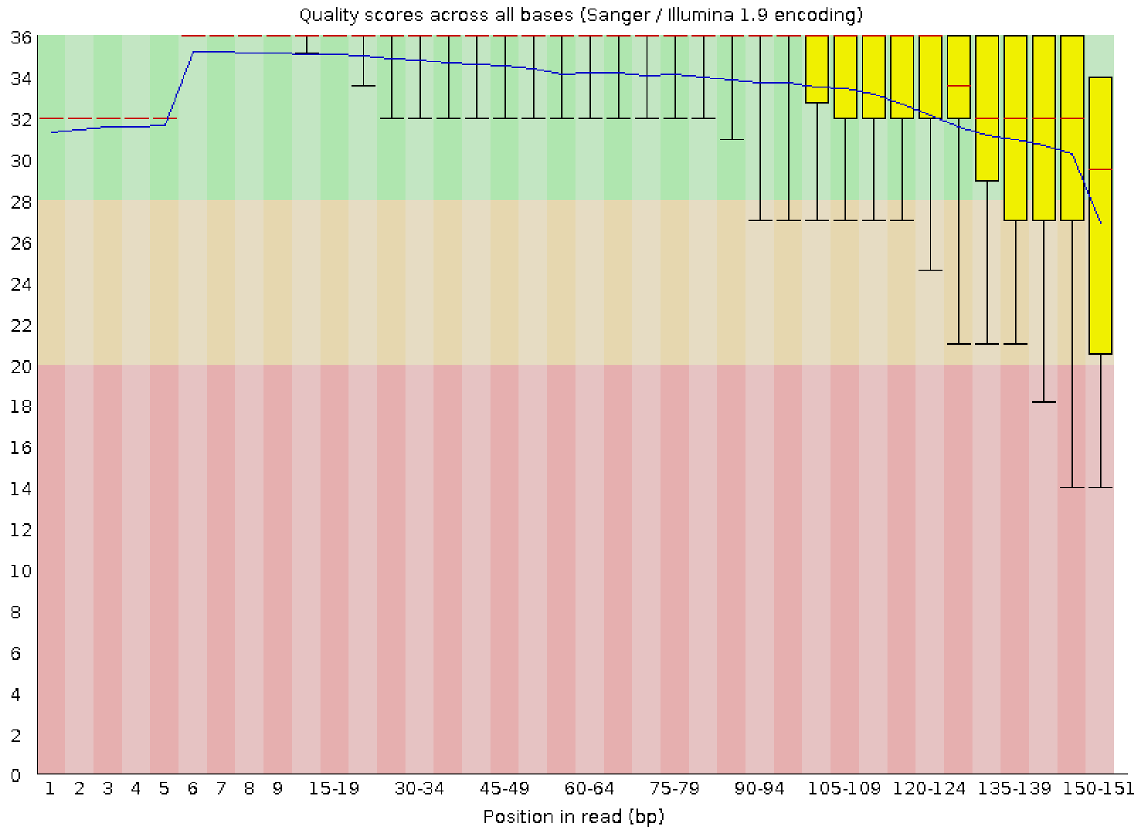

Quality scores across all bases (Sanger / Illumina 1.9 encoding)
36
34
32
30
28
26
24
22
20
18
16
14
12
10
8
6
4
2
0
1 2 3 4 5 6 7 8 9 15-19 30-34 45-49 60-64 75-79 90-94 105-109 120-124 135-139 150-151
Position in read (bp)


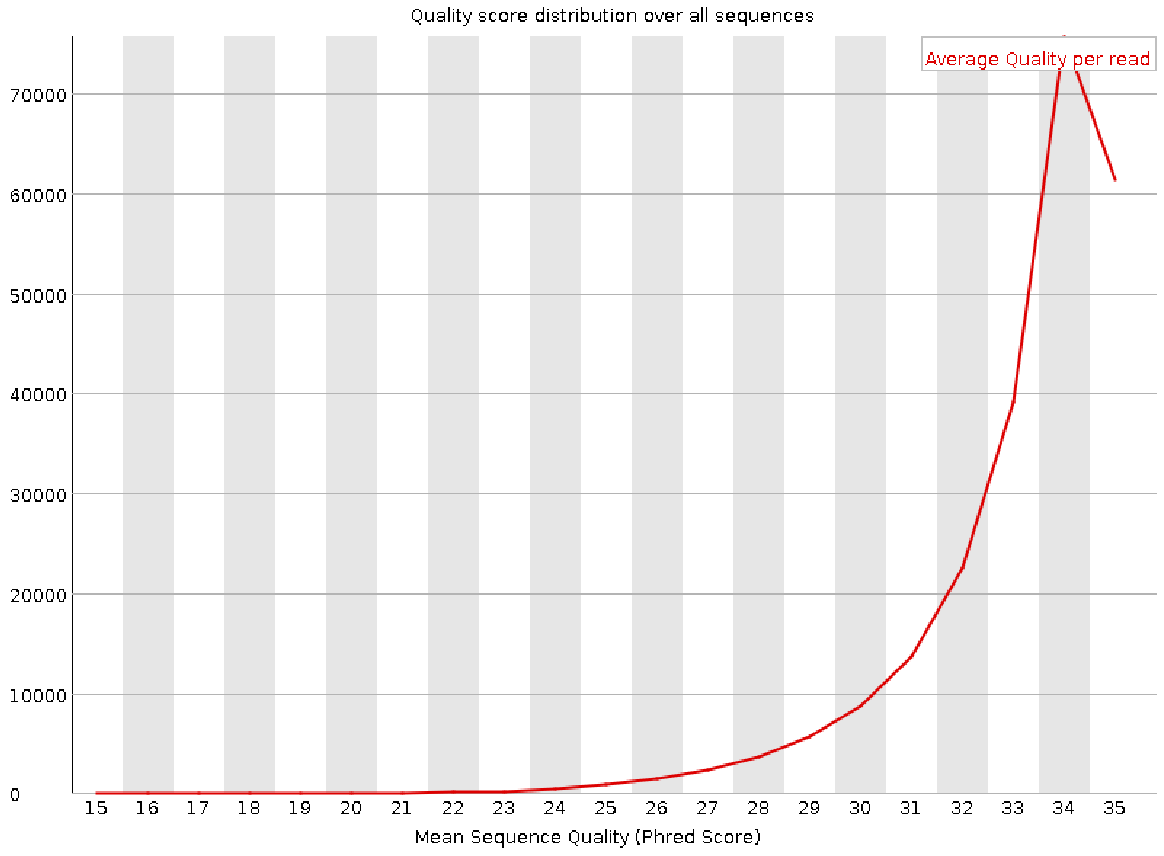

Quality score distribution over all sequences
Average Quality per read
70000
60000
50000
40000
30000
20000
10000
0
15 16 17 18 19 20 21 22 23 24 25 26 27 28 29 30 31 32 33 34 35
Mean Sequence Quality (Phred Score)
