## Supplementary file 3 for "Host transcriptional responses and SARS-CoV-2 isolates from the nasopharyngeal samples of Bangladeshi COVID-19 patients"

**Supplementary file 3: Isolate-wise variation information of the four SARS-CoV-2 isolates used in this study.**

| Isolate S2 | | | | | | | | |
|---|---|---|---|---|---|---|---|---|
| Start position | End position | Base | Ref base | Ref annotation position | Annotation Type | Protein.Position.Amino acids change | Gene.Position.Codons | Impact |
| 29742 | 29742 | A | REF:G | START:3'UTR | downstream gene variant | QHI42199.1 | gene-ORF10 | MODIFIER; DISTANCE=68 |
| 7 | 7 | C | REF:G | START:5'UTR | intergenic variant | - | - | MODIFIER |
| 12 | 12 | T | REF:A | START:5'UTR | intergenic variant | - | - | MODIFIER |
| 29893 | 29893 | G | REF:A | START:3'UTR | intergenic variant | - | - | MODIFIER |
| 29895 | 29895 | T | REF:A | START:3'UTR | intergenic variant | - | - | MODIFIER |
| 29896 | 29896 | C | REF:A | START:3'UTR | intergenic variant | - | - | MODIFIER |
| 29897 | 29897 | G | REF:A | START:3'UTR | intergenic variant | - | - | MODIFIER |
| 29898 | 29898 | G | REF:A | START:3'UTR | intergenic variant | - | - | MODIFIER |
| 29901 | 29901 | G | REF:A | START:3'UTR | intergenic variant | - | - | MODIFIER |
| 2 | 2 | TA | REF:T | START:5'UTR | intergenic variant | - | - | MODIFIER |
| 9 | 9 | TTTTCGC | REF:T | START:5'UTR | intergenic variant | - | - | MODIFIER |
| 29864 | 29891 | G | REF:GAATGACAAAAAAAAAAAAAAAAAAAAAA | START:3'UTR | intergenic variant | - | - | MODIFIER |
| 29903 | 29903 | GCCGTCGT | REF:A | START:3'UTR | intergenic variant | - | - | MODIFIER |
| 10329 | 10329 | G | REF:A | START:gene-orf1ab | missense variant | QHD43415.1:p.3355D>G | gene-orf1ab:c.10064gAt>gGt | MODERATE |
| 28144 | 28144 | C | REF:T | START:gene-ORF8 | missense variant | QHD43422.1:p.84L>S | gene-ORF8:c.251tTa>tCa | MODERATE |
| 28878 | 28878 | A | REF:G | START:gene-N | missense variant | QHD43423.2:p.202S>N | gene-N:c.605aGt>aAt | MODERATE |
| 29392 | 29392 | T | REF:G | START:gene-N | missense variant | QHD43423.2:p.373K>N | gene-N:c.1119aaG>aaT | MODERATE |

| | | | | | | | | |
|---|---|---|---|---|---|---|---|---|
| 601 | 601 | T | REF:C | START:gene-orf1ab | synonymous variant | QHD43415.1:p.112G | gene-orf1ab:c.336ggC>ggT | LOW |
| 8782 | 8782 | T | REF:C | START:gene-orf1ab | synonymous variant | QHD43415.1:p.2839S | gene-orf1ab:c.8517agC>agT | LOW |
| 15324 | 15324 | T | REF:C | START:gene-orf1ab | synonymous variant | QHD43415.1:p.5020N | gene-orf1ab:c.15060aaC>aaT | LOW |
| 22468 | 22468 | T | REF:G | START:gene-S | synonymous variant | QHD43416.1:p.302T | gene-S:c.906acG>acT | LOW |

#### Isolate S3

| Start position | End position | Base | Ref base | Ref annotation position | Annotation Type | Protein.Position. Amino acids change | Gene.Position. Codons | Impact |
|---|---|---|---|---|---|---|---|---|
| 29742 | 29742 | A | REF:G | START:3'UTR | downstream gene variant | QHI42199.1 | gene-ORF10 | MODIFIER; DISTANCE=68 |
| 1 | 12 | - | REF:ATTAAAGGTTTA | START:5'UTR | intergenic variant | - | - | MODIFIER |
| 29870 | 29903 | C | REF:CAAAAAAAAAAAAAAAAAAAAAAAAAAAAAAAAA | START:3'UTR | intergenic variant | - | - | MODIFIER |
| 10329 | 10329 | G | REF:A | START:gene-orf1ab | missense variant | QHD43415.1:p.3355D>G | gene-orf1ab:c.10064gAt>gGt | MODERATE |
| 28144 | 28144 | C | REF:T | START:gene-ORF8 | missense variant | QHD43422.1:p.84L>S | gene-ORF8:c.251tTa>tCa | MODERATE |
| 28878 | 28878 | A | REF:G | START:gene-N | missense variant | QHD43423.2:p.202S>N | gene-N:c.605aGt>aAt | MODERATE |
| 29392 | 29392 | T | REF:G | START:gene-N | missense variant | QHD43423.2:p.373K>N | gene-N:c.1119aaG>aaT | MODERATE |
| 601 | 601 | T | REF:C | START:gene-orf1ab | synonymous variant | QHD43415.1:p.112G | gene-orf1ab:c.336ggC>ggT | LOW |
| 8782 | 8782 | T | REF:C | START:gene-orf1ab | synonymous variant | QHD43415.1:p.2839S | gene-orf1ab:c.8517agC>agT | LOW |

| | | | | | | | | |
|---|---|---|---|---|---|---|---|---|
| 15324 | 15324 | T | REF:C | START:gene-orf1ab | synonymous variant | QHD43415.1:p.5020N | gene-orf1ab:c.15060aaC>aaT | LOW |
| 22468 | 22468 | T | REF:G | START:gene-S | synonymous variant | QHD43416.1:p.302T | gene-S:c.906acG>acT | LOW |

#### Isolate S4

| Start position | End position | Base | Ref base | Ref annotation position | Annotation Type | Protein.Position.Amino acids change | Gene.Position.Codons | Impact |
|---|---|---|---|---|---|---|---|---|
| 29742 | 29742 | A | REF:G | START:3'UTR | downstream gene variant | QHI42199.1 | gene-ORF10 | MODIFIER; DISTANCE=68 |
| 4 | 4 | T | REF:A | START:5'UTR | intergenic variant | - | - | MODIFIER |
| 13 | 13 | C | REF:T | START:5'UTR | intergenic variant | - | - | MODIFIER |
| 29870 | 29870 | G | REF:C | START:3'UTR | intergenic variant | - | - | MODIFIER |
| 29872 | 29872 | T | REF:A | START:3'UTR | intergenic variant | - | - | MODIFIER |
| 29873 | 29873 | C | REF:A | START:3'UTR | intergenic variant | - | - | MODIFIER |
| 29874 | 29874 | G | REF:A | START:3'UTR | intergenic variant | - | - | MODIFIER |
| 29875 | 29875 | G | REF:A | START:3'UTR | intergenic variant | - | - | MODIFIER |
| 29878 | 29878 | T | REF:A | START:3'UTR | intergenic variant | - | - | MODIFIER |
| 29880 | 29880 | G | REF:A | START:3'UTR | intergenic variant | - | - | MODIFIER |
| 29882 | 29882 | G | REF:A | START:3'UTR | intergenic variant | - | - | MODIFIER |
| 29883 | 29883 | T | REF:A | START:3'UTR | intergenic variant | - | - | MODIFIER |
| 29884 | 29884 | C | REF:A | START:3'UTR | intergenic variant | - | - | MODIFIER |
| 29885 | 29885 | G | REF:A | START:3'UTR | intergenic variant | - | - | MODIFIER |
| 29886 | 29886 | T | REF:A | START:3'UTR | intergenic variant | - | - | MODIFIER |
| 29887 | 29887 | G | REF:A | START:3'UTR | intergenic variant | - | - | MODIFIER |
| 29888 | 29888 | T | REF:A | START:3'UTR | intergenic variant | - | - | MODIFIER |
| 29890 | 29890 | G | REF:A | START:3'UTR | intergenic variant | - | - | MODIFIER |
| 29891 | 29891 | G | REF:A | START:3'UTR | intergenic variant | - | - | MODIFIER |

| | | | | | | | | |
|---|---|---|---|---|---|---|---|---|
| 29892 | 29892 | G | REF:A | START:3'UTR | intergenic variant | - | - | MODIFIER |
| 29896 | 29896 | G | REF:A | START:3'UTR | intergenic variant | - | - | MODIFIER |
| 29898 | 29898 | G | REF:A | START:3'UTR | intergenic variant | - | - | MODIFIER |
| 29900 | 29900 | G | REF:A | START:3'UTR | intergenic variant | - | - | MODIFIER |
| 2 | 2 | TTTCAAAGATCAAGTCA | REF:T | START:5'UTR | intergenic variant | - | - | MODIFIER |
| 29901 | 29903 | A | REF:AAA | START:3'UTR | intergenic variant | - | - | MODIFIER |
| 10329 | 10329 | G | REF:A | START:gene-orf1ab | missense variant | QHD43415.1:p.3355D>G | gene-orf1ab:c.10064gAt>gGt | MODERATE |
| 12119 | 12119 | T | REF:C | START:gene-orf1ab | missense variant | QHD43415.1:p.3952P>S | gene-orf1ab:c.11854Cca>Tca | MODERATE |
| 19414 | 19414 | A | REF:G | START:gene-orf1ab | missense variant | QHD43415.1:p.6384V>I | gene-orf1ab:c.19150Gta>Ata | MODERATE |
| 28144 | 28144 | C | REF:T | START:gene-ORF8 | missense variant | QHD43422.1:p.84L>S | gene-ORF8:c.251tTa>tCa | MODERATE |
| 28878 | 28878 | A | REF:G | START:gene-N | missense variant | QHD43423.2:p.202S>N | gene-N:c.605aGt>aAt | MODERATE |
| 280 | 280 | T | REF:C | START:gene-orf1ab | synonymous variant | QHD43415.1:p.5V | gene-orf1ab:c.15gtC>gtT | LOW |
| 8782 | 8782 | T | REF:C | START:gene-orf1ab | synonymous variant | QHD43415.1:p.2839S | gene-orf1ab:c.8517agC>agT | LOW |
| 10870 | 10870 | T | REF:G | START:gene-orf1ab | synonymous variant | QHD43415.1:p.3535L | gene-orf1ab:c.10605ctG>ctT | LOW |
| 15324 | 15324 | T | REF:C | START:gene-orf1ab | synonymous variant | QHD43415.1:p.5020N | gene-orf1ab:c.15060aaC>aaT | LOW |
| 22468 | 22468 | T | REF:G | START:gene-S | synonymous variant | QHD43416.1:p.302T | gene-S:c.906acG>acT | LOW |

### Isolate S9

| Start position | End position | Base | Ref base | Ref annotation position | Annotation Type | Protein.Position.Amino acids change | Gene.Position.Codons | Impact |
|---|---|---|---|---|---|---|---|---|
| 29742 | 29742 | A | REF:G | START:3'UTR | downstream gene variant | QHI42199.1 | gene-ORF10 | MODIFIER; DISTANCE=68 |
| 29856 | 29856 | A | REF:T | START:3'UTR | intergenic variant | - | - | MODIFIER |
| 29858 | 29858 | A | REF:T | START:3'UTR | intergenic variant | - | - | MODIFIER |
| 29895 | 29895 | T | REF:A | START:3'UTR | intergenic variant | - | - | MODIFIER |
| 29897 | 29897 | G | REF:A | START:3'UTR | intergenic variant | - | - | MODIFIER |
| 29898 | 29898 | T | REF:A | START:3'UTR | intergenic variant | - | - | MODIFIER |
| 29901 | 29901 | G | REF:A | START:3'UTR | intergenic variant | - | - | MODIFIER |
| 1 | 14 | - | REF:ATTAAAGGTTTATA | START:5'UTR | intergenic variant | - | - | MODIFIER |
| 29864 | 29893 | T | REF:GAATGACAAAAAAAAAAAAAAAAAAAAAAA | START:3'UTR | intergenic variant | - | - | MODIFIER |
| 29903 | 29903 | GCGTCGTGT | REF:A | START:3'UTR | intergenic variant | - | - | MODIFIER |
| 10323 | 10323 | G | REF:A | START:gene-orf1ab | missense variant | QHD43415.1:p.3353K>R | gene-orf1ab:c.10058aAg>aGg | MODERATE |
| 25505 | 25505 | T | REF:A | START:gene-ORF3a | missense variant | QHD43417.1:p.38Q>L | gene-ORF3a:c.113cAa>cTa | MODERATE |
| 28087 | 28087 | T | REF:C | START:gene-ORF8 | missense variant | QHD43422.1:p.65A>V | gene-ORF8:c.194gCt>gTt | MODERATE |
| 28144 | 28144 | C | REF:T | START:gene-ORF8 | missense variant | QHD43422.1:p.84L>S | gene-ORF8:c.251tTa>tCa | MODERATE |
| 28878 | 28878 | A | REF:G | START:gene-N | missense variant | QHD43423.2:p.202S>N | gene-N:c.605aGt>aAt | MODERATE |
| 8782 | 8782 | T | REF:C | START:gene-orf1ab | synonymous variant | QHD43415.1:p.2839S | gene-orf1ab:c.8517agC>agT | LOW |
| 22468 | 22468 | T | REF:G | START:gene-S | synonymous variant | QHD43416.1:p.302T | gene-S:c.906acG>acT | LOW |

| | | | | | | | | |
|---|---|---|---|---|---|---|---|---|
| 23320 | 23320 | T | REF:C | START:gene-S | synonymous variant | QHD43416.1:p.586D | gene-S:c.1758gaC>gaT | LOW |
