## Supplementary file 4 for "Host transcriptional responses and SARS-CoV-2 isolates from the nasopharyngeal samples of Bangladeshi COVID-19 patients"

**Supplementary file 4: Differentially expressed genes found in the Nasal samples of Bangladeshi COVID-19 patient.**

| Up-regulated | Down-regulated | Differentially expressed |
|---|---|---|
| RNA5S14 | LINC00963 | RNA5S14 |
| RNA5S11 | ADGRG1 | RNA5S11 |
| RNA5S8 | SRSF5 | RNA5S8 |
| RNA5S1 | ACTN4 | RNA5S1 |
| RNA5S10 | EWSR1 | RNA5S10 |
| RNA5S13 | HK1 | RNA5S13 |
| RNA5S15 | TMED2 | RNA5S15 |
| RNA5S2 | CCZ1B | RNA5S2 |
| RNA5S16 | PSMD2 | RNA5S16 |
| RNA5S3 | VAMP3 | RNA5S3 |
| RNA5S6 | CTNND1 | RNA5S6 |
| RNA5S17 | ARRDC3 | RNA5S17 |
| RNA5S7 | RDH10 | RNA5S7 |
| RNA5S12 | EIF3CL | RNA5S12 |
| RNA5S5 | PRKAR1A | RNA5S5 |
| RNA5S4 | PPP2CB | RNA5S4 |
| RNU1-28P | TSPYL1 | RNU1-28P |
| RNVU1-18 | SRP9 | RNVU1-18 |
| RNU1-4 | SMARCA2 | RNU1-4 |
| RNVU1-29 | GLUL | RNVU1-29 |
| RNU1-2 | RBM25 | RNU1-2 |
| RNU1-1 | TNFRSF21 | RNU1-1 |
| RNU1-27P | SELENBP1 | RNU1-27P |
| RNU1-3 | DHCR24 | RNU1-3 |
| RNVU1-7 | MAT2A | RNVU1-7 |
| RNY1 | EIF3C | RNY1 |
| RNU4-2 | SCARB2 | RNU4-2 |
| RNA5S9 | TMPRSS4 | RNA5S9 |
| FP671120.2 | EIF4G1 | FP671120.2 |
| CR392039.1 | ENC1 | CR392039.1 |
| VTRNA1-1 | HNRNPH1 | VTRNA1-1 |
| RNA5SP370 | WDR1 | RNA5SP370 |
| FP236383.6 | ADSS2 | FP236383.6 |
| RNA5SP202 | TMEM123 | RNA5SP202 |
| RNVU1-28 | IL33 | RNVU1-28 |
| CDR1 | CCZ1 | CDR1 |
| SNORA73B | FBXL5 | SNORA73B |
| RNU1-11P | ZBTB7A | RNU1-11P |
| RNY4 | HNRNPK | RNY4 |
| FP236383.1 | PLEKHB2 | FP236383.1 |
| RNA5SP389 | HNRNPU | RNA5SP389 |
| RNU4-1 | ARL8B | RNU4-1 |
| FP236383.9 | DHX15 | FP236383.9 |
| H4C5 | UBQLN1 | H4C5 |
| MIR3648-1 | ANXA7 | MIR3648-1 |
| SMR3B | NDRG2 | SMR3B |
| H4C11 | TOMM20 | H4C11 |
| RNA5-8SN2 | GANAB | RNA5-8SN2 |
| RNA5-8SN3 | CTBP2 | RNA5-8SN3 |
| FP671120.4 | CBWD6 | FP671120.4 |
| RNA5-8SN1 | ITGB1 | RNA5-8SN1 |
| FP236383.3 | ASPH | FP236383.3 |
| MIR3648-2 | ATP8B1 | MIR3648-2 |
| FP671120.1 | PFN2 | FP671120.1 |
| FP236383.10 | DDB1 | FP236383.10 |
| RN7SL5P | ARGLU1 | RN7SL5P |
| RNA28S5 | SF3B1 | RNA28S5 |
| PRB4 | ANPEP | PRB4 |

| | | |
|---|---|---|
| PRB3 | DDX1 | PRB3 |
| RNA5SP298 | NCSTN | RNA5SP298 |
| FP236383.5 | DNAJC10 | FP236383.5 |
| FP236383.4 | DDX5 | FP236383.4 |
| FP236383.12 | IFT57 | FP236383.12 |
| FP671120.7 | PGK1 | FP671120.7 |
| RN7SL4P | MIA3 | RN7SL4P |
| RNA5SP226 | SDC4 | RNA5SP226 |
| RN7SL752P | VPS35 | RN7SL752P |
| RNY3 | GALNT7 | RNY3 |
| AC079601.1 | NAP1L4 | AC079601.1 |
| RNA5SP145 | SRSF10 | RNA5SP145 |
| RNA5SP149 | COPB1 | RNA5SP149 |
| RNA5SP429 | DDX17 | RNA5SP429 |
| AL135938.1 | RBM39 | AL135938.1 |
| RNA5SP74 | TAX1BP1 | RNA5SP74 |
| CRNN | ANKRD12 | CRNN |
| RNY3P1 | LMBRD1 | RNY3P1 |
| FAM27E2 | CALM2 | FAM27E2 |
| FP236383.7 | SLC37A3 | FP236383.7 |
| RN7SL396P | RDX | RN7SL396P |
| FP671120.5 | ABCD3 | FP671120.5 |
| H1-4 | CKAP4 | H1-4 |
| AL162581.1 | TMEM41B | AL162581.1 |
| CRCT1 | NFE2L1 | CRCT1 |
| RNA5SP335 | EPRS1 | RNA5SP335 |
| RNA5-8SP6 | UBE2Q1 | RNA5-8SP6 |
| AC024051.11 | ERBB3 | AC024051.11 |
| AC024051.5 | DIMT1 | AC024051.5 |
| AC024051.12 | RAC1 | AC024051.12 |
| AC024051.1 | CALM1 | AC024051.1 |
| FP236383.8 | PRSS23 | FP236383.8 |
| RNU2-1 | SFPQ | RNU2-1 |
| AC024051.2 | EXOC3 | AC024051.2 |
| SNORD17 | ADAM9 | SNORD17 |
| AC024051.7 | XPO1 | AC024051.7 |
| AC024051.10 | ACSL5 | AC024051.10 |
| AC024051.3 | CCT2 | AC024051.3 |
| AC024051.8 | KDELR1 | AC024051.8 |
| AC024051.6 | PCMTD2 | AC024051.6 |
| RNA5SP161 | ACTR2 | RNA5SP161 |
| AC024051.4 | ALDH3A2 | AC024051.4 |
| AL161626.1 | RTF1 | AL161626.1 |
| LINC01783 | SH3YL1 | LINC01783 |
| CXCL11 | ATP6V1A | CXCL11 |
| H4C8 | CBX3 | H4C8 |
| FABP6-AS1 | MAP2K2 | FABP6-AS1 |
| RN7SKP71 | CLSTN1 | RN7SKP71 |
| RN7SKP203 | ZDHHC13 | RN7SKP203 |
| UBQLNL | LAPTM4A | UBQLNL |
| CTA-384D8.31 | ANXA2 | CTA-384D8.31 |
| AC092299.7 | SEPTIN2 | AC092299.7 |
| BPIFB2 | SLC6A6 | BPIFB2 |
| SIGLEC16 | CLTC | SIGLEC16 |
| TRPV1 | STIM2 | TRPV1 |
| H2BC8 | RCBTB1 | H2BC8 |
| FAM27E3 | PSMD14 | FAM27E3 |
| RNA5SP481 | PCYOX1 | RNA5SP481 |
| H4C2 | SACM1L | H4C2 |
| LINC02176 | ST14 | LINC02176 |
| MIR663AHG | ACVR1B | MIR663AHG |

| | | |
|---|---|---|
| SCARNA7 | SEC61A1 | SCARNA7 |
| C1QB | SYNGR2 | C1QB |
| GDF5 | IL13RA1 | GDF5 |
| SIGLEC1 | CAPZA2 | SIGLEC1 |
| CA6 | XPO7 | CA6 |
| SCARNA6 | CANX | SCARNA6 |
| MZB1 | GALNT1 | MZB1 |
| CD8A | SDC1 | CD8A |
| HTN3 | MFSD14B | HTN3 |
| AC087276.3 | DDX18 | AC087276.3 |
| MOCS1 | CCNDBP1 | MOCS1 |
| AC084082.1 | ADGRF1 | AC084082.1 |
| RN7SKP80 | UBE2N | RN7SKP80 |
| CARNS1 | MDFIC | CARNS1 |
| MT-RNR2 | SSR1 | MT-RNR2 |
| SIGLEC11 | CNOT8 | SIGLEC11 |
| AP001462.1 | SDCBP | AP001462.1 |
| THSD4-AS1 | RAB5C | THSD4-AS1 |
| AL354919.1 | ERP44 | AL354919.1 |
| STATH | MPZL2 | STATH |
| AL137779.1 | HMGXB3 | AL137779.1 |
| BSN | RSRC2 | BSN |
| CYP2D8P | KPNA1 | CYP2D8P |
| H2AC8 | HSPA9 | H2AC8 |
| SKAP2 | AKR1C3 | SKAP2 |
| SNX29P1 | ALCAM | SNX29P1 |
| G2E3-AS1 | CTNNB1 | G2E3-AS1 |
| AC016168.4 | GAPDH | AC016168.4 |
| IKZF3 | GSN | IKZF3 |
| PPP1R1A | TFCP2L1 | PPP1R1A |
| H2BC4 | PDCD4 | H2BC4 |
| CHAC1 | DYNLT1 | CHAC1 |
| AL596325.2 | QSOX1 | AL596325.2 |
| MAP1LC3C | GOLGA1 | MAP1LC3C |
| HOPX | ESYT2 | HOPX |
| IFIT2 | HSP90AA1 | IFIT2 |
| RN7SKP255 | RBBP9 | RN7SKP255 |
| CXCL10 | AAGAB | CXCL10 |
| KRT78 | CD46 | KRT78 |
| AC068987.2 | TAP1 | AC068987.2 |
| RN7SL391P | HIGD1A | RN7SL391P |
| ARHGAP40 | CLDN7 | ARHGAP40 |
| ADRA2A | GM2A | ADRA2A |
| CNFN | KIF21A | CNFN |
| TERB1 | MARK3 | TERB1 |
| TRIM34 | PAICS | TRIM34 |
| SLC9C2 | NFKB1 | SLC9C2 |
| CACNG8 | HNRNPA2B1 | CACNG8 |
| CLVS1 | EIF5B | CLVS1 |
| H1-2 | WEE1 | H1-2 |
| SPRR2D | CDC42SE2 | SPRR2D |
| SLC23A3 | USP48 | SLC23A3 |
| AC005921.4 | ANKRD36B | AC005921.4 |
| LMX1B | HNRNPL | LMX1B |
| PGBD5 | RCAN3 | PGBD5 |
| C8orf34 | ALDH3A1 | C8orf34 |
| SSPO | SRPRA | SSPO |
| FLG2 | FMO3 | FLG2 |
| AP000708.1 | CLCN3 | AP000708.1 |
| AC006435.4 | UGP2 | AC006435.4 |
| H2BC5 | NIPAL2 | H2BC5 |

| | | |
|---|---|---|
| FGF17 | NCKAP1 | FGF17 |
| ABCA8 | RAB5A | ABCA8 |
| LINC01562 | SLC38A2 | LINC01562 |
| FXYD6 | EIF1AX | FXYD6 |
| ARHGAP15 | DECR1 | ARHGAP15 |
| THRB-IT1 | SLC25A3 | THRB-IT1 |
| LINC01460 | SLC44A2 | LINC01460 |
| AC073571.1 | CP | AC073571.1 |
| MUC5B | PSAP | MUC5B |
| RN7SL274P | GRN | RN7SL274P |
| COL9A3 | SLC20A1 | COL9A3 |
| CROCC2 | MARCHF5 | CROCC2 |
| AC005696.4 | IGFBP3 | AC005696.4 |
| ABI3 | SON | ABI3 |
| BACH2 | GOT1 | BACH2 |
| MAL | SLC15A2 | MAL |
| AC091132.5 | ACADM | AC091132.5 |
| ZBP1 | SLC30A7 | ZBP1 |
| AL445665.1 | CCT5 | AL445665.1 |
| SSUH2 | SORT1 | SSUH2 |
| RGL4 | TMEM87B | RGL4 |
| PPP1R1B | VAV1 | PPP1R1B |
| AP001207.3 | PLRG1 | AP001207.3 |
| NR1I3 | FBP1 | NR1I3 |
| POU2F2 | EIF2S3 | POU2F2 |
| LINC01134 | IARS2 | LINC01134 |
| AC138866.2 | TMBIM6 | AC138866.2 |
| AC079949.1 | MAOA | AC079949.1 |
| AC138866.1 | HSPA5 | AC138866.1 |
| IFIT1 | B4GALT5 | IFIT1 |
| LINC02832 | COX6C | LINC02832 |
| CRIP1 | CHL1 | CRIP1 |
| RBM34 | MFSD1 | RBM34 |
| MIR99AHG | LINC01578 | MIR99AHG |
| IRF8 | BUB3 | IRF8 |
| IL15RA | TRA2A | IL15RA |
| DYSF | ATP5PB | DYSF |
| AC114498.1 | POLR2B | AC114498.1 |
| LINGO4 | ACSL3 | LINGO4 |
| AC011466.4 | SRSF3 | AC011466.4 |
| AC015967.2 | DHX29 | AC015967.2 |
| STAC2 | RAB1A | STAC2 |
| TGM3 | SCNN1A | TGM3 |
| BCL2L14 | WASHC2C | BCL2L14 |
| C10orf82 | IMPAD1 | C10orf82 |
| AP001020.2 | GALNT12 | AP001020.2 |
| CDYL2 | LAMP2 | CDYL2 |
| AC099489.1 | ATXN10 | AC099489.1 |
| FRMD6-AS1 | SPTLC1 | FRMD6-AS1 |
| OASL | FDX1 | OASL |
| LETM2 | TM4SF1 | LETM2 |
| MX2 | TMEM50B | MX2 |
| AL691477.1 | ATP5F1A | AL691477.1 |
| AL162258.2 | RBM6 | AL162258.2 |
| PLEKHD1 | B2M | PLEKHD1 |
| VASH2 | NPC2 | VASH2 |
| HERC5 | PERP | HERC5 |
| AC004815.1 | ELOC | AC004815.1 |
| RNF139-AS1 | ANXA3 | RNF139-AS1 |
| ZFHX2 | SLBP | ZFHX2 |
| CMPK2 | TMED10 | CMPK2 |

| | | |
|---|---|---|
| RUSC2 | CPA4 | RUSC2 |
| CPED1 | EFTUD2 | CPED1 |
| ZBTB8A | LDHB | ZBTB8A |
| ODF3B | MMUT | ODF3B |
| IFITM1 | EEF1A1P6 | IFITM1 |
| ECM1 | SRI | ECM1 |
| CD53 | LINC00511 | CD53 |
| IFIT3 | SSB | IFIT3 |
| AC016590.1 | SKP1 | AC016590.1 |
| FCER1G | AKIRIN1 | FCER1G |
| HTN1 | DERL1 | HTN1 |
| GUSBP3 | ATRX | GUSBP3 |
| ZNF324B | UNC5B | ZNF324B |
| AEN | TMEM30B | AEN |
| NHLRC4 | SLC44A1 | NHLRC4 |
| CHDC2 | LTA4H | CHDC2 |
| FABP6 | SEPHS2 | FABP6 |
| H2AC20 | ACTG1 | H2AC20 |
| ZNF250 | IRF2BPL | ZNF250 |
| PTGER2 | AQP5 | PTGER2 |
| IFITM3 | NDUFS8 | IFITM3 |
| PLEKHM3 | STT3B | PLEKHM3 |
| SMAD9 | A4GALT | SMAD9 |
| SPRR2A | TUBA1A | SPRR2A |
| TBX6 | COPS8 | TBX6 |
| ZC3H3 | NDUFA10 | ZC3H3 |
| CD37 | PRXL2A | CD37 |
| IFI44L | GDE1 | IFI44L |
| CROCC | DARS1 | CROCC |
| AL031282.2 | ANXA1 | AL031282.2 |
| PIK3AP1 | NUCB2 | PIK3AP1 |
| SLC2A5 | ID1 | SLC2A5 |
| ISG15 | FMO2 | ISG15 |
| ZNF579 | COG2 | ZNF579 |
| RSAD2 | PSMC5 | RSAD2 |
| SYT12 | RSRP1 | SYT12 |
| AC127164.1 | NDFIP1 | AC127164.1 |
| AKNA | B4GALT1 | AKNA |
| AC025580.3 | RPL6P27 | AC025580.3 |
| WDR49 | KIFAP3 | WDR49 |
| XRRA1 | F3 | XRRA1 |
| AC027243.1 | KARS1 | AC027243.1 |
| GUSBP2 | OSTC | GUSBP2 |
| ING1 | IDH1 | ING1 |
| H4C14 | JAG1 | H4C14 |
| AC020741.1 | GLT8D1 | AC020741.1 |
| RAMP2-AS1 | KYNU | RAMP2-AS1 |
| C1orf229 | CMTM6 | C1orf229 |
| H4C15 | SDR16C5 | H4C15 |
| NUPR1 | EMB | NUPR1 |
| AC009646.2 | CCDC47 | AC009646.2 |
| BCDIN3D | PROM1 | BCDIN3D |
| ISG20 | APP | ISG20 |
| MT2A | MIPEP | MT2A |
| CFAP46 | C2CD2 | CFAP46 |
| MUC5AC | TMEM33 | MUC5AC |
| AC111149.2 | CD164 | AC111149.2 |
| CHP2 | MDH1 | CHP2 |
| SPTB | TFCP2 | SPTB |
| CCL5 | TMPRSS11D | CCL5 |
| DLEC1 | NR2F2 | DLEC1 |

| | | |
|---|---|---|
| RNF222 | VPS26A | RNF222 |
| LBHD1 | CAST | LBHD1 |
| NLRC3 | GSTK1 | NLRC3 |
| IL1RN | YWHAB | IL1RN |
| P2RX7 | TXNDC17 | P2RX7 |
| EPHA10 | CCN2 | EPHA10 |
| PNRC1 | CKMT1A | PNRC1 |
| PLPPR2 | PSMD11 | PLPPR2 |
| FER1L5 | PKM | FER1L5 |
| MAPK12 | METTL21A | MAPK12 |
| RP11-706O15.5 | LAPTM4B | RP11-706O15.5 |
| AL592211.1 | PMPCB | AL592211.1 |
| PML | PRNP | PML |
| PLAAT2 | SEC63 | PLAAT2 |
| AC138932.1 | TMEM165 | AC138932.1 |
| CA5A | CAV2 | CA5A |
| USP18 | PYGL | USP18 |
| SPI1 | ARPC5 | SPI1 |
| HNRNPA1P40 | TMEM150C | HNRNPA1P40 |
| SPRR2E | CD63 | SPRR2E |
| MCF2L | PLS3 | MCF2L |
| FGR | HLF | FGR |
| PPP1R16B | AC087473.1 | PPP1R16B |
| CCDC33 | ATP6V0D1 | CCDC33 |
| ENKD1 | PSMB3 | ENKD1 |
| MARCKSL1 | NOMO1 | MARCKSL1 |
| AC026523.2 | GTF2E2 | AC026523.2 |
| CFAP74 | OAT | CFAP74 |
| HMOX1 | CD151 | HMOX1 |
| GPR65 | ALDH2 | GPR65 |
| NINL | FBXO3 | NINL |
| AP001107.1 | MORN2 | AP001107.1 |
| MICB | CTBS | MICB |
| OAS2 | STOM | OAS2 |
| UBE2T | RNF149 | UBE2T |
| MAP4K2 | MAP3K5 | MAP4K2 |
| RPL37 | ERLIN1 | RPL37 |
| IRF7 | ERMP1 | IRF7 |
| NTAN1 | SMIM15 | NTAN1 |
| IFI27 | CCDC80 | IFI27 |
| DTX2P1 | HSP90B1 | DTX2P1 |
| AC027290.3 | CKMT1B | AC027290.3 |
| ABHD8 | TSPAN3 | ABHD8 |
| UBE2L6 | STAM2 | UBE2L6 |
| AC019117.3 | ABHD5 | AC019117.3 |
| TEKT2 | MSMO1 | TEKT2 |
| AC118344.4 | RBM5 | AC118344.4 |
| NATD1 | TM9SF3 | NATD1 |
| BEX2 | ATP1B1 | BEX2 |
| GNL3L | SARAF | GNL3L |
| SRCIN1 | AMFR | SRCIN1 |
| SAMD9 | DYNC2LI1 | SAMD9 |
| LINC01551 | H3-3A | LINC01551 |
| SGTB | RNF145 | SGTB |
| TNFRSF14-AS1 | ERG28 | TNFRSF14-AS1 |
| SERPING1 | TUBA1C | SERPING1 |
| BICD2 | RBM3 | BICD2 |
| SRGAP3-AS2 | GLB1 | SRGAP3-AS2 |
| DUSP5 | PGD | DUSP5 |
| PRR29 | USP47 | PRR29 |
| AC008079.1 | TAGLN2 | AC008079.1 |

| | | |
|---|---|---|
| CD96 | RETREG2 | CD96 |
| RP11-706O15.3 | TMX4 | RP11-706O15.3 |
| RN7SL718P | EXOC1 | RN7SL718P |
| AHNAK2 | MAT2B | AHNAK2 |
| NIN | VPS25 | NIN |
| OCEL1 | NUP107 | OCEL1 |
| CYP2F1 | ADAM15 | CYP2F1 |
| CCND3 | ARPC3 | CCND3 |
| RPS15 | SYPL1 | RPS15 |
| MUC13 | TGFBR1 | MUC13 |
| AKAP12 | DSG2 | AKAP12 |
| GUSBP1 | RPL4P4 | GUSBP1 |
| SPINK5 | SELENOP | SPINK5 |
| ZNF76 | EPCAM | ZNF76 |
| PTCHD4 | S100A6 | PTCHD4 |
| CRYBG3 | COG6 | CRYBG3 |
| ZFP36 | ANKRD10 | ZFP36 |
| CARD16 | WDR33 | CARD16 |
| DHX35 | LRRC8D | DHX35 |
| SAMD9L | MMP14 | SAMD9L |
| ST3GAL2 | RWDD4 | ST3GAL2 |
| PPDPF | SDF4 | PPDPF |
| EYA1 | DLD | EYA1 |
| SP2 | FHL2 | SP2 |
| CCDC40 | SLC35F5 | CCDC40 |
| IFIH1 | TACSTD2 | IFIH1 |
| AMOTL2 | ASAH1 | AMOTL2 |
| NUMA1 | GUF1 | NUMA1 |
| FAM222A | ENO1 | FAM222A |
| CCDC88C | LUC7L3 | CCDC88C |
| RND1 | MTDH | RND1 |
| UBA52 | JKAMP | UBA52 |
| ARHGAP39 | H2AZ2 | ARHGAP39 |
| WDR62 | ERAP1 | WDR62 |
| AC134407.2 | TMEM192 | AC134407.2 |
| SCGB1A1 | RCN1 | SCGB1A1 |
| ATF3 | WASH6P | ATF3 |
| ZNF500 | ITGB5 | ZNF500 |
| ALPK3 | SLC35A2 | ALPK3 |
| RP11-589F5.3 | TTC19 | RP11-589F5.3 |
| TICAM1 | OCIAD1 | TICAM1 |
| UBAP2 | ADAM28 | UBAP2 |
| CEP135 | P4HB | CEP135 |
| ZNF329 | SFN | ZNF329 |
| CCDC78 | SLC39A7 | CCDC78 |
| C12orf50 | EIF4A3 | C12orf50 |
| EVI2B | TMEM30A | EVI2B |
| YY1AP1 | GPD2 | YY1AP1 |
| TRIM14 | CD59 | TRIM14 |
| NKX3-1 | GTF2H2B | NKX3-1 |
| CCDC159 | PON2 | CCDC159 |
| JPX | HACD3 | JPX |
| ZNF335 | GSS | ZNF335 |
| MRTFA | TUFM | MRTFA |
| FOXG1 | ATP2C1 | FOXG1 |
| CCDC106 | CACHD1 | CCDC106 |
| SYTL5 | CA12 | SYTL5 |
| COTL1 | MBOAT2 | COTL1 |
| OAS1 | THYN1 | OAS1 |
| H2AC6 | PDIA6 | H2AC6 |
| AC004151.1 | LEMD3 | AC004151.1 |

| | | |
|---|---|---|
| MUC21 | RPN1 | MUC21 |
| HOOK1 | PTTG1IP | HOOK1 |
| H2BC18 | PRDX1 | H2BC18 |
| OAS3 | GAA | OAS3 |
| PREX1 | ABCC3 | PREX1 |
| CCDC88B | IER3IP1 | CCDC88B |
| FTL | EGFR | FTL |
| FOSB | ANXA4 | FOSB |
| WNK2 | IGFBP2 | WNK2 |
| MAST3 | SLC27A2 | MAST3 |
| EPSTI1 | TRAPPC3 | EPSTI1 |
| PCNT | OS9 | PCNT |
| NFATC3 | SLC18B1 | NFATC3 |
| HIGD2A | ALOX15 | HIGD2A |
| KIAA0040 | GNS | KIAA0040 |
| C15orf62 | RRN3 | C15orf62 |
| CEP112 | PIGX | CEP112 |
| ARHGAP26 | ARL6IP1 | ARHGAP26 |
| SGSM1 | RPL7AP66 | SGSM1 |
| MVB12B | AC004069.1 | MVB12B |
| SPAG9 | LGR4 | SPAG9 |
| SYNPO | LRG1 | SYNPO |
| PKD1P5 | SPARCL1 | PKD1P5 |
| SHANK2 | TMEM87A | SHANK2 |
| CLEC16A | ALDH1A1 | CLEC16A |
| ELF4 | LMAN2 | ELF4 |
| NCCRP1 | AC007318.1 | NCCRP1 |
| H2AC18 | DDOST | H2AC18 |
| HELB | CHPF | HELB |
| FCGR2A | CTSB | FCGR2A |
| ATP5F1E | RASA1 | ATP5F1E |
| SIPA1L3 | SLC2A1 | SIPA1L3 |
| H2AC19 | CFH | H2AC19 |
| SEC24A | TM9SF2 | SEC24A |
| POLR2M | EML3 | POLR2M |
| GNG5 | CTSH | GNG5 |
| KAT2B | HNRNPA1P4 | KAT2B |
| A2ML1 | ERLIN2 | A2ML1 |
| TNFAIP3 | MTRNR2L9 | TNFAIP3 |
| HSF1 | LRP5 | HSF1 |
| MFN1 | SEMA3A | MFN1 |
| AC126755.1 | GFM2 | AC126755.1 |
| ALMS1 | HNRNPA1L2 | ALMS1 |
| FBXO48 | PIGG | FBXO48 |
| RABIF | RRM1 | RABIF |
| SLC25A23 | TSPAN13 | SLC25A23 |
| MCUB | TMEM68 | MCUB |
| SOBP | KRT5 | SOBP |
| HELZ2 | ACVR1 | HELZ2 |
| CBX6 | ATP1A1 | CBX6 |
| SRCAP | PRODH | SRCAP |
| TACC2 | RPAP2 | TACC2 |
| FAM193A | MFSD11 | FAM193A |
| SERPINB8 | FDFT1 | SERPINB8 |
| PRPF3 | MFSD14C | PRPF3 |
| SYTL2 | ANKRD36 | SYTL2 |
| MBNL1 | SGK1 | MBNL1 |
| PATL1 | NEU1 | PATL1 |
| ZMIZ2 | CLDN4 | ZMIZ2 |
| ZNF358 | SCGB2A1 | ZNF358 |
| BLZF1 | ERLEC1 | BLZF1 |

| | | |
|---|---|---|
| STK10 | EEF1A1P5 | STK10 |
| NIBAN1 | AL391121.1 | NIBAN1 |
| CDK18 | POR | CDK18 |
| PHC3 | RAB3IP | PHC3 |
| FYCO1 | FAM3B | FYCO1 |
| LGALS9B | CALR | LGALS9B |
| TEX9 | AC073333.1 | TEX9 |
| MX1 | EBPL | MX1 |
| FNDC3B | NPTN | FNDC3B |
| CLPB | HSD17B13 | CLPB |
| PRRC2A | RPS21 | PRRC2A |
| APOL3 | NIFK | APOL3 |
| CENPBD1P1 | ALS2CL | CENPBD1P1 |
| RAB8A | PSEN2 | RAB8A |
| DAP | RCN2 | DAP |
| ZNF592 | CHKA | ZNF592 |
| GTDC1 | GTF2H2C | GTDC1 |
| MOB3A | TMA7 | MOB3A |
| TADA2B | ALDH7A1 | TADA2B |
| SH3KBP1 | CD81 | SH3KBP1 |
| SAMD4B | RTN4 | SAMD4B |
| TRAF3IP2 | DHX36 | TRAF3IP2 |
| SPDEF | SNRPA1 | SPDEF |
| CDKN2B | DNAJC1 | CDKN2B |
| RBMS2 | HNRNPA1P35 | RBMS2 |
| CIZ1 | FTH1P10 | CIZ1 |
| DUSP3 | HNRNPAB | DUSP3 |
| RFX5 | FUCA2 | RFX5 |
| MAP3K11 | ADK | MAP3K11 |
| TENT5C | FMO5 | TENT5C |
| TBC1D15 | RTN3 | TBC1D15 |
| RPL36AL | ASCC3 | RPL36AL |
| R3HDM2 | PSME1 | R3HDM2 |
| MLPH | WASH4P | MLPH |
| S100A11 | TFB2M | S100A11 |
| RND3 | REEP5 | RND3 |
| UBB | TMEM51 | UBB |
| PAK4 | WSB1 | PAK4 |
| RASSF9 | SREK1 | RASSF9 |
| ATXN7 | RTN3P1 | ATXN7 |
| RUNDC1 | RARS1 | RUNDC1 |
| IQCE | HADH | IQCE |
| R3HDM1 | MBOAT1 | R3HDM1 |
| NEK6 | PGAP4 | NEK6 |
| JADE2 | FAAH2 | JADE2 |
| TCOF1 | SMG1P4 | TCOF1 |
| CSNK1G2 | AC008810.1 | CSNK1G2 |
| TOB2 | TLR2 | TOB2 |
| ATN1 | ZMPSTE24 | ATN1 |
| NCOR2 | HNRNPA1P10 | NCOR2 |
| IL1R1 | PAPSS2 | IL1R1 |
| SPECC1 | AGL | SPECC1 |
| ANKRD17 | TMEM9 | ANKRD17 |
| BICDL1 | TMEM9B | BICDL1 |
| RAB3B | TSPAN1 | RAB3B |
| VPS37B | CLN5 | VPS37B |
| CRY2 | TMEM59 | CRY2 |
| ACSS1 | AL592114.1 | ACSS1 |
| WARS1 | GUSB | WARS1 |
| OPTN | ANAPC4 | OPTN |
| ZNF609 | COMMD7 | ZNF609 |

| | | |
|---|---|---|
| PDE4DIP | MAP3K6 | PDE4DIP |
| FCHSD2 | ANKRD66 | FCHSD2 |
| SP3 | DSE | SP3 |
| WWC1 | GCLC | WWC1 |
| MLXIP | PRSS8 | MLXIP |
| ZFP36L2 | CYP4F12 | ZFP36L2 |
| SORBS3 | ABCE1 | SORBS3 |
| SND1 | ENPP4 | SND1 |
| FOXK1 | ABCA5 | FOXK1 |
| GTPBP1 | GLUD2 | GTPBP1 |
| CCDC69 | CALM2P2 | CCDC69 |
| C6orf132 | KLHL42 | C6orf132 |
| | PRMT7 | LINC00963 |
| | ITGB6 | ADGRG1 |
| | SESN1 | SRSF5 |
| | FTH1P2 | ACTN4 |
| | CAMK2G | EWSR1 |
| | STAM | HK1 |
| | MANF | TMED2 |
| | THOC3 | CCZ1B |
| | MSTO1 | PSMD2 |
| | ZPR1 | VAMP3 |
| | INPP1 | CTNND1 |
| | NOMO2 | ARRDC3 |
| | LRRN1 | RDH10 |
| | SRSF11 | EIF3CL |
| | SELENOI | PRKAR1A |
| | TOPORS | PPP2CB |
| | PLTP | TSPYL1 |
| | SLC39A6 | SRP9 |
| | TMEM106C | SMARCA2 |
| | CD47 | GLUL |
| | SORD2P | RBM25 |
| | AL158206.1 | TNFRSF21 |
| | KRT10 | SELENBP1 |
| | EIF4A1P4 | DHCR24 |
| | UNC93B3 | MAT2A |
| | CXADR | EIF3C |
| | CUEDC1 | SCARB2 |
| | IL10RB | TMPRSS4 |
| | YWHAZP4 | EIF4G1 |
| | ATP5F1B | ENC1 |
| | PRKAR2B | HNRNPH1 |
| | SMC4 | WDR1 |
| | PTDSS1 | ADSS2 |
| | COPG2 | TMEM123 |
| | GTF2F2 | IL33 |
| | LONRF1 | CCZ1 |
| | RPL10P9 | FBXL5 |
| | HLA-B | ZBTB7A |
| | RFNG | HNRNPK |
| | MAN1B1 | PLEKHB2 |
| | EIF4HP1 | HNRNPU |
| | H3P36 | ARL8B |
| | LACTB | DHX15 |
| | CNIH1 | UBQLN1 |
| | MAPKAPK3 | ANXA7 |
| | JPT1 | NDRG2 |
| | SMAP1 | TOMM20 |
| | CYP3A5 | GANAB |
| | SPTSSA | CTBP2 |

| | | |
|---|---|---|
| | TTC29 | CBWD6 |
| | C1D | ITGB1 |
| | PA2G4P6 | ASPH |
| | DPAGT1 | ATP8B1 |
| | SMARCAD1 | PFN2 |
| | RPL7AP6 | DDB1 |
| | BX679664.3 | ARGLU1 |
| | PLS1 | SF3B1 |
| | F11R | ANPEP |
| | RMDN3 | DDX1 |
| | DDAH2 | NCSTN |
| | BCAP29 | DNAJC10 |
| | FSCN1 | DDX5 |
| | AKAP1 | IFT57 |
| | ITFG1 | PGK1 |
| | ARVCF | MIA3 |
| | LOXL4 | SDC4 |
| | AC090498.1 | VPS35 |
| | FAM3D | GALNT7 |
| | HNRNPA1P7 | NAP1L4 |
| | HNRNPA1P12 | SRSF10 |
| | RHBDL2 | COPB1 |
| | MPP7 | DDX17 |
| | UBE2E3 | RBM39 |
| | SCPEP1 | TAX1BP1 |
| | DNAJB11 | ANKRD12 |
| | SDHA | LMBRD1 |
| | LAMB3 | CALM2 |
| | VARS1 | SLC37A3 |
| | NT5DC1 | RDX |
| | PBXIP1 | ABCD3 |
| | TAP2 | CKAP4 |
| | HLA-F | TMEM41B |
| | CDH1 | NFE2L1 |
| | PRELID1 | EPRS1 |
| | UPK1B | UBE2Q1 |
| | PSMD1 | ERBB3 |
| | TMTC4 | DIMT1 |
| | ITGAV | RAC1 |
| | NUDT12 | CALM1 |
| | PGAM4 | PRSS23 |
| | RNF26 | SFPQ |
| | CCDC65 | EXOC3 |
| | FARSB | ADAM9 |
| | PARP2 | XPO1 |
| | VIPR1 | ACSL5 |
| | ARL3 | CCT2 |
| | ENY2 | KDELR1 |
| | SDCBPP3 | PCMTD2 |
| | BCAP31 | ACTR2 |
| | B4GALT4 | ALDH3A2 |
| | RAMAC | RTF1 |
| | C5orf15 | SH3YL1 |
| | FTH1P8 | ATP6V1A |
| | MAD2L1BP | CBX3 |
| | RAC1P2 | MAP2K2 |
| | NAAA | CLSTN1 |
| | GLULP4 | ZDHHC13 |
| | EXOSC8 | LAPTM4A |
| | MT-TY | ANXA2 |
| | BLOC1S4 | SEPTIN2 |

| | | |
|---|---|---|
| | AC144530.1 | SLC6A6 |
| | TMEM45B | CLTC |
| | ABHD3 | STIM2 |
| | HLA-L | RCBTB1 |
| | AC113935.1 | PSMD14 |
| | PRPF39 | PCYOX1 |
| | CTNNAL1 | SACM1L |
| | VPS35P1 | ST14 |
| | HSP90AB2P | ACVR1B |
| | UNC93B7 | SEC61A1 |
| | CHPT1 | SYNGR2 |
| | TNC | IL13RA1 |
| | DDX50 | CAPZA2 |
| | MLYCD | XPO7 |
| | AQP3 | CANX |
| | LRRCC1 | GALNT1 |
| | TPBG | SDC1 |
| | GJB3 | MFSD14B |
| | PABPC1P4 | DDX18 |
| | SLC25A17 | CCNDBP1 |
| | SLC44A3 | ADGRF1 |
| | SLC26A4 | UBE2N |
| | PRDX4 | MDFIC |
| | GGH | SSR1 |
| | HNRNPA1P8 | CNOT8 |
| | LSAMP | SDCBP |
| | CDS2 | RAB5C |
| | FAT1 | ERP44 |
| | ATP6AP2 | MPZL2 |
| | MRAP2 | HMGXB3 |
| | G6PC3 | RSRC2 |
| | ERO1B | KPNA1 |
| | CLK1 | HSPA9 |
| | SNX14 | AKR1C3 |
| | PTGES3P3 | ALCAM |
| | CCT6A | CTNNB1 |
| | AC006511.4 | GAPDH |
| | KLF9 | GSN |
| | FKBP1C | TFCP2L1 |
| | PIPSL | PDCD4 |
| | FAM162A | DYNLT1 |
| | AC024293.1 | QSOX1 |
| | TMEM147 | GOLGA1 |
| | ALG1 | ESYT2 |
| | CCDC59 | HSP90AA1 |
| | DBP | RBBP9 |
| | TF | AAGAB |
| | FKBP9 | CD46 |
| | RPL14P1 | TAP1 |
| | TMEM43 | HIGD1A |
| | ACAD10 | CLDN7 |
| | BORCS5 | GM2A |
| | EMC7 | KIF21A |
| | ACTA1 | MARK3 |
| | UNC93B6 | PAICS |
| | EDEM2 | NFKB1 |
| | PTK7 | HNRNPA2B1 |
| | ARSDP1 | EIF5B |
| | TMED7 | WEE1 |
| | TUSC3 | CDC42SE2 |
| | PGRMC1 | USP48 |

| | | |
|---|---|---|
| | UBLCP1 | ANKRD36B |
| | AC026271.1 | HNRNPL |
| | BMI1 | RCAN3 |
| | DYNC1I2P1 | ALDH3A1 |
| | ARMT1 | SRPRA |
| | TLR1 | FMO3 |
| | GPC1 | CLCN3 |
| | VDAC1P1 | UGP2 |
| | HNRNPKP4 | NIPAL2 |
| | RRAGA | NCKAP1 |
| | KIT | RAB5A |
| | ST13P3 | SLC38A2 |
| | AC136632.1 | EIF1AX |
| | AL391244.2 | DECR1 |
| | ETHE1 | SLC25A3 |
| | TMED1 | SLC44A2 |
| | RPL7P10 | CP |
| | HEXB | PSAP |
| | MOSPD1 | GRN |
| | SLC52A2 | SLC20A1 |
| | KDSR | MARCHF5 |
| | SELENOS | IGFBP3 |
| | HLA-A | SON |
| | ADH1A | GOT1 |
| | RPL22P1 | SLC15A2 |
| | DNAJB9 | ACADM |
| | AC083873.1 | SLC30A7 |
| | DENND10P1 | CCT5 |
| | FUCA1 | SORT1 |
| | HSPA8P5 | TMEM87B |
| | LPCAT3 | VAV1 |
| | SLC35B2 | PLRG1 |
| | EXOSC9 | FBP1 |
| | TAGLN2P1 | EIF2S3 |
| | ATP5MC3 | IARS2 |
| | EIF3FP3 | TMBIM6 |
| | COL6A3 | MAOA |
| | CHCHD1 | HSPA5 |
| | TMCO1 | B4GALT5 |
| | PABPC3 | COX6C |
| | TP63 | CHL1 |
| | CTSC | MFSD1 |
| | HSD17B12 | LINC01578 |
| | CEACAM3 | BUB3 |
| | SMG1P1 | TRA2A |
| | PSMC1P1 | ATP5PB |
| | ELMO3 | POLR2B |
| | AL158801.6 | ACSL3 |
| | UQCRC1 | SRSF3 |
| | HLA-V | DHX29 |
| | MGST1 | RAB1A |
| | NTS | SCNN1A |
| | AL133477.1 | WASHC2C |
| | H3P44 | IMPAD1 |
| | PSENEN | GALNT12 |
| | RPL7AP34 | LAMP2 |
| | PSPC1 | ATXN10 |
| | LAMC1 | SPTLC1 |
| | AL121769.1 | FDX1 |
| | NDUFA12 | TM4SF1 |
| | MT-TL1 | TMEM50B |

| | | |
|---|---|---|
| | AC106795.1 | ATP5F1A |
| | AC022968.1 | RBM6 |
| | LRRIQ1 | B2M |
| | LRRC17 | NPC2 |
| | UFD1 | PERP |
| | UXS1 | ELOC |
| | AC008065.1 | ANXA3 |
| | KRT6B | SLBP |
| | HMGN2P17 | TMED10 |
| | ALOX15P1 | CPA4 |
| | RPL37AP1 | EFTUD2 |
| | ATP6AP1 | LDHB |
| | AC099670.1 | MMUT |
| | BMP3 | EEF1A1P6 |
| | FKSG70 | SRI |
| | FAM171A1 | LINC00511 |
| | RPN2 | SSB |
| | HLA-H | SKP1 |
| | EEF1A1P38 | AKIRIN1 |
| | THNSL2 | DERL1 |
| | PDIA3 | ATRX |
| | AC064799.1 | UNC5B |
| | KTN1 | TMEM30B |
| | AC209007.1 | SLC44A1 |
| | AC104619.3 | LTA4H |
| | SYT8 | SEPHS2 |
| | COQ5 | ACTG1 |
| | MSH2 | IRF2BPL |
| | NPC1 | AQP5 |
| | UQCRFS1P1 | NDUFS8 |
| | MORF4L1P1 | STT3B |
| | UBBP4 | A4GALT |
| | GPR89A | TUBA1A |
| | MFSD5 | COPS8 |
| | SERPINB4 | NDUFA10 |
| | HNRNPA1P48 | PRXL2A |
| | AP000936.3 | GDE1 |
| | SIRT3 | DARS1 |
| | AC092115.2 | ANXA1 |
| | RPS27P29 | NUCB2 |
| | CYP26A1 | ID1 |
| | ATP13A5 | FMO2 |
| | AL109918.1 | COG2 |
| | PIGO | PSMC5 |
| | ITM2B | RSRP1 |
| | RPL9P32 | NDFIP1 |
| | PFN1P1 | B4GALT1 |
| | EIF2S2P4 | RPL6P27 |
| | HNRNPA3P6 | KIFAP3 |
| | HSP90AB3P | F3 |
| | PLLP | KARS1 |
| | CLDN1 | OSTC |
| | ANXA8L1 | IDH1 |
| | RPL23P8 | JAG1 |
| | AC113404.3 | GLT8D1 |
| | HSP90AA6P | KYNU |
| | RPL7P47 | CMTM6 |
| | SETP20 | SDR16C5 |
| | PHF14 | EMB |
| | SRD5A3 | CCDC47 |
| | SCAMP3 | PROM1 |

| | | |
|---|---|---|
| | AC244034.1 | APP |
| | TRIAP1 | MIPEP |
| | F2RL1 | C2CD2 |
| | PPIAP87 | TMEM33 |
| | AP002784.2 | CD164 |
| | SUMO2P1 | MDH1 |
| | AC005000.1 | TFCP2 |
| | HLA-C | TMPRSS11D |
| | MTATP8P1 | NR2F2 |
| | KRT18P16 | VPS26A |
| | SERBP1P5 | CAST |
| | RPL4P5 | GSTK1 |
| | DCAF13 | YWHAB |
| | SRSF2 | TXNDC17 |
| | SETSIP | CCN2 |
| | FKBP9P1 | CKMT1A |
| | PRCP | PSMD11 |
| | CD9 | PKM |
| | IFNGR1 | METTL21A |
| | RPL7P9 | LAPTM4B |
| | TMEM212 | PMPCB |
| | EIF4BP3 | PRNP |
| | PSCA | SEC63 |
| | AL049597.1 | TMEM165 |
| | RPL7P32 | CAV2 |
| | YWHAZP5 | PYGL |
| | AC012085.1 | ARPC5 |
| | HNRNPCP2 | TMEM150C |
| | FTLP3 | CD63 |
| | APLP2 | PLS3 |
| | SETP14 | HLF |
| | UBE2I | AC087473.1 |
| | CDC42P6 | ATP6V0D1 |
| | ANXA8 | PSMB3 |
| | AL354702.1 | NOMO1 |
| | FGFR3 | GTF2E2 |
| | H3-5 | OAT |
| | TSPAN6 | CD151 |
| | EIF4A1P2 | ALDH2 |
| | RPL7P1 | FBXO3 |
| | GAPDHP65 | MORN2 |
| | AC105250.1 | CTBS |
| | EPHA1 | STOM |
| | AC002075.2 | RNF149 |
| | RPS7P10 | MAP3K5 |
| | EEF1A1P19 | ERLIN1 |
| | EIF5AL1 | ERMP1 |
| | RPL10AP2 | SMIM15 |
| | ARSD | CCDC80 |
| | XRCC6P2 | HSP90B1 |
| | RPL13AP20 | CKMT1B |
| | AC092597.1 | TSPAN3 |
| | EIF4BP6 | STAM2 |
| | AC099560.2 | ABHD5 |
| | RPL12P38 | MSMO1 |
| | RPS26P6 | RBM5 |
| | LDHBP2 | TM9SF3 |
| | KRT18P11 | ATP1B1 |
| | AC004057.1 | SARAF |
| | RPL34P26 | AMFR |
| | BZW1P2 | DYNC2LI1 |

| | | |
|---|---|---|
| | AC092683.1 | H3-3A |
| | S100A4 | RNF145 |
| | HLA-DRB6 | ERG28 |
| | C1GALT1C1 | TUBA1C |
| | RPS26P31 | RBM3 |
| | SLC5A8 | GLB1 |
| | MTCO1P40 | PGD |
| | GAPDHP44 | USP47 |
| | RPL7AP11 | TAGLN2 |
| | PPIC | RETREG2 |
| | TMX1 | TMX4 |
| | ITGA6 | EXOC1 |
| | GAPDHP61 | MAT2B |
| | PSMC1P5 | VPS25 |
| | EIF4BP7 | NUP107 |
| | PPIAP16 | ADAM15 |
| | EEF1A1P7 | ARPC3 |
| | RPSAP19 | SYPL1 |
| | AC005480.2 | TGFBR1 |
| | UNC50 | DSG2 |
| | MIR22HG | RPL4P4 |
| | RPS26P8 | SELENOP |
| | AC115223.1 | EPCAM |
| | HSPA8P1 | S100A6 |
| | DPYD | COG6 |
| | AC126120.1 | ANKRD10 |
| | AC016734.1 | WDR33 |
| | RARRES1 | LRRC8D |
| | MTCO3P12 | MMP14 |
| | DPY30 | RWDD4 |
| | ALG5 | SDF4 |
| | PPIAP66 | DLD |
| | PPIAP43 | FHL2 |
| | AC112187.1 | SLC35F5 |
| | PPIAL4C | TACSTD2 |
| | MTND4P12 | ASAH1 |
| | TMEM183B | GUF1 |
| | ACTBP2 | ENO1 |
| | HLA-G | LUC7L3 |
| | RPS7P11 | MTDH |
| | NACA3P | JKAMP |
| | EEF1A1P29 | H2AZ2 |
| | AC068522.1 | ERAP1 |
| | RPS26P11 | TMEM192 |
| | AC034236.1 | RCN1 |
| | MTCO2P2 | WASH6P |
| | AC020898.1 | ITGB5 |
| | CLCA4 | SLC35A2 |
| | LYPD3 | TTC19 |
| | EEF1A1P4 | OCIAD1 |
| | AC004552.1 | ADAM28 |
| | NAMPTP1 | P4HB |
| | PPIAP13 | SFN |
| | RPS4XP22 | SLC39A7 |
| | EEF1A1P25 | EIF4A3 |
| | TUBAP2 | TMEM30A |
| | CROT | GPD2 |
| | AP000281.2 | CD59 |
| | RPL10P12 | GTF2H2B |
| | FTH1P15 | PON2 |
| | RPL3P4 | HACD3 |

| | | |
|---|---|---|
| | AC104339.1 | GSS |
| | RPS23P8 | TUFM |
| | PPIAP31 | ATP2C1 |
| | AC135178.7 | CACHD1 |
| | RPL10P4 | CA12 |
| | AC104563.1 | MBOAT2 |
| | RPS26P47 | THYN1 |
| | H3P16 | PDIA6 |
| | FTH1P11 | LEMD3 |
| | EEF1A1P16 | RPN1 |
| | YWHAZP3 | PTTG1IP |
| | GAPDHP73 | PRDX1 |
| | GAPDHP63 | GAA |
| | AL627402.1 | ABCC3 |
| | AC009245.1 | IER3IP1 |
| | RPS3AP5 | EGFR |
| | H3P6 | ANXA4 |
| | RPL27AP5 | IGFBP2 |
| | RPS26P15 | SLC27A2 |
| | H3P47 | TRAPPC3 |
| | AC078819.1 | OS9 |
| | TCN1 | SLC18B1 |
| | HSP90AA2P | ALOX15 |
| | FTH1P7 | GNS |
| | RPS7P1 | RRN3 |
| | RPS27AP16 | PIGX |
| | RPL7P19 | ARL6IP1 |
| | TMSB4XP2 | RPL7AP66 |
| | RPS26P28 | AC004069.1 |
| | RPL10AP6 | LGR4 |
| | PPIAP22 | LRG1 |
| | EEF1A1P8 | SPARCL1 |
| | ADH1B | TMEM87A |
| | ATP1B3 | ALDH1A1 |
| | PPIAP6 | LMAN2 |
| | RPL3P2 | AC007318.1 |
| | RPL13AP25 | DDOST |
| | MTND6P4 | CHPF |
| | RPS24P8 | CTSB |
| | AC092865.1 | RASA1 |
| | RPL15P20 | SLC2A1 |
| | AL133260.1 | CFH |
| | FTH1P16 | TM9SF2 |
| | EEF1A1P11 | EML3 |
| | RPS15AP1 | CTSH |
| | FTH1P3 | HNRNPA1P4 |
| | APOD | ERLIN2 |
| | AL596275.1 | MTRNR2L9 |
| | RPL15P18 | LRP5 |
| | RPL17P22 | SEMA3A |
| | KRT6C | GFM2 |
| | EIF4A1P10 | HNRNPA1L2 |
| | PDIA3P1 | PIGG |
| | FTH1P20 | RRM1 |
| | EEF1A1P22 | TSPAN13 |
| | ANXA2P2 | TMEM68 |
| | AC073072.1 | KRT5 |
| | AC025518.1 | ACVR1 |
| | S100A2 | ATP1A1 |
| | RPL7P23 | PRODH |
| | MTND6P3 | RPAP2 |

| | | |
|---|---|---|
| | AL009174.1 | MFSD11 |
| | AC092670.1 | FDFT1 |
| | PPIAP29 | MFSD14C |
| | FTH1P12 | ANKRD36 |
| | AC090543.3 | SGK1 |
| | H3C9P | NEU1 |
| | RPL17P36 | CLDN4 |
| | MTND5P11 | SCGB2A1 |
| | MT-TE | ERLEC1 |
| | MTCO2P12 | EEF1A1P5 |
| | MTRNR2L1 | AL391121.1 |
| | HLA-J | POR |
| | EEF1A1P12 | RAB3IP |
| | EEF1A1P14 | FAM3B |
| | AC091429.1 | CALR |
| | FTH1P5 | AC073333.1 |
| | TPT1P9 | EBPL |
| | EEF1A1P13 | NPTN |
| | AC006386.2 | HSD17B13 |
| | AC012005.1 | RPS21 |
| | RPL41P2 | NIFK |
| | MT-TA | ALS2CL |
| | | PSEN2 |
| | | RCN2 |
| | | CHKA |
| | | GTF2H2C |
| | | TMA7 |
| | | ALDH7A1 |
| | | CD81 |
| | | RTN4 |
| | | DHX36 |
| | | SNRPA1 |
| | | DNAJC1 |
| | | HNRNPA1P35 |
| | | FTH1P10 |
| | | HNRNPAB |
| | | FUCA2 |
| | | ADK |
| | | FMO5 |
| | | RTN3 |
| | | ASCC3 |
| | | PSME1 |
| | | WASH4P |
| | | TFB2M |
| | | REEP5 |
| | | TMEM51 |
| | | WSB1 |
| | | SREK1 |
| | | RTN3P1 |
| | | RARS1 |
| | | HADH |
| | | MBOAT1 |
| | | PGAP4 |
| | | FAAH2 |
| | | SMG1P4 |
| | | AC008810.1 |
| | | TLR2 |
| | | ZMPSTE24 |
| | | HNRNPA1P10 |
| | | PAPSS2 |
| | | AGL |

| | | |
|---|---|---|
| | | TMEM9 |
| | | TMEM9B |
| | | TSPAN1 |
| | | CLN5 |
| | | TMEM59 |
| | | AL592114.1 |
| | | GUSB |
| | | ANAPC4 |
| | | COMMD7 |
| | | MAP3K6 |
| | | ANKRD66 |
| | | DSE |
| | | GCLC |
| | | PRSS8 |
| | | CYP4F12 |
| | | ABCE1 |
| | | ENPP4 |
| | | ABCA5 |
| | | GLUD2 |
| | | CALM2P2 |
| | | KLHL42 |
| | | PRMT7 |
| | | ITGB6 |
| | | SESN1 |
| | | FTH1P2 |
| | | CAMK2G |
| | | STAM |
| | | MANF |
| | | THOC3 |
| | | MSTO1 |
| | | ZPR1 |
| | | INPP1 |
| | | NOMO2 |
| | | LRRN1 |
| | | SRSF11 |
| | | SELENOI |
| | | TOPORS |
| | | PLTP |
| | | SLC39A6 |
| | | TMEM106C |
| | | CD47 |
| | | SORD2P |
| | | AL158206.1 |
| | | KRT10 |
| | | EIF4A1P4 |
| | | UNC93B3 |
| | | CXADR |
| | | CUEDC1 |
| | | IL10RB |
| | | YWHAZP4 |
| | | ATP5F1B |
| | | PRKAR2B |
| | | SMC4 |
| | | PTDSS1 |
| | | COPG2 |
| | | GTF2F2 |
| | | LONRF1 |
| | | RPL10P9 |
| | | HLA-B |
| | | RFNG |
| | | MAN1B1 |

| | | |
|---|---|---|
| | | EIF4HP1 |
| | | H3P36 |
| | | LACTB |
| | | CNIH1 |
| | | MAPKAPK3 |
| | | JPT1 |
| | | SMAP1 |
| | | CYP3A5 |
| | | SPTSSA |
| | | TTC29 |
| | | C1D |
| | | PA2G4P6 |
| | | DPAGT1 |
| | | SMARCAD1 |
| | | RPL7AP6 |
| | | BX679664.3 |
| | | PLS1 |
| | | F11R |
| | | RMDN3 |
| | | DDAH2 |
| | | BCAP29 |
| | | FSCN1 |
| | | AKAP1 |
| | | ITFG1 |
| | | ARVCF |
| | | LOXL4 |
| | | AC090498.1 |
| | | FAM3D |
| | | HNRNPA1P7 |
| | | HNRNPA1P12 |
| | | RHBDL2 |
| | | MPP7 |
| | | UBE2E3 |
| | | SCPEP1 |
| | | DNAJB11 |
| | | SDHA |
| | | LAMB3 |
| | | VARS1 |
| | | NT5DC1 |
| | | PBXIP1 |
| | | TAP2 |
| | | HLA-F |
| | | CDH1 |
| | | PRELID1 |
| | | UPK1B |
| | | PSMD1 |
| | | TMTC4 |
| | | ITGAV |
| | | NUDT12 |
| | | PGAM4 |
| | | RNF26 |
| | | CCDC65 |
| | | FARSB |
| | | PARP2 |
| | | VIPR1 |
| | | ARL3 |
| | | ENY2 |
| | | SDCBPP3 |
| | | BCAP31 |
| | | B4GALT4 |
| | | RAMAC |

| | | |
|---|---|---|
| | | C5orf15 |
| | | FTH1P8 |
| | | MAD2L1BP |
| | | RAC1P2 |
| | | NAAA |
| | | GLULP4 |
| | | EXOSC8 |
| | | MT-TY |
| | | BLOC1S4 |
| | | AC144530.1 |
| | | TMEM45B |
| | | ABHD3 |
| | | HLA-L |
| | | AC113935.1 |
| | | PRPF39 |
| | | CTNNAL1 |
| | | VPS35P1 |
| | | HSP90AB2P |
| | | UNC93B7 |
| | | CHPT1 |
| | | TNC |
| | | DDX50 |
| | | MLYCD |
| | | AQP3 |
| | | LRRCC1 |
| | | TPBG |
| | | GJB3 |
| | | PABPC1P4 |
| | | SLC25A17 |
| | | SLC44A3 |
| | | SLC26A4 |
| | | PRDX4 |
| | | GGH |
| | | HNRNPA1P8 |
| | | LSAMP |
| | | CDS2 |
| | | FAT1 |
| | | ATP6AP2 |
| | | MRAP2 |
| | | G6PC3 |
| | | ERO1B |
| | | CLK1 |
| | | SNX14 |
| | | PTGES3P3 |
| | | CCT6A |
| | | AC006511.4 |
| | | KLF9 |
| | | FKBP1C |
| | | PIPSL |
| | | FAM162A |
| | | AC024293.1 |
| | | TMEM147 |
| | | ALG1 |
| | | CCDC59 |
| | | DBP |
| | | TF |
| | | FKBP9 |
| | | RPL14P1 |
| | | TMEM43 |
| | | ACAD10 |
| | | BORCS5 |

| | | |
|---|---|---|
| | | EMC7 |
| | | ACTA1 |
| | | UNC93B6 |
| | | EDEM2 |
| | | PTK7 |
| | | ARSDP1 |
| | | TMED7 |
| | | TUSC3 |
| | | PGRMC1 |
| | | UBLCP1 |
| | | AC026271.1 |
| | | BMI1 |
| | | DYNC1I2P1 |
| | | ARMT1 |
| | | TLR1 |
| | | GPC1 |
| | | VDAC1P1 |
| | | HNRNPKP4 |
| | | RRAGA |
| | | KIT |
| | | ST13P3 |
| | | AC136632.1 |
| | | AL391244.2 |
| | | ETHE1 |
| | | TMED1 |
| | | RPL7P10 |
| | | HEXB |
| | | MOSPD1 |
| | | SLC52A2 |
| | | KDSR |
| | | SELENOS |
| | | HLA-A |
| | | ADH1A |
| | | RPL22P1 |
| | | DNAJB9 |
| | | AC083873.1 |
| | | DENND10P1 |
| | | FUCA1 |
| | | HSPA8P5 |
| | | LPCAT3 |
| | | SLC35B2 |
| | | EXOSC9 |
| | | TAGLN2P1 |
| | | ATP5MC3 |
| | | EIF3FP3 |
| | | COL6A3 |
| | | CHCHD1 |
| | | TMCO1 |
| | | PABPC3 |
| | | TP63 |
| | | CTSC |
| | | HSD17B12 |
| | | CEACAM3 |
| | | SMG1P1 |
| | | PSMC1P1 |
| | | ELMO3 |
| | | AL158801.6 |
| | | UQCRC1 |
| | | HLA-V |
| | | MGST1 |
| | | NTS |

| | | |
|---|---|---|
| | | AL133477.1 |
| | | H3P44 |
| | | PSENEN |
| | | RPL7AP34 |
| | | PSPC1 |
| | | LAMC1 |
| | | AL121769.1 |
| | | NDUFA12 |
| | | MT-TL1 |
| | | AC106795.1 |
| | | AC022968.1 |
| | | LRRIQ1 |
| | | LRRC17 |
| | | UFD1 |
| | | UXS1 |
| | | AC008065.1 |
| | | KRT6B |
| | | HMGN2P17 |
| | | ALOX15P1 |
| | | RPL37AP1 |
| | | ATP6AP1 |
| | | AC099670.1 |
| | | BMP3 |
| | | FKSG70 |
| | | FAM171A1 |
| | | RPN2 |
| | | HLA-H |
| | | EEF1A1P38 |
| | | THNSL2 |
| | | PDIA3 |
| | | AC064799.1 |
| | | KTN1 |
| | | AC209007.1 |
| | | AC104619.3 |
| | | SYT8 |
| | | COQ5 |
| | | MSH2 |
| | | NPC1 |
| | | UQCRFS1P1 |
| | | MORF4L1P1 |
| | | UBBP4 |
| | | GPR89A |
| | | MFSD5 |
| | | SERPINB4 |
| | | HNRNPA1P48 |
| | | AP000936.3 |
| | | SIRT3 |
| | | AC092115.2 |
| | | RPS27P29 |
| | | CYP26A1 |
| | | ATP13A5 |
| | | AL109918.1 |
| | | PIGO |
| | | ITM2B |
| | | RPL9P32 |
| | | PFN1P1 |
| | | EIF2S2P4 |
| | | HNRNPA3P6 |
| | | HSP90AB3P |
| | | PLLP |
| | | CLDN1 |

| | | |
|---|---|---|
| | | ANXA8L1 |
| | | RPL23P8 |
| | | AC113404.3 |
| | | HSP90AA6P |
| | | RPL7P47 |
| | | SETP20 |
| | | PHF14 |
| | | SRD5A3 |
| | | SCAMP3 |
| | | AC244034.1 |
| | | TRIAP1 |
| | | F2RL1 |
| | | PPIAP87 |
| | | AP002784.2 |
| | | SUMO2P1 |
| | | AC005000.1 |
| | | HLA-C |
| | | MTATP8P1 |
| | | KRT18P16 |
| | | SERBP1P5 |
| | | RPL4P5 |
| | | DCAF13 |
| | | SRSF2 |
| | | SETSIP |
| | | FKBP9P1 |
| | | PRCP |
| | | CD9 |
| | | IFNGR1 |
| | | RPL7P9 |
| | | TMEM212 |
| | | EIF4BP3 |
| | | PSCA |
| | | AL049597.1 |
| | | RPL7P32 |
| | | YWHAZP5 |
| | | AC012085.1 |
| | | HNRNPCP2 |
| | | FTLP3 |
| | | APLP2 |
| | | SETP14 |
| | | UBE2I |
| | | CDC42P6 |
| | | ANXA8 |
| | | AL354702.1 |
| | | FGFR3 |
| | | H3-5 |
| | | TSPAN6 |
| | | EIF4A1P2 |
| | | RPL7P1 |
| | | GAPDHP65 |
| | | AC105250.1 |
| | | EPHA1 |
| | | AC002075.2 |
| | | RPS7P10 |
| | | EEF1A1P19 |
| | | EIF5AL1 |
| | | RPL10AP2 |
| | | ARSD |
| | | XRCC6P2 |
| | | RPL13AP20 |
| | | AC092597.1 |

| | | |
|---|---|---|
| | | EIF4BP6 |
| | | AC099560.2 |
| | | RPL12P38 |
| | | RPS26P6 |
| | | LDHBP2 |
| | | KRT18P11 |
| | | AC004057.1 |
| | | RPL34P26 |
| | | BZW1P2 |
| | | AC092683.1 |
| | | S100A4 |
| | | HLA-DRB6 |
| | | C1GALT1C1 |
| | | RPS26P31 |
| | | SLC5A8 |
| | | MTCO1P40 |
| | | GAPDHP44 |
| | | RPL7AP11 |
| | | PPIC |
| | | TMX1 |
| | | ITGA6 |
| | | GAPDHP61 |
| | | PSMC1P5 |
| | | EIF4BP7 |
| | | PPIAP16 |
| | | EEF1A1P7 |
| | | RPSAP19 |
| | | AC005480.2 |
| | | UNC50 |
| | | MIR22HG |
| | | RPS26P8 |
| | | AC115223.1 |
| | | HSPA8P1 |
| | | DPYD |
| | | AC126120.1 |
| | | AC016734.1 |
| | | RARRES1 |
| | | MTCO3P12 |
| | | DPY30 |
| | | ALG5 |
| | | PPIAP66 |
| | | PPIAP43 |
| | | AC112187.1 |
| | | PPIAL4C |
| | | MTND4P12 |
| | | TMEM183B |
| | | ACTBP2 |
| | | HLA-G |
| | | RPS7P11 |
| | | NACA3P |
| | | EEF1A1P29 |
| | | AC068522.1 |
| | | RPS26P11 |
| | | AC034236.1 |
| | | MTCO2P2 |
| | | AC020898.1 |
| | | CLCA4 |
| | | LYPD3 |
| | | EEF1A1P4 |
| | | AC004552.1 |
| | | NAMPTP1 |

| | | |
|---|---|---|
| | | PPIAP13 |
| | | RPS4XP22 |
| | | EEF1A1P25 |
| | | TUBAP2 |
| | | CROT |
| | | AP000281.2 |
| | | RPL10P12 |
| | | FTH1P15 |
| | | RPL3P4 |
| | | AC104339.1 |
| | | RPS23P8 |
| | | PPIAP31 |
| | | AC135178.7 |
| | | RPL10P4 |
| | | AC104563.1 |
| | | RPS26P47 |
| | | H3P16 |
| | | FTH1P11 |
| | | EEF1A1P16 |
| | | YWHAZP3 |
| | | GAPDHP73 |
| | | GAPDHP63 |
| | | AL627402.1 |
| | | AC009245.1 |
| | | RPS3AP5 |
| | | H3P6 |
| | | RPL27AP5 |
| | | RPS26P15 |
| | | H3P47 |
| | | AC078819.1 |
| | | TCN1 |
| | | HSP90AA2P |
| | | FTH1P7 |
| | | RPS7P1 |
| | | RPS27AP16 |
| | | RPL7P19 |
| | | TMSB4XP2 |
| | | RPS26P28 |
| | | RPL10AP6 |
| | | PPIAP22 |
| | | EEF1A1P8 |
| | | ADH1B |
| | | ATP1B3 |
| | | PPIAP6 |
| | | RPL3P2 |
| | | RPL13AP25 |
| | | MTND6P4 |
| | | RPS24P8 |
| | | AC092865.1 |
| | | RPL15P20 |
| | | AL133260.1 |
| | | FTH1P16 |
| | | EEF1A1P11 |
| | | RPS15AP1 |
| | | FTH1P3 |
| | | APOD |
| | | AL596275.1 |
| | | RPL15P18 |
| | | RPL17P22 |
| | | KRT6C |
| | | EIF4A1P10 |

| | | |
|---|---|---|
| | | PDIA3P1 |
| | | FTH1P20 |
| | | EEF1A1P22 |
| | | ANXA2P2 |
| | | AC073072.1 |
| | | AC025518.1 |
| | | S100A2 |
| | | RPL7P23 |
| | | MTND6P3 |
| | | AL009174.1 |
| | | AC092670.1 |
| | | PPIAP29 |
| | | FTH1P12 |
| | | AC090543.3 |
| | | H3C9P |
| | | RPL17P36 |
| | | MTND5P11 |
| | | MT-TE |
| | | MTCO2P12 |
| | | MTRNR2L1 |
| | | HLA-J |
| | | EEF1A1P12 |
| | | EEF1A1P14 |
| | | AC091429.1 |
| | | FTH1P5 |
| | | TPT1P9 |
| | | EEF1A1P13 |
| | | AC006386.2 |
| | | AC012005.1 |
| | | RPL41P2 |
| | | MT-TA |
