## Supplementary file 5 for "Host transcriptional responses and SARS-CoV-2 isolates from the nasopharyngeal samples of Bangladeshi COVID-19 patients"

**Supplementary file 5: Differentially expressed genes in different SARS-CoV-2 infected cell types.**

| BD_Nasal | GSE147507_Lung | GSE150316_Lung | GSE147507_NHBE | GSE148729_CALU3 |
|---|---|---|---|---|
| RNA5S14 | AAK1 | IGHV4-34 | AC006058.4 | ZC3HAV1 |
| RNA5S11 | AAMP | IGHV4-59 | AC092964.1 | DDX58 |
| RNA5S8 | AAR2 | MIR205HG | AC099336.2 | OAS3 |
| RNA5S1 | AARS1 | IGKV2-24 | AC144530.1 | OAS2 |
| RNA5S10 | AASS | IGLV3-27 | AL445490.1 | GBP1 |
| RNA5S13 | AATF | IGHV3-73 | ALDH1A3 | IFIT3 |
| RNA5S15 | ABCA1 | IGHM | ANXA3 | IFIT2 |
| RNA5S2 | ABCB10 | IGLV1-47 | AOAH | ZNFX1 |
| RNA5S16 | ABCB11 | FAM26F | AP001324.1 | HELZ2 |
| RNA5S3 | ABCB7 | IGLV3-21 | BCL2A1 | TRIM22 |
| RNA5S6 | ABCC10 | IGLV9-49 | BCL2L11 | RSAD2 |
| RNA5S17 | ABCC2 | IGHV5-51 | C1QTNF1 | CMPK2 |
| RNA5S7 | ABCC3 | APLNR | C1R | HERC5 |
| RNA5S12 | ABCD4 | IGLV3-25 | C3 | MX1 |
| RNA5S5 | ABCF1 | IGKV1-12 | CAMK2D | GBP4 |
| RNA5S4 | ABCF2 | CD3D | CCDC14 | IFI27 |
| RNU1-28P | ABCF3 | IGLL5 | CCL20 | TAP1 |
| RNVU1-18 | ABCG2 | LAG3 | CCNA2 | CXCL10 |
| RNU1-4 | ABHD12B | IGHV1-18 | CD3E | ISG20 |
| RNVU1-29 | ABHD13 | IGHV3-74 | CENPA | PARP14 |
| RNU1-2 | ABHD14B | IGKV1-39 | CEP55 | MX2 |
| RNU1-1 | ABHD17A | DERL3 | CES1 | IFIT1 |
| RNU1-27P | ABHD2 | KRT4 | CES1P1 | APOL6 |
| RNU1-3 | ABHD3 | IGHV3-48 | CFB | WARS1 |
| RNVU1-7 | ABHD5 | HIST1H3G | CLEC5A | IDO1 |
| RNY1 | ABHD6 | IGHV3-49 | COL8A1 | LAMP3 |
| RNU4-2 | ABI1 | FOLR2 | CPA4 | IFI35 |
| RNA5S9 | ABL1 | IGHG4 | CSF3 | ISG15 |
| FP671120.2 | ABL2 | IGHGP | CSTB | TRANK1 |
| CR392039.1 | ABLIM1 | HIST1H2BL | CTSH | OAS1 |
| VTRNA1-1 | ABLIM3 | IGHV3-53 | CXCL16 | TNFSF10 |
| RNA5SP370 | ABR | CD27 | CXCL5 | IFITM1 |
| FP236383.6 | ABRACL | HIST1H3C | CXCL9 | PMAIP1 |
| RNA5SP202 | ABT1 | IGHV3-15 | DUSP4 | IRF1 |
| RNVU1-28 | ABTB1 | HIST1H2BF | EDN1 | IRF7 |
| CDR1 | AC000089.1 | HIST1H2AH | EEF1A1P13 | SAMD9 |
| SNORA73B | AC002075.2 | LTB | EFNA1 | SAMHD1 |
| RNU1-11P | AC004057.1 | HIST1H2BO | EIF4BP3 | JUN |
| RNY4 | AC004151.1 | KIF2C | EIF4BP6 | TXNIP |
| FP236383.1 | AC004158.1 | HIST1H3F | FAT2 | PARP10 |
| RNA5SP389 | AC004453.1 | HIST1H2BH | FBXW7 | GSDMB |
| RNU4-1 | AC004552.1 | RAMP3 | FCGBP | IFIT5 |
| FP236383.9 | AC004846.1 | APOL4 | GAS6-AS1 | HCP5 |
| H4C5 | AC004846.2 | HIST1H2BI | GCLC | PPP1R15A |
| MIR3648-1 | AC004847.1 | HIST1H2AL | HELZ2 | DHX58 |
| SMR3B | AC004877.2 | PSMB10 | HK3 | CCL5 |
| H4C11 | AC005005.4 | SNORA13 | HSH2D | UBE2L6 |
| RNA5-8SN2 | AC005083.1 | CCNB1 | HSPB1P1 | RNF213 |
| RNA5-8SN3 | AC005229.4 | EXOC3L1 | ICAM1 | HLA-B |
| FP671120.4 | AC005261.2 | RNU1-28P | IFI27 | PLEKHA4 |
| RNA5-8SN1 | AC005332.6 | ITM2C | IFI44L | HLA-F |
| FP236383.3 | AC005336.1 | LTF | IFI6 | STAT2 |
| MIR3648-2 | AC005912.1 | C12orf75 | IFITM1 | CX3CL1 |
| FP671120.1 | AC006001.4 | HIST1H1B | IFITM10 | XAF1 |
| FP236383.10 | AC006252.1 | HIST1H3I | IKBKE | TAP2 |
| RN7SL5P | AC006262.1 | HIST1H3B | IKZF1 | DDX60 |

| | | | | |
|---|---|---|---|---|
| RNA28S5 | AC006386.2 | MYBL2 | IL10 | USP18 |
| PRB4 | AC006435.2 | CLEC3B | IL6 | LGALS9 |
| PRB3 | AC006511.4 | PLVAP | IRF7 | IFI16 |
| RNA5SP298 | AC006548.28 | KIFC1 | ITGA5 | NFKBIA |
| FP236383.5 | AC007336.1 | HIST1H2AG | IVL | HLA-E |
| FP236383.4 | AC007384.1 | HIST1H2BN | KB-1507C5.4 | PTGER4 |
| FP236383.12 | AC007686.3 | VAMP5 | KRT6B | SAMD9L |
| FP671120.7 | AC007743.1 | MMP28 | LINC00504 | DDX60L |
| RN7SL4P | AC007881.1 | DLGAP5 | LTB | IFI44 |
| RNA5SP226 | AC007906.2 | ICAM2 | MAFF | AL669918.1 |
| RN7SL752P | AC008147.2 | RARRES3 | MAP3K8 | HERC6 |
| RNY3 | AC008537.3 | HIST2H2AB | MAP7D2 | KLF4 |
| AC079601.1 | AC008555.2 | IFI27L2 | MCM4 | TIPARP |
| RNA5SP145 | AC008676.1 | HIST1H3D | MIR155HG | DTX3L |
| RNA5SP149 | AC008687.2 | LIMD2 | MIR3142HG | CXCL8 |
| RNA5SP429 | AC008760.2 | BUB1 | MME | RTP4 |
| AL135938.1 | AC008764.2 | STYXL1 | MMP9 | TENT5A |
| RNA5SP74 | AC008969.1 | HIST1H2AE | MX1 | NFKBIZ |
| CRNN | AC009005.1 | CKS2 | MX2 | MAP3K8 |
| RNY3P1 | AC009054.2 | BANK1 | MYLK | PARP12 |
| FAM27E2 | AC009126.1 | MRPL2 | NANOS1 | OASL |
| FP236383.7 | AC009245.1 | NCF1 | NCCRP1 | STAT1 |
| RN7SL396P | AC009275.1 | RAMP2 | NEDD9 | PLAAT4 |
| FP671120.5 | AC009283.1 | SP140 | NHBE_DE | IFNL1 |
| H1-4 | AC009303.4 | HIST1H2BG | NID1 | NLRC5 |
| AL162581.1 | AC009318.1 | NDUFA3 | NT5DC3 | SP110 |
| CRCT1 | AC009961.1 | CIITA | OAS1 | CEACAM1 |
| RNA5SP335 | AC010173.1 | THYN1 | OR7E47P | CCL20 |
| RNA5-8SP6 | AC010226.1 | ITGB4 | OSBP2 | BATF2 |
| AC024051.11 | AC010326.4 | GIMAP7 | P2RY6 | PARP9 |
| AC024051.5 | AC010970.1 | HDAC9 | PARP12 | UBA7 |
| AC024051.12 | AC011472.2 | CORO1A | PCDH17 | PSMB8 |
| AC024051.1 | AC011498.7 | CLEC14A | PDCD4 | TEP1 |
| FP236383.8 | AC011933.1 | CRIP1 | PDGFB | ETS2 |
| RNU2-1 | AC012085.1 | PSD4 | PDZK1IP1 | CFB |
| AC024051.2 | AC012368.1 | FCGRT | PGLYRP4 | VIM |
| SNORD17 | AC015912.3 | TXNDC5 | PLA1A | NMI |
| AC024051.7 | AC015922.3 | HN1 | PLAT | TNFAIP2 |
| AC024051.10 | AC016142.1 | SNRPD2 | PLSCR1 | GBP3 |
| AC024051.3 | AC016745.2 | HIST1H2BC | POU2F2 | HLA-C |
| AC024051.8 | AC016747.1 | CXorf36 | PPARGC1A | PMEPA1 |
| AC024051.6 | AC016957.2 | PLCG2 | PRELID1 | APOL1 |
| RNA5SP161 | AC018638.1 | HIST1H2BD | PRELID2 | NCOA7 |
| AC024051.4 | AC018638.2 | LMAN2 | PRF1 | TRIM5 |
| AL161626.1 | AC018868.1 | PSME1 | RAB15 | IFIH1 |
| LINC01783 | AC019322.6 | NCAPD2 | RAB7B | PML |
| CXCL11 | AC020612.3 | CLU | RFTN1 | HLA-A |
| H4C8 | AC020765.2 | RFX5 | RND1 | IFITM2 |
| FABP6-AS1 | AC020916.1 | ATPIF1 | RPL10AP6 | AHNAK |
| RN7SKP71 | AC021106.1 | PRDX5 | RPL21P16 | SNORD3A |
| RN7SKP203 | AC022149.1 | PPP1CA | RPL37P2 | JAK2 |
| UBQLNL | AC022211.4 | PSMD4 | RPL7AP50 | LAP3 |
| CTA-384D8.31 | AC022274.1 | SPAG9 | RPL9 | DUSP1 |
| AC092299.7 | AC022400.9 | SERTAD2 | RPS2P55 | XRN1 |
| BPIFB2 | AC022613.1 | ROCK2 | S100P | SLC15A3 |
| SIGLEC16 | AC022916.2 | CREB3L2 | SAA1 | BBC3 |
| TRPV1 | AC022929.2 | FAM168B | SAA2 | B2M |
| H2BC8 | AC022966.1 | NABP1 | SEC14L4 | LY6E |
| FAM27E3 | AC023157.1 | PIK3R1 | SERPINB1 | KDM6A |
| RNA5SP481 | AC023157.3 | AFF4 | SMTN | BIRC3 |

| | | | | |
|---|---|---|---|---|
| H4C2 | AC023590.1 | DDX21 | SOD2 | FNDC3A |
| LINC02176 | AC025518.1 | SNX13 | SPRR2B | NEAT1 |
| MIR663AHG | AC025580.2 | PDZD8 | SPRR2D | SCD |
| SCARNA7 | AC025884.1 | BCL2L2 | SPRR2E | HLA-H |
| C1QB | AC027801.1 | SLIT3 | ST3GAL5 | IL1A |
| GDF5 | AC034236.1 | CHSY1 | STAT5A | OPTN |
| SIGLEC1 | AC040160.1 | LRRTM2 | STRBP | RN7SL3 |
| CA6 | AC040162.1 | FOXO3 | SUSD6 | PPM1K |
| SCARNA6 | AC040970.1 | YBX3 | TCF4 | NUB1 |
| MZB1 | AC055733.1 | LPP | TCIM | CD74 |
| CD8A | AC060766.4 | CLIC4 | TFCP2L1 | NFKB2 |
| HTN3 | AC060766.7 | FNIP1 | TGM1 | MLEC |
| AC087276.3 | AC060814.2 | TNFAIP3 | TNF | TRIM21 |
| MOCS1 | AC064799.1 | ELK4 | TNFAIP2 | CXCL11 |
| AC084082.1 | AC067930.4 | NUPL1 | TNFAIP3 | KLF6 |
| RN7SKP80 | AC067945.3 | SPRY4 | TNFSF14 | SAT1 |
| CARNS1 | AC068039.4 | TTC19 | TREML4 | ABHD2 |
| MT-RNR2 | AC068448.2 | RLIM | TRIM47 | ETV7 |
| SIGLEC11 | AC068888.1 | AGFG1 | VGLL4 | CFTR |
| AP001462.1 | AC068987.2 | PHLDA1 | VNN1 | EGR1 |
| THSD4-AS1 | AC069547.1 | SLC6A6 | VNN3 | MXD1 |
| AL354919.1 | AC073072.1 | FOXO1 | XAF1 | AFAP1 |
| STATH | AC073343.1 | IMPAD1 | XDH | SLC1A3 |
| AL137779.1 | AC073349.2 | OSBPL8 | YTHDC2 | SNORA73B |
| BSN | AC073548.1 | MAPK6 | ZC3H12A | APOL2 |
| CYP2D8P | AC073861.1 | ZFAND5 | ZCCHC10 | EPSTI1 |
| H2AC8 | AC073869.1 | PPP1R3B | ZNF488 | ATF3 |
| SKAP2 | AC078819.1 | CTB-89H12.4 | | IL6 |
| SNX29P1 | AC079228.1 | PAG1 | | ZFP36 |
| G2E3-AS1 | AC079250.1 | KLF6 | | TRIM38 |
| AC016168.4 | AC079922.1 | NEDD9 | | HLA-DRA |
| IKZF3 | AC079949.1 | TXNRD1 | | CYLD |
| PPP1R1A | AC080023.1 | ENPP4 | | RIPOR2 |
| H2BC4 | AC083843.3 | RP11-463O12.5 | | BTN3A1 |
| CHAC1 | AC083899.1 | KITLG | | OGFR |
| AL596325.2 | AC084871.1 | FLJ42393 | | ERAP2 |
| MAP1LC3C | AC084880.1 | UPP1 | | PLAUR |
| HOPX | AC087473.1 | CSGALNACT2 | | AL645922.1 |
| IFIT2 | AC087623.2 | SLC16A7 | | TMEM140 |
| RN7SKP255 | AC090114.2 | NRCAM | | SNORA73A |
| CXCL10 | AC090543.3 | GK | | DNAJB4 |
| KRT78 | AC090559.1 | EMP1 | | NFE2L3 |
| AC068987.2 | AC090587.2 | BHLHE40 | | NT5C3A |
| RN7SL391P | AC091230.1 | RP11-107E5.3 | | IKZF3 |
| ARHGAP40 | AC091304.1 | AKAP12 | | ARRDC3 |
| ADRA2A | AC092118.1 | NUDT16 | | TDRD7 |
| CNFN | AC092597.1 | GBE1 | | TRIM69 |
| TERB1 | AC092670.1 | ADAMTS9 | | SERPINB2 |
| TRIM34 | AC092683.1 | TMED7 | | TPM1 |
| SLC9C2 | AC092718.4 | VGLL3 | | RASGRP3 |
| CACNG8 | AC092865.1 | SLC6A8 | | TAPBPL |
| CLVS1 | AC092919.3 | PTP4A1 | | DICER1 |
| H1-2 | AC092964.1 | ATP13A3 | | TOP2A |
| SPRR2D | AC093010.3 | NAMPT | | PRKD2 |
| SLC23A3 | AC093323.1 | ADAMTS1 | | SHFL |
| AC005921.4 | AC093690.1 | SLC2A3 | | BST2 |
| LMX1B | AC093752.1 | RHOQ | | PNPT1 |
| PGBD5 | AC097263.1 | PDLIM3 | | PIK3AP1 |
| C8orf34 | AC097523.1 | ERRFI1 | | PTGS2 |
| SSPO | AC098583.1 | BEST1 | | TYMP |

| | | | | |
|---|---|---|---|---|
| FLG2 | AC098851.1 | IER3 | | TRIM26 |
| AP000708.1 | AC099336.2 | SLC7A2 | | NEURL3 |
| AC006435.4 | AC099343.4 | ADAMTS15 | | HSH2D |
| H2BC5 | AC099524.1 | ZNF638-IT1 | | PTX3 |
| FGF17 | AC099560.2 | IGSF10 | | ADAR |
| ABCA8 | AC099789.1 | KLF9 | | MED12L |
| LINC01562 | AC104339.1 | NME9 | | HIP1R |
| FXYD6 | AC104563.1 | CD300E | | CYP1A1 |
| ARHGAP15 | AC104791.2 | KIAA1614 | | CCNL1 |
| THRB-IT1 | AC104981.1 | NCR3LG1 | | MB21D2 |
| LINC01460 | AC105250.1 | IL6 | | SEMA3A |
| AC073571.1 | AC105942.1 | RP11-415J8.3 | | ALCAM |
| MUC5B | AC106795.1 | ADAMTS9-AS1 | | MYD88 |
| RN7SL274P | AC107375.1 | RP11-752L20.3 | | SOD2 |
| COL9A3 | AC108058.1 | SPDYA | | CFAP54 |
| CROCC2 | AC108134.1 | GADD45A | | SP100 |
| AC005696.4 | AC108925.1 | HIF1A-AS2 | | KIF20A |
| ABI3 | AC109322.1 | USP53 | | SYNE1 |
| BACH2 | AC109829.2 | EGLN3 | | SCN3A |
| MAL | AC110285.6 | FAM110C | | INHBA |
| AC091132.5 | AC110749.1 | C11orf96 | | NR3C1 |
| ZBP1 | AC112777.1 | NAV2-AS5 | | OTUD1 |
| AL445665.1 | AC113398.1 | CTD-2033D15.2 | | SLC12A7 |
| SSUH2 | AC113935.1 | GLDN | | FOS |
| RGL4 | AC114498.1 | SLED1 | | IL32 |
| PPP1R1B | AC114728.1 | NAMPTP1 | | TFRC |
| AP001207.3 | AC115223.1 | APOLD1 | | LINC00944 |
| NR1I3 | AC116049.1 | HILPDA | | SCARNA5 |
| POU2F2 | AC116407.2 | FOSB | | FZD4 |
| LINC01134 | AC116533.1 | CYR61 | | FST |
| AC138866.2 | AC116914.2 | EGR1 | | IRAK2 |
| AC079949.1 | AC121761.1 | RP11-417F21.1 | | B4GALT5 |
| AC138866.1 | AC124067.4 | NR4A3 | | GCA |
| IFIT1 | AC125437.1 | SLC19A2 | | SQLE |
| LINC02832 | AC125807.1 | PTX3 | | MT2A |
| CRIP1 | AC125807.2 | PTGS2 | | NUAK2 |
| RBM34 | AC127502.1 | LIF | | TRAFD1 |
| MIR99AHG | AC127502.2 | BRD7P4 | | NFKBIE |
| IRF8 | AC129507.1 | CTD-2026K11.6 | | CFH |
| IL15RA | AC131011.2 | RP11-212E4.1 | | KYNU |
| DYSF | AC131235.2 | FGG | | CNP |
| AC114498.1 | AC132192.2 | RP1-309I22.2 | | GRB10 |
| LINGO4 | AC132812.1 | AC007278.3 | | CD274 |
| AC011466.4 | AC133134.1 | RP11-212I21.3 | | PSMB9 |
| AC015967.2 | AC136475.5 | RP1-29C18.10 | | CSF1 |
| STAC2 | AC136475.9 | RP11-314N13.9 | | RPS6KC1 |
| TGM3 | AC138305.3 | AC007278.2 | | TRIM31 |
| BCL2L14 | AC144530.1 | RP11-635L1.2 | | BTN3A3 |
| C10orf82 | AC145285.2 | SLCO4A1-AS1 | | APOL3 |
| AP001020.2 | AC145350.2 | RP11-393I2.2 | | ANXA4 |
| CDYL2 | AC146944.3 | RP11-291I6.2 | | TGM2 |
| AC099489.1 | AC159540.2 | CTD-2184D3.5 | | MOV10 |
| FRMD6-AS1 | AC211476.10 | SDCBPP1 | | SNHG3 |
| OASL | AC211476.5 | EDN2 | | CTSS |
| LETM2 | AC234781.1 | CCL20 | | HDAC9 |
| MX2 | AC240565.3 | SLC5A8 | | LGMN |
| AL691477.1 | AC240565.4 | | | BRD2 |
| AL162258.2 | AC241952.2 | | | PRKDC |
| PLEKHD1 | AC243964.4 | | | REL |
| VASH2 | AC244669.1 | | | KMT2C |

| | | | | |
|---|---|---|---|---|
| HERC5 | AC245297.1 | | | TNIP1 |
| AC004815.1 | AC245297.4 | | | RMRP |
| RNF139-AS1 | AC246787.1 | | | RMRP |
| ZFHX2 | ACAA1 | | | TNFRSF10B |
| CMPK2 | ACACB | | | CYP24A1 |
| RUSC2 | ACAD8 | | | ZBP1 |
| CPED1 | ACADVL | | | SNORD17 |
| ZBTB8A | ACAP3 | | | CHD2 |
| ODF3B | ACAT2 | | | SPOCK2 |
| IFITM1 | ACBD5 | | | ACACA |
| ECM1 | ACBD6 | | | PGK1 |
| CD53 | ACBD7 | | | FAP |
| IFIT3 | ACD | | | LGALS3BP |
| AC016590.1 | ACHE | | | PLSCR1 |
| FCER1G | ACKR3 | | | DHCR24 |
| HTN1 | ACO2 | | | AL049839.2 |
| GUSBP3 | ACOT1 | | | AF117829.1 |
| ZNF324B | ACOT13 | | | PELI1 |
| AEN | ACOT2 | | | AMBRA1 |
| NHLRC4 | ACOT8 | | | PGAP3 |
| CHDC2 | ACOT9 | | | ADRB2 |
| FABP6 | ACP2 | | | LINC02068 |
| H2AC20 | ACP5 | | | SRPK2 |
| ZNF250 | ACRBP | | | CXCL2 |
| PTGER2 | ACSF2 | | | AREG |
| IFITM3 | ACSF3 | | | CLTRN |
| PLEKHM3 | ACSL1 | | | CBX6 |
| SMAD9 | ACSS1 | | | TNFAIP3 |
| SPRR2A | ACTB | | | CEBPB |
| TBX6 | ACTG1P17 | | | SERPINA3 |
| ZC3H3 | ACTL6A | | | NEDD4 |
| CD37 | ACTN1 | | | F2RL1 |
| IFI44L | ACTN4 | | | SEMA7A |
| CROCC | ACTR1A | | | HMGCR |
| AL031282.2 | ACTR1B | | | UBN2 |
| PIK3AP1 | ACTR3 | | | ARL14 |
| SLC2A5 | ACTR3B | | | TLR3 |
| ISG15 | ACTR5 | | | PIK3R3 |
| ZNF579 | ACVR1 | | | KRT17 |
| RSAD2 | ACVR1B | | | LIF |
| SYT12 | ACVR1C | | | CFLAR |
| AC127164.1 | ACVRL1 | | | SPAG9 |
| AKNA | ADA | | | RCAN1 |
| AC025580.3 | ADA2 | | | PSME2 |
| WDR49 | ADAM11 | | | MIR155HG |
| XRRA1 | ADAM15 | | | TM4SF4 |
| AC027243.1 | ADAM17 | | | CREB5 |
| GUSBP2 | ADAM19 | | | ABL2 |
| ING1 | ADAM1A | | | ATP9A |
| H4C14 | ADAM8 | | | ST3GAL2 |
| AC020741.1 | ADAMTS1 | | | FOSB |
| RAMP2-AS1 | ADAMTS2 | | | APOBEC3G |
| C1orf229 | ADAMTS3 | | | PLXNA2 |
| H4C15 | ADAMTS9 | | | CD59 |
| NUPR1 | ADAMTS9-AS1 | | | CD24 |
| AC009646.2 | ADAMTSL3 | | | SP140L |
| BCDIN3D | ADAMTSL4-AS1 | | | RNVU1-28 |
| ISG20 | ADAMTSL5 | | | GADD45B |
| MT2A | ADAR | | | RIPK2 |
| CFAP46 | ADARB1 | | | ETV3 |

| | | | | |
|---|---|---|---|---|
| MUC5AC | ADAT2 | | | IFITM3 |
| AC111149.2 | ADCK2 | | | SNORA63 |
| CHP2 | ADCY3 | | | LASP1 |
| SPTB | ADCY4 | | | HDLBP |
| CCL5 | ADCY5 | | | ACO1 |
| DLEC1 | ADCY6 | | | CMTM4 |
| RNF222 | ADCY9 | | | SORT1 |
| LBHD1 | ADD1 | | | TSPAN8 |
| NLRC3 | ADD3 | | | MAP4K4 |
| IL1RN | ADGRD1 | | | UNC93B1 |
| P2RX7 | ADGRD2 | | | IRF1-AS1 |
| EPHA10 | ADGRE1 | | | TMBIM6 |
| PNRC1 | ADGRE2 | | | CASC19 |
| PLPPR2 | ADGRE5 | | | VAT1 |
| FER1L5 | ADGRF2 | | | MYH9 |
| MAPK12 | ADGRF5P2 | | | RNF19B |
| RP11-706O15.5 | ADGRG1 | | | BRI3BP |
| AL592211.1 | ADGRG3 | | | TRAF1 |
| PML | ADGRL2 | | | THEMIS2 |
| PLAAT2 | ADH5 | | | BAZ2B |
| AC138932.1 | ADH7 | | | LDHA |
| CA5A | ADIPOR1 | | | HSPA5 |
| USP18 | ADIPOR2 | | | LGALS4 |
| SPI1 | ADIRF-AS1 | | | RELB |
| HNRNPA1P40 | ADK | | | SLC12A2 |
| SPRR2E | ADM5 | | | SEPTIN9 |
| MCF2L | ADNP | | | XBP1 |
| FGR | ADNP2 | | | PLS1 |
| PPP1R16B | ADPRM | | | TNFSF15 |
| CCDC33 | ADRA2A | | | KIAA0100 |
| ENKD1 | ADRM1 | | | NCAPD2 |
| MARCKSL1 | ADSL | | | ATP1A1 |
| AC026523.2 | ADSS1 | | | CCNB2 |
| CFAP74 | ADSS2 | | | SDC1 |
| HMOX1 | AEBP1 | | | SMURF1 |
| GPR65 | AEBP2 | | | ENDOD1 |
| NINL | AEN | | | ZEB2 |
| AP001107.1 | AF165147.1 | | | SLC3A2 |
| MICB | AFAP1 | | | ITGB1 |
| OAS2 | AFAP1L1 | | | BACH2 |
| UBE2T | AFDN | | | ZBTB20 |
| MAP4K2 | AFF4 | | | MCL1 |
| RPL37 | AFG3L1P | | | DCP1A |
| IRF7 | AFG3L2 | | | FRS2 |
| NTAN1 | AFTPH | | | NUDT3 |
| IFI27 | AGAP1 | | | HIVEP2 |
| DTX2P1 | AGAP10P | | | NUCKS1 |
| AC027290.3 | AGAP3 | | | EPHA4 |
| ABHD8 | AGAP4 | | | SDC4 |
| UBE2L6 | AGAP6 | | | N4BP1 |
| AC019117.3 | AGAP7P | | | C3 |
| TEKT2 | AGBL5 | | | MYCBP2 |
| AC118344.4 | AGER | | | IL15RA |
| NATD1 | AGFG1 | | | NEDD9 |
| BEX2 | AGFG2 | | | ACKR4 |
| GNL3L | AGK | | | GPBP1 |
| SRCIN1 | AGMO | | | MSMO1 |
| SAMD9 | AGO1 | | | UHRF1BP1 |
| LINC01551 | AGPAT2 | | | DHCR7 |
| SGTB | AGPAT4 | | | NFAT5 |

| | | | | |
|---|---|---|---|---|
| TNFRSF14-AS1 | AGRN | | | DAB2IP |
| SERPING1 | AGTR1 | | | CMTR1 |
| BICD2 | AGTRAP | | | YBX1 |
| SRGAP3-AS2 | AHCTF1 | | | EEF1A1 |
| DUSP5 | AHDC1 | | | SPP1 |
| PRR29 | AHNAK | | | ARL6IP1 |
| AC008079.1 | AHR | | | MLKL |
| CD96 | AHRR | | | TRIM25 |
| RP11-706O15.3 | AHSA2P | | | ANXA2P2 |
| RN7SL718P | AIDA | | | SEC16B |
| AHNAK2 | AIF1L | | | SMCHD1 |
| NIN | AIFM2 | | | RN7SL4P |
| OCEL1 | AIM2 | | | BTN2A2 |
| CYP2F1 | AIMP1 | | | SNORA74A |
| CCND3 | AIMP2 | | | ARHGEF28 |
| RPS15 | AIP | | | NAPA |
| MUC13 | AJ003147.3 | | | H2BU1 |
| AKAP12 | AJM1 | | | RSRC2 |
| GUSBP1 | AJUBA | | | TRIM56 |
| SPINK5 | AK3 | | | AC107959.3 |
| ZNF76 | AK7 | | | TOP2B |
| PTCHD4 | AKAP1 | | | FDPS |
| CRYBG3 | AKAP10 | | | ASPM |
| ZFP36 | AKAP11 | | | IER5 |
| CARD16 | AKAP12 | | | EGOT |
| DHX35 | AKAP13 | | | CPEB3 |
| SAMD9L | AKAP17A | | | KPNA2 |
| ST3GAL2 | AKAP5 | | | SCARNA6 |
| PPDPF | AKAP8L | | | DUSP16 |
| EYA1 | AKIRIN1 | | | PER1 |
| SP2 | AKIRIN2 | | | GDF15 |
| CCDC40 | AKNA | | | GBP5 |
| IFIH1 | AKR1A1 | | | RARRES1 |
| AMOTL2 | AKR1B1 | | | LBR |
| NUMA1 | AKR1B10 | | | ZNF462 |
| FAM222A | AKR1B15 | | | RN7SL2 |
| CCDC88C | AKR1C2 | | | ZBED5 |
| RND1 | AKR1C3 | | | TAPBP |
| UBA52 | AKR1C7P | | | IDI1 |
| ARHGAP39 | AKR7A2 | | | NCAPG |
| WDR62 | AKT1 | | | SECTM1 |
| AC134407.2 | AKT2 | | | PATL1 |
| SCGB1A1 | AKT3 | | | DAPP1 |
| ATF3 | AL008729.1 | | | CHST9 |
| ZNF500 | AL009174.1 | | | ANXA5 |
| ALPK3 | AL022322.2 | | | HMMR |
| RP11-589F5.3 | AL031058.1 | | | ARHGAP26 |
| TICAM1 | AL031283.1 | | | CCNB1 |
| UBAP2 | AL031595.3 | | | FDFT1 |
| CEP135 | AL031777.1 | | | ERAP1 |
| ZNF329 | AL033397.1 | | | EHD4 |
| CCDC78 | AL033519.3 | | | AC109326.1 |
| C12orf50 | AL034379.1 | | | IRF9 |
| EVI2B | AL034397.3 | | | TAAR3P |
| YY1AP1 | AL035071.1 | | | RND1 |
| TRIM14 | AL035661.1 | | | AHNAK2 |
| NKX3-1 | AL049555.1 | | | AC015688.4 |
| CCDC159 | AL049840.6 | | | RICTOR |
| JPX | AL049873.1 | | | BTN3A2 |
| ZNF335 | AL109806.1 | | | GTPBP1 |

| | | | | |
|---|---|---|---|---|
| MRTFA | AL118516.1 | | | UBE2Z |
| FOXG1 | AL121603.2 | | | DUSP5 |
| CCDC106 | AL121895.1 | | | ZNF274 |
| SYTL5 | AL121944.2 | | | ASAP2 |
| COTL1 | AL121949.1 | | | AP2M1 |
| OAS1 | AL133260.1 | | | CASP1 |
| H2AC6 | AL133335.1 | | | NAV2 |
| AC004151.1 | AL135745.1 | | | UBD |
| MUC21 | AL135818.1 | | | H2AZ2 |
| HOOK1 | AL158050.1 | | | BICC1 |
| H2BC18 | AL158801.6 | | | NOCT |
| OAS3 | AL159141.1 | | | BLZF1 |
| PREX1 | AL161431.1 | | | DPYSL2 |
| CCDC88B | AL161626.1 | | | MDH1 |
| FTL | AL161787.1 | | | CRYZL2P-SEC16B |
| FOSB | AL162151.2 | | | SPRY2 |
| WNK2 | AL162231.2 | | | HERPUD1 |
| MAST3 | AL353804.1 | | | HLA-J |
| EPSTI1 | AL355309.1 | | | DUOX1 |
| PCNT | AL355355.2 | | | FSCN1 |
| NFATC3 | AL356273.6 | | | RPLP0 |
| HIGD2A | AL356489.2 | | | NAMPT |
| KIAA0040 | AL391427.1 | | | XRCC5 |
| C15orf62 | AL445363.3 | | | IDH1 |
| CEP112 | AL445490.1 | | | PPP1R10 |
| ARHGAP26 | AL445524.1 | | | PHACTR4 |
| SGSM1 | AL445931.1 | | | CSRNP1 |
| MVB12B | AL512306.2 | | | SERPINB9 |
| SPAG9 | AL512408.1 | | | ANXA2 |
| SYNPO | AL513314.2 | | | PERP |
| PKD1P5 | AL589743.1 | | | PSME1 |
| SHANK2 | AL590705.3 | | | NEDD1 |
| CLEC16A | AL590867.2 | | | CLUHP3 |
| ELF4 | AL591846.2 | | | H1-0 |
| NCCRP1 | AL592211.1 | | | SH3KBP1 |
| H2AC18 | AL596202.1 | | | CTSD |
| HELB | AL627309.5 | | | RASGEF1B |
| FCGR2A | AL627309.6 | | | KDM2A |
| ATP5F1E | AL669831.1 | | | CYP2J2 |
| SIPA1L3 | AL691432.2 | | | CRYZ |
| H2AC19 | AL691447.2 | | | SOCS1 |
| SEC24A | AL732372.2 | | | UPK1B |
| POLR2M | AL772337.3 | | | GLCCI1 |
| GNG5 | AL807752.1 | | | PLEKHA7 |
| KAT2B | AL807757.2 | | | GNG12 |
| A2ML1 | AL845472.1 | | | TSC22D2 |
| TNFAIP3 | ALAS1 | | | HLTF |
| HSF1 | ALDH1A1 | | | TMEM62 |
| MFN1 | ALDH1A3 | | | JUNB |
| AC126755.1 | ALDH3A1 | | | H2AZ1 |
| ALMS1 | ALDH3B1 | | | SOCS3 |
| FBXO48 | ALDH3B2 | | | AHR |
| RABIF | ALDOA | | | CAT |
| SLC25A23 | ALDOC | | | STK10 |
| MCUB | ALG1L13P | | | TRAM1 |
| SOBP | ALG3 | | | KCNV1 |
| HELZ2 | ALG5 | | | AL356488.2 |
| CBX6 | ALG6 | | | SCARNA2 |
| SRCAP | ALKBH2 | | | NR2F1-AS1 |
| TACC2 | ALKBH5 | | | DSEL |

| | | | | |
|---|---|---|---|---|
| FAM193A | ALKBH6 | | | ACAT2 |
| SERPINB8 | ALKBH7 | | | TET2 |
| PRPF3 | ALOX5AP | | | CFAP57 |
| SYTL2 | ALPK1 | | | ADM |
| MBNL1 | ALPL | | | BACH1 |
| PATL1 | ALS2CL | | | FUT4 |
| ZMIZ2 | ALYREF | | | IARS2 |
| ZNF358 | AMBRA1 | | | GMPR |
| BLZF1 | AMDHD1 | | | TGFBI |
| STK10 | AMER1 | | | GBP1P1 |
| NIBAN1 | AMFR | | | SNORA70 |
| CDK18 | AMIGO2 | | | DND1 |
| PHC3 | AMMECR1L | | | PDZD2 |
| FYCO1 | AMN1 | | | SNORA12 |
| LGALS9B | AMOTL1 | | | GALNT10 |
| TEX9 | AMPD2 | | | CD47 |
| MX1 | AMT | | | ZNF503 |
| FNDC3B | AMTN | | | CD46 |
| CLPB | ANAPC1 | | | AC019117.4 |
| PRRC2A | ANAPC11 | | | IL4I1 |
| APOL3 | ANAPC13 | | | COL11A2 |
| CENPBD1P1 | ANAPC2 | | | IFNB1 |
| RAB8A | ANAPC5 | | | TNFSF13B |
| DAP | ANAPC7 | | | RND3 |
| ZNF592 | ANGEL1 | | | SCARNA13 |
| GTDC1 | ANGPTL2 | | | TRIM14 |
| MOB3A | ANGPTL4 | | | RPS25 |
| TADA2B | ANK3 | | | ZNF200 |
| SH3KBP1 | ANKDD1A | | | CEP350 |
| SAMD4B | ANKDD1B | | | HMGB3 |
| TRAF3IP2 | ANKFY1 | | | VPS41 |
| SPDEF | ANKH | | | ZCCHC2 |
| CDKN2B | ANKLE2 | | | CTSH |
| RBMS2 | ANKMY1 | | | RNU1-67P |
| CIZ1 | ANKMY2 | | | SKIV2L |
| DUSP3 | ANKRA2 | | | EEF1A1P5 |
| RFX5 | ANKRD10 | | | HMGB1 |
| MAP3K11 | ANKRD11 | | | ZC3H12A |
| TENT5C | ANKRD17 | | | PPP1R15B |
| TBC1D15 | ANKRD22 | | | APLP2 |
| RPL36AL | ANKRD27 | | | ARAP2 |
| R3HDM2 | ANKRD29 | | | NEB |
| MLPH | ANKRD36 | | | WTAP |
| S100A11 | ANKRD36B | | | MAFF |
| RND3 | ANKRD37 | | | HACD3 |
| UBB | ANKRD42 | | | RPS3 |
| PAK4 | ANKRD46 | | | CSKMT |
| RASSF9 | ANKRD49 | | | SNORD3B-2 |
| ATXN7 | ANKRD50 | | | HSD17B7P2 |
| RUNDC1 | ANKRD54 | | | CENPF |
| IQCE | ANKS1A | | | SEPTIN2 |
| R3HDM1 | ANKZF1 | | | SLFN5 |
| NEK6 | ANO6 | | | EPC1 |
| JADE2 | ANP32A | | | ANKS1A |
| TCOF1 | ANPEP | | | S100A6 |
| CSNK1G2 | ANXA1 | | | CAPN2 |
| TOB2 | ANXA2 | | | DBI |
| ATN1 | ANXA2R | | | BIRC5 |
| NCOR2 | ANXA3 | | | IFNL2 |
| IL1R1 | ANXA5 | | | ENO1 |

| | | | | |
|---|---|---|---|---|
| SPECC1 | ANXA6 | | | BAZ1A |
| ANKRD17 | ANXA8 | | | ARID4B |
| BICDL1 | AOC3 | | | CPD |
| RAB3B | AOPEP | | | PSMB8-AS1 |
| VPS37B | AP000446.1 | | | PSMB10 |
| CRY2 | AP000487.1 | | | HCG27 |
| ACSS1 | AP000640.2 | | | TAGLN2 |
| WARS1 | AP000769.1 | | | CANX |
| OPTN | AP000844.2 | | | PATL2 |
| ZNF609 | AP000892.4 | | | DNAH17 |
| PDE4DIP | AP000936.3 | | | IQGAP3 |
| FCHSD2 | AP000974.1 | | | KIF4A |
| SP3 | AP001324.1 | | | SLC5A4-AS1 |
| WWC1 | AP001830.1 | | | RBMS2 |
| MLXIP | AP001972.5 | | | ACTB |
| ZFP36L2 | AP002784.2 | | | PRKACB |
| SORBS3 | AP003068.2 | | | MED13 |
| SND1 | AP006621.2 | | | ARL6IP5 |
| FOXK1 | AP006623.1 | | | INTS6 |
| GTPBP1 | AP1B1 | | | GABBR1 |
| CCDC69 | AP1M1 | | | GOLIM4 |
| C6orf132 | AP1M2 | | | RBCK1 |
| LINC00963 | AP1S2 | | | NECTIN2 |
| ADGRG1 | AP2A1 | | | FOSL1 |
| SRSF5 | AP2A2 | | | RNF149 |
| ACTN4 | AP2M1 | | | PPP4R4 |
| EWSR1 | AP3B1 | | | GLO1 |
| HK1 | AP3D1 | | | STING1 |
| TMED2 | AP3M1 | | | FAS |
| CCZ1B | AP3M2 | | | H2AW |
| PSMD2 | AP3S1 | | | FADS1 |
| VAMP3 | AP3S2 | | | RBM39 |
| CTNND1 | AP4M1 | | | CITED2 |
| ARRDC3 | AP5Z1 | | | BNIP3L |
| RDH10 | APBA3 | | | CIITA |
| EIF3CL | APBB1 | | | DIP2B |
| PRKAR1A | APBB1IP | | | MFSD14A |
| PPP2CB | APBB3 | | | SUN2 |
| TSPYL1 | APC | | | CAPS |
| SRP9 | APEH | | | NT5E |
| SMARCA2 | APEX2 | | | VTRNA1-3 |
| GLUL | APH1A | | | STMN1 |
| RBM25 | API5 | | | AL136295.5 |
| TNFRSF21 | APLF | | | TSN |
| SELENBP1 | APLP1 | | | SERPINB10 |
| DHCR24 | APLP2 | | | MAOB |
| MAT2A | APOBEC3A | | | TPX2 |
| EIF3C | APOBEC3C | | | AL021578.1 |
| SCARB2 | APOBEC3G | | | KIAA0040 |
| TMPRSS4 | APOBR | | | GABRP |
| EIF4G1 | APOC1 | | | IFNL3 |
| ENC1 | APOD | | | FOXO1 |
| HNRNPH1 | APOL1 | | | ARL5B |
| WDR1 | APOL3 | | | IFI6 |
| ADSS2 | APOL6 | | | PTTG1IP |
| TMEM123 | APOLD1 | | | VAV3 |
| IL33 | APOO | | | CXCL1 |
| CCZ1 | APP | | | CD2AP |
| FBXL5 | APPL1 | | | CDKN1A |
| ZBTB7A | APRT | | | HLA-DQB1 |

| | | | | |
|---|---|---|---|---|
| HNRNPK | APTR | | | CRIP2 |
| PLEKHB2 | APTX | | | REV3L |
| HNRNPU | AQP1 | | | KCNJ15 |
| ARL8B | AQP3 | | | LRP2 |
| DHX15 | AQP9 | | | CLK1 |
| UBQLN1 | AQR | | | HLA-DOB |
| ANXA7 | ARAP1 | | | PLCG2 |
| NDRG2 | ARAP3 | | | CXXC4 |
| TOMM20 | ARC | | | GTPBP2 |
| GANAB | AREG | | | EEF1G |
| CTBP2 | ARF3 | | | ARHGEF38 |
| CBWD6 | ARF5 | | | MYH10 |
| ITGB1 | ARFGAP1 | | | NCEH1 |
| ASPH | ARFGAP2 | | | PNP |
| ATP8B1 | ARFGAP3 | | | UBC |
| PFN2 | ARFGEF1 | | | AC006064.5 |
| DDB1 | ARFGEF3 | | | ITGB6 |
| ARGLU1 | ARFIP1 | | | SCARNA10 |
| SF3B1 | ARFIP2 | | | SETD5 |
| ANPEP | ARFRP1 | | | PTTG1 |
| DDX1 | ARGLU1 | | | SIPA1L2 |
| NCSTN | ARHGAP1 | | | CYP51A1 |
| DNAJC10 | ARHGAP10 | | | KDM6B |
| DDX5 | ARHGAP15 | | | DDIT3 |
| IFT57 | ARHGAP17 | | | SOD2 |
| PGK1 | ARHGAP21 | | | GPSM2 |
| MIA3 | ARHGAP23 | | | MBD1 |
| SDC4 | ARHGAP26 | | | COL7A1 |
| VPS35 | ARHGAP27P1-BPTFP1-KPNA2P3 | | | DUSP4 |
| GALNT7 | ARHGAP29 | | | IDH2 |
| NAP1L4 | ARHGAP32 | | | EMP2 |
| SRSF10 | ARHGAP35 | | | C1R |
| COPB1 | ARHGAP4 | | | HAS2 |
| DDX17 | ARHGAP42 | | | CD40 |
| RBM39 | ARHGAP44 | | | TNFRSF10A |
| TAX1BP1 | ARHGAP9 | | | AC002480.1 |
| ANKRD12 | ARHGDIA | | | EXT1 |
| LMBRD1 | ARHGDIB | | | PARP8 |
| CALM2 | ARHGEF1 | | | AC000120.3 |
| SLC37A3 | ARHGEF10 | | | AP002990.1 |
| RDX | ARHGEF10L | | | DHFR |
| ABCD3 | ARHGEF16 | | | HLA-V |
| CKAP4 | ARHGEF17 | | | AMOTL2 |
| TMEM41B | ARHGEF25 | | | DEPDC1 |
| NFE2L1 | ARHGEF3 | | | SC5D |
| EPRS1 | ARHGEF34P | | | ITPKC |
| UBE2Q1 | ARHGEF37 | | | NFKB1 |
| ERBB3 | ARHGEF5 | | | CASP8 |
| DIMT1 | ARHGEF7 | | | VTCN1 |
| RAC1 | ARHGEF9 | | | KSR1 |
| CALM1 | ARID2 | | | CDK12 |
| PRSS23 | ARID3A | | | RAB24 |
| SFPQ | ARID3B | | | AURKA |
| EXOC3 | ARID5A | | | AC068580.4 |
| ADAM9 | ARIH2 | | | TUBA1A |
| XPO1 | ARL10 | | | RTN3 |
| ACSL5 | ARL13B | | | ITGAV |
| CCT2 | ARL14EP | | | RPL3 |
| KDELR1 | ARL2 | | | AEN |
| PCMTD2 | ARL3 | | | TTC28 |

| | | | | |
|---|---|---|---|---|
| ACTR2 | ARL5B | | | IL10RA |
| ALDH3A2 | ARL8A | | | PIK3R1 |
| RTF1 | ARMC10 | | | POLD2 |
| SH3YL1 | ARMC5 | | | TNFRSF9 |
| ATP6V1A | ARMC6 | | | ANLN |
| CBX3 | ARMC8 | | | AP5Z1 |
| MAP2K2 | ARMC9 | | | NOL6 |
| CLSTN1 | ARMCX4 | | | RYR2 |
| ZDHHC13 | ARMCX5-GPRASP2 | | | RBM5 |
| LAPTM4A | ARMCX7P | | | PTPRF |
| ANXA2 | ARMH3 | | | SGO2 |
| SEPTIN2 | ARNT2 | | | NEK2 |
| SLC6A6 | ARPC1B | | | SHROOM2 |
| CLTC | ARPC2 | | | ZNF263 |
| STIM2 | ARPC3 | | | GBP2 |
| RCBTB1 | ARPC5 | | | ANKRD33B |
| PSMD14 | ARPC5L | | | EID1 |
| PCYOX1 | ARPP19 | | | PSAP |
| SACM1L | ARRB1 | | | IQGAP1 |
| ST14 | ARRB2 | | | DUSP10 |
| ACVR1B | ARRDC1 | | | LAD1 |
| SEC61A1 | ARRDC2 | | | SFT2D2 |
| SYNGR2 | ARRDC3 | | | SERBP1 |
| IL13RA1 | ARRDC4 | | | PALMD |
| CAPZA2 | ARSB | | | HSD17B14 |
| XPO7 | ARSD | | | TBL1XR1 |
| CANX | ARSJ | | | GPCPD1 |
| GALNT1 | ARSL | | | GJB1 |
| SDC1 | ARX | | | HBEGF |
| MFSD14B | ASAP3 | | | HACD2 |
| DDX18 | ASB1 | | | ZNF385C |
| CCNDBP1 | ASB13 | | | ZC3H12C |
| ADGRF1 | ASB16 | | | USF1 |
| UBE2N | ASB6 | | | PCGF5 |
| MDFIC | ASB7 | | | MAGT1 |
| SSR1 | ASB8 | | | SNHG12 |
| CNOT8 | ASCC2 | | | FN1 |
| SDCBP | ASCC3 | | | ASAH1 |
| RAB5C | ASF1A | | | TP53BP2 |
| ERP44 | ASL | | | MDK |
| MPZL2 | ASMTL | | | LPP |
| HMGXB3 | ASPSCR1 | | | PFKFB3 |
| RSRC2 | ASS1 | | | AC118553.2 |
| KPNA1 | ASXL1 | | | CD55 |
| HSPA9 | ASXL2 | | | SEMA3D |
| AKR1C3 | ATAD3A | | | IFI44L |
| ALCAM | ATAD3B | | | LINC02701 |
| CTNNB1 | ATF1 | | | LIPA |
| GAPDH | ATF3 | | | NNT |
| GSN | ATF4 | | | ANP32E |
| TFCP2L1 | ATF6B | | | CIT |
| PDCD4 | ATF7 | | | ACAA2 |
| DYNLT1 | ATG101 | | | JMJD1C |
| QSOX1 | ATG16L1 | | | SNPH |
| GOLGA1 | ATG16L2 | | | CD63 |
| ESYT2 | ATG2A | | | SARAF |
| HSP90AA1 | ATG2B | | | ODAM |
| RBBP9 | ATG4A | | | DARS2 |
| AAGAB | ATG4B | | | MAP4 |
| CD46 | ATG4C | | | MED1 |

| | | | | |
|---|---|---|---|---|
| TAP1 | ATG9A | | | SERINC5 |
| HIGD1A | ATN1 | | | SUCO |
| CLDN7 | ATOX1 | | | CXCL3 |
| GM2A | ATP10A | | | PA2G4P4 |
| KIF21A | ATP11A | | | PCLO |
| MARK3 | ATP12A | | | FA2H |
| PAICS | ATP13A1 | | | GALNT1 |
| NFKB1 | ATP13A2 | | | DUSP8 |
| HNRNPA2B1 | ATP1A1 | | | BCL2L14 |
| EIF5B | ATP1A3 | | | H3-3B |
| WEE1 | ATP2A3 | | | PPP2R3A |
| CDC42SE2 | ATP2B1 | | | LMAN1 |
| USP48 | ATP2B1-AS1 | | | PSMA6 |
| ANKRD36B | ATP2B4 | | | NKX3-1 |
| HNRNPL | ATP2C2 | | | DEPP1 |
| RCAN3 | ATP5F1A | | | AHCYL2 |
| ALDH3A1 | ATP5F1B | | | GTF2B |
| SRPRA | ATP5F1D | | | SNIP1 |
| FMO3 | ATP5F1E | | | MORF4L1 |
| CLCN3 | ATP5IF1 | | | ZNF697 |
| UGP2 | ATP5MD | | | SREBF2 |
| NIPAL2 | ATP5MF | | | PDXK |
| NCKAP1 | ATP5MPL | | | H2BC14 |
| RAB5A | ATP5PB | | | RPL7 |
| SLC38A2 | ATP5PD | | | HES4 |
| EIF1AX | ATP5PF | | | LSS |
| DECR1 | ATP5PO | | | HLA-DRB1 |
| SLC25A3 | ATP6V0A1 | | | GPT2 |
| SLC44A2 | ATP6V0A2 | | | ARHGAP11A |
| CP | ATP6V0A4 | | | PRR15 |
| PSAP | ATP6V0D1 | | | SNORA68 |
| GRN | ATP6V0E1 | | | IGFBP4 |
| SLC20A1 | ATP6V1B2 | | | MKI67 |
| MARCHF5 | ATP6V1C1 | | | SEMA3E |
| IGFBP3 | ATP6V1F | | | GPI |
| SON | ATP6V1G1 | | | PLCH1 |
| GOT1 | ATP6V1H | | | ABCD3 |
| SLC15A2 | ATP7A | | | ERBB2 |
| ACADM | ATP8B2 | | | HK2 |
| SLC30A7 | ATRN | | | STC2 |
| CCT5 | ATRX | | | ZFP36L2 |
| SORT1 | ATXN10 | | | KRT19 |
| TMEM87B | ATXN1L | | | PPP1CB |
| VAV1 | ATXN2 | | | BAG1 |
| PLRG1 | ATXN2L | | | IER2 |
| FBP1 | ATXN3 | | | CELSR3 |
| EIF2S3 | ATXN7L3 | | | MASTL |
| IARS2 | AUNIP | | | PDGFRL |
| TMBIM6 | AUP1 | | | RPL27A |
| MAOA | AURKAIP1 | | | LIFR |
| HSPA5 | AURKB | | | NOD1 |
| B4GALT5 | AUTS2 | | | NT5DC2 |
| COX6C | AXIN1 | | | PAM |
| CHL1 | AXIN2 | | | C1S |
| MFSD1 | AXL | | | PCYOX1 |
| LINC01578 | AZIN1 | | | CLDN10 |
| BUB3 | B2M | | | H1-10 |
| TRA2A | B3GAT3 | | | STON2 |
| ATP5PB | B3GNT2 | | | CD68 |
| POLR2B | B3GNT5 | | | ATP1B1 |

| | | | | |
|---|---|---|---|---|
| ACSL3 | B3GNT6 | | | ADAM9 |
| SRSF3 | B4GALT2 | | | SLIT2 |
| DHX29 | B4GALT3 | | | RPS6 |
| RAB1A | B4GALT7 | | | IGF2 |
| SCNN1A | BAALC | | | MYO1C |
| WASHC2C | BACE1 | | | RFX5 |
| IMPAD1 | BACE2 | | | NAMPTP1 |
| GALNT12 | BAG1 | | | MNS1 |
| LAMP2 | BAG3 | | | AFDN |
| ATXN10 | BAG6 | | | ABTB2 |
| SPTLC1 | BAHD1 | | | BUB1B |
| FDX1 | BAIAP2 | | | NIBAN2 |
| TM4SF1 | BAIAP2L1 | | | IL7R |
| TMEM50B | BAK1 | | | LMO4 |
| ATP5F1A | BANF1 | | | AC083837.1 |
| RBM6 | BANP | | | SRI |
| B2M | BAP1 | | | RN7SL1 |
| NPC2 | BARD1 | | | SUMF1 |
| PERP | BASP1 | | | COL16A1 |
| ELOC | BATF2 | | | TSPAN6 |
| ANXA3 | BAX | | | LDLR |
| SLBP | BAZ1B | | | DENND4A |
| TMED10 | BAZ2A | | | TMA7 |
| CPA4 | BAZ2B | | | HNRNPR |
| EFTUD2 | BBOF1 | | | FAM53C |
| LDHB | BBS1 | | | NCMAP |
| MMUT | BBS12 | | | OR52K3P |
| EEF1A1P6 | BBS2 | | | PSMA4 |
| SRI | BBS4 | | | MIR3609 |
| LINC00511 | BCAP29 | | | PRC1 |
| SSB | BCAP31 | | | INSIG1 |
| SKP1 | BCAR1 | | | USP5 |
| AKIRIN1 | BCAR3 | | | LINC02542 |
| DERL1 | BCAS1 | | | ZNF620 |
| ATRX | BCAT1 | | | AC132217.2 |
| UNC5B | BCAT2 | | | IRAK1 |
| TMEM30B | BCDIN3D-AS1 | | | ICAM1 |
| SLC44A1 | BCKDK | | | SLC25A28 |
| LTA4H | BCL11A | | | HELB |
| SEPHS2 | BCL2A1 | | | MAP3K20 |
| ACTG1 | BCL2L1 | | | VDAC1 |
| IRF2BPL | BCL2L13 | | | HLA-DMB |
| AQP5 | BCL2L14 | | | TGOLN2 |
| NDUFS8 | BCL2L2 | | | PBXIP1 |
| STT3B | BCL3 | | | CDCA3 |
| A4GALT | BCL6 | | | SNHG1 |
| TUBA1A | BCL7A | | | CHAC1 |
| COPS8 | BCL7B | | | OVOL1 |
| NDUFA10 | BCL7C | | | AIF1L |
| PRXL2A | BCL9 | | | ARRDC4 |
| GDE1 | BCL9L | | | AC022034.1 |
| DARS1 | BCLAF3 | | | ARC |
| ANXA1 | BCOR | | | VCL |
| NUCB2 | BCR | | | AL109918.1 |
| ID1 | BCS1L | | | ANXA11 |
| FMO2 | BDKRB2 | | | ARSD |
| COG2 | BECN1 | | | VPS35 |
| PSMC5 | BEND7 | | | LMNB1 |
| RSRP1 | BEST1 | | | LYZ |
| NDFIP1 | BET1L | | | C15orf48 |

| | | | | |
|---|---|---|---|---|
| B4GALT1 | BEX1 | | | AC107959.1 |
| RPL6P27 | BEX4 | | | CPEB4 |
| KIFAP3 | BHLHE40 | | | AKR1C3 |
| F3 | BICD1 | | | FGFR3 |
| KARS1 | BICD2 | | | PNRC1 |
| OSTC | BICRAL | | | GRINA |
| IDH1 | BID | | | ATP5F1A |
| JAG1 | BIN1 | | | RGMB |
| GLT8D1 | BIN2 | | | SNORA71A |
| KYNU | BIN3 | | | AFF4 |
| CMTM6 | BIRC5 | | | HSPB8 |
| SDR16C5 | BIRC6 | | | SAR1A |
| EMB | BLCAP | | | SH3BGRL2 |
| CCDC47 | BLMH | | | RNF128 |
| PROM1 | BLOC1S1 | | | AC025580.2 |
| APP | BLVRB | | | GCNT4 |
| MIPEP | BLZF1 | | | CTDSPL |
| C2CD2 | BMERB1 | | | PAXIP1-AS2 |
| TMEM33 | BMP1 | | | AC018644.1 |
| CD164 | BMP2 | | | AC106886.6 |
| MDH1 | BMPR1AP2 | | | DDB1 |
| TFCP2 | BMS1 | | | AGO1 |
| TMPRSS11D | BMS1P1 | | | PLEKHB1 |
| NR2F2 | BMS1P17 | | | ANXA9 |
| VPS26A | BMS1P22 | | | IKBKE |
| CAST | BMS1P8 | | | AHCY |
| GSTK1 | BNIP1 | | | ALDH7A1 |
| YWHAB | BNIP3P1 | | | PRDX5 |
| TXNDC17 | BNIP3P11 | | | MYC |
| CCN2 | BOD1 | | | OAZ1 |
| CKMT1A | BOLA3 | | | DAG1 |
| PSMD11 | BRAF | | | GNB4 |
| PKM | BRAT1 | | | IFRD1 |
| METTL21A | BRCA2 | | | LARS1 |
| LAPTM4B | BRD1 | | | ERRFI1 |
| PMPCB | BRD2 | | | MARCKS |
| PRNP | BRD3 | | | HRH1 |
| SEC63 | BRD3OS | | | C4A |
| TMEM165 | BRD8 | | | PODXL |
| CAV2 | BRD9 | | | EZR |
| PYGL | BRI3 | | | NCOA3 |
| ARPC5 | BRK1 | | | ROR1 |
| TMEM150C | BRMS1 | | | SMARCC1 |
| CD63 | BRWD1 | | | GPAT4 |
| PLS3 | BRWD3 | | | PTPRS |
| HLF | BSDC1 | | | PRR11 |
| AC087473.1 | BST1 | | | CH25H |
| ATP6V0D1 | BTBD1 | | | ITGAM |
| PSMB3 | BTBD19 | | | PTMA |
| NOMO1 | BTBD3 | | | SQSTM1 |
| GTF2E2 | BTBD7 | | | SAMD4A |
| OAT | BTF3L4 | | | INO80D |
| CD151 | BTF3L4P2 | | | TRIO |
| ALDH2 | BTG2 | | | HLA-DPA1 |
| FBXO3 | BTN2A2 | | | SLC44A4 |
| MORN2 | BTN3A2 | | | CTNNA1 |
| CTBS | BTNL9 | | | RTL9 |
| STOM | BUB3 | | | MBNL3 |
| RNF149 | BUD13 | | | SLC9A7 |
| MAP3K5 | BUD23 | | | FTL |

| | | | | |
|---|---|---|---|---|
| ERLIN1 | BX248409.1 | | | SLC25A25 |
| ERMP1 | BX284668.2 | | | FOSL2 |
| SMIM15 | BX679664.1 | | | RPS4X |
| CCDC80 | BX679664.3 | | | IL12A |
| HSP90B1 | BYSL | | | SNRPD2 |
| CKMT1B | BZW1P2 | | | RBL2 |
| TSPAN3 | C10orf91 | | | PDGFA |
| STAM2 | C11orf1 | | | CNN3 |
| ABHD5 | C11orf24 | | | KIAA1217 |
| MSMO1 | C11orf49 | | | TCN1 |
| RBM5 | C11orf71 | | | ARID5B |
| TM9SF3 | C11orf86 | | | CHD8 |
| ATP1B1 | C11orf95 | | | MPC2 |
| SARAF | C11orf96 | | | CASP7 |
| AMFR | C12orf10 | | | B3GNT5 |
| DYNC2LI1 | C12orf43 | | | CHEK2 |
| H3-3A | C12orf49 | | | Z93241.1 |
| RNF145 | C12orf57 | | | AC005515.1 |
| ERG28 | C12orf75 | | | SLC26A2 |
| TUBA1C | C14orf132 | | | ZNF217 |
| RBM3 | C15orf39 | | | AL162581.1 |
| GLB1 | C15orf48 | | | RN7SK |
| PGD | C16orf58 | | | SNORA49 |
| USP47 | C16orf91 | | | HLA-K |
| TAGLN2 | C17orf58 | | | PIP5K1A |
| RETREG2 | C17orf80 | | | SYNPO |
| TMX4 | C17orf97 | | | NUSAP1 |
| EXOC1 | C18orf21 | | | MSH6 |
| MAT2B | C19orf25 | | | TMEM59 |
| VPS25 | C19orf48 | | | SPAG5 |
| NUP107 | C19orf84 | | | DPYSL3 |
| ADAM15 | C1GALT1 | | | STT3B |
| ARPC3 | C1orf109 | | | H2BC11 |
| SYPL1 | C1orf112 | | | PFN2 |
| TGFBR1 | C1orf116 | | | HTATSF1 |
| DSG2 | C1orf162 | | | UQCRH |
| RPL4P4 | C1orf194 | | | PDGFD |
| SELENOP | C1orf198 | | | CLINT1 |
| EPCAM | C1orf21 | | | AL121594.1 |
| S100A6 | C1orf216 | | | PDE1C |
| COG6 | C1QBP | | | SNORD10 |
| ANKRD10 | C1QL4 | | | CUX1 |
| WDR33 | C1QTNF1 | | | ZNF787 |
| LRRC8D | C1QTNF6 | | | DHRS2 |
| MMP14 | C1S | | | SCARNA9 |
| RWDD4 | C2 | | | OPHN1 |
| SDF4 | C20orf194 | | | ITPRIPL2 |
| DLD | C20orf27 | | | YWHAQ |
| FHL2 | C20orf96 | | | NXF1 |
| SLC35F5 | C21orf91 | | | BMP2 |
| TACSTD2 | C22orf39 | | | INPP4B |
| ASAH1 | C22orf46 | | | ARPP19 |
| GUF1 | C2CD2 | | | DNAJA4 |
| ENO1 | C2CD4A | | | DNAJC22 |
| LUC7L3 | C2orf42 | | | SCP2 |
| MTDH | C3 | | | SNAI1 |
| JKAMP | C3AR1 | | | KLF10 |
| H2AZ2 | C3orf38 | | | CPEB2 |
| ERAP1 | C3orf62 | | | LINC00472 |
| TMEM192 | C4A | | | PGRMC1 |

| | | | | |
|---|---|---|---|---|
| RCN1 | C4B | | | OTUD4 |
| WASH6P | C4orf3 | | | MAP3K13 |
| ITGB5 | C4orf33 | | | SCARNA12 |
| SLC35A2 | C5AR1 | | | PLS3 |
| TTC19 | C5AR2 | | | APOBEC3F |
| OCIAD1 | C5orf30 | | | STX19 |
| ADAM28 | C5orf51 | | | APOL4 |
| P4HB | C6orf120 | | | USP43 |
| SFN | C6orf132 | | | C3orf52 |
| SLC39A7 | C6orf136 | | | ROS1 |
| EIF4A3 | C6orf47 | | | TPM4 |
| TMEM30A | C6orf62 | | | DDX39B |
| GPD2 | C7orf25 | | | ZNF441 |
| CD59 | C7orf26 | | | HOMER2 |
| GTF2H2B | C7orf50 | | | RNF111 |
| PON2 | C8orf33 | | | HLA-L |
| HACD3 | C8orf34 | | | HSD17B4 |
| GSS | C8orf37 | | | NCALD |
| TUFM | C8orf44 | | | PLK3 |
| ATP2C1 | C8orf58 | | | SLC39A10 |
| CACHD1 | C8orf82 | | | PTP4A2 |
| CA12 | C9orf116 | | | PHF11 |
| MBOAT2 | C9orf64 | | | PIWIL4 |
| THYN1 | C9orf72 | | | ADAM8 |
| PDIA6 | CA1 | | | NFIA |
| LEMD3 | CA12 | | | RPS3A |
| RPN1 | CAB39 | | | ITGB5 |
| PTTG1IP | CAB39L | | | CDK1 |
| PRDX1 | CABIN1 | | | TMSB4X |
| GAA | CABLES1 | | | CYP2B6 |
| ABCC3 | CACNA1A | | | AGRN |
| IER3IP1 | CACNA1C | | | ERLIN2 |
| EGFR | CACNG6 | | | AC027290.2 |
| ANXA4 | CACNG7 | | | ATP13A3 |
| IGFBP2 | CACTIN | | | CHMP3 |
| SLC27A2 | CACUL1 | | | GSR |
| TRAPPC3 | CAD | | | MOB3C |
| OS9 | CADM1 | | | RPL41 |
| SLC18B1 | CADM4 | | | DNPEP |
| ALOX15 | CADPS2 | | | SBNO2 |
| GNS | CALCOCO1 | | | MMAB |
| RRN3 | CALCOCO2 | | | MSI2 |
| PIGX | CALHM2 | | | C15orf39 |
| ARL6IP1 | CALHM6 | | | PRKAR2A |
| RPL7AP66 | CALML3-AS1 | | | TTC26 |
| AC004069.1 | CALR | | | CD83 |
| LGR4 | CAMKK1 | | | ESYT1 |
| LRG1 | CAMKK2 | | | AC245033.1 |
| SPARCL1 | CAMP | | | AC073548.1 |
| TMEM87A | CAMSAP1 | | | MAP2 |
| ALDH1A1 | CAMSAP2 | | | TRA2A |
| LMAN2 | CAMTA1 | | | HS2ST1 |
| AC007318.1 | CAMTA2 | | | PAICS |
| DDOST | CANT1 | | | BTN2A1 |
| CHPF | CANX | | | HNMT |
| CTSB | CAP1 | | | PTPN11 |
| RASA1 | CAPN10 | | | EPS8 |
| SLC2A1 | CAPN8 | | | DEPDC1B |
| CFH | CAPRIN2 | | | HDGF |
| TM9SF2 | CAPZA1 | | | GSN |

| | | | | |
|---|---|---|---|---|
| EML3 | CAPZB | | | UGGT1 |
| CTSH | CARD16 | | | CPSF1P1 |
| HNRNPA1P4 | CARD17 | | | SRFBP1 |
| ERLIN2 | CARD6 | | | SNORA21 |
| MTRNR2L9 | CARD8 | | | ZNF627 |
| LRP5 | CARHSP1 | | | AKAP8L |
| SEMA3A | CARS2 | | | ZNF292 |
| GFM2 | CASC3 | | | RCN2 |
| HNRNPA1L2 | CASC7 | | | JPT2 |
| PIGG | CASC9 | | | AC007952.4 |
| RRM1 | CASKIN1 | | | ACHE |
| TSPAN13 | CASP1 | | | ZNF704 |
| TMEM68 | CASP10 | | | SAPCD2 |
| KRT5 | CASP4 | | | RPS8 |
| ACVR1 | CASP5 | | | LAMA2 |
| ATP1A1 | CASP7 | | | MBTPS1 |
| PRODH | CASP8 | | | SRCAP |
| RPAP2 | CASP9 | | | ZNF317 |
| MFSD11 | CASTOR3 | | | MLF1 |
| FDFT1 | CAT | | | PIEZO1 |
| MFSD14C | CAV1 | | | MRPL3 |
| ANKRD36 | CAV2 | | | ARPC5 |
| SGK1 | CAVIN1 | | | MYLK |
| NEU1 | CAVIN3 | | | NR4A1 |
| CLDN4 | CBFA2T2 | | | CTH |
| SCGB2A1 | CBFA2T3 | | | AC093010.3 |
| ERLEC1 | CBL | | | ZNF442 |
| EEF1A1P5 | CBLB | | | ZBTB21 |
| AL391121.1 | CBLN3 | | | SYDE1 |
| POR | CBR1 | | | ECH1 |
| RAB3IP | CBWD5 | | | JADE2 |
| FAM3B | CBX4 | | | HMGCS1 |
| CALR | CBX6 | | | ODF3B |
| AC073333.1 | CBX7 | | | NR2F1 |
| EBPL | CBY1 | | | SPAG17 |
| NPTN | CC2D1A | | | H2BC18 |
| HSD17B13 | CC2D1B | | | HADHA |
| RPS21 | CCAR1 | | | ASNS |
| NIFK | CCAR2 | | | STARD5 |
| ALS2CL | CCDC102B | | | ZFYVE26 |
| PSEN2 | CCDC113 | | | RPL5 |
| RCN2 | CCDC114 | | | JUND |
| CHKA | CCDC115 | | | GIGYF2 |
| GTF2H2C | CCDC12 | | | CASP4 |
| TMA7 | CCDC124 | | | HIPK3 |
| ALDH7A1 | CCDC130 | | | TBC1D22B |
| CD81 | CCDC137 | | | FLRT3 |
| RTN4 | CCDC14 | | | IFT122 |
| DHX36 | CCDC148 | | | CHSY3 |
| SNRPA1 | CCDC151 | | | EBP |
| DNAJC1 | CCDC184 | | | LUCAT1 |
| HNRNPA1P35 | CCDC186 | | | ACTN2 |
| FTH1P10 | CCDC22 | | | AL358075.4 |
| HNRNPAB | CCDC3 | | | ARHGAP24 |
| FUCA2 | CCDC30 | | | SPC25 |
| ADK | CCDC38 | | | ABLIM1 |
| FMO5 | CCDC43 | | | SYNPO2 |
| RTN3 | CCDC57 | | | DCLRE1C |
| ASCC3 | CCDC59 | | | CLK4 |
| PSME1 | CCDC60 | | | LGALS3 |

| | | | | |
|---|---|---|---|---|
| WASH4P | CCDC66 | | | IFNL4 |
| TFB2M | CCDC68 | | | CYFIP2 |
| REEP5 | CCDC69 | | | TMED10 |
| TMEM51 | CCDC71 | | | PAPOLA |
| WSB1 | CCDC71L | | | ITGA6 |
| SREK1 | CCDC74A | | | MTF2 |
| RTN3P1 | CCDC84 | | | IRF2 |
| RARS1 | CCDC85C | | | ETFB |
| HADH | CCDC86 | | | RRM1 |
| MBOAT1 | CCDC88A | | | ATP5PB |
| PGAP4 | CCDC9 | | | C4B |
| FAAH2 | CCHCR1 | | | AC097625.2 |
| SMG1P4 | CCK | | | BDP1 |
| AC008810.1 | CCL2 | | | SPATS2L |
| TLR2 | CCL3L3 | | | TMEM41B |
| ZMPSTE24 | CCL4 | | | IFT80 |
| HNRNPA1P10 | CCL7 | | | CDKN2B |
| PAPSS2 | CCL8 | | | PLA2G4C |
| AGL | CCM2 | | | CREBRF |
| TMEM9 | CCN2 | | | RPS24 |
| TMEM9B | CCN5 | | | MAFG |
| TSPAN1 | CCNA1 | | | ERMP1 |
| CLN5 | CCNB1 | | | HEXD |
| TMEM59 | CCND1 | | | EGR2 |
| AL592114.1 | CCND3 | | | TEX15 |
| GUSB | CCNDBP1 | | | AVPI1 |
| ANAPC4 | CCNG2 | | | GCH1 |
| COMMD7 | CCNI | | | ALDH1A3 |
| MAP3K6 | CCNJ | | | RACGAP1 |
| ANKRD66 | CCNJL | | | DDX17 |
| DSE | CCNL1 | | | GPR160 |
| GCLC | CCNL2 | | | RUBCN |
| PRSS8 | CCNT1 | | | AC068831.8 |
| CYP4F12 | CCNT2 | | | SPTLC2 |
| ABCE1 | CCNT2-AS1 | | | NDUFV1 |
| ENPP4 | CCPG1 | | | DDX5 |
| ABCA5 | CCR1 | | | KIF14 |
| GLUD2 | CCR5AS | | | CCDC85C |
| CALM2P2 | CCR7 | | | FGFR4 |
| KLHL42 | CCRL2 | | | ARPC2 |
| PRMT7 | CCS | | | KIF23 |
| ITGB6 | CCT4 | | | PRXL2A |
| SESN1 | CCT5 | | | ST13 |
| FTH1P2 | CCT7 | | | C1orf116 |
| CAMK2G | CCT8 | | | RPL6 |
| STAM | CCZ1 | | | OXCT1 |
| MANF | CCZ1B | | | PCDH9 |
| THOC3 | CD151 | | | RAB11A |
| MSTO1 | CD164 | | | CASP10 |
| ZPR1 | CD24 | | | XRCC6 |
| INPP1 | CD248 | | | USP13 |
| NOMO2 | CD27-AS1 | | | ITM2A |
| LRRN1 | CD2BP2 | | | ATP6AP2 |
| SRSF11 | CD320 | | | MDGA1 |
| SELENOI | CD37 | | | AC040162.1 |
| TOPORS | CD38 | | | ENAH |
| PLTP | CD4 | | | TNF |
| SLC39A6 | CD47 | | | RALBP1 |
| TMEM106C | CD53 | | | KLK6 |
| CD47 | CD55 | | | AC099489.1 |

| | | | | |
|---|---|---|---|---|
| SORD2P | CD59 | | | CDC14A |
| AL158206.1 | CD63 | | | SLC38A1 |
| KRT10 | CD74 | | | TANK |
| EIF4A1P4 | CD82 | | | EIF4EBP2 |
| UNC93B3 | CD86 | | | NPLOC4 |
| CXADR | CD8A | | | G0S2 |
| CUEDC1 | CD9 | | | KIF15 |
| IL10RB | CD93 | | | FRYL |
| YWHAZP4 | CD99L2 | | | PCSK9 |
| ATP5F1B | CDA | | | RBBP6 |
| PRKAR2B | CDADC1 | | | SLC16A7 |
| SMC4 | CDC14A | | | BTF3 |
| PTDSS1 | CDC16 | | | AC005070.3 |
| COPG2 | CDC25B | | | PJA2 |
| GTF2F2 | CDC27 | | | PPTC7 |
| LONRF1 | CDC34 | | | LINC02432 |
| RPL10P9 | CDC37 | | | TMEM167A |
| HLA-B | CDC40 | | | CARHSP1 |
| RFNG | CDC42 | | | YEATS2 |
| MAN1B1 | CDC42BPA | | | ACTN1 |
| EIF4HP1 | CDC42BPB | | | RPS20 |
| H3P36 | CDC42EP1 | | | ELF1 |
| LACTB | CDC42EP2 | | | TRAF4 |
| CNIH1 | CDC42EP4 | | | NAALADL2 |
| MAPKAPK3 | CDC7 | | | SNORA71D |
| JPT1 | CDCA2 | | | FAM168A |
| SMAP1 | CDCA7 | | | YRDC |
| CYP3A5 | CDH11 | | | ZBTB10 |
| SPTSSA | CDH17 | | | ARID3B |
| TTC29 | CDH2 | | | SAA2 |
| C1D | CDH23 | | | FAM168B |
| PA2G4P6 | CDH5 | | | ZBTB43 |
| DPAGT1 | CDHR4 | | | CYB5A |
| SMARCAD1 | CDHR5 | | | PLBD1 |
| RPL7AP6 | CDIP1 | | | CORO1C |
| BX679664.3 | CDK10 | | | RNVU1-31 |
| PLS1 | CDK11B | | | CERK |
| F11R | CDK13 | | | AC022506.1 |
| RMDN3 | CDK17 | | | PLK1 |
| DDAH2 | CDK18 | | | IFI30 |
| BCAP29 | CDK4 | | | COX7C |
| FSCN1 | CDK5RAP1 | | | MVD |
| AKAP1 | CDK5RAP3 | | | UHMK1 |
| ITFG1 | CDK6 | | | PLEC |
| ARVCF | CDK7 | | | NPFFR2 |
| LOXL4 | CDK9 | | | RPS17 |
| AC090498.1 | CDKAL1 | | | HDX |
| FAM3D | CDKL3 | | | CWC25 |
| HNRNPA1P7 | CDKN1A | | | PVR |
| HNRNPA1P12 | CDKN1B | | | MGAM2 |
| RHBDL2 | CDKN2AIP | | | MAZ |
| MPP7 | CDKN2B | | | ZNF7 |
| UBE2E3 | CDR1 | | | MAST4 |
| SCPEP1 | CDR2L | | | EREG |
| DNAJB11 | CDV3 | | | ARPIN |
| SDHA | CEACAM1 | | | FNBP4 |
| LAMB3 | CEACAM3 | | | HMGA1 |
| VARS1 | CEACAM6 | | | ARL14EPL |
| NT5DC1 | CELF1 | | | FLNA |
| PBXIP1 | CELSR1 | | | SNORA57 |

| | | | | |
|---|---|---|---|---|
| TAP2 | CEMIP2 | | | TKT |
| HLA-F | CENPB | | | EEF1B2 |
| CDH1 | CENPC | | | TOR1B |
| PRELID1 | CENPN | | | ATP5F1C |
| UPK1B | CENPT | | | ZNF92 |
| PSMD1 | CENPX | | | RBBP7 |
| TMTC4 | CEP120 | | | ANGPT2 |
| ITGAV | CEP164 | | | NIPAL3 |
| NUDT12 | CEP170 | | | SNORA74D |
| PGAM4 | CEP19 | | | LDHB |
| RNF26 | CEP192 | | | CAPZB |
| CCDC65 | CEP250 | | | HMGN3 |
| FARSB | CEP295 | | | MTHFD2 |
| PARP2 | CEP63 | | | OLFM4 |
| VIPR1 | CEP68 | | | COPA |
| ARL3 | CEP83 | | | IRS2 |
| ENY2 | CEP85L | | | SLC28A3 |
| SDCBPP3 | CEP95 | | | SOX4 |
| BCAP31 | CERK | | | ERGIC1 |
| B4GALT4 | CERS2 | | | HDAC1 |
| RAMAC | CERS3 | | | SUPT6H |
| C5orf15 | CERS5 | | | CBX3 |
| FTH1P8 | CERS6 | | | HSCB |
| MAD2L1BP | CERT1 | | | MTF1 |
| RAC1P2 | CFAP161 | | | CDKN3 |
| NAAA | CFAP20 | | | CCND3 |
| GLULP4 | CFAP300 | | | ANP32A |
| EXOSC8 | CFAP36 | | | ARHGAP27 |
| MT-TY | CFAP52 | | | ANGPTL4 |
| BLOC1S4 | CFAP58-DT | | | ACP6 |
| AC144530.1 | CFAP77 | | | SNORA48 |
| TMEM45B | CFAP97 | | | ATF4 |
| ABHD3 | CFDP1 | | | SLC3A1 |
| HLA-L | CFL1 | | | RNU4-2 |
| AC113935.1 | CFLAR | | | ICMT |
| PRPF39 | CGB2 | | | ATP5PD |
| CTNNAL1 | CGB5 | | | CCNG1 |
| VPS35P1 | CGN | | | AC020916.1 |
| HSP90AB2P | CGRRF1 | | | NDUFB6 |
| UNC93B7 | CHAC1 | | | RACK1 |
| CHPT1 | CHCHD1 | | | CHP1 |
| TNC | CHCHD10 | | | TUBA1B |
| DDX50 | CHCHD2 | | | PARK7 |
| MLYCD | CHCHD3 | | | SLC1A5 |
| AQP3 | CHCHD4 | | | TRIB1 |
| LRRCC1 | CHCHD6 | | | HGSNAT |
| TPBG | CHCHD7 | | | DNM1 |
| GJB3 | CHD1 | | | CYP1B1 |
| PABPC1P4 | CHD2 | | | SYT16 |
| SLC25A17 | CHD3 | | | PRKCI |
| SLC44A3 | CHD4 | | | AL049629.2 |
| SLC26A4 | CHD6 | | | TRHDE |
| PRDX4 | CHD7 | | | AP2B1 |
| GGH | CHD8 | | | RPS9 |
| HNRNPA1P8 | CHFR | | | TJAP1 |
| LSAMP | CHI3L1 | | | CPXM1 |
| CDS2 | CHI3L2 | | | WDR74 |
| FAT1 | CHID1 | | | NIPSNAP2 |
| ATP6AP2 | CHIT1 | | | ARHGDIB |
| MRAP2 | CHKA | | | PROM1 |

| | | | | |
|---|---|---|---|---|
| G6PC3 | CHL1 | | | C6orf58 |
| ERO1B | CHMP1A | | | BCAS1 |
| CLK1 | CHMP1B | | | CCNA2 |
| SNX14 | CHMP4B | | | RNU4-1 |
| PTGES3P3 | CHMP5 | | | PLAAT2 |
| CCT6A | CHMP6 | | | LINC02605 |
| AC006511.4 | CHN1 | | | KHDC4 |
| KLF9 | CHPF | | | TMEM181 |
| FKBP1C | CHPF2 | | | FSD1L |
| PIPSL | chr22-38_28785274-29006793.1 | | | SH2B3 |
| FAM162A | CHRAC1 | | | ANXA1 |
| AC024293.1 | CHRNB4 | | | KANSL1 |
| TMEM147 | CHROMR | | | WFDC5 |
| ALG1 | CHST10 | | | SPTA1 |
| CCDC59 | CHST11 | | | TGIF2 |
| DBP | CHST2 | | | TLCD4 |
| TF | CHST6 | | | PCM1 |
| FKBP9 | CHSY1 | | | FRAS1 |
| RPL14P1 | CHTOP | | | CDC20 |
| TMEM43 | CHUK | | | GSPT1 |
| ACAD10 | CIAO1 | | | VANGL1 |
| BORCS5 | CIAO2A | | | ARFGEF1 |
| EMC7 | CIAPIN1 | | | TICAM1 |
| ACTA1 | CIB1 | | | USP15 |
| UNC93B6 | CIC | | | LMO2 |
| EDEM2 | CICP23 | | | KIF1C |
| PTK7 | CIITA | | | CERT1 |
| ARSDP1 | CIPC | | | DSC3 |
| TMED7 | CIR1 | | | FNBP1L |
| TUSC3 | CIRBP | | | AKR1B10 |
| PGRMC1 | CISD2 | | | THRA |
| UBLCP1 | CISD3 | | | GPR107 |
| AC026271.1 | CITED2 | | | MTHFD1 |
| BMI1 | CKB | | | CALM1 |
| DYNC1I2P1 | CKS1B | | | CENPI |
| ARMT1 | CLCA3P | | | RNU6ATAC |
| TLR1 | CLCF1 | | | CDK17 |
| GPC1 | CLCN2 | | | PLEKHB2 |
| VDAC1P1 | CLCN3 | | | NUDT21 |
| HNRNPKP4 | CLCN6 | | | CTSE |
| RRAGA | CLCN7 | | | NAP1L1 |
| KIT | CLDN1 | | | CHD3 |
| ST13P3 | CLDN2 | | | CTNNB1 |
| AC136632.1 | CLDN4 | | | AC013394.1 |
| AL391244.2 | CLDN5 | | | SET |
| ETHE1 | CLDN7 | | | PFKFB2 |
| TMED1 | CLDND1 | | | PLAC8 |
| RPL7P10 | CLEC1A | | | H1-5 |
| HEXB | CLEC4A | | | IL15 |
| MOSPD1 | CLEC4E | | | GALNT2 |
| SLC52A2 | CLEC7A | | | AC087721.2 |
| KDSR | CLIC3 | | | RASGRF2 |
| SELENOS | CLIC4 | | | RNU2-2P |
| HLA-A | CLIP2 | | | PER2 |
| ADH1A | CLIP4 | | | INO80 |
| RPL22P1 | CLK1 | | | CACYBP |
| DNAJB9 | CLK3 | | | FARP1 |
| AC083873.1 | CLK4 | | | TRIB3 |
| DENND10P1 | CLMP | | | ETFA |
| FUCA1 | CLPP | | | CTTN |

| | | | | |
|---|---|---|---|---|
| HSPA8P5 | CLPTM1L | | | B3GNT2 |
| LPCAT3 | CLPX | | | MRPS16 |
| SLC35B2 | CLSTN1 | | | SMC2 |
| EXOSC9 | CLSTN3 | | | ATP5F1E |
| TAGLN2P1 | CLTB | | | PHKB |
| ATP5MC3 | CLUH | | | HARS1 |
| EIF3FP3 | CLUHP3 | | | ATL3 |
| COL6A3 | CMAHP | | | GIMAP2 |
| CHCHD1 | CMC1 | | | TFPI |
| TMCO1 | CMC2 | | | SLC30A9 |
| PABPC3 | CMIP | | | CKAP5 |
| TP63 | CMPK2 | | | NDUFA4 |
| CTSC | CMTM3 | | | SNORD3B-1 |
| HSD17B12 | CMTR2 | | | RPL35A |
| CEACAM3 | CMYA5 | | | BCL3 |
| SMG1P1 | CNBP | | | TTLL11 |
| PSMC1P1 | CNGB1 | | | HNF4A |
| ELMO3 | CNIH4 | | | TBC1D5 |
| AL158801.6 | CNKSR1 | | | DIAPH2-AS1 |
| UQCRC1 | CNOT1 | | | G6PD |
| HLA-V | CNOT2 | | | CEP55 |
| MGST1 | CNOT3 | | | KLHL29 |
| NTS | CNOT7 | | | AC104389.5 |
| AL133477.1 | CNPY2 | | | RPS21 |
| H3P44 | CNRIP1 | | | SCARNA7 |
| PSENEN | CNST | | | PLEKHG7 |
| RPL7AP34 | CNTLN | | | ZNF207 |
| PSPC1 | CNTN1 | | | DESI1 |
| LAMC1 | CNTROB | | | AK4 |
| AL121769.1 | COA5 | | | GM2A |
| NDUFA12 | COASY | | | VILL |
| MT-TL1 | COG1 | | | H2BC6 |
| AC106795.1 | COG3 | | | LMCD1 |
| AC022968.1 | COG4 | | | OR4C6 |
| LRRIQ1 | COG6 | | | PLOD2 |
| LRRC17 | COG7 | | | TSPAN13 |
| UFD1 | COL12A1 | | | EIF4B |
| UXS1 | COL16A1 | | | PPP2R5D |
| AC008065.1 | COL1A1 | | | MFSD12 |
| KRT6B | COL20A1 | | | TNFRSF1B |
| HMGN2P17 | COL4A1 | | | SPNS2 |
| ALOX15P1 | COL4A4 | | | PGD |
| RPL37AP1 | COL5A1 | | | PRDX3 |
| ATP6AP1 | COL5A2 | | | SEPHS1 |
| AC099670.1 | COL5A3 | | | CBX5 |
| BMP3 | COL8A1 | | | TCF12 |
| FKSG70 | COL9A2 | | | ACTN4 |
| FAM171A1 | COLGALT1 | | | ACSF2 |
| RPN2 | COMMD10 | | | ANKRD10 |
| HLA-H | COMMD3 | | | ARHGAP44 |
| EEF1A1P38 | COMMD4 | | | PARP3 |
| THNSL2 | COMMD7 | | | ERVK3-1 |
| PDIA3 | COMTD1 | | | SLC25A6 |
| AC064799.1 | COP1 | | | LYST |
| KTN1 | COPA | | | CAV1 |
| AC209007.1 | COPB1 | | | ATP11B |
| AC104619.3 | COPE | | | PRDX2 |
| SYT8 | COPS3 | | | SPRY4 |
| COQ5 | COPS5 | | | PPP1CA |
| MSH2 | COPS7A | | | SNORA23 |

| | | | | |
|---|---|---|---|---|
| NPC1 | COPS7B | | | NIN |
| UQCRFS1P1 | COPZ1 | | | CMPK1 |
| MORF4L1P1 | COPZ2 | | | CDCA7 |
| UBBP4 | COQ2 | | | SNORA54 |
| GPR89A | COQ4 | | | TNNT1 |
| MFSD5 | COQ8A | | | ZNF689 |
| SERPINB4 | COQ9 | | | XPO6 |
| HNRNPA1P48 | CORIN | | | AC010197.2 |
| AP000936.3 | CORO1C | | | GPC4 |
| SIRT3 | COX10-AS1 | | | ZNF300P1 |
| AC092115.2 | COX16 | | | AC124319.3 |
| RPS27P29 | COX18 | | | FKRP |
| CYP26A1 | COX4I1 | | | SELENOF |
| ATP13A5 | COX5A | | | MBOAT2 |
| AL109918.1 | COX5B | | | TTN |
| PIGO | COX6A1 | | | TRIOBP |
| ITM2B | COX6A1P2 | | | GEM |
| RPL9P32 | COX6B1 | | | PPP2R5C |
| PFN1P1 | COX6C | | | TPGS2 |
| EIF2S2P4 | COX7A1 | | | PBK |
| HNRNPA3P6 | COX7B | | | SHMT1 |
| HSP90AB3P | COX7C | | | GNPNAT1 |
| PLLP | COX8A | | | NECTIN3 |
| CLDN1 | CPAMD8 | | | PHC2 |
| ANXA8L1 | CPD | | | RNF144A |
| RPL23P8 | CPE | | | CLMN |
| AC113404.3 | CPLX1 | | | SERF2 |
| HSP90AA6P | CPLX2 | | | ATP8B1 |
| RPL7P47 | CPNE3 | | | LCP1 |
| SETP20 | CPNE8 | | | DBN1 |
| PHF14 | CPSF1 | | | AKIRIN1 |
| SRD5A3 | CPSF3 | | | FYB1 |
| SCAMP3 | CPSF4 | | | LMO7 |
| AC244034.1 | CPSF6 | | | NIPSNAP1 |
| TRIAP1 | CPSF7 | | | LMNB2 |
| F2RL1 | CR1 | | | NRIR |
| PPIAP87 | CR392039.1 | | | SLC9A8 |
| AP002784.2 | CRABP2 | | | SERPING1 |
| SUMO2P1 | CRACR2B | | | ATP5MG |
| AC005000.1 | CRAMP1 | | | GSDMD |
| HLA-C | CRAT | | | ECE1 |
| MTATP8P1 | CRCT1 | | | TCIM |
| KRT18P16 | CREB3 | | | CXADR |
| SERBP1P5 | CREB3L2 | | | MAFA |
| RPL4P5 | CREBZF | | | SUMF2 |
| DCAF13 | CRELD2 | | | DTYMK |
| SRSF2 | CREM | | | SLC25A5 |
| SETSIP | CRIP2 | | | ATRAID |
| FKBP9P1 | CRIPAK | | | ARHGAP5 |
| PRCP | CRISP3 | | | ELK4 |
| CD9 | CRK | | | APPL1 |
| IFNGR1 | CRNDE | | | MORF4L1P1 |
| RPL7P9 | CROCC | | | LPGAT1 |
| TMEM212 | CRTC2 | | | KAT6A |
| EIF4BP3 | CRTC3 | | | RIPK1 |
| PSCA | CRY2 | | | NPNT |
| AL049597.1 | CRYAB | | | CCDC18 |
| RPL7P32 | CRYBG3 | | | AC079781.5 |
| YWHAZP5 | CRYL1 | | | HEXIM1 |
| AC012085.1 | CRYZL2P | | | AC008038.1 |

| | | | | |
|---|---|---|---|---|
| HNRNPCP2 | CS | | | DCAF11 |
| FTLP3 | CSDE1 | | | CDCA7L |
| APLP2 | CSE1L | | | AC016831.1 |
| SETP14 | CSF2RA | | | RANBP3L |
| UBE2I | CSF2RB | | | MYL6 |
| CDC42P6 | CSF3R | | | MGAT4B |
| ANXA8 | CSNK1A1 | | | RAPGEF5 |
| AL354702.1 | CSNK1D | | | GLUD1 |
| FGFR3 | CSNK1E | | | SSRP1 |
| H3-5 | CSNK2A2 | | | VDAC3 |
| TSPAN6 | CSNK2B | | | FANCD2 |
| EIF4A1P2 | CSPG4 | | | TNS3 |
| RPL7P1 | CSRNP1 | | | CDC42 |
| GAPDHP65 | CST7 | | | IRS1 |
| AC105250.1 | CSTA | | | NKILA |
| EPHA1 | CSTB | | | TMEM132A |
| AC002075.2 | CSTF2 | | | SH3TC1 |
| RPS7P10 | CTA-342B11.3 | | | MICB |
| EEF1A1P19 | CTBP1 | | | CCDC88A |
| EIF5AL1 | CTBP1-DT | | | MIR100HG |
| RPL10AP2 | CTBS | | | AC245014.3 |
| ARSD | CTC1 | | | LSM3 |
| XRCC6P2 | CTCF | | | RNU12 |
| RPL13AP20 | CTD-2369P2.2 | | | CALM2 |
| AC092597.1 | CTD-2521M24.9 | | | SLC25A22 |
| EIF4BP6 | CTD-3195I5.1 | | | COA3 |
| AC099560.2 | CTDNEP1 | | | LCLAT1 |
| RPL12P38 | CTDP1 | | | MMD2 |
| RPS26P6 | CTDSP1 | | | ADIRF-AS1 |
| LDHBP2 | CTDSP2 | | | HTATIP2 |
| KRT18P11 | CTDSPL | | | WASHC5 |
| AC004057.1 | CTDSPL2 | | | MCM6 |
| RPL34P26 | CTF1 | | | CENPA |
| BZW1P2 | CTNNA1 | | | LSM4 |
| AC092683.1 | CTNNB1 | | | ERN1 |
| S100A4 | CTNNBIP1 | | | DLC1 |
| HLA-DRB6 | CTNNBL1 | | | SETD2 |
| C1GALT1C1 | CTNND1 | | | SERTAD1 |
| RPS26P31 | CTPS1 | | | GBF1 |
| SLC5A8 | CTPS2 | | | FZD8 |
| MTCO1P40 | CTR9 | | | MARCKSL1 |
| GAPDHP44 | CTSA | | | ANO9 |
| RPL7AP11 | CTSC | | | ACTL6A |
| PPIC | CTSE | | | OIP5-AS1 |
| TMX1 | CTSH | | | FAM20B |
| ITGA6 | CTSL | | | CCR1 |
| GAPDHP61 | CTSS | | | OAF |
| PSMC1P5 | CTSZ | | | SP140 |
| EIF4BP7 | CTTN | | | PRSS12 |
| PPIAP16 | CTXN1 | | | ZUP1 |
| EEF1A1P7 | CUL1 | | | RPL18 |
| RPSAP19 | CUL4A | | | ZNF3 |
| AC005480.2 | CUL9 | | | CYB5B |
| UNC50 | CUTALP | | | BCL2L13 |
| MIR22HG | CWC22 | | | STK19 |
| RPS26P8 | CWF19L2 | | | SYNE2 |
| AC115223.1 | CXCL10 | | | ZBTB38 |
| HSPA8P1 | CXCL11 | | | DSC2 |
| DPYD | CXCL16 | | | JUP |
| AC126120.1 | CXCL17 | | | MEG8 |

| | | | | |
|---|---|---|---|---|
| AC016734.1 | CXCL2 | | | CD38 |
| RARRES1 | CXCL6 | | | PAH |
| MTCO3P12 | CXCR2 | | | CXXC5 |
| DPY30 | CXorf21 | | | GNPAT |
| ALG5 | CXorf40A | | | RNPEP |
| PPIAP66 | CXorf40B | | | EXOC3L4 |
| PPIAP43 | CXXC1 | | | GANAB |
| AC112187.1 | CXXC5 | | | AC084816.1 |
| PPIAL4C | CYB561 | | | E2F7 |
| MTND4P12 | CYB561A3 | | | SCARB2 |
| TMEM183B | CYB5A | | | KIF12 |
| ACTBP2 | CYB5B | | | GNAS |
| HLA-G | CYB5D2 | | | STRBP |
| RPS7P11 | CYB5R1 | | | VMA21 |
| NACA3P | CYB5R2 | | | MFSD2A |
| EEF1A1P29 | CYBB | | | IFNL3P1 |
| AC068522.1 | CYBRD1 | | | LRRC20 |
| RPS26P11 | CYC1 | | | TUFT1 |
| AC034236.1 | CYFIP1 | | | APEH |
| MTCO2P2 | CYFIP2 | | | HOGA1 |
| AC020898.1 | CYHR1 | | | TUBB |
| CLCA4 | CYP19A1 | | | SLC16A9 |
| LYPD3 | CYP1A1 | | | ATP5MPL |
| EEF1A1P4 | CYP24A1 | | | USP25 |
| AC004552.1 | CYP2J2 | | | MYO9A |
| NAMPTP1 | CYP2R1 | | | MICAL3 |
| PPIAP13 | CYP2S1 | | | MAPRE1 |
| RPS4XP22 | CYP3A5 | | | TPI1 |
| EEF1A1P25 | CYP4F11 | | | SFSWAP |
| TUBAP2 | CYP4F3 | | | UROD |
| CROT | CYP7B1 | | | LONP2 |
| AP000281.2 | CYREN | | | TLK2 |
| RPL10P12 | CYSLTR1 | | | SIX4 |
| FTH1P15 | CYSTM1 | | | GALNT12 |
| RPL3P4 | CYTH1 | | | TNFRSF10D |
| AC104339.1 | CYTH3 | | | RNVU1-27 |
| RPS23P8 | CYTIP | | | BTC |
| PPIAP31 | DAAM1 | | | CHROMR |
| AC135178.7 | DAAM2 | | | C5 |
| RPL10P4 | DAB2IP | | | NR4A3 |
| AC104563.1 | DACT1 | | | KNL1 |
| RPS26P47 | DAG1 | | | IFNAR1 |
| H3P16 | DAGLB | | | ZFC3H1 |
| FTH1P11 | DANCR | | | GAS5 |
| EEF1A1P16 | DAP | | | TRIM2 |
| YWHAZP3 | DAP3 | | | EIF4A1 |
| GAPDHP73 | DAPK2 | | | ZHX2 |
| GAPDHP63 | DAPK3 | | | CREBL2 |
| AL627402.1 | DAPP1 | | | AGR2 |
| AC009245.1 | DARS1 | | | AL645929.1 |
| RPS3AP5 | DARS-AS1 | | | MAD2L1 |
| H3P6 | DAXX | | | DNAH9 |
| RPL27AP5 | DAZAP1 | | | HNRNPUL2 |
| RPS26P15 | DAZAP2 | | | KRT7 |
| H3P47 | DBF4B | | | UBASH3B |
| AC078819.1 | DBN1 | | | FBN2 |
| TCN1 | DBNDD1 | | | STIM2 |
| HSP90AA2P | DBNL | | | DIRAS3 |
| FTH1P7 | DBR1 | | | OR1F1 |
| RPS7P1 | DCAF10 | | | PRKAR1A |

| | | | | |
|---|---|---|---|---|
| RPS27AP16 | DCAF11 | | | TMPO |
| RPL7P19 | DCAF13 | | | C11orf58 |
| TMSB4XP2 | DCAF15 | | | DMBX1 |
| RPS26P28 | DCAF16 | | | RPS10-NUDT3 |
| RPL10AP6 | DCAF4 | | | TNRC6A |
| PPIAP22 | DCAF5 | | | GRIP2 |
| EEF1A1P8 | DCAF6 | | | FYB2 |
| ADH1B | DCAF8 | | | JAKMIP3 |
| ATP1B3 | DCAKD | | | TC2N |
| PPIAP6 | DCBLD1 | | | CHSY1 |
| RPL3P2 | DCHS1 | | | GABRE |
| RPL13AP25 | DCLRE1A | | | IL22RA1 |
| MTND6P4 | DCP1A | | | MARS1 |
| RPS24P8 | DCP1B | | | KLHL3 |
| AC092865.1 | DCPS | | | VAMP5 |
| RPL15P20 | DCTD | | | RUFY4 |
| AL133260.1 | DCTN2 | | | ACOT13 |
| FTH1P16 | DCTN4 | | | TM7SF2 |
| EEF1A1P11 | DCTN5 | | | ASPH |
| RPS15AP1 | DCTN6 | | | GRWD1 |
| FTH1P3 | DCUN1D3 | | | NECAP1 |
| APOD | DCUN1D4 | | | FEM1A |
| AL596275.1 | DDAH1 | | | KMO |
| RPL15P18 | DDB1 | | | PKP4 |
| RPL17P22 | DDB2 | | | GGCT |
| KRT6C | DDHD2 | | | AC091133.1 |
| EIF4A1P10 | DDIT4 | | | ADPRHL2 |
| PDIA3P1 | DDOST | | | FAM149A |
| FTH1P20 | DDR2 | | | AMER1 |
| EEF1A1P22 | DDT | | | LNPK |
| ANXA2P2 | DDTL | | | SESN3 |
| AC073072.1 | DDX1 | | | TNFSF9 |
| AC025518.1 | DDX17 | | | AC015849.1 |
| S100A2 | DDX23 | | | U62317.1 |
| RPL7P23 | DDX31 | | | TMEM41A |
| MTND6P3 | DDX39A | | | IK |
| AL009174.1 | DDX39B | | | PBX1 |
| AC092670.1 | DDX41 | | | SNORD13 |
| PPIAP29 | DDX42 | | | B4GALT4 |
| FTH1P12 | DDX5 | | | NOTCH3 |
| AC090543.3 | DDX51 | | | TMEM245 |
| H3C9P | DDX54 | | | CSDE1 |
| RPL17P36 | DDX55 | | | AIFM2 |
| MTND5P11 | DDX56 | | | NUP62 |
| MT-TE | DDX58 | | | ANAPC16 |
| MTCO2P12 | DDX6 | | | CEP57 |
| MTRNR2L1 | DDX60 | | | NYNRIN |
| HLA-J | DDX60L | | | AL355075.4 |
| EEF1A1P12 | DEAF1 | | | RPPH1 |
| EEF1A1P14 | DEF6 | | | PPIAP22 |
| AC091429.1 | DEF8 | | | GBP7 |
| FTH1P5 | DELE1 | | | GMFB |
| TPT1P9 | DENND11 | | | YWHAE |
| EEF1A1P13 | DENND2A | | | HMGXB3 |
| AC006386.2 | DENND2B | | | EGLN1 |
| AC012005.1 | DENND2C | | | UGT1A10 |
| RPL41P2 | DENND2D | | | ARL4A |
| MT-TA | DENND3 | | | G3BP2 |
| | DENND4B | | | AL021707.6 |
| | DENND4C | | | HOOK1 |

| | | | | |
|---|---|---|---|---|
| | DENND6A | | | AC008760.2 |
| | DENR | | | SERPINA4 |
| | DEPDC5 | | | HNRNPU |
| | DEPP1 | | | FUT8 |
| | DERL2 | | | TLN1 |
| | DESI1 | | | ZNF433 |
| | DESI2 | | | IPO8 |
| | DGAT1 | | | SAMD1 |
| | DGCR2 | | | CPNE2 |
| | DGCR6 | | | WDR75 |
| | DGCR8 | | | RMND5A |
| | DGCR9 | | | IL1RN |
| | DGKD | | | AL138787.2 |
| | DGKE | | | TTYH3 |
| | DGKG | | | ADD3 |
| | DGUOK | | | MRPL24 |
| | DHCR24 | | | OBSL1 |
| | DHDDS | | | ZNF44 |
| | DHODH | | | ZNF532 |
| | DHPS | | | DENND4B |
| | DHRS12 | | | NBEA |
| | DHRS4L1 | | | EEF1A2 |
| | DHRS4L2 | | | RNF5 |
| | DHRS7 | | | CDC25C |
| | DHRS9 | | | HGD |
| | DHRSX | | | SBNO1 |
| | DHX15 | | | LSM5 |
| | DHX16 | | | SMCR8 |
| | DHX29 | | | ZSWIM8 |
| | DHX30 | | | PDCL3 |
| | DHX32 | | | IL11 |
| | DHX35 | | | KLHDC7A |
| | DHX38 | | | DUSP2 |
| | DHX57 | | | OPA1 |
| | DHX8 | | | BCAR1 |
| | DHX9 | | | UPF1 |
| | DIAPH1 | | | THUMPD3-AS1 |
| | DIDO1 | | | AL591222.1 |
| | DIP2A | | | RAD21 |
| | DIP2C | | | CXCL9 |
| | DIPK2A | | | AL645929.3 |
| | DIS3 | | | SNRPD3 |
| | DIS3L2 | | | SPATA2 |
| | DISP1 | | | RPS10 |
| | DKC1 | | | KLC4 |
| | DKK1 | | | HP1BP3 |
| | DKK3 | | | DNMBP |
| | DLC1 | | | EPB41L1 |
| | DLEU1 | | | KHNYN |
| | DLG4 | | | ACLY |
| | DLGAP1-AS1 | | | ITGA5 |
| | DLL1 | | | POLR3D |
| | DLL3 | | | RAET1L |
| | DLL4 | | | PLAT |
| | DLST | | | CALU |
| | DLX1 | | | PYGB |
| | DM1-AS | | | ZEB1 |
| | DMAC2 | | | AC083862.2 |
| | DMBT1 | | | RECQL5 |
| | DMBX1 | | | ZNF222 |

| | | | | |
|---|---|---|---|---|
| | DMKN | | | LEPROT |
| | DMPK | | | PXMP4 |
| | DMXL2 | | | ARHGAP42 |
| | DNAAF1 | | | SLC39A7 |
| | DNAAF2 | | | UNC119B |
| | DNAAF3 | | | PSMA7 |
| | DNAH1 | | | COPS9 |
| | DNAH14 | | | SNRNP200 |
| | DNAH7 | | | PRR3 |
| | DNAJA3 | | | ATP2C1 |
| | DNAJB12 | | | ADH5 |
| | DNAJB14 | | | HDAC2 |
| | DNAJB2 | | | GATA6 |
| | DNAJB5 | | | RAPGEF2 |
| | DNAJC10 | | | SNCA |
| | DNAJC11 | | | SYAP1 |
| | DNAJC16 | | | TBCC |
| | DNAJC18 | | | MAD2L1BP |
| | DNAJC2 | | | HK1 |
| | DNAJC3 | | | MIER3 |
| | DNAJC4 | | | CDC42BPA |
| | DNAJC5 | | | CCDC34 |
| | DNAJC7 | | | ERI3 |
| | DNALI1 | | | EP300 |
| | DNASE1 | | | UGT2B7 |
| | DNM1P51 | | | ZFAS1 |
| | DNM2 | | | DYNC2H1 |
| | DNMBP | | | ZFX |
| | DNMT1 | | | ODF2 |
| | DNPEP | | | GET4 |
| | DNTTIP1 | | | LINC00641 |
| | DNTTIP2 | | | LINC00265 |
| | DOCK1 | | | LRRFIP1 |
| | DOCK10 | | | CHCHD2 |
| | DOCK2 | | | IKBIP |
| | DOCK4 | | | JAZF1 |
| | DOCK5 | | | HSBP1 |
| | DOCK6 | | | WIF1 |
| | DOCK8 | | | OR10AB1P |
| | DOCK8-AS1 | | | FNIP1 |
| | DOCK9 | | | NR4A2 |
| | DOHH | | | BBX |
| | DOK1 | | | EXPH5 |
| | DOK3 | | | FARSB |
| | DOK4 | | | LYSMD2 |
| | DOLK | | | FH |
| | DONSON | | | IER3 |
| | DOP1A | | | MIR6891 |
| | DPF2 | | | RNVU1-15 |
| | DPH1 | | | DYRK1A |
| | DPH2 | | | HSP90AA1 |
| | DPH5 | | | SEPTIN11 |
| | DPH7 | | | MIR320A |
| | DPM1 | | | ACOX1 |
| | DPM3 | | | RPS23 |
| | DPP7 | | | DCAF6 |
| | DPT | | | TRPC4 |
| | DPY19L1 | | | RBBP4 |
| | DPY30 | | | LRRN3 |
| | DPYSL2 | | | HFE |

| | | | | |
|---|---|---|---|---|
| | DPYSL3 | | | TENT5C |
| | DPYSL5 | | | CMBL |
| | DRAP1 | | | KMT2E |
| | DRG1 | | | SEC24D |
| | DROSHA | | | MMUT |
| | DSCC1 | | | RPL15 |
| | DST | | | AAR2 |
| | DSTN | | | C2CD4A |
| | DSTYK | | | SLC9A1 |
| | DTWD1 | | | ZBED5-AS1 |
| | DTX2 | | | SMARCC2 |
| | DTX3 | | | RNVU1-6 |
| | DTX3L | | | EIF3L |
| | DTX4 | | | NQO1 |
| | DUOX1 | | | FAM107B |
| | DUOX2 | | | RANBP17 |
| | DUS2 | | | REPS2 |
| | DUS3L | | | BCL10 |
| | DUSP1 | | | SYT12 |
| | DUSP10 | | | QARS1 |
| | DUSP16 | | | LMAN2 |
| | DUSP18 | | | SUPT7L |
| | DUSP23 | | | SND1 |
| | DUSP3 | | | HOXB6 |
| | DUSP7 | | | CD109 |
| | DUSP8 | | | PLEKHA2 |
| | DUX4L2 | | | SSTR2 |
| | DUX4L5 | | | TREX1 |
| | DUX4L6 | | | NDC1 |
| | DUX4L7 | | | HLA-G |
| | DUXAP9 | | | SNORA80B |
| | DVL1 | | | AKT1S1 |
| | DYM | | | ANKFY1 |
| | DYNC1H1 | | | RHBDF2 |
| | DYNC2LI1 | | | ZNF629 |
| | DYNLL1 | | | H2AC12 |
| | DYNLRB1 | | | C12orf49 |
| | DYNLT1 | | | LEKR1 |
| | DYRK2 | | | THEM6 |
| | DYRK3 | | | TACC1 |
| | DYSF | | | CCDC117 |
| | DZIP1L | | | CCDC174 |
| | E2F4 | | | DNAJA1 |
| | E2F8 | | | NUP98 |
| | E4F1 | | | SCCPDH |
| | EBNA1BP2 | | | PHKA2 |
| | EBP | | | RPS19 |
| | ECE1 | | | METRNL |
| | ECHDC1 | | | TIPRL |
| | ECHDC2 | | | CBX4 |
| | ECHDC3 | | | MAP4K5 |
| | ECHS1 | | | PSMD10 |
| | ECI2 | | | PIGK |
| | ECM1 | | | TMEM97 |
| | ECM2 | | | PTPRB |
| | ECPAS | | | EFNA1 |
| | ECSIT | | | ADPGK |
| | EDC3 | | | UQCRQ |
| | EDC4 | | | HSPA6 |
| | EDEM2 | | | PPM1L |

| | | | | |
|---|---|---|---|---|
| | EDF1 | | | PSORS1C1 |
| | EDNRA | | | DGLUCY |
| | EDRF1 | | | SMARCA5 |
| | EED | | | DAZAP2 |
| | EEF1A1 | | | TPD52L1 |
| | EEF1A1P5 | | | BSDC1 |
| | EEF1A1P6 | | | PIP4K2B |
| | EEF1AKMT1 | | | E2F8 |
| | EEF1AKMT3 | | | NCAPD3 |
| | EEF1B2 | | | PRPF8 |
| | EEF1D | | | VPS13C |
| | EEF2 | | | COPG2 |
| | EFCAB11 | | | G2E3 |
| | EFEMP2 | | | NFX1 |
| | EFHC1 | | | CHD1 |
| | EFHD2 | | | TTC3 |
| | EFL1 | | | SWAP70 |
| | EFNA1 | | | FDCSP |
| | EFNA2 | | | PPFIBP1 |
| | EFNB1 | | | AF127936.5 |
| | EFNB2 | | | PPP2R5A |
| | EFTUD2 | | | AC068299.2 |
| | EGFL7 | | | LINC02044 |
| | EGFLAM | | | HCFC1R1 |
| | EGLN1 | | | HAPLN3 |
| | EGLN2 | | | SCARNA21 |
| | EGR1 | | | TASOR2 |
| | EGR2 | | | EHMT2 |
| | EHBP1 | | | ASAP1 |
| | EHBP1L1 | | | STRAP |
| | EHD2 | | | TTC39C |
| | EHD4 | | | GTSE1 |
| | EHMT1 | | | BRAP |
| | EIF1 | | | UFC1 |
| | EIF1AD | | | RASAL1 |
| | EIF1B | | | BRPF1 |
| | EIF2AK1 | | | KCNN1 |
| | EIF2AK2 | | | AL645608.7 |
| | EIF2AK4 | | | EIF4H |
| | EIF2B1 | | | MMGT1 |
| | EIF2B4 | | | H2AC13 |
| | EIF2B5 | | | ZFP36L1 |
| | EIF2D | | | SKAP2 |
| | EIF2S3 | | | DENND5A |
| | EIF3A | | | AKT3 |
| | EIF3B | | | HCG4B |
| | EIF3C | | | AP1M2 |
| | EIF3CL | | | TUBB6 |
| | EIF3D | | | KIAA1549L |
| | EIF3F | | | LRCH3 |
| | EIF3G | | | PDHA1 |
| | EIF3H | | | URI1 |
| | EIF3J-DT | | | MVP |
| | EIF3K | | | H4C8 |
| | EIF3L | | | PAIP2B |
| | EIF4A1 | | | AASS |
| | EIF4A2 | | | MYORG |
| | EIF4A3 | | | SYTL5 |
| | EIF4E2 | | | PINLYP |
| | EIF4E3 | | | HEATR9 |

| | | | | |
|---|---|---|---|---|
| | EIF4EBP1 | | | FILIP1L |
| | EIF4ENIF1 | | | ARFIP2 |
| | EIF4G1 | | | GATA4 |
| | EIF4G3 | | | KNSTRN |
| | EIF4H | | | PLA2R1 |
| | EIF5A | | | STEAP4 |
| | EIF5AL1 | | | SOCS2 |
| | EIF6 | | | RUVBL1 |
| | EIPR1 | | | SMARCA1 |
| | ELAC2 | | | SFXN3 |
| | ELF1 | | | RSU1 |
| | ELF2 | | | INTS5 |
| | ELF4 | | | UACA |
| | ELK1 | | | CD69 |
| | ELK3 | | | STK38 |
| | ELMOD2 | | | BROX |
| | ELMOD3 | | | ANKRD1 |
| | ELMSAN1 | | | SREK1 |
| | ELOB | | | RBBP8 |
| | ELOVL1 | | | HLA-DRB5 |
| | ELP1 | | | HSPA2 |
| | ELP3 | | | TNIK |
| | EMB | | | RPL31 |
| | EMC3 | | | MAPK3 |
| | EMC4 | | | PKN1 |
| | EMC7 | | | NADSYN1 |
| | EMC9 | | | TTL |
| | EME2 | | | ACSL4 |
| | EMG1 | | | STX16 |
| | EMILIN2 | | | ATP5MF |
| | EML3 | | | RPLP0P6 |
| | EMP1 | | | CEP70 |
| | EMP2 | | | PPL |
| | EMP3 | | | F3 |
| | ENC1 | | | DDX52 |
| | ENDOD1 | | | PAPSS1 |
| | ENDOV | | | EXOC4 |
| | ENG | | | WDR73 |
| | ENGASE | | | C18orf25 |
| | ENKD1 | | | AL137077.2 |
| | ENKUR | | | IGF2BP3 |
| | ENO1 | | | SELENOK |
| | ENO2 | | | TRAK2 |
| | ENO3 | | | POLR2B |
| | ENOSF1 | | | NR1D2 |
| | ENOX2 | | | CADPS2 |
| | ENSA | | | WDFY1 |
| | ENTPD4 | | | PARPBP |
| | ENTPD6 | | | RAB1B |
| | ENTPD7 | | | STK38L |
| | ENTR1 | | | MAVS |
| | EOMES | | | ARMC10 |
| | EP400 | | | CCDC167 |
| | EP400P1 | | | ARHGEF9 |
| | EPAS1 | | | RAB8B |
| | EPB41L1 | | | HELLS |
| | EPB41L2 | | | FBXO6 |
| | EPB41L4A | | | SNX14 |
| | EPB41L4A-AS1 | | | GGA3 |
| | EPC1 | | | KDM5A |

| | | | | |
|---|---|---|---|---|
| | EPDR1 | | | PXK |
| | EPGN | | | RPS3AP6 |
| | EPHA2 | | | EIF3E |
| | EPHA4 | | | EXTL3 |
| | EPHB4 | | | TMEM230 |
| | EPM2AIP1 | | | NR1D1 |
| | EPN1 | | | PSMD2 |
| | EPN2 | | | STAT3 |
| | EPN3 | | | CCNT2 |
| | EPOP | | | SRP9 |
| | EPOR | | | ZNF267 |
| | EPS8L2 | | | SPINK1 |
| | EPS8L3 | | | SMOX |
| | EPSTI1 | | | PIMREG |
| | ERBB2 | | | GALM |
| | ERC1 | | | RABGAP1L |
| | ERCC1 | | | LAPTM4B |
| | ERCC5 | | | MYEOV |
| | ERGIC1 | | | TNFAIP8L1 |
| | ERH | | | RAB10 |
| | ERI1 | | | NRDC |
| | ERN1 | | | APPL2 |
| | ERP44 | | | MXRA5 |
| | ERRFI1 | | | H4C2 |
| | ERVK13-1 | | | AC012020.1 |
| | ESF1 | | | STARD8 |
| | ESS2 | | | PLAU |
| | ESYT1 | | | SNORA84 |
| | ETS1 | | | MIR3651 |
| | ETS2 | | | CREBBP |
| | ETV1 | | | DNAJC10 |
| | ETV3 | | | NOSTRIN |
| | ETV6 | | | AC243919.1 |
| | EVA1C | | | ANO10 |
| | EVI2B | | | MIB1 |
| | EVI5 | | | ZNF140 |
| | EVL | | | CLIP1 |
| | EWSR1 | | | MAP4K3 |
| | EXO1 | | | PDK1 |
| | EXOC2 | | | NSDHL |
| | EXOC3 | | | TRERF1 |
| | EXOC4 | | | AC140912.1 |
| | EXOC6B | | | HLA-DMA |
| | EXOSC3 | | | DHX40 |
| | EXOSC4 | | | SCD5 |
| | EXOSC7 | | | ANO6 |
| | EXOSC8 | | | TOMM20 |
| | EXOSC9 | | | TSPAN17 |
| | EXT2 | | | SEC14L2 |
| | EXTL3 | | | AC008079.1 |
| | EYA1 | | | PCDH1 |
| | EYA3 | | | UBE2C |
| | EZR | | | GRIN3A |
| | F11R | | | TRHDE-AS1 |
| | F2R | | | AC119150.1 |
| | F5 | | | PCNP |
| | F8A1 | | | C19orf48 |
| | F8A3 | | | LY75 |
| | FAAH2 | | | SNHG20 |
| | FAAP100 | | | RIMKLB |

| | | | | |
|---|---|---|---|---|
| | FABP3 | | | DDIT4 |
| | FABP6 | | | ZFPM2 |
| | FADS1 | | | RDH11 |
| | FADS3 | | | CENPU |
| | FAHD2A | | | EPS8L2 |
| | FAHD2B | | | MCCC1 |
| | FAM104A | | | SPSB1 |
| | FAM110B | | | COL8A1 |
| | FAM111A | | | CHPF2 |
| | FAM114A1 | | | ZDHHC2 |
| | FAM118A | | | PIGA |
| | FAM118B | | | FAM122C |
| | FAM120B | | | MEGF6 |
| | FAM122A | | | IP6K2 |
| | FAM122B | | | ABCF1 |
| | FAM126A | | | TMEM19 |
| | FAM126B | | | SEC24A |
| | FAM131A | | | NCAPH |
| | FAM135A | | | AGPS |
| | FAM13A-AS1 | | | AP5B1 |
| | FAM13B | | | HSD17B12 |
| | FAM151B | | | EDC3 |
| | FAM156A | | | EBF4 |
| | FAM156B | | | SEPTIN10 |
| | FAM157B | | | MIPEP |
| | FAM157C | | | TESK2 |
| | FAM160A2 | | | UQCRB |
| | FAM160B1 | | | NUMA1 |
| | FAM160B2 | | | COMMD9 |
| | FAM162A | | | HAS3 |
| | FAM167A | | | RBMS3 |
| | FAM168A | | | HAVCR1 |
| | FAM168B | | | SESN2 |
| | FAM171A2 | | | RUVBL2 |
| | FAM171B | | | PLCE1 |
| | FAM174B | | | VBP1 |
| | FAM174C | | | PPIP5K2 |
| | FAM177A1 | | | VEGFA |
| | FAM177B | | | FKBP9 |
| | FAM183A | | | TEX261 |
| | FAM189A2 | | | KIAA0930 |
| | FAM193A | | | UBE2Q2 |
| | FAM198B-AS1 | | | NEK6 |
| | FAM199X | | | AC008957.3 |
| | FAM207A | | | ENC1 |
| | FAM20B | | | NCAM1 |
| | FAM20C | | | IARS1 |
| | FAM214A | | | ZNF562 |
| | FAM214B | | | HELZ |
| | FAM219B | | | HMGB1P5 |
| | FAM220A | | | PTPRU |
| | FAM222B | | | SNRPE |
| | FAM231D | | | TMEM219 |
| | FAM234A | | | FAM169A |
| | FAM234B | | | DPY30 |
| | FAM32A | | | CARD16 |
| | FAM3A | | | LCORL |
| | FAM3C | | | ANKRD17 |
| | FAM49A | | | ATRIP |
| | FAM49B | | | CEBPD |

| | | | | |
|---|---|---|---|---|
| | FAM50A | | | NMRAL2P |
| | FAM53B | | | LSR |
| | FAM72A | | | DENND3 |
| | FAM72B | | | PEBP1 |
| | FAM83B | | | AC138811.2 |
| | FAM83G | | | METTL7B |
| | FAM86JP | | | RLF |
| | FAM8A1 | | | NDST1 |
| | FAM92A | | | NPC2 |
| | FANCA | | | FAM43A |
| | FANCC | | | CTNND2 |
| | FAR2 | | | TMC4 |
| | FARP1 | | | MYO6 |
| | FARP2 | | | IRGQ |
| | FARS2 | | | OSBPL8 |
| | FARSA | | | IRF6 |
| | FARSB | | | NFE2L2 |
| | FAS | | | RNU1-2 |
| | FASN | | | SNHG15 |
| | FASTK | | | NEDD8 |
| | FASTKD2 | | | PEX16 |
| | FASTKD5 | | | MGST3 |
| | FAT4 | | | CEP78 |
| | FAU | | | RELA |
| | FAXDC2 | | | BX119927.1 |
| | FBH1 | | | PATJ |
| | FBL | | | HADH |
| | FBLN1 | | | RNU4ATAC |
| | FBLN5 | | | DNTTIP2 |
| | FBN2 | | | SMG7 |
| | FBRS | | | OAZ2 |
| | FBXL12 | | | NAV3 |
| | FBXL15 | | | DHX30 |
| | FBXL3 | | | CCDC47 |
| | FBXL4 | | | FYTTD1 |
| | FBXL5 | | | DMAC1 |
| | FBXL7 | | | RSF1 |
| | FBXO21 | | | ID3 |
| | FBXO25 | | | KCNH8 |
| | FBXO27 | | | RPL34 |
| | FBXO31 | | | COX7A2 |
| | FBXO32 | | | BLVRB |
| | FBXO39 | | | RPL22 |
| | FBXO42 | | | SGK2 |
| | FBXO6 | | | UBXN6 |
| | FBXO9 | | | PMVK |
| | FBXW11 | | | STRN4 |
| | FBXW2 | | | AC010733.2 |
| | FBXW4 | | | AFF1 |
| | FBXW5 | | | SPDEF |
| | FCER1G | | | RNF11 |
| | FCGR1A | | | PSAT1 |
| | FCGR1B | | | ACTG1 |
| | FCGR2A | | | ANKRD9 |
| | FCGR2B | | | NDC80 |
| | FCGR2C | | | SDCBP2 |
| | FCGR3A | | | CDKN2AIP |
| | FCHO2 | | | LYN |
| | FCRLA | | | LIG1 |
| | FDCSP | | | STK32A |

| | | | | |
|---|---|---|---|---|
| | FDFT1 | | | CHML |
| | FDPS | | | TYMS |
| | FEM1C | | | IMPDH1 |
| | FEN1 | | | RNVU1-29 |
| | FER | | | AC007192.1 |
| | FERMT2 | | | KPNA1 |
| | FES | | | AL365436.2 |
| | FEZ1 | | | DSTN |
| | FFAR2 | | | FANCI |
| | FGD4 | | | QKI |
| | FGD5 | | | KMT2B |
| | FGD5-AS1 | | | KCNC4 |
| | FGF1 | | | PPCS |
| | FGF2 | | | LAMTOR5 |
| | FGF5 | | | ANKLE2 |
| | FGF7P6 | | | ILRUN |
| | FGFR1 | | | COX7A2L |
| | FGFR1OP2 | | | PRKCA |
| | FGFR2 | | | CNOT6LP1 |
| | FGFR4 | | | HMGB2 |
| | FGG | | | NDUFS1 |
| | FGL1 | | | ALDH3A2 |
| | FGR | | | ANKS6 |
| | FHL1 | | | SIK2 |
| | FHL3 | | | MAP3K14 |
| | FHOD1 | | | AP002381.2 |
| | FIBIN | | | TMEM9 |
| | FIBP | | | RPS7P1 |
| | FILIP1 | | | SLC7A5 |
| | FIP1L1 | | | SLC35G2 |
| | FIS1 | | | CIAO2A |
| | FIZ1 | | | LINC01963 |
| | FJX1 | | | CCN1 |
| | FKBP10 | | | WAC |
| | FKBP14 | | | SLC44A2 |
| | FKBP1A | | | STT3A |
| | FKBP1C | | | RPS18 |
| | FKBP4 | | | VGLL1 |
| | FKBP8 | | | ANK3 |
| | FKSG62 | | | BBS9 |
| | FKTN | | | ITSN2 |
| | FLAD1 | | | RPTOR |
| | FLG | | | SUCLG1 |
| | FLG2 | | | BZW1 |
| | FLI1 | | | STX10 |
| | FLII | | | TINF2 |
| | FLNA | | | TSPYL2 |
| | FLNB | | | SLC25A23 |
| | FLNC | | | C11orf96 |
| | FLOT1 | | | LAMC1 |
| | FLRT2 | | | FGD5-AS1 |
| | FLT1 | | | AKAP13 |
| | FLT4 | | | BAHCC1 |
| | FLVCR2 | | | AC069224.2 |
| | FLYWCH1 | | | MGST1 |
| | FLYWCH2 | | | SAA1 |
| | FMR1 | | | RPL13 |
| | FNBP4 | | | AC016831.6 |
| | FNDC3B | | | USP11 |
| | FNDC4 | | | ALDH1A2 |

| | | | | |
|---|---|---|---|---|
| | FOS | | | KDM7A |
| | FOSB | | | BCAP31 |
| | FOSL1 | | | SNORD3C |
| | FOXA1 | | | RAB5B |
| | FOXD1 | | | MYOM1 |
| | FOXD3-AS1 | | | SQOR |
| | FOXE1 | | | GORAB |
| | FOXJ2 | | | THBS1 |
| | FOXJ3 | | | COPB2 |
| | FOXK2 | | | INS-IGF2 |
| | FOXN2 | | | CYBA |
| | FOXN3 | | | USH1C |
| | FOXO1 | | | OSER1 |
| | FOXO4 | | | PPARA |
| | FOXP4-AS1 | | | RBM33 |
| | FOXRED1 | | | STK39 |
| | FOXRED2 | | | C12orf75 |
| | FP236383.1 | | | EPC2 |
| | FP236383.9 | | | TACC3 |
| | FP565260.1 | | | CAP1 |
| | FP565260.3 | | | SLC40A1 |
| | FP565260.6 | | | RAN |
| | FP671120.2 | | | CKAP4 |
| | FP671120.6 | | | GPATCH8 |
| | FPGS | | | AP000873.2 |
| | FPR1 | | | MPHOSPH9 |
| | FREM2 | | | GGH |
| | FRMD4A | | | ACP2 |
| | FRMD6 | | | BCO1 |
| | FRMD8 | | | TRAPPC6A |
| | FRS2 | | | DNPH1 |
| | FRYL | | | RNF114 |
| | FSCN1 | | | PNPLA2 |
| | FSIP2 | | | LATS2 |
| | FSTL3 | | | TPRKB |
| | FTH1 | | | SORBS2 |
| | FTH1P10 | | | SNRK |
| | FTH1P11 | | | HOMER1 |
| | FTH1P12 | | | RHOA |
| | FTH1P15 | | | AL590867.2 |
| | FTH1P16 | | | CARS1 |
| | FTH1P2 | | | REEP3 |
| | FTH1P20 | | | IGF2-AS |
| | FTH1P23 | | | CD151 |
| | FTH1P3 | | | MIR22HG |
| | FTH1P4 | | | TYW5 |
| | FTH1P5 | | | LAMB3 |
| | FTH1P7 | | | RN7SL396P |
| | FTH1P8 | | | PROCR |
| | FTL | | | KRT23 |
| | FTLP3 | | | KIFC1 |
| | FTO | | | TTC9B |
| | FTSJ3 | | | NHSL2 |
| | FUBP1 | | | ZNF296 |
| | FUBP3 | | | NSUN6 |
| | FUCA1 | | | PPT1 |
| | FUNDC2 | | | RPL24 |
| | FURIN | | | MRPL30 |
| | FUT1 | | | CEBPA |
| | FXR1 | | | MEG9 |

| | | | | |
|---|---|---|---|---|
| | FXR2 | | | MRPL33 |
| | FXYD6 | | | PELO |
| | FYB1 | | | PDE5A |
| | FYCO1 | | | IL18R1 |
| | FYN | | | MYO5C |
| | FYTTD1 | | | STC1 |
| | FZD4 | | | AC079594.2 |
| | FZD8 | | | GULP1 |
| | FZR1 | | | LRP8 |
| | G0S2 | | | HIRA |
| | G2E3 | | | ATXN2L |
| | G6PD | | | ZMPSTE24 |
| | GABARAPL1 | | | CD99L2 |
| | GABARAPL2 | | | H1-2 |
| | GABBR1 | | | SKA2 |
| | GABPB1 | | | RPL13A |
| | GABPB1-AS1 | | | SNHG19 |
| | GABRA5 | | | ITM2C |
| | GABRB3 | | | SIK3 |
| | GABRE | | | PIH1D1 |
| | GABRP | | | ZMIZ2 |
| | GABRQ | | | MAN1A2 |
| | GADD45A | | | FAM161A |
| | GADD45B | | | NPTX1 |
| | GADD45G | | | PAK3 |
| | GADD45GIP1 | | | FKBP8 |
| | GAK | | | SRSF6 |
| | GAL | | | DHX37 |
| | GALC | | | KDELR2 |
| | GALNT1 | | | POLK |
| | GALNT10 | | | ADAMTS1 |
| | GALNT11 | | | UQCR10 |
| | GALNT18 | | | TPM2 |
| | GALNT2 | | | SRD5A3-AS1 |
| | GALT | | | METTL7A |
| | GAN | | | COL18A1 |
| | GAPDH | | | GATA3 |
| | GAPVD1 | | | TMEM47 |
| | GARS-DT | | | RLIM |
| | GART | | | IFNA7 |
| | GAS1 | | | ABCC4 |
| | GAS2L3 | | | AC244517.11 |
| | GAS5 | | | NME1 |
| | GAS6 | | | TCEAL4 |
| | GATA2 | | | UBASH3A |
| | GATA6 | | | HNRNPUL1 |
| | GATD1 | | | ZBTB17 |
| | GATD3A | | | MED15 |
| | GBA | | | GHITM |
| | GBA2 | | | TSKU |
| | GBF1 | | | NFIX |
| | GBGT1 | | | NDUFB3 |
| | GBP1 | | | EIF3K |
| | GBP1P1 | | | CLCN2 |
| | GBP2 | | | PDZK1IP1 |
| | GBP3 | | | SYS1 |
| | GBP4 | | | CMTM6 |
| | GBP5 | | | CDCA2 |
| | GCA | | | ADAP1 |
| | GCC1 | | | USF3 |

| | | | | |
|---|---|---|---|---|
| | GCFC2 | | | ID1 |
| | GCH1 | | | SLC16A3 |
| | GCN1 | | | MPZL1 |
| | GCNT2 | | | SPA17 |
| | GCNT3 | | | RDX |
| | GDE1 | | | GSTM3 |
| | GDF11 | | | CAPN5 |
| | GDF15 | | | LST1 |
| | GDI1 | | | TBC1D16 |
| | GDPD5 | | | PLEKHG6 |
| | GEM | | | SENP5 |
| | GEMIN2 | | | FOPNL |
| | GET3 | | | CEP250 |
| | GFER | | | TTC39B |
| | GFOD1 | | | BTG2 |
| | GFOD2 | | | GSTP1 |
| | GGA1 | | | MRPL51 |
| | GGA2 | | | AJ003147.2 |
| | GGA3 | | | AC005332.6 |
| | GGH | | | BISPR |
| | GGNBP2 | | | AC105105.4 |
| | GID8 | | | ABCA5 |
| | GIGYF1 | | | KBTBD8 |
| | GIMAP5 | | | AC092117.1 |
| | GIPC1 | | | DTNA |
| | GIT1 | | | NELFE |
| | GIT2 | | | VANGL2 |
| | GJA4 | | | C4orf3 |
| | GJA5 | | | MTFR1 |
| | GJC1 | | | SVIP |
| | GK | | | VOPP1 |
| | GK5 | | | SELL |
| | GKN2 | | | ERG28 |
| | GLDN | | | EML4 |
| | GLG1 | | | SUDS3 |
| | GLI3 | | | DCAF13 |
| | GLIPR1 | | | BRF2 |
| | GLMP | | | C11orf24 |
| | GLOD4 | | | CRY2 |
| | GLRX | | | SSR1 |
| | GLT8D1 | | | ADK |
| | GMCL1 | | | SEC62 |
| | GMDS | | | TMED3 |
| | GMEB2 | | | RPL7A |
| | GMFG | | | NOD2 |
| | GMPPA | | | IPO5 |
| | GMPPB | | | OSGIN1 |
| | GMPR | | | HMGN2P46 |
| | GMPS | | | ESRP1 |
| | GNA11 | | | FAM214A |
| | GNA12 | | | KIF2C |
| | GNAI3 | | | RPA1 |
| | GNAS | | | RNU11 |
| | GNB1 | | | FNDC3B |
| | GNB2 | | | COX11 |
| | GNB4 | | | PPARGC1A |
| | GNB5 | | | EIF2S3 |
| | GNG11 | | | MST1R |
| | GNG2 | | | SRD5A3 |
| | GNG5 | | | STK17B |

| | | | | |
|---|---|---|---|---|
| | GNG5P2 | | | BAZ2A |
| | GNL1 | | | PARP1 |
| | GNL2 | | | LSMEM1 |
| | GNL3 | | | AC073957.3 |
| | GNL3L | | | CTSV |
| | GNLY | | | H1-6 |
| | GNPAT | | | DCTN1 |
| | GNPDA2 | | | FAT1 |
| | GNPNAT1 | | | FSIP2 |
| | GNPTG | | | ABCD1 |
| | GNS | | | RNF130 |
| | GOLGA1 | | | FAM172A |
| | GOLGA2 | | | SLC9A3R1 |
| | GOLGA3 | | | KLHDC7B |
| | GOLGA4 | | | CHMP4B |
| | GOLGA6L4 | | | MAPK13 |
| | GOLGA6L5P | | | DENR |
| | GOLGA6L9 | | | SLC7A1 |
| | GOLGA7 | | | CACNA1D |
| | GOLGA8A | | | NGRN |
| | GOLGA8B | | | F11R |
| | GOLGB1 | | | CCDC71L |
| | GOLM1 | | | KNOP1 |
| | GOLPH3L | | | TMEM106C |
| | GON4L | | | TBK1 |
| | GOT2 | | | RAB4A |
| | GPAA1 | | | PPFIA1 |
| | GPANK1 | | | BAAT |
| | GPAT4 | | | SLC25A40 |
| | GPATCH11 | | | SAE1 |
| | GPATCH2 | | | AC011466.3 |
| | GPATCH3 | | | BTG1 |
| | GPATCH4 | | | RHBDD3 |
| | GPATCH8 | | | FLCN |
| | GPBP1 | | | LPIN2 |
| | GPBP1L1 | | | IQCE |
| | GPC3 | | | ROMO1 |
| | GPD1 | | | DIP2C |
| | GPD1L | | | APIP |
| | GPD2 | | | ATP10D |
| | GPKOW | | | CERS6 |
| | GPLD1 | | | APRT |
| | GPM6B | | | AZIN2 |
| | GPN1 | | | TRIM36 |
| | GPN2 | | | HNF1B |
| | GPN3 | | | HLA-DRB6 |
| | GPNMB | | | AGPAT1 |
| | GPR107 | | | HSPE1 |
| | GPR108 | | | AL049766.1 |
| | GPR132 | | | BAIAP2L2 |
| | GPR141 | | | HINT1 |
| | GPR155 | | | BDNF |
| | GPR160 | | | SATB2 |
| | GPR161 | | | TBCA |
| | GPR180 | | | DOCK1 |
| | GPR37L1 | | | TERC |
| | GPR65 | | | REC8 |
| | GPR68 | | | MXD4 |
| | GPR84 | | | EPHA2 |
| | GPR89A | | | CAV2 |

| | | | | |
|---|---|---|---|---|
| | GPR89B | | | POLR1C |
| | GPRC5A | | | BTN2A3P |
| | GPRC5B | | | SMC1A |
| | GPS1 | | | SBF2 |
| | GPSM3 | | | RAD54L |
| | GPX1 | | | DHRS7 |
| | GPX2 | | | ZNF473 |
| | GPX4 | | | IL6-AS1 |
| | GRAMD1A | | | IRGM |
| | GRASP | | | UBXN2B |
| | GRB10 | | | HIP1 |
| | GRB7 | | | BMPR2 |
| | GRHL1 | | | AC007991.2 |
| | GRHPR | | | KCTD3 |
| | GRIN2D | | | PTGER2 |
| | GRINA | | | AL360012.1 |
| | GRK3 | | | CUL7 |
| | GRK5 | | | MORF4L2 |
| | GRPEL2 | | | TMEM239 |
| | GRSF1 | | | IL18 |
| | GRWD1 | | | MAML3 |
| | GS1-358P8.4 | | | SEMA6A |
| | GSAP | | | RPS7 |
| | GSDMD | | | PAGR1 |
| | GSDME | | | CC2D1B |
| | GSK3A | | | INTS6L |
| | GSN | | | FOCAD |
| | GSN-AS1 | | | PSRC1 |
| | GSR | | | PSMD3 |
| | GSS | | | CPNE3 |
| | GSTA4 | | | RNF216 |
| | GSTM3 | | | SNX12 |
| | GSTO1 | | | CHRDL1 |
| | GSTP1 | | | ATPSCKMT |
| | GSTT2B | | | EVPL |
| | GTDC1 | | | RASD1 |
| | GTF2B | | | CSAG3 |
| | GTF2F1 | | | AL031777.1 |
| | GTF2H1 | | | MAU2 |
| | GTF2H2 | | | TOMM40L |
| | GTF2H2B | | | ERO1B |
| | GTF2H2C | | | MICU1 |
| | GTF2I | | | HMGN5 |
| | GTF2IP1 | | | GTF2F1 |
| | GTF2IP13 | | | SCAMP1-AS1 |
| | GTF2IP4 | | | TNFRSF12A |
| | GTF2IP7 | | | TNFRSF11B |
| | GTF3C1 | | | MAP3K10 |
| | GTF3C3 | | | ARL3 |
| | GTF3C5 | | | BTBD19 |
| | GTPBP1 | | | CYFIP1 |
| | GTPBP10 | | | HMGN1 |
| | GTPBP2 | | | OSTM1 |
| | GTPBP4 | | | MOB3B |
| | GTPBP6 | | | MCM2 |
| | GTPBP8 | | | COX17 |
| | GUCD1 | | | NAA60 |
| | GUCY1A2 | | | SEMA4G |
| | GUF1 | | | HPRT1 |
| | GUK1 | | | NHP2 |

| | | | | |
|---|---|---|---|---|
| | GUSB | | | RPL12 |
| | GYG1 | | | WNT7B |
| | GYPC | | | FZD5 |
| | GYS1 | | | LINC02100 |
| | H1-0 | | | TAOK3 |
| | H1-10 | | | TTK |
| | H19 | | | DTX4 |
| | H2AC11 | | | POC5 |
| | H2AC18 | | | ZNF79 |
| | H2AC19 | | | DLX2 |
| | H2AC20 | | | RGS20 |
| | H2AC6 | | | SMAD5 |
| | H2AC7 | | | STIP1 |
| | H2AJ | | | AC006042.2 |
| | H2AX | | | MIS18BP1 |
| | H2BC12 | | | STX3 |
| | H2BC18 | | | SRP14 |
| | H2BC21 | | | DNAJB5 |
| | H2BC4 | | | MTCH2 |
| | H2BC5 | | | SMAD2 |
| | H2BP1 | | | AC104073.1 |
| | H2BU1 | | | SLC25A1 |
| | H3-3A | | | PALM3 |
| | H3-3B | | | AC104532.1 |
| | H3-5 | | | PKD2 |
| | H3C10 | | | HID1 |
| | H3C4 | | | NAPSB |
| | H3P14 | | | USP30-AS1 |
| | H3P16 | | | PRDX1 |
| | H3P36 | | | MTPN |
| | H3P6 | | | SRSF7 |
| | H4C14 | | | NDUFA1 |
| | H4C15 | | | ZBTB44 |
| | H4C9 | | | COG3 |
| | HABP4 | | | LAMB1 |
| | HACD2 | | | NFKBIB |
| | HACD3 | | | APAF1 |
| | HACL1 | | | WDR24 |
| | HAGH | | | WSB2 |
| | HAPLN3 | | | PIK3C2B |
| | HARS1 | | | ACAT1 |
| | HARS2 | | | ZNF75A |
| | HAS2 | | | NACC2 |
| | HAT1 | | | NT5DC1 |
| | HAUS2 | | | AC068860.1 |
| | HAVCR1 | | | RN7SKP71 |
| | HBA1 | | | SYCP2 |
| | HBA2 | | | NDUFB5 |
| | HBEGF | | | IPO11 |
| | HBP1 | | | MYO5A |
| | HBS1L | | | MBTD1 |
| | HCAR1 | | | HNRNPM |
| | HCAR2 | | | RRAD |
| | HCFC1R1 | | | ZMYND11 |
| | HCG18 | | | CBX1 |
| | HCK | | | USP42 |
| | HCLS1 | | | OGFRL1 |
| | HCST | | | WDR54 |
| | HDAC11 | | | VDR |
| | HDAC3 | | | TXNDC12 |

| | | | | |
|---|---|---|---|---|
| | HDAC4 | | | SACS |
| | HDAC5 | | | MTCO1P11 |
| | HDAC6 | | | AC022149.1 |
| | HDAC7 | | | C11orf68 |
| | HDGF | | | AC005062.1 |
| | HDHD2 | | | KIAA0355 |
| | HDHD5 | | | HMGB1P6 |
| | HDLBP | | | NDUFB1 |
| | HEATR3 | | | AC078802.2 |
| | HECA | | | SLC2A4RG |
| | HECTD1 | | | SMARCA2 |
| | HECTD3 | | | ATAD2B |
| | HECTD4 | | | STYK1 |
| | HELB | | | C1orf61 |
| | HELLS | | | SPTBN2 |
| | HELZ | | | TINAG |
| | HELZ2 | | | ZNF672 |
| | HERC1 | | | MMP1 |
| | HERC2 | | | TASOR |
| | HERC2P2 | | | TNFRSF19 |
| | HERC2P9 | | | SLC2A6 |
| | HERC3 | | | RYK |
| | HERC5 | | | AC004130.2 |
| | HERC6 | | | GUK1 |
| | HERPUD1 | | | HS3ST1 |
| | HES1 | | | MYLIP |
| | HES2 | | | GNL3L |
| | HES7 | | | AF279873.2 |
| | HESX1 | | | LYRM7 |
| | HEXA | | | FADS2 |
| | HEXB | | | DRAM1 |
| | HEXIM1 | | | BMPR1B |
| | HEYL | | | WHAMM |
| | HGD | | | COTL1 |
| | HGH1 | | | ARRB2 |
| | HGS | | | AL021707.3 |
| | HHLA1 | | | H2BC19P |
| | HHLA2 | | | THAP12 |
| | HID1 | | | OS9 |
| | HIF1A-AS1 | | | LY75-CD302 |
| | HIF1AN | | | AC011287.1 |
| | HIGD2A | | | SRPK1 |
| | HIKESHI | | | GPD1L |
| | HILPDA | | | AC110079.1 |
| | HINFP | | | PRMT5 |
| | HINT3 | | | NECAP2 |
| | HIP1R | | | CFI |
| | HIPK1 | | | SNORA79B |
| | HIPK2 | | | TRIQK |
| | HIPK3 | | | PDZD8 |
| | HIRA | | | NR2F2 |
| | HIRIP3 | | | SERPINH1 |
| | HIST2H2BC | | | SAV1 |
| | HK1 | | | SART3 |
| | HLA-A | | | CASC4 |
| | HLA-B | | | PCCB |
| | HLA-DOB | | | PKM |
| | HLA-DPA1 | | | FAF2 |
| | HLA-DPB1 | | | SNORD3D |
| | HLA-DQA1 | | | SNORD3D |

| | | | | |
|---|---|---|---|---|
| | HLA-DQB1 | | | ADAM28 |
| | HLA-DRB1 | | | GPX8 |
| | HLA-DRB5 | | | UTP6 |
| | HLA-H | | | NBPF10 |
| | HLA-L | | | AC092153.1 |
| | HLCS | | | HIGD1A |
| | HLTF | | | ATP4A |
| | HLX | | | VASP |
| | HM13 | | | TECR |
| | HMBOX1 | | | ZRSR2P1 |
| | HMCES | | | ST6GAL1 |
| | HMG20B | | | TCF3 |
| | HMGB1P6 | | | PCBP1-AS1 |
| | HMGB2 | | | UBOX5 |
| | HMGCS1 | | | C16orf58 |
| | HMGN2 | | | IFT27 |
| | HMGN3 | | | WDR1 |
| | HMGXB3 | | | CSE1L |
| | HMOX1 | | | ACTR1B |
| | HMOX2 | | | LMNA |
| | HNF1A | | | ARAP3 |
| | HNF4G | | | KLHL20 |
| | HNRNPA0 | | | CXCL16 |
| | HNRNPA1 | | | PPP1R7 |
| | HNRNPA1P48 | | | SGSM2 |
| | HNRNPA1P7 | | | SPIN1 |
| | HNRNPA2B1 | | | PPM1H |
| | HNRNPA3P6 | | | HNRNPH2 |
| | HNRNPAB | | | DBF4B |
| | HNRNPC | | | INTS13 |
| | HNRNPD | | | RELT |
| | HNRNPDL | | | DLD |
| | HNRNPH1 | | | SNORA7B |
| | HNRNPH3 | | | MYSM1 |
| | HNRNPK | | | IFT57 |
| | HNRNPL | | | VPS26B |
| | HNRNPLL | | | PSIP1 |
| | HNRNPM | | | PGM2 |
| | HNRNPR | | | C7orf31 |
| | HNRNPUL1 | | | NUF2 |
| | HNRNPUL2 | | | FER1L4 |
| | HNRNPUL2-BSCL2 | | | SH3RF1 |
| | HOMER3 | | | PLD1 |
| | HOOK2 | | | RPL30 |
| | HOOK3 | | | CNOT4 |
| | HOPX | | | ESD |
| | HOTAIRM1 | | | MAFB |
| | HOXA5 | | | RPL24P4 |
| | HOXC10 | | | IMPAD1 |
| | HOXC13 | | | COX6B1 |
| | HP | | | ZNF436 |
| | HPCAL1 | | | CYTH1 |
| | HPF1 | | | CDKN2A |
| | HPGDS | | | ZNF646 |
| | HPR | | | CACNA1I |
| | HPRT1 | | | MYH14 |
| | HPS3 | | | COL4A1 |
| | HPS4 | | | SLMAP |
| | HPSE | | | FSTL3 |
| | HRAS | | | NRAS |

| | | | | |
|---|---|---|---|---|
| | HS1BP3 | | | PCF11-AS1 |
| | HS2ST1 | | | WDR25 |
| | HS3ST3A1 | | | RHOB |
| | HS6ST1 | | | IP6K1 |
| | HS6ST1P1 | | | PISD |
| | HSBP1 | | | CRIM1 |
| | HSD17B10 | | | CIB1 |
| | HSD17B13 | | | TMEM171 |
| | HSD17B7 | | | DCHS1 |
| | HSD17B8 | | | TSPAN3 |
| | HSDL1 | | | HSPG2 |
| | HSF1 | | | RPS15A |
| | HSH2D | | | TSTD1 |
| | HSP90AB1 | | | TLE4 |
| | HSP90AB3P | | | CCT7 |
| | HSPA12A | | | MED18 |
| | HSPA12B | | | BUB3 |
| | HSPA1B | | | RNF31 |
| | HSPA2 | | | RC3H1 |
| | HSPA4 | | | BCL9L |
| | HSPA4L | | | PITPNM1 |
| | HSPA5 | | | NUP155 |
| | HSPA6 | | | TMEM237 |
| | HSPA7 | | | AC009220.1 |
| | HSPA8 | | | PACSIN2 |
| | HSPB6 | | | ANKRD11 |
| | HSPB7 | | | RGL2 |
| | HSPB8 | | | EFHC1 |
| | HSPD1 | | | CLTC |
| | HSPG2 | | | SLC6A9 |
| | HSPH1 | | | LINC01184 |
| | HTATIP2 | | | VKORC1 |
| | HTATSF1 | | | CCL3 |
| | HTRA1 | | | LGALS8 |
| | HTRA2 | | | SMOC1 |
| | HTRA3 | | | GNAI2 |
| | HTT | | | NDUFB2 |
| | HUS1 | | | NARS1 |
| | HYAL1 | | | CPVL |
| | HYAL2 | | | MGAT5 |
| | HYI | | | KLF7 |
| | HYKK | | | LRCH1 |
| | HYOU1 | | | SLC4A4 |
| | IARS2 | | | MYADM |
| | IBTK | | | AC087239.1 |
| | ICA1 | | | IGSF8 |
| | ICAM1 | | | MAGI1 |
| | ICAM2 | | | GALNT11 |
| | ICE1 | | | MRPS27 |
| | ICOSLG | | | BRD4 |
| | ID1 | | | STAP2 |
| | ID2 | | | CTPS2 |
| | ID3 | | | NDUFA2 |
| | ID4 | | | PPA2 |
| | IDH2 | | | NSRP1 |
| | IDH3A | | | NCAPG2 |
| | IDO1 | | | CLK3 |
| | IER2 | | | PLK2 |
| | IER3 | | | ARID1A |
| | IER5L | | | ASH1L |

| | | | | |
|---|---|---|---|---|
| | IFFO1 | | | NFKBIL1 |
| | IFFO2 | | | MED24 |
| | IFI16 | | | GCC1 |
| | IFI27L1 | | | BTAF1 |
| | IFI27L2 | | | C16orf70 |
| | IFI44 | | | ADD1 |
| | IFI44L | | | MAP3K21 |
| | IFI6 | | | VWA2 |
| | IFIH1 | | | RINT1 |
| | IFIT1 | | | NDE1 |
| | IFIT2 | | | STAT4 |
| | IFIT3 | | | LINC02032 |
| | IFIT5 | | | CENPE |
| | IFITM1 | | | GET1 |
| | IFITM10 | | | SLC4A7 |
| | IFITM2 | | | SAMM50 |
| | IFITM3 | | | ELL2 |
| | IFNAR2 | | | SERTAD2 |
| | IFNGR1 | | | POFUT1 |
| | IFNGR2 | | | TMCO3 |
| | IFRD2 | | | ZNF543 |
| | IFT172 | | | HNRNPA1 |
| | IFT43 | | | AC022966.1 |
| | IFT46 | | | AC011939.1 |
| | IFT57 | | | SZT2 |
| | IGF1R | | | B3GALNT1 |
| | IGF2BP1 | | | PHTF2 |
| | IGF2R | | | AC010168.2 |
| | IGFBP1 | | | NADK |
| | IGFBP2 | | | CEP192 |
| | IGFBP3 | | | AC099343.4 |
| | IGFBP6 | | | ZNF707 |
| | IGFL1 | | | MMP7 |
| | IGFL2-AS1 | | | CRYZL2P |
| | IGFL4 | | | NBPF14 |
| | IGFLR1 | | | ABCC9 |
| | IGHMBP2 | | | MUC4 |
| | IGIP | | | MTMR12 |
| | IGSF22 | | | ALDH18A1 |
| | IGSF3 | | | NAALADL1 |
| | IGSF6 | | | APBA3 |
| | IK | | | FBXO9 |
| | IKBKB | | | AC009093.4 |
| | IKBKGP1 | | | AC132192.2 |
| | IL10RB-DT | | | RWDD1 |
| | IL11 | | | N4BP3 |
| | IL11RA | | | GK5 |
| | IL12RB2 | | | ZNF318 |
| | IL13RA1 | | | PKHD1L1 |
| | IL15 | | | MYRF |
| | IL15RA | | | TRAF7 |
| | IL17C | | | BEX3 |
| | IL18BP | | | CCT6P1 |
| | IL18RAP | | | AC006458.1 |
| | IL1A | | | USF2 |
| | IL1B | | | CCDC85B |
| | IL1R1 | | | PHLDB3 |
| | IL1R2 | | | MFSD1 |
| | IL1RL1 | | | HIVEP1 |
| | IL1RN | | | DCAF7 |

| | | | | |
|---|---|---|---|---|
| | IL27RA | | | SMAD3 |
| | IL2RG | | | RPL21 |
| | IL3RA | | | AC116407.2 |
| | IL4R | | | RSAD1 |
| | IL6ST | | | LRRC45 |
| | ILF3 | | | ILVBL |
| | ILF3-DT | | | PFDN5 |
| | ILKAP | | | USP53 |
| | ILRUN | | | SRSF1 |
| | ILVBL | | | CKLF |
| | IMMT | | | SIPA1L1 |
| | IMP3 | | | GADD45A |
| | IMP4 | | | ZSCAN32 |
| | IMPAD1 | | | UVRAG |
| | IMPDH1 | | | MAP1LC3B |
| | IMPDH2 | | | PYGL |
| | INA | | | ELOB |
| | INAFM1 | | | ARL1 |
| | INF2 | | | KALRN |
| | ING2 | | | SLC8A2 |
| | ING3 | | | CLIC1 |
| | ING4 | | | NSD2 |
| | ING5 | | | NLN |
| | INHBB | | | RPL9 |
| | INIP | | | LINC02328 |
| | INKA2 | | | TPI1P1 |
| | INMT | | | PRTFDC1 |
| | INO80 | | | CLIP4 |
| | INO80D | | | C2 |
| | INO80E | | | NSA2 |
| | INPP1 | | | PRRC2A |
| | INPP4A | | | FAM162A |
| | INPP5A | | | TRAF2 |
| | INPP5K | | | NSUN4 |
| | INSIG1 | | | IL6ST |
| | INSIG2 | | | LRP1 |
| | INSR | | | GALNT5 |
| | INSYN2B | | | PLA1A |
| | INTS1 | | | NEDD8-MDP1 |
| | INTS10 | | | EGLN3 |
| | INTS11 | | | ZNF324 |
| | INTS12 | | | SCARNA16 |
| | INTS13 | | | GCNT2 |
| | INTS3 | | | ADI1 |
| | INTS5 | | | SLCO3A1 |
| | INTS6L | | | KDM3B |
| | INTS7 | | | TMEM165 |
| | IP6K1 | | | KCNMB4 |
| | IP6K2 | | | IER5L |
| | IPO5 | | | TXNRD1 |
| | IPO8 | | | ERBB4 |
| | IQCA1 | | | DPY19L4 |
| | IQCB1 | | | CLDN23 |
| | IQCE | | | SNRPG |
| | IQGAP1 | | | LIN54 |
| | IQUB | | | NOP2 |
| | IRAK3 | | | CIC |
| | IRF1-AS1 | | | SYNJ1 |
| | IRF2 | | | HSDL2 |
| | IRF2BP1 | | | PITX1 |

| | | | | |
|---|---|---|---|---|
| | IRF2BPL | | | SPATA6L |
| | IRF5 | | | COX20 |
| | IRF7 | | | RNVU1-14 |
| | IRF9 | | | SAMD4B |
| | IRGQ | | | KANSL2 |
| | IRS2 | | | DEDD2 |
| | IRX3 | | | AC010967.1 |
| | ISCA1 | | | SPCS3 |
| | ISCU | | | AL356234.1 |
| | ISG15 | | | TMEM170B |
| | ISG20L2 | | | MGST2 |
| | ISOC2 | | | PHB |
| | IST1 | | | LIMS1 |
| | ISY1 | | | AC009133.1 |
| | ISYNA1 | | | LTA |
| | ITCH | | | CACTIN |
| | ITGA10 | | | ARPC3 |
| | ITGA2 | | | ZNF106 |
| | ITGA3 | | | BSG |
| | ITGA5 | | | SOX9 |
| | ITGA7 | | | MYO3B |
| | ITGAE | | | KDM5B |
| | ITGAM | | | PPP1CC |
| | ITGAX | | | UNC45A |
| | ITGB1BP1 | | | CCT5 |
| | ITGB2-AS1 | | | ZNF619 |
| | ITGB3 | | | LACTB2 |
| | ITGB3BP | | | DOCK11 |
| | ITGB4 | | | HOOK3 |
| | ITM2A | | | AGPAT5 |
| | ITM2C | | | DNAJB9 |
| | ITPA | | | ECHDC1 |
| | ITPKB | | | PCYOX1L |
| | ITPKC | | | DDX6 |
| | ITPR1 | | | SYK |
| | ITPR3 | | | MFF |
| | ITPRID2 | | | KIDINS220 |
| | ITPRIP | | | RALGAPA1 |
| | ITSN1 | | | CSF3 |
| | IVD | | | IKZF2 |
| | IVL | | | AL592437.2 |
| | IVNS1ABP | | | TDP2 |
| | IWS1 | | | AC093510.1 |
| | IYD | | | CHD6 |
| | JADE1 | | | BAK1 |
| | JAG1 | | | SLC2A12 |
| | JAG2 | | | FBXW2 |
| | JAK2 | | | THSD7A |
| | JAK3 | | | AP000640.2 |
| | JAM3 | | | VEPH1 |
| | JCAD | | | NDUFS5 |
| | JMJD1C | | | SPOP |
| | JMJD1C-AS1 | | | ZNF14 |
| | JMJD6 | | | DGKH |
| | JMY | | | TRMT112 |
| | JOSD1 | | | PLA2G6 |
| | JOSD2 | | | UGP2 |
| | JPT1 | | | NTN4 |
| | JPX | | | AL353795.3 |
| | JUN | | | S100A13 |

| | | | | |
|---|---|---|---|---|
| | JUNB | | | MIEN1 |
| | JUND | | | PDE10A |
| | KALRN | | | GADD45G |
| | KANK2 | | | CUEDC2 |
| | KANK3 | | | PPP1R18 |
| | KANSL1 | | | NRP1 |
| | KANSL2 | | | SETD7 |
| | KANSL3 | | | MIR1291 |
| | KARS1 | | | DCTPP1 |
| | KAT2A | | | PMS2 |
| | KAT5 | | | SLC41A2 |
| | KAT8 | | | KLHL24 |
| | KATNB1 | | | GTF2IP13 |
| | KATNBL1 | | | CSRNP2 |
| | KAZN | | | TTC19 |
| | KBTBD11 | | | MZT2B |
| | KBTBD2 | | | SCNN1A |
| | KCMF1 | | | SIRPB2 |
| | KCNAB2 | | | HMCES |
| | KCNAB3 | | | STX11 |
| | KCNC4 | | | TCTN1 |
| | KCND1 | | | PLB1 |
| | KCNE1 | | | SLC16A5 |
| | KCNE1B | | | UGT1A6 |
| | KCNE4 | | | SEH1L |
| | KCNJ2 | | | HSD17B11 |
| | KCNJ2-AS1 | | | MATN2 |
| | KCNK1 | | | CDK18 |
| | KCNK3 | | | RPL35 |
| | KCNK5 | | | CDC42EP3 |
| | KCNMA1 | | | GMPS |
| | KCNQ3 | | | ZFAND2A |
| | KCNS3 | | | IER3IP1 |
| | KCTD10 | | | IQCK |
| | KCTD11 | | | ADAMTS9 |
| | KCTD12 | | | SGK1 |
| | KCTD15 | | | NDRG2 |
| | KCTD17 | | | KDM1A |
| | KCTD18 | | | PHF3 |
| | KCTD2 | | | MME |
| | KCTD20 | | | CETN3 |
| | KCTD3 | | | DPP4 |
| | KCTD5 | | | WDHD1 |
| | KCTD7 | | | TMTC3 |
| | KDELR1 | | | LANCL1 |
| | KDELR3 | | | VWA1 |
| | KDM1A | | | SPAG16 |
| | KDM2A | | | EIF3A |
| | KDM3A | | | LINC00886 |
| | KDM3B | | | DCP2 |
| | KDM4C | | | ATP2B1 |
| | KDM5C | | | SLC52A2 |
| | KDM6B | | | ARL4D |
| | KDSR | | | UBA6 |
| | KEAP1 | | | ARMC1 |
| | KHDRBS1 | | | IL17RB |
| | KHDRBS3 | | | TRIM52 |
| | KHNYN | | | DENND4C |
| | KHSRP | | | NDUFA13 |
| | KIAA0100 | | | SUCLG2 |

| | | | | |
|---|---|---|---|---|
| | KIAA0355 | | | ISG20L2 |
| | KIAA0513 | | | CDH1 |
| | KIAA0556 | | | MRPS35 |
| | KIAA0895L | | | EYA1 |
| | KIAA0930 | | | MTMR3 |
| | KIAA1217 | | | CNIH4 |
| | KIAA1324 | | | PROX1 |
| | KIAA1328 | | | RBPJL |
| | KIAA1522 | | | RPL3L |
| | KIAA1671 | | | UBE2E3 |
| | KIF14 | | | RSPRY1 |
| | KIF18B | | | IFNGR2 |
| | KIF1C | | | YOD1 |
| | KIF21B | | | CFAP36 |
| | KIF22 | | | MRFAP1 |
| | KIF23 | | | MDH2 |
| | KIF3A | | | RPP38 |
| | KIF3B | | | RBMS1 |
| | KIF5B | | | HEBP2 |
| | KIF9-AS1 | | | ZNF184 |
| | KIFBP | | | PIGN |
| | KIFC2 | | | SUGT1P1 |
| | KIFC3 | | | TBCD |
| | KIRREL1 | | | PTPN18 |
| | KIZ | | | BAIAP2-DT |
| | KLC1 | | | MTCH1 |
| | KLC4 | | | ARSB |
| | KLF10 | | | GOLGA2P10 |
| | KLF11 | | | RBMX |
| | KLF12 | | | STK26 |
| | KLF13 | | | AC002059.3 |
| | KLF4 | | | PRR15L |
| | KLF6 | | | MATR3 |
| | KLF7 | | | VEZF1 |
| | KLF9 | | | IFNW1 |
| | KLHDC2 | | | KDM5C |
| | KLHDC3 | | | H6PD |
| | KLHDC4 | | | GTPBP4 |
| | KLHDC7B | | | RAB6A |
| | KLHL21 | | | ZNF618 |
| | KLHL22 | | | AC018521.1 |
| | KLHL23 | | | PPP2R1A |
| | KLHL24 | | | RETREG2 |
| | KLHL25 | | | ZNF746 |
| | KLHL3 | | | TESC |
| | KLHL36 | | | ALDH4A1 |
| | KLHL42 | | | HSPA1B |
| | KLHL9 | | | DDX18 |
| | KLK8 | | | IMPACT |
| | KLRC2 | | | PCK2 |
| | KMT2A | | | SGPL1 |
| | KMT2B | | | AGFG2 |
| | KMT2E | | | AL671883.2 |
| | KMT5A | | | AC020978.5 |
| | KMT5B | | | AC021205.1 |
| | KPNA2 | | | LAMB2 |
| | KPNA5 | | | HOXB7 |
| | KPNA6 | | | ELOVL7 |
| | KPNB1 | | | YTHDC1 |
| | KPRP | | | SHCBP1 |

| | | | | |
|---|---|---|---|---|
| | KRBOX4 | | | DHX15 |
| | KREMEN2 | | | CNKSR3 |
| | KRI1 | | | TIMM23B-AGAP6 |
| | KRIT1 | | | LXN |
| | KRR1 | | | PNPO |
| | KRT15 | | | TMEM135 |
| | KRT16P1 | | | NDUFA6 |
| | KRT16P2 | | | RRBP1 |
| | KRT17 | | | TFDP2 |
| | KRT17P6 | | | AC020915.1 |
| | KRT18 | | | TXLNA |
| | KRT19 | | | ADGRF1 |
| | KRT23 | | | MIGA1 |
| | KRT31 | | | LRIG3 |
| | KRT42P | | | DPF2 |
| | KRT81 | | | YTHDF1 |
| | KSR1 | | | FLT3LG |
| | KTI12 | | | MIR1248 |
| | KXD1 | | | ZFAND5 |
| | KYNU | | | AMOT |
| | L1CAM | | | SLC35F5 |
| | L2HGDH | | | IDS |
| | L3HYPDH | | | NDUFA5 |
| | L3MBTL2 | | | WDR34 |
| | L3MBTL3 | | | PRDX4 |
| | LACC1 | | | UBL5 |
| | LAGE3 | | | RNF25 |
| | LAMA2 | | | TBCB |
| | LAMA4 | | | GRIPAP1 |
| | LAMB1 | | | AC007728.2 |
| | LAMB2 | | | PLOD3 |
| | LAMC3 | | | MATR3 |
| | LAMP2 | | | GAPDH |
| | LAMP3 | | | PFDN4 |
| | LAMTOR1 | | | SLC39A6 |
| | LAMTOR2 | | | SNORA2C |
| | LAMTOR4 | | | TRMT13 |
| | LAMTOR5 | | | CENPW |
| | LAMTOR5-AS1 | | | CSF2 |
| | LANCL3 | | | ACSM3 |
| | LAPTM4A | | | ARHGEF2 |
| | LAPTM5 | | | TSPAN15 |
| | LARGE1 | | | DNAJC8 |
| | LARP1 | | | RPL4 |
| | LARP1B | | | NPC1 |
| | LARP4 | | | INCENP |
| | LARP4B | | | GOLPH3L |
| | LARP6 | | | CXorf38 |
| | LAS1L | | | ZNF408 |
| | LASP1 | | | PRDM1 |
| | LAYN | | | LTA4H |
| | LBH | | | NACA |
| | LBR | | | AL021155.5 |
| | LCAT | | | EIF2AK2 |
| | LCE1C | | | GBP6 |
| | LCN2 | | | RAB27B |
| | LCP1 | | | LIMCH1 |
| | LCP2 | | | PHF6 |
| | LDB1 | | | CDC27 |
| | LDHA | | | MARK3 |

| | | | | |
|---|---|---|---|---|
| | LDHAP4 | | | SESTD1 |
| | LDHAP7 | | | AC022916.2 |
| | LDLR | | | PHGDH |
| | LDLRAD3 | | | CNOT7 |
| | LDLRAP1 | | | BCLAF1 |
| | LENG1 | | | SLC31A2 |
| | LENG8 | | | S100P |
| | LEPROTL1 | | | KLHDC2 |
| | LERFS | | | EXOC3L1 |
| | LETM1 | | | TPRA1 |
| | LFNG | | | PON1 |
| | LGALS1 | | | DDX3X |
| | LGALS3 | | | TULP4 |
| | LGALS3BP | | | OFD1 |
| | LGALS9 | | | TBC1D8B |
| | LGALS9B | | | OSTC |
| | LGALS9C | | | STAU2 |
| | LGI4 | | | IMPDH2 |
| | LGMN | | | LINC01004 |
| | LGR4 | | | ZNF335 |
| | LHFPL6 | | | ZFHX2 |
| | LIF | | | RFPL1 |
| | LIFR | | | GNA13 |
| | LILRA6 | | | EGR4 |
| | LILRB2 | | | GUCY1B1 |
| | LILRB3 | | | RNU5F-1 |
| | LILRB5 | | | ARFGEF3 |
| | LIMA1 | | | CALM3 |
| | LIN54 | | | COX8A |
| | LINC00243 | | | GPATCH4 |
| | LINC00312 | | | CEACAMP10 |
| | LINC00342 | | | RNU6-531P |
| | LINC00470 | | | HROB |
| | LINC00504 | | | ING3 |
| | LINC00539 | | | UQCRHL |
| | LINC00589 | | | ALDH3B1 |
| | LINC00605 | | | EXOC5 |
| | LINC00623 | | | NEU4 |
| | LINC00641 | | | AC108134.2 |
| | LINC00649 | | | EHBP1L1 |
| | LINC00665 | | | TMEM268 |
| | LINC00667 | | | DDX56 |
| | LINC00907 | | | CDK19 |
| | LINC00908 | | | RHOBTB1 |
| | LINC00923 | | | RPL13AP5 |
| | LINC00937 | | | FNIP2 |
| | LINC00958 | | | ATXN10 |
| | LINC01001 | | | AKAP11 |
| | LINC01002 | | | E2F3 |
| | LINC01126 | | | TTBK2 |
| | LINC01128 | | | EPPK1 |
| | LINC01136 | | | RNF103-CHMP3 |
| | LINC01138 | | | RCAN3 |
| | LINC01145 | | | KCTD20 |
| | LINC01234 | | | AL645941.2 |
| | LINC01270 | | | UBR2 |
| | LINC01278 | | | EIF4E3 |
| | LINC01303 | | | PTPRJ |
| | LINC01347 | | | PDGFC |
| | LINC01366 | | | TM7SF3 |

| | | | | |
|---|---|---|---|---|
| | LINC01436 | | | TSC22D1 |
| | LINC01521 | | | MED26 |
| | LINC01550 | | | AC115223.1 |
| | LINC01559 | | | TMOD3 |
| | LINC01684 | | | ENPP4 |
| | LINC01695 | | | DNAH3 |
| | LINC01783 | | | SNORA81 |
| | LINC01806 | | | CRK |
| | LINC01814 | | | RNA5SP221 |
| | LINC01833 | | | MOCOS |
| | LINC01836 | | | RNF141 |
| | LINC01842 | | | HAP1 |
| | LINC01873 | | | FAM120A |
| | LINC01979 | | | LPA |
| | LINC02068 | | | MCPH1-AS1 |
| | LINC02352 | | | CDK2AP1 |
| | LINC02362 | | | AP003498.3 |
| | LINC02422 | | | FOXC1 |
| | LINC02535 | | | AC025263.1 |
| | LINC02541 | | | ZSWIM6 |
| | LINC02560 | | | SPTBN5 |
| | LINC02582 | | | DDX1 |
| | LINC02649 | | | IVNS1ABP |
| | LINC02728 | | | AZI2 |
| | LINC02731 | | | ATG2A |
| | LINC02762 | | | FBXO7 |
| | LINC02821 | | | RPGRIP1L |
| | LINC02861 | | | ATP2A2 |
| | LINP1 | | | SEL1L |
| | LINS1 | | | RNVU1-7 |
| | LIPA | | | IGHMBP2 |
| | LITAF | | | TAF8 |
| | LLGL2 | | | AC107959.2 |
| | LMAN2 | | | PIF1 |
| | LMAN2L | | | SNORA17B |
| | LMBR1 | | | LINC00921 |
| | LMBR1L | | | TWF1 |
| | LMBRD1 | | | ZNF850 |
| | LMCD1 | | | TMEM139 |
| | LMNA | | | GPATCH2 |
| | LMNB1 | | | COX16 |
| | LMNB2 | | | TMEM265 |
| | LMO1 | | | CNTRL |
| | LMO2 | | | AP3B2 |
| | LMO7 | | | H2AC21 |
| | LMOD1 | | | HKDC1 |
| | LMTK2 | | | B4GALT1 |
| | LNPEP | | | SUMO1 |
| | LNPK | | | KCNT2 |
| | LNX2 | | | DNAJC13 |
| | LOH12CR2 | | | ATP5PO |
| | LONP1 | | | EDN1 |
| | LONRF1 | | | CRTAP |
| | LPAR6 | | | PM20D2 |
| | LPCAT1 | | | E2F4 |
| | LPCAT3 | | | TLR5 |
| | LPCAT4 | | | RASSF8 |
| | LPIN1 | | | RASGRP1 |
| | LPXN | | | SERPINB1 |
| | LRATD2 | | | BCYRN1 |

| | | | | |
|---|---|---|---|---|
| | LRCH1 | | | GYS1 |
| | LRCH3 | | | ATXN7 |
| | LRFN1 | | | LINC00887 |
| | LRFN3 | | | SRSF5 |
| | LRIG1 | | | ALG14 |
| | LRIG2-DT | | | AK2 |
| | LRIG3 | | | ARHGAP26-IT1 |
| | LRP4 | | | EEF2K |
| | LRP5 | | | GNPTAB |
| | LRP5L | | | TIMM23B |
| | LRP6 | | | SLC39A8 |
| | LRPAP1 | | | AC011195.1 |
| | LRRC1 | | | AC018638.4 |
| | LRRC15 | | | RN7SL5P |
| | LRRC23 | | | BCL2L11 |
| | LRRC27 | | | PLAA |
| | LRRC32 | | | VTI1B |
| | LRRC37A9P | | | TSLP |
| | LRRC37BP1 | | | ASPHD2 |
| | LRRC40 | | | AC068305.2 |
| | LRRC42 | | | TMEM229B |
| | LRRC47 | | | AL662844.4 |
| | LRRC59 | | | ZWINT |
| | LRRC8A | | | FAM219A |
| | LRRC8E | | | AC020915.5 |
| | LRRFIP1 | | | IVD |
| | LRRFIP2 | | | RHNO1 |
| | LRRK2 | | | HAND1 |
| | LRRK2-DT | | | H2AC4 |
| | LRSAM1 | | | FGF2 |
| | LSG1 | | | MYL6B |
| | LSM1 | | | AC245033.4 |
| | LSM10 | | | USP22 |
| | LSM12 | | | ATP5MC3 |
| | LSM14A | | | SPTLC1 |
| | LSM3 | | | KIAA0586 |
| | LSM4 | | | INTS12 |
| | LSM7 | | | SLC35B2 |
| | LSS | | | SLC50A1 |
| | LST1 | | | ABCG1 |
| | LTA4H | | | ZNF285 |
| | LTB4R | | | DDHD2 |
| | LTBP1 | | | RARS1 |
| | LTBP2 | | | SSPN |
| | LTBP3 | | | MRPL45 |
| | LTF | | | PPP4C |
| | LTO1 | | | ZADH2 |
| | LUARIS | | | CEBPZOS |
| | LUC7L | | | ALDH6A1 |
| | LUC7L2 | | | FAM171B |
| | LUC7L3 | | | MAGED2 |
| | LURAP1L | | | MARCHF5 |
| | LUZP1 | | | GINS2 |
| | LY6E | | | PDCD1LG2 |
| | LY6G5B | | | AL078599.3 |
| | LY6G5C | | | RP1 |
| | LY75 | | | PI4KB |
| | LY96 | | | AC244197.3 |
| | LYAR | | | PRPF31 |
| | LYN | | | AC119674.1 |

| | | | | |
|---|---|---|---|---|
| | LYPLA1 | | | EXOC8 |
| | LYPLA2 | | | C21orf91 |
| | LYRM1 | | | HCG4 |
| | LYSMD2 | | | RNF20 |
| | LZTR1 | | | Z93930.2 |
| | LZTS1 | | | GOLM1 |
| | LZTS2 | | | PTGFRN |
| | LZTS3 | | | TSPAN12 |
| | MACF1 | | | LRRC8E |
| | MACO1 | | | GGT1 |
| | MAD2L2 | | | LYRM2 |
| | MADD | | | SNORA17B |
| | MAF1 | | | PIM3 |
| | MAFF | | | AXL |
| | MAFG | | | SNX17 |
| | MAFK | | | RERG |
| | MAGEF1 | | | ATP7A |
| | MAIP1 | | | GPN3 |
| | MAL | | | ABCA2 |
| | MALL | | | ARL4C |
| | MALSU1 | | | VGLL3 |
| | MAMDC2 | | | ACTR1A |
| | MAML1 | | | CYBRD1 |
| | MAMLD1 | | | SUGT1 |
| | MAN1B1 | | | CTSA |
| | MAN2A1 | | | SKP2 |
| | MAN2A2 | | | SERINC3 |
| | MAN2C1 | | | CERCAM |
| | MANBA | | | HJURP |
| | MAOA | | | POLA1 |
| | MAP1LC3B | | | MRPL19 |
| | MAP1LC3B2 | | | ADAM22 |
| | MAP1S | | | SLC41A1 |
| | MAP2K2 | | | ADM2 |
| | MAP2K3 | | | RNFT2 |
| | MAP2K5 | | | FYTTD1P1 |
| | MAP2K6 | | | MICAL2 |
| | MAP2K7 | | | CAB39 |
| | MAP3K12 | | | PLEKHA3 |
| | MAP3K13 | | | SHC3 |
| | MAP3K14 | | | MUC13 |
| | MAP3K20 | | | PDE4B |
| | MAP3K2-DT | | | NAA50 |
| | MAP3K3 | | | HMG20B |
| | MAP3K6 | | | COL22A1 |
| | MAP3K7CL | | | TANC2 |
| | MAP4 | | | TBILA |
| | MAP4K5 | | | CYP27A1 |
| | MAP7D1 | | | KLF2 |
| | MAP7D2 | | | TIMMDC1 |
| | MAPK1 | | | VCPIP1 |
| | MAPK11 | | | FBXW10 |
| | MAPK12 | | | COA4 |
| | MAPK14 | | | MYO18A |
| | MAPK3 | | | TTC37 |
| | MAPK7 | | | DCPS |
| | MAPK8 | | | INF2 |
| | MAPK9 | | | CHDH |
| | MAPKAP1 | | | TMEM134 |
| | MAPKAPK2 | | | AL354740.1 |

| | | | | |
|---|---|---|---|---|
| | MAPKAPK3 | | | SLC44A3 |
| | MAPKAPK5 | | | AC090541.1 |
| | MAPKBP1 | | | ERGIC2 |
| | MAPRE1 | | | GAREM1 |
| | MAPRE3-AS1 | | | CDK16 |
| | MARCHF1 | | | KMT2A |
| | MARCHF2 | | | UQCRC1 |
| | MARCHF4 | | | TYRO3 |
| | MARCHF5 | | | USP34 |
| | MARCHF7 | | | AEBP2 |
| | MARCKS | | | IGHEP2 |
| | MARF1 | | | MRPL11 |
| | MARK3 | | | LRRC8A |
| | MARK4 | | | CYP4F3 |
| | MARS1 | | | TES |
| | MAST1 | | | SCAF1 |
| | MAST2 | | | BET1L |
| | MAT2A | | | ECI2 |
| | MAU2 | | | APP |
| | MAX | | | PAFAH1B2 |
| | MBD1 | | | AMIGO2 |
| | MBD2 | | | SUMO2 |
| | MBD3 | | | RBIS |
| | MBIP | | | THSD1 |
| | MBNL1 | | | HCCS |
| | MBNL3 | | | AC005622.1 |
| | MBOAT7 | | | LNPEP |
| | MBP | | | AMZ2 |
| | MBTD1 | | | BX276092.9 |
| | MBTPS2 | | | TFCP2L1 |
| | MCAM | | | MIR210HG |
| | MCAT | | | PPIA |
| | MCC | | | LCAL1 |
| | MCCC1 | | | PRMT2 |
| | MCCC2 | | | EGR3 |
| | MCF2L | | | CNIH1 |
| | MCFD2 | | | DPH5 |
| | MCIDAS | | | OPRK1 |
| | MCL1 | | | PROS1 |
| | MCM3AP | | | AL022311.1 |
| | MCM7 | | | NME1-NME2 |
| | MCOLN1 | | | AC125807.2 |
| | MCOLN2 | | | CHPT1 |
| | MCRIP1 | | | BTG3 |
| | MCRIP2 | | | CYTH3 |
| | MCTP2 | | | ISOC1 |
| | MCTS1 | | | P3H2 |
| | MDC1 | | | CCDC115 |
| | MDFIC | | | N4BP2L2-IT2 |
| | MDH1 | | | RBM22 |
| | MDH1B | | | GNL1 |
| | MDH2 | | | TEPSIN |
| | MDK | | | LSM7 |
| | MDM1 | | | NUP160 |
| | MDM2 | | | POLR3A |
| | MDN1 | | | MYRFL |
| | ME2 | | | CCDC86 |
| | ME3 | | | BCL7C |
| | MECP2 | | | AC020907.3 |
| | MED1 | | | AC005840.4 |

| | | | | |
|---|---|---|---|---|
| | MED10 | | | TTC30A |
| | MED11 | | | AC092868.2 |
| | MED12 | | | PAFAH1B3 |
| | MED13L | | | FAHD1 |
| | MED15 | | | AC097059.1 |
| | MED16 | | | MSANTD4 |
| | MED17 | | | AC083880.1 |
| | MED18 | | | AC084018.2 |
| | MED20 | | | VPS29 |
| | MED22 | | | GPC3 |
| | MED23 | | | USP49 |
| | MED24 | | | PPP4R1L |
| | MED27 | | | CEACAM20 |
| | MED4 | | | HYOU1 |
| | MED6 | | | SP8 |
| | MED8 | | | BIN1 |
| | MED9 | | | CCND1 |
| | MEF2D | | | ARL14EP |
| | MEG3 | | | TXNDC11 |
| | MEIS1 | | | SRPRA |
| | MEIS2 | | | DNAH11 |
| | MEPCE | | | LINC00431 |
| | MERTK | | | ZNF638 |
| | MESD | | | ZNF654 |
| | METRNL | | | PHYH |
| | METTL16 | | | LIMA1 |
| | METTL17 | | | AL671277.1 |
| | METTL23 | | | MECOM |
| | METTL26 | | | ANXA13 |
| | METTL2A | | | GPR161 |
| | METTL3 | | | KLHDC3 |
| | METTL5 | | | CARNMT1 |
| | METTL7A | | | UBXN4 |
| | METTL7B | | | EPHB2 |
| | METTL9 | | | PLEKHG5 |
| | MFAP1 | | | LINC00881 |
| | MFGE8 | | | RPP25 |
| | MFHAS1 | | | REXO1 |
| | MFN2 | | | AL031587.5 |
| | MFNG | | | FN3KRP |
| | MFSD1 | | | CCDC171 |
| | MFSD10 | | | GCNA |
| | MFSD14A | | | LAMP2 |
| | MFSD2A | | | AL450384.2 |
| | MFSD5 | | | SOAT1 |
| | MFSD6 | | | MIS12 |
| | MGA | | | SNX16 |
| | MGAM | | | HECTD1 |
| | MGAT1 | | | NLRP3 |
| | MGAT4B | | | RHOF |
| | MGAT4C | | | C5orf24 |
| | MGAT5 | | | ALS2 |
| | MGLL | | | NDUFC2 |
| | MGMT | | | PRXL2B |
| | MGRN1 | | | CAPN1 |
| | MGST3 | | | FREM2 |
| | MIA2 | | | RNU6-9 |
| | MIA3 | | | PVT1 |
| | MIB1 | | | ACTR2 |
| | MICA | | | ZKSCAN2 |

| | | | | |
|---|---|---|---|---|
| | MICAL1 | | | ZNF702P |
| | MICAL3 | | | DNHD1 |
| | MICALL2 | | | MTG2 |
| | MICB | | | GMEB2 |
| | MICOS10 | | | SUSD3 |
| | MICOS13 | | | AP001160.1 |
| | MID1 | | | CCDC130 |
| | MID1IP1 | | | SMARCD2 |
| | MID2 | | | MIER1 |
| | MIDN | | | PET100 |
| | MIEF1 | | | ADAMTSL4-AS1 |
| | MIEN1 | | | KRT18 |
| | MIER1 | | | DYNLT1 |
| | MIER2 | | | AC009495.3 |
| | MIF4GD | | | WASF2 |
| | MIGA2 | | | EPB41L2 |
| | MILR1 | | | FAH |
| | MINDY3 | | | AC126283.2 |
| | MINK1 | | | AIG1 |
| | MIOS | | | TPMT |
| | MIPOL1 | | | CRLF2 |
| | MIR22HG | | | AHCYL1 |
| | MIR29B2CHG | | | CAPZA2 |
| | MIR3189 | | | CCDC191 |
| | MIR3945HG | | | YWHAB |
| | MIR4697HG | | | MPI |
| | MISP | | | MTDH |
| | MKI67 | | | STX5 |
| | MKNK2 | | | ADGRF4 |
| | MKRN2 | | | AC024293.1 |
| | MLEC | | | TXNDC17 |
| | MLF2 | | | RHEBL1 |
| | MLH1 | | | RPL17-C18orf32 |
| | MLH3 | | | TCN2 |
| | MLKL | | | AC025539.1 |
| | MLLT1 | | | GRK6 |
| | MLLT10 | | | CRYBG2 |
| | MLLT11 | | | ZMAT3 |
| | MLNR | | | STAT6 |
| | MLST8 | | | ATP6V1C1 |
| | MLX | | | SH3BGRL |
| | MLXIP | | | PIK3IP1 |
| | MLXIPL | | | PLAGL2 |
| | MMAA | | | SNX33 |
| | MMAB | | | PHLDB2 |
| | MMADHC | | | SCARNA22 |
| | MMP1 | | | OIP5 |
| | MMP10 | | | AP003499.4 |
| | MMP12 | | | TMEM44-AS1 |
| | MMP14 | | | NAA38 |
| | MMP24OS | | | TOP3B |
| | MMP25 | | | ZKSCAN8 |
| | MMP25-AS1 | | | PBDC1 |
| | MMP28 | | | WDFY2 |
| | MMP3 | | | BMP2K |
| | MMS19 | | | ODF2L |
| | MNAT1 | | | CSF1R |
| | MND1 | | | ABCA6 |
| | MOAP1 | | | RB1CC1 |
| | MOB1A | | | GPD2 |

| | | | | |
|---|---|---|---|---|
| | MOB1B | | | FBXL12 |
| | MOB2 | | | SLC43A2 |
| | MOB3C | | | RAB40B |
| | MOCOS | | | ACNATP |
| | MOCS1 | | | TET3 |
| | MOGS | | | PPP2R2A |
| | MON1A | | | AKTIP |
| | MON1B | | | KRCC1 |
| | MON2 | | | LRIF1 |
| | MORC2 | | | XDH |
| | MORC3 | | | MROH8 |
| | MORC4 | | | FAM225B |
| | MOSMO | | | LOX |
| | MOSPD1 | | | MRPL47 |
| | MOSPD2 | | | NFATC4 |
| | MOV10 | | | ZNF652 |
| | MPDU1 | | | PCYT2 |
| | MPG | | | FBXO17 |
| | MPHOSPH10 | | | CDK5RAP3 |
| | MPHOSPH8 | | | IGF1R |
| | MPLKIP | | | AKAP10 |
| | MPP1 | | | TAF4B |
| | MPP3 | | | UBTF |
| | MPP4 | | | NUP58 |
| | MPPE1 | | | HMCN1 |
| | MPRIP | | | POU2F2 |
| | MPST | | | VPS28 |
| | MPV17 | | | ZFP62 |
| | MPZ | | | NDUFAB1 |
| | MPZL1 | | | HIBADH |
| | MPZL2 | | | MALAT1 |
| | MR1 | | | ACAD10 |
| | MRE11 | | | MGLL |
| | MRFAP1 | | | NFKBID |
| | MRFAP1L1 | | | SIAH2 |
| | MRGBP | | | AL450338.1 |
| | MRI1 | | | Z95114.1 |
| | MRM2 | | | GNS |
| | MRM3 | | | ABCA10 |
| | MRNIP | | | PARP4 |
| | MRO | | | SH3TC2 |
| | MROH1 | | | ZNF597 |
| | MROH8 | | | TRDMT1 |
| | MRPL1 | | | DMKN |
| | MRPL10 | | | TMEM205 |
| | MRPL11 | | | HEXB |
| | MRPL14 | | | MIR3143 |
| | MRPL17 | | | SPAG1 |
| | MRPL18 | | | FGFR2 |
| | MRPL2 | | | RPE |
| | MRPL20 | | | OASL2P |
| | MRPL22 | | | MKKS |
| | MRPL23 | | | PDCD4 |
| | MRPL28 | | | YBX1P10 |
| | MRPL3 | | | NABP2 |
| | MRPL30 | | | TNKS2 |
| | MRPL32 | | | CEP97 |
| | MRPL33 | | | FITM2 |
| | MRPL34 | | | PLXNA3 |
| | MRPL37 | | | RPL36A-HNRNPH2 |

| | | | | |
|---|---|---|---|---|
| | MRPL39 | | | MRTFA |
| | MRPL4 | | | RANGAP1 |
| | MRPL40 | | | BIRC2 |
| | MRPL41 | | | PBX2 |
| | MRPL44 | | | NSD3 |
| | MRPL48 | | | AC009779.4 |
| | MRPL49 | | | ALDOA |
| | MRPL51 | | | KCTD5 |
| | MRPL52 | | | SCO2 |
| | MRPL54 | | | REXO2 |
| | MRPL55 | | | ATP5ME |
| | MRPS10 | | | NUDT16 |
| | MRPS15 | | | HSD17B2 |
| | MRPS18A | | | PRCP |
| | MRPS2 | | | ARSL |
| | MRPS21 | | | ZNF764 |
| | MRPS26 | | | SEC16A |
| | MRPS33 | | | AC108010.1 |
| | MRPS34 | | | RIOK3 |
| | MRPS5 | | | TMBIM1 |
| | MRPS7 | | | UNK |
| | MRS2 | | | BZW2 |
| | MRTFA | | | ZFPM2-AS1 |
| | MRTFB | | | PTPRK |
| | MSANTD3 | | | LINC01554 |
| | MSANTD4 | | | AC004477.1 |
| | MSC | | | ZNF711 |
| | MSH5 | | | ZNF888 |
| | MSH6 | | | NFIB |
| | MSL3 | | | STK40 |
| | MSMO1 | | | SLPI |
| | MSR1 | | | C2orf42 |
| | MSRB2 | | | IL1B |
| | MSTO1 | | | UHRF1 |
| | MSX2 | | | SLC16A6 |
| | MT1XP1 | | | BRAF |
| | MT2A | | | APMAP |
| | MT2P1 | | | RPS29 |
| | MTA2 | | | CA12 |
| | MTA3 | | | GDF11 |
| | MTCH1 | | | OGT |
| | MTCH2 | | | OAT |
| | MT-CO1 | | | SLC25A39 |
| | MTCO1P12 | | | P4HA1 |
| | MT-CO2 | | | MIR7-3HG |
| | MT-CO3 | | | ALX1 |
| | MT-CYB | | | AL445685.3 |
| | MTDH | | | ADGRL2 |
| | MTERF2 | | | AC005747.1 |
| | MTF1 | | | ZNF891 |
| | MTFR1L | | | WDR72 |
| | MTHFD1 | | | CKS2 |
| | MTHFD1L | | | ERBIN |
| | MTHFD2 | | | RPL10A |
| | MTHFD2L | | | AGBL5 |
| | MTHFR | | | SLC39A9 |
| | MTHFSD | | | BRCC3 |
| | MTIF3 | | | AC051619.7 |
| | MTLN | | | SLC9A6 |
| | MTM1 | | | C4BPB |

| | | | | |
|---|---|---|---|---|
| | MTMR10 | | | POLA2 |
| | MTMR12 | | | FAM71A |
| | MTMR14 | | | DHFRP1 |
| | MTMR2 | | | AC009133.6 |
| | MTMR9LP | | | LINP1 |
| | MT-ND1 | | | NT5C3AP1 |
| | MTND2P28 | | | SCO2 |
| | MT-ND6 | | | KLHL23 |
| | MTOR | | | PSMA5 |
| | MTR | | | CA8 |
| | MTREX | | | CHD9 |
| | MTRF1L | | | EPHA7 |
| | MTRNR2L12 | | | ALDH2 |
| | MTRNR2L8 | | | CASK |
| | MTSS2 | | | TTC17 |
| | MT-TC | | | SLC19A3 |
| | MT-TP | | | OXTR |
| | MTURN | | | DPCD |
| | MTX1 | | | CYB561 |
| | MTX1P1 | | | EPHX1 |
| | MTX3 | | | RPN2 |
| | MUC1 | | | RPP14 |
| | MUC20-OT1 | | | UBR4 |
| | MUC4 | | | POLD4 |
| | MUC5AC | | | AC099518.3 |
| | MUC5B | | | ATP6V1G2-DDX39B |
| | MUL1 | | | AC008750.8 |
| | MVB12B | | | FBL |
| | MVD | | | LINC01612 |
| | MVP | | | PGAM1P7 |
| | MX1 | | | OLFM3 |
| | MX2 | | | NUDCD1 |
| | MXD1 | | | KIAA1143 |
| | MXD4 | | | IPMK |
| | MXI1 | | | CTDSP2 |
| | MXRA5 | | | SEC22C |
| | MXRA7 | | | ARHGAP19 |
| | MXRA8 | | | ARID5A |
| | MYADM | | | PPP3CA |
| | MYBBP1A | | | RTF2 |
| | MYC | | | TBC1D15 |
| | MYCBP2 | | | DFFA |
| | MYCT1 | | | ALG8 |
| | MYD88 | | | CLNS1A |
| | MYDGF | | | TNFRSF21 |
| | MYH14 | | | FBXO22 |
| | MYL12A | | | AC243772.2 |
| | MYL12B | | | GPBAR1 |
| | MYL6 | | | MYO19 |
| | MYL6B | | | LPCAT2 |
| | MYLK | | | LINC01358 |
| | MYO10 | | | G3BP1 |
| | MYO15B | | | GID8 |
| | MYO18A | | | AC005154.5 |
| | MYO1C | | | GATM |
| | MYO1D | | | SGPP1 |
| | MYO1G | | | PLS3-AS1 |
| | MYO6 | | | TMPRSS3 |
| | MYO9A | | | SCAF4 |
| | MYO9B | | | CLU |

| | | | | |
|---|---|---|---|---|
| | MYOF | | | NCK1 |
| | MYOM2 | | | SH3RF3 |
| | MYZAP | | | SRP72 |
| | MZB1 | | | MUC20 |
| | MZT2A | | | SPCS1 |
| | MZT2B | | | ZNF212 |
| | N4BP1 | | | POLG |
| | N4BP3 | | | SNHG17 |
| | NAA10 | | | C6orf62 |
| | NAA20 | | | TMC7 |
| | NAA35 | | | PDXDC1 |
| | NAA38 | | | SAP30BP |
| | NAA60 | | | ZCRB1 |
| | NAB1 | | | CCNB1IP1 |
| | NAB2 | | | PPP1R9A |
| | NACA | | | NFRKB |
| | NADK | | | EIF2AK1 |
| | NADK2 | | | NME4 |
| | NADSYN1 | | | CA11 |
| | NAGK | | | DENND11 |
| | NAIF1 | | | USP19 |
| | NAMPT | | | C22orf39 |
| | NAMPTP1 | | | ATP6V0E1 |
| | NANP | | | RAF1 |
| | NAP1L4 | | | LINC00992 |
| | NAP1L5 | | | BTD |
| | NAPA | | | NUDT2 |
| | NAPG | | | RHOC |
| | NAPSA | | | AL023755.1 |
| | NARF | | | ASS1P11 |
| | NARS1 | | | AGBL1-AS1 |
| | NASP | | | CMYA5 |
| | NAT10 | | | AC073869.1 |
| | NAT14 | | | MRPL42 |
| | NAT9 | | | TMEM168 |
| | NATD1 | | | AC007923.1 |
| | NAXD | | | CETN2 |
| | NAXE | | | TMEM129 |
| | NBAS | | | AC104837.2 |
| | NBL1 | | | OSR2 |
| | NBN | | | AC006960.4 |
| | NBPF11 | | | HTD2 |
| | NBPF14 | | | IDH3B |
| | NBPF15 | | | HABP4 |
| | NBPF19 | | | TLN2 |
| | NBPF20 | | | RNVU1-18 |
| | NBPF25P | | | PRPF38B |
| | NBPF26 | | | RPL37A |
| | NBPF8 | | | LRPAP1 |
| | NBPF9 | | | OR4V1P |
| | NBR1 | | | SLC25A24 |
| | NCALD | | | DSG2 |
| | NCAPH2 | | | CACNG4 |
| | NCBP2AS2 | | | FRMD4B |
| | NCBP3 | | | PACERR |
| | NCCRP1 | | | THYN1 |
| | NCDN | | | GTF3A |
| | NCEH1 | | | USP47 |
| | NCF2 | | | KIFC2 |
| | NCF4 | | | AC091167.2 |

| | | | | |
|---|---|---|---|---|
| | NCKIPSD | | | BDH1 |
| | NCL | | | ALDH5A1 |
| | NCOA3 | | | OLA1 |
| | NCOA4 | | | AC245041.2 |
| | NCOA7 | | | SUCLA2 |
| | NCOR2 | | | VSIG1 |
| | NCS1 | | | CHKB-CPT1B |
| | NCSTN | | | MYO1D |
| | NDEL1 | | | LINC-PINT |
| | NDN | | | PTPRG-AS1 |
| | NDNF | | | TRIM67 |
| | NDRG1 | | | DUSP15 |
| | NDRG2 | | | AL451165.1 |
| | NDRG3 | | | AL603832.3 |
| | NDST1 | | | TMEM64 |
| | NDUFA10 | | | STAC |
| | NDUFA12 | | | ANKRD28 |
| | NDUFA4 | | | INTS1 |
| | NDUFA4L2 | | | CAPRIN2 |
| | NDUFA5 | | | DDX55 |
| | NDUFAB1 | | | ACTR3B |
| | NDUFAF2 | | | PRPF38A |
| | NDUFAF5 | | | LTB |
| | NDUFAF8 | | | ARF5 |
| | NDUFB1 | | | PGM1 |
| | NDUFB10 | | | SELENOT |
| | NDUFB3 | | | SNORD12B |
| | NDUFB4 | | | GRB14 |
| | NDUFB4P12 | | | ARMC5 |
| | NDUFB8 | | | TMEM213 |
| | NDUFB9 | | | MGAT1 |
| | NDUFC2 | | | ERCC6L2 |
| | NDUFS2 | | | DBR1 |
| | NDUFS4 | | | PAX8-AS1 |
| | NDUFS5 | | | RNVU1-19 |
| | NDUFS6 | | | TK1 |
| | NDUFS7 | | | MRPL44 |
| | NDUFS8 | | | MIR222HG |
| | NDUFV1 | | | AC079466.1 |
| | NDUFV2-AS1 | | | MYBL2 |
| | NEAT1 | | | FBRS |
| | NEBL | | | USP10 |
| | NECAP1 | | | JADE1 |
| | NECAP2 | | | AL136164.3 |
| | NECTIN2 | | | ARHGAP1 |
| | NEDD1 | | | GLB1 |
| | NEDD4L | | | H2BC8 |
| | NEDD8 | | | PARP6 |
| | NEDD9 | | | RRP9 |
| | NEIL2 | | | C17orf75 |
| | NEK3 | | | PEPD |
| | NEK5 | | | DDC |
| | NEK7 | | | ZNF382 |
| | NEK9 | | | NSUN5 |
| | NELFA | | | RAD9A |
| | NELFB | | | PDLIM1 |
| | NELFCD | | | WNT2B |
| | NELFE | | | ALAD |
| | NEMP2 | | | PCLAF |
| | NENF | | | RAB13 |

| | | | | |
|---|---|---|---|---|
| | NES | | | FRMD3 |
| | NEURL1B | | | TACSTD2 |
| | NEXN | | | BRK1 |
| | NF2 | | | PCMTD2 |
| | NFASC | | | ZMYND8 |
| | NFATC1 | | | ECHDC2 |
| | NFATC2 | | | ZFP69B |
| | NFATC4 | | | GANC |
| | NFE2L1 | | | AIFM1 |
| | NFE2L3 | | | KLHL18 |
| | NFIB | | | AP005482.2 |
| | NFIL3 | | | TBC1D32 |
| | NFIX | | | LAMTOR4 |
| | NFKBID | | | MDFIC |
| | NFKBIL1 | | | NCOA5 |
| | NFRKB | | | RPL11P3 |
| | NFX1 | | | RNF187 |
| | NFYC | | | PAQR4 |
| | NGRN | | | CCNDBP1 |
| | NHLRC2 | | | HOXB3 |
| | NHP2 | | | SNORA20 |
| | NIBAN1 | | | YBX1P1 |
| | NIBAN2 | | | CAVIN1 |
| | NID1 | | | MEF2D |
| | NIF3L1 | | | FKBP3 |
| | NIN | | | AGTPBP1 |
| | NINJ1 | | | ATRN |
| | NIPA2 | | | SLC10A7 |
| | NIPAL3 | | | NIPBL |
| | NIPAL4 | | | SERPINB8 |
| | NIPBL | | | MMP24OS |
| | NISCH | | | FBXO38 |
| | NIT1 | | | USP9X |
| | NKD1 | | | ZNF280C |
| | NKIRAS2 | | | ZNF592 |
| | NKTR | | | GINS4 |
| | NLE1 | | | GATD1 |
| | NLN | | | YDJC |
| | NLRP1 | | | PPP1R13L |
| | NLRP3 | | | HSPB11 |
| | NMD3 | | | ZSWIM3 |
| | NME1 | | | CLEC4O |
| | NME3 | | | MYO1E |
| | NME5 | | | MRPL43 |
| | NME8 | | | TPD52 |
| | NMI | | | SERPINB5 |
| | NMRAL1 | | | CHMP2A |
| | NMRAL2P | | | TFAM |
| | NMT1 | | | EIF3I |
| | NMT2 | | | SNHG8 |
| | NMU | | | MAGED1 |
| | NNT | | | COMMD4 |
| | NOC2L | | | TOPBP1 |
| | NOC3L | | | SLC38A6 |
| | NOC4L | | | JDP2 |
| | NOD1 | | | DDR1 |
| | NOL6 | | | CGAS |
| | NOL8 | | | TMED9 |
| | NOLC1 | | | ARFRP1 |
| | NOMO3 | | | FOXN3 |

| | | | | |
|---|---|---|---|---|
| | NOP10 | | | SLCO5A1 |
| | NOP16 | | | AP000331.1 |
| | NOP2 | | | CDH18 |
| | NOP53 | | | SMG1 |
| | NOP56 | | | GYG2 |
| | NOP58 | | | C1orf21 |
| | NOP9 | | | ITGB1BP1 |
| | NORAD | | | STOML2 |
| | NOSIP | | | IMPDH1P10 |
| | NOSTRIN | | | STYX |
| | NOTCH1 | | | POU2F1 |
| | NOTCH2 | | | AC141586.1 |
| | NOTCH4 | | | PLPBP |
| | NOVA2 | | | KHDRBS1 |
| | NOXRED1 | | | DEK |
| | NPAS1 | | | FUT10 |
| | NPC1 | | | GALC |
| | NPC2 | | | AMFR |
| | NPHS1 | | | HIGD2A |
| | NPIPA1 | | | TCEA1 |
| | NPIPA3 | | | POU5F1 |
| | NPIPB11 | | | C11orf54 |
| | NPIPB12 | | | H2BC5 |
| | NPIPB13 | | | LCN2 |
| | NPIPB3 | | | HSPA8 |
| | NPIPB4 | | | C1orf52 |
| | NPIPB5 | | | TMEM141 |
| | NPIPP1 | | | AP000944.5 |
| | NPL | | | RGS2 |
| | NPLOC4 | | | NCS1 |
| | NPM1 | | | BRD1 |
| | NPR1 | | | HIBCH |
| | NPR2 | | | MYNN |
| | NPR3 | | | IGFBP2 |
| | NPRL2 | | | E2F1 |
| | NQO1 | | | ZC3H4 |
| | NQO2 | | | NDUFA10 |
| | NR1D2 | | | LPAR1 |
| | NR1H2 | | | TNFRSF14 |
| | NR1H3 | | | AC112907.3 |
| | NR2C2 | | | AC007255.1 |
| | NR2C2AP | | | TXNL4A |
| | NR2F1-AS1 | | | DYNC1I2 |
| | NR4A1 | | | SERTAD3 |
| | NR4A2 | | | PON2 |
| | NR5A2 | | | MAFK |
| | NRBP1 | | | SAYSD1 |
| | NRCAM | | | BBS4 |
| | NRDC | | | SELENOH |
| | NRF1 | | | CHCHD1 |
| | NRGN | | | TSNARE1 |
| | NRIP1 | | | DCBLD2 |
| | NSD2 | | | UTP18 |
| | NSF | | | MCM5 |
| | NSMAF | | | IL1R1 |
| | NSMCE1 | | | PRELID3B |
| | NSMCE4A | | | SDE2 |
| | NSMF | | | TIMM23 |
| | NSRP1 | | | EDEM1 |
| | NSUN2 | | | RPL27 |

| | | | | |
|---|---|---|---|---|
| | NSUN4 | | | MYCL |
| | NSUN5 | | | C2CD5 |
| | NSUN5P1 | | | MMP24 |
| | NSUN7 | | | CSTF3 |
| | NT5C | | | TMEM217 |
| | NT5C3A | | | LINC00842 |
| | NT5C3AP1 | | | TDRKH |
| | NT5C3B | | | ANKRD18B |
| | NT5DC2 | | | CCDC25 |
| | NTHL1 | | | GLT8D1 |
| | NTMT1 | | | FCF1 |
| | NTNG2 | | | STK25 |
| | NUAK1 | | | SCAMP1 |
| | NUB1 | | | GSDMC |
| | NUBP1 | | | CERS2 |
| | NUCKS1 | | | DNAH12 |
| | NUDC | | | EPB41L4A-AS1 |
| | NUDCD3 | | | AC245041.1 |
| | NUDT12 | | | HIST1H3B |
| | NUDT16L1 | | | CDC42P6 |
| | NUDT4 | | | XPO7 |
| | NUMA1 | | | HPS5 |
| | NUMB | | | SNORA7A |
| | NUP133 | | | MCM3AP |
| | NUP153 | | | TNPO1 |
| | NUP155 | | | MCM3AP-AS1 |
| | NUP188 | | | LATS1 |
| | NUP43 | | | CKLF-CMTM1 |
| | NUP62CL | | | MEFV |
| | NUP85 | | | UGCG |
| | NUPR1 | | | ALKBH1 |
| | NUS1 | | | HELQ |
| | NUTF2 | | | ENO3 |
| | NVL | | | ZNF35 |
| | NXF1 | | | MPRIP |
| | NXNL2 | | | RPL17 |
| | NXT1 | | | H4C13 |
| | NXT2 | | | PTK7 |
| | NYNRIN | | | CTSL3P |
| | OAF | | | KDM4A |
| | OARD1 | | | PPP1R14B |
| | OAS1 | | | COMMD7 |
| | OAS2 | | | TAF2 |
| | OAS3 | | | STRIP1 |
| | OASL | | | DMD |
| | OAT | | | PANX1 |
| | OAZ2 | | | RAP1A |
| | OBSCN | | | TTLL1 |
| | ODC1 | | | MRPL23 |
| | ODF2 | | | ATM |
| | ODF2L | | | SCRN3 |
| | ODF3B | | | SNORA21B |
| | OGA | | | HNRNPK |
| | OGDH | | | CCDC152 |
| | OGFOD1 | | | FUNDC2 |
| | OGG1 | | | ACBD5 |
| | OLA1 | | | AF196969.1 |
| | OLIG1 | | | SETDB1 |
| | OLR1 | | | MED8 |
| | OMA1 | | | ECD |

| | | | | |
|---|---|---|---|---|
| | OPHN1 | | | TPT1 |
| | OR7E14P | | | FYN |
| | OR7E36P | | | TMEM72-AS1 |
| | OR7E37P | | | AL365203.2 |
| | OR7E38P | | | AKAP2 |
| | ORAI1 | | | RNPEPL1 |
| | ORAI2 | | | RAB42 |
| | ORC2 | | | AC091271.1 |
| | ORC3 | | | MZF1-AS1 |
| | ORMDL3 | | | RO60 |
| | OS9 | | | MARS2 |
| | OSBPL10 | | | SLC17A9 |
| | OSBPL1A | | | CXCR4 |
| | OSBPL2 | | | NKIRAS2 |
| | OSBPL5 | | | MYBBP1A |
| | OSBPL7 | | | CPNE8 |
| | OSBPL9 | | | ANO3 |
| | OSER1 | | | ID2 |
| | OSGEP | | | TOGARAM1 |
| | OSGIN2 | | | BRCA1 |
| | OTUB1 | | | TNFRSF8 |
| | OTUD3 | | | CCNF |
| | OTUD4 | | | CCR4 |
| | OTUD7B | | | OR1F2P |
| | OTULINL | | | CLEC7A |
| | OTX1 | | | ZFAND6 |
| | OXA1L | | | PUS7 |
| | OXCT1 | | | LNX1 |
| | OXR1 | | | ATP5MC2 |
| | OXSR1 | | | TBRG4 |
| | P2RX4 | | | YIF1B |
| | P3H1 | | | RPL6P27 |
| | P3H2 | | | DHX38 |
| | P3H3 | | | RPL7P1 |
| | P3H4 | | | ADCY7 |
| | P4HA2 | | | GLOD4 |
| | P4HB | | | SNX27 |
| | PA2G4 | | | SCGN |
| | PAAF1 | | | ROGDI |
| | PABPC1 | | | FAM199X |
| | PACC1 | | | NELFB |
| | PACS2 | | | MTR |
| | PACSIN2 | | | SCIN |
| | PACSIN3 | | | CHRNA5 |
| | PADI2 | | | POLR2L |
| | PAF1 | | | METTL5 |
| | PAFAH1B2 | | | ZNF395 |
| | PAIP1 | | | KIF3B |
| | PAK1 | | | STXBP6 |
| | PAK2 | | | CYP21A1P |
| | PALD1 | | | HSPA14 |
| | PALLD | | | TGFBR1 |
| | PALM | | | UGT8 |
| | PALMD | | | METRN |
| | PAM | | | GTF2A2 |
| | PAMR1 | | | EWSR1 |
| | PAN2 | | | ZDHHC14 |
| | PAN3 | | | VSIG10 |
| | PANK3 | | | ZBTB40 |
| | PANK4 | | | MALSU1 |

| | | | | |
|---|---|---|---|---|
| | PAPOLA | | | PPIL4 |
| | PAPPA | | | GSTK1 |
| | PAPSS2 | | | RAB1A |
| | PAQR6 | | | YIF1A |
| | PAQR9 | | | IFT172 |
| | PARD3 | | | AGGF1 |
| | PARG | | | FBLN1 |
| | PARK7 | | | TMEM14B |
| | PARL | | | VIM-AS1 |
| | PARN | | | ERP29 |
| | PARP11 | | | TMCO1 |
| | PARP12 | | | STK3 |
| | PARP14 | | | ADAM20 |
| | PARP15 | | | LINC02827 |
| | PARP16 | | | NCDN |
| | PARP2 | | | SPCS2 |
| | PARP3 | | | PCED1A |
| | PARP4 | | | BMI1 |
| | PARP4P2 | | | WDFY3 |
| | PARP8 | | | METTL1 |
| | PARP9 | | | AC022106.1 |
| | PART1 | | | YARS1 |
| | PARVA | | | AC010619.3 |
| | PARVB | | | NGDN |
| | PARVG | | | LINC02842 |
| | PASK | | | ABCC10 |
| | PATL1 | | | BORCS8 |
| | PATL2 | | | SLC35A5 |
| | PAX8-AS1 | | | PRR34-AS1 |
| | PAX9 | | | NRBF2 |
| | PAXBP1 | | | CDH3 |
| | PAXBP1-AS1 | | | SLC35B1 |
| | PAXX | | | PLCB1 |
| | PBRM1 | | | NPHP1 |
| | PBX1 | | | NABP1 |
| | PBX2 | | | SNHG22 |
| | PBXIP1 | | | CEP68 |
| | PCBD1 | | | AC092117.2 |
| | PCBP1 | | | GMNN |
| | PCBP1-AS1 | | | TBX18 |
| | PCBP2 | | | NRIP1 |
| | PCBP4 | | | PITPNB |
| | PCCA | | | BCR |
| | PCCB | | | GFPT2 |
| | PCDH1 | | | POF1B |
| | PCDH12 | | | H2BC13 |
| | PCDHA11 | | | SNUPN |
| | PCDHGB5 | | | KLHL36 |
| | PCED1A | | | NUCB1 |
| | PCED1B | | | WWP1 |
| | PCED1B-AS1 | | | RASA1 |
| | PCGF2 | | | MTX3 |
| | PCGF3 | | | BLACAT1 |
| | PCGF5 | | | CD302 |
| | PCID2 | | | LAMTOR2 |
| | PCMT1 | | | ZNF316 |
| | PCNX1 | | | LRPPRC |
| | PCNX2 | | | PCDHAC2 |
| | PCNX3 | | | PRPF6 |
| | PCP4L1 | | | URB1 |

| | | | | |
|---|---|---|---|---|
| | PCSK9 | | | MED13L |
| | PCTP | | | UCHL5 |
| | PCYOX1 | | | DOP1A |
| | PCYT1A | | | GDE1 |
| | PCYT1B | | | COL12A1 |
| | PDCD11 | | | MAP3K4 |
| | PDCD1LG2 | | | COMMD6 |
| | PDCD2 | | | CRY1 |
| | PDCD4 | | | MPP7 |
| | PDCD4-AS1 | | | ATXN7L1 |
| | PDCD5 | | | NDUFS6 |
| | PDCD6 | | | SPDL1 |
| | PDCL | | | CLSPN |
| | PDCL3 | | | SH3RF2 |
| | PDE11A | | | TCF19 |
| | PDE1C | | | ATP7B |
| | PDE2A | | | PAXBP1-AS1 |
| | PDE4B | | | PHF19 |
| | PDE4DIP | | | AL353719.1 |
| | PDE5A | | | RABGGTA |
| | PDE6D | | | STX1B |
| | PDE8B | | | MAPKAP1 |
| | PDE9A | | | PAFAH2 |
| | PDGFB | | | COPZ1 |
| | PDGFC | | | NDOR1 |
| | PDGFRA | | | TUBB2A |
| | PDGFRB | | | ACSL5 |
| | PDGFRL | | | ATP5MF-PTCD1 |
| | PDIA4 | | | XPO1 |
| | PDIA5 | | | KHSRP |
| | PDIA6 | | | RBM23 |
| | PDK4 | | | POC1B-GALNT4 |
| | PDLIM2 | | | DDX28 |
| | PDLIM3 | | | GTF2E2 |
| | PDLIM4 | | | BNIP5 |
| | PDLIM5 | | | PCMTD1 |
| | PDPK1 | | | RALGAPA1P1 |
| | PDPN | | | TEFM |
| | PDRG1 | | | TBX3 |
| | PDS5A | | | NAB1 |
| | PDSS1 | | | RPN1 |
| | PDXDC1 | | | AC073109.1 |
| | PDXK | | | CAPN15 |
| | PDZD11 | | | H2AC15 |
| | PDZD8 | | | ABI2 |
| | PEAK1 | | | ZDHHC20 |
| | PEAR1 | | | PPRC1 |
| | PEBP1 | | | C3orf38 |
| | PELI1 | | | RBMXL1 |
| | PELO | | | DYNLL1 |
| | PELP1 | | | AL732366.1 |
| | PEPD | | | LMBR1L |
| | PER1 | | | CHST4 |
| | PER2 | | | AC005090.1 |
| | PER3 | | | LIMK2 |
| | PES1 | | | SIRPA |
| | PEX1 | | | MSL3 |
| | PEX11B | | | ATP6V0E2 |
| | PEX16 | | | MMP25-AS1 |
| | PEX2 | | | PPM1A |

| | | | | |
|---|---|---|---|---|
| | PEX5 | | | AC092279.1 |
| | PFAS | | | BCKDHB |
| | PFDN1 | | | FECH |
| | PFDN2 | | | PPAN |
| | PFDN5 | | | LAGE3 |
| | PFDN6 | | | MANEA |
| | PFKFB4 | | | KCNK5 |
| | PFKL | | | COX6A1 |
| | PFKM | | | TKFC |
| | PFKP | | | ARFIP1 |
| | PFN1 | | | AC006064.6 |
| | PGAM4 | | | SLC2A13 |
| | PGAP3 | | | THAP6 |
| | PGAP4 | | | NUDT5 |
| | PGAP6 | | | OSBPL1A |
| | PGBD1 | | | MARK2 |
| | PGBD4 | | | AL645608.8 |
| | PGC | | | RNASEL |
| | PGD | | | ACSS2 |
| | PGGHG | | | AC135050.2 |
| | PGLYRP1 | | | POLR2A |
| | PGM5 | | | GSK3A |
| | PHB | | | CASP3 |
| | PHB2 | | | BCCIP |
| | PHC1P1 | | | CNPPD1 |
| | PHC2 | | | ARHGEF11 |
| | PHETA2 | | | NHLRC2 |
| | PHF10 | | | LINC01588 |
| | PHF11 | | | ETV1 |
| | PHF12 | | | RAB2A |
| | PHF13 | | | GLRX5 |
| | PHF20 | | | ATXN7 |
| | PHF20L1 | | | MIRLET7A1HG |
| | PHF23 | | | RPS27A |
| | PHF5A | | | AC022400.3 |
| | PHF6 | | | BOC |
| | PHKB | | | AC074050.2 |
| | PHKG2 | | | AKAP7 |
| | PHLDA1 | | | COA5 |
| | PHLDA2 | | | PTPRZ1 |
| | PHLDA3 | | | ATP5F1B |
| | PHLDB1 | | | AC093425.1 |
| | PHLPP1 | | | THBS3 |
| | PHLPP2 | | | FOXK1 |
| | PHPT1 | | | CXCL6 |
| | PHRF1 | | | SH3GLB2 |
| | PHTF2 | | | QSOX1 |
| | PI4K2A | | | TROAP |
| | PI4K2B | | | KRBA1 |
| | PI4KA | | | TERF2IP |
| | PI4KAP1 | | | AC009951.4 |
| | PI4KAP2 | | | TRIM29 |
| | PI4KB | | | AC008074.2 |
| | PIAS2 | | | DIAPH1 |
| | PIAS3 | | | GALNT4 |
| | PIEZO1 | | | RBAK |
| | PIGC | | | RNU5D-1 |
| | PIGL | | | AC093512.2 |
| | PIGO | | | RFFL |
| | PIGR | | | ELP2 |

| | | | | |
|---|---|---|---|---|
| | PIGS | | | SEPSECS |
| | PIGU | | | OTUB1 |
| | PIGV | | | MEA1 |
| | PIH1D1 | | | RYBP |
| | PIK3AP1 | | | UBA1 |
| | PIK3C2B | | | TYSND1 |
| | PIK3IP1 | | | FAM217B |
| | PIK3R1 | | | PHRF1 |
| | PIK3R4 | | | RPA3 |
| | PILRA | | | HEATR1 |
| | PIM1 | | | CNKSR1 |
| | PIM2 | | | SLC66A3 |
| | PIN4 | | | DCTN2 |
| | PINX1 | | | MRPL18 |
| | PIP | | | CCDC186 |
| | PIP4P1 | | | PKIB |
| | PIP4P2 | | | PDK3 |
| | PIP5K1A | | | KIF21A |
| | PIR | | | STMP1 |
| | PISD | | | BCKDK |
| | PITHD1 | | | ETS1 |
| | PITPNC1 | | | IFNA21 |
| | PITPNM1 | | | CLEC17A |
| | PITPNM2 | | | LINC02158 |
| | PITRM1 | | | PANK3 |
| | PITX1 | | | RAB11FIP4 |
| | PJA1 | | | COG4 |
| | PJA2 | | | SMYD3 |
| | PKD2 | | | ELOVL3 |
| | PKIA | | | TCTEX1D2 |
| | PKIG | | | NPM1P27 |
| | PKN1 | | | DDAH2 |
| | PKN3 | | | ZNF175 |
| | PKP4 | | | ATP6V0A4 |
| | PLA2G15 | | | ULK4 |
| | PLA2G4B | | | MIR300 |
| | PLA2G7 | | | MAF1 |
| | PLAAT2 | | | FGD4 |
| | PLAAT3 | | | MPND |
| | PLAAT4 | | | POLR2E |
| | PLAC8 | | | SPINDOC |
| | PLAC9 | | | TCFL5 |
| | PLAGL1 | | | MAP3K2 |
| | PLAUR | | | REEP6 |
| | PLBD1 | | | DCTN5 |
| | PLBD2 | | | BRWD1P2 |
| | PLCB1 | | | IPPK |
| | PLCB2 | | | EIF4E |
| | PLCB3 | | | LAMA3 |
| | PLCD1 | | | SLC5A3 |
| | PLCG1 | | | SEC61G |
| | PLCG2 | | | RPL23A |
| | PLCL2 | | | AC090527.3 |
| | PLD2 | | | RAB3D |
| | PLD5 | | | SIM2 |
| | PLEC | | | FLII |
| | PLEK | | | AQP4 |
| | PLEKHA2 | | | CRPPA |
| | PLEKHA3 | | | SYNGR1 |
| | PLEKHA4 | | | RTKN2 |

| | | | | |
|---|---|---|---|---|
| | PLEKHA5 | | | PLCD1 |
| | PLEKHA7 | | | LMF2 |
| | PLEKHB2 | | | GABPA |
| | PLEKHG1 | | | NAA25 |
| | PLEKHG2 | | | ZCCHC8 |
| | PLEKHG3 | | | CAVIN2 |
| | PLEKHH2 | | | NUDT14 |
| | PLEKHJ1 | | | CIP2A |
| | PLEKHM1 | | | RPAP1 |
| | PLEKHM2 | | | TRAPPC2L |
| | PLEKHO1 | | | LINC02029 |
| | PLEKHO2 | | | RB1 |
| | PLGRKT | | | RMDN1 |
| | PLIN2 | | | KLF5 |
| | PLK2 | | | AL021707.1 |
| | PLK3 | | | TP53I11 |
| | PLLP | | | ATP13A4 |
| | PLOD1 | | | AOC3 |
| | PLP2 | | | CDKAL1 |
| | PLPBP | | | ELOVL5 |
| | PLPP1 | | | ABCB10 |
| | PLPP3 | | | CASP6 |
| | PLPPR2 | | | ZNF484 |
| | PLSCR1 | | | SRP19 |
| | PLSCR4 | | | TWSG1 |
| | PLXNA1 | | | AL096711.2 |
| | PLXNA2 | | | C2CD4B |
| | PLXNA3 | | | FAM193B |
| | PLXNB1 | | | AC092803.1 |
| | PLXNB2 | | | MIR3685 |
| | PLXNC1 | | | COLEC11 |
| | PLXND1 | | | AC131011.2 |
| | PMEPA1 | | | PDK2 |
| | PML | | | UBE2S |
| | PMPCA | | | TCAP |
| | PMPCB | | | DXO |
| | PMS1 | | | CAPN12 |
| | PMS2CL | | | PSMD1 |
| | PMS2P6 | | | MAL2 |
| | PMS2P7 | | | RSL1D1 |
| | PMVK | | | MIR3142HG |
| | PNISR | | | PLPP3 |
| | PNLDC1 | | | DARS1 |
| | PNN | | | DIS3L |
| | PNP | | | CHMP4C |
| | PNPLA2 | | | TIMM13 |
| | PNPLA4 | | | KANK2 |
| | PNPLA6 | | | DNAH5 |
| | PNPO | | | MICOS13 |
| | PNPT1 | | | TMEM87B |
| | PNRC1 | | | SETD1A |
| | PODXL | | | PRADC1 |
| | POF1B | | | EMC4 |
| | POFUT2 | | | MIF4GD |
| | POGLUT1 | | | HNRNPH3 |
| | POGZ | | | PSMB6 |
| | POLA1 | | | TNKS1BP1 |
| | POLA2 | | | NUDT16L1 |
| | POLB | | | ZNF529 |
| | POLD2 | | | ACVR1 |

| | | | | |
|---|---|---|---|---|
| | POLDIP3 | | | CDC42BPG |
| | POLE | | | GRAMD1B |
| | POLG | | | WASHC4 |
| | POLM | | | MCFD2 |
| | POLR1A | | | ZBTB39 |
| | POLR1D | | | DCP1B |
| | POLR1E | | | ANKRD45 |
| | POLR2B | | | NPL |
| | POLR2D | | | IGSF3 |
| | POLR2E | | | DHX34 |
| | POLR2G | | | AL355987.3 |
| | POLR2K | | | SNORA74B |
| | POLR2L | | | HMG20A |
| | POLR2M | | | PRDM6 |
| | POLR3C | | | PITPNM3 |
| | POLR3GL | | | SPINT2 |
| | POM121 | | | MGMT |
| | POM121C | | | ACP1 |
| | POMGNT1 | | | MAPK1 |
| | POMT1 | | | AL358472.5 |
| | POMT2 | | | ALDH16A1 |
| | POMZP3 | | | PLXNA1 |
| | POP4 | | | SERPINB6 |
| | POP5 | | | GPRC5B |
| | POP7 | | | ITGB4 |
| | POPDC3 | | | COP1 |
| | POR | | | AP000944.4 |
| | POU2AF1 | | | L3MBTL3 |
| | POU2F2 | | | CRACD |
| | POU6F1 | | | CCDC157 |
| | PPA1 | | | STX7 |
| | PPARA | | | CLN5 |
| | PPARD | | | CDC123 |
| | PPARG | | | RBFOX2 |
| | PPARGC1B | | | ABCA3 |
| | PPDPF | | | SLC8B1 |
| | PPFIA1 | | | APEX1 |
| | PPFIA4 | | | FAM86C2P |
| | PPFIBP2 | | | FAM210B |
| | PPHLN1 | | | AC011939.2 |
| | PPIA | | | LSM11 |
| | PPIB | | | RIC1 |
| | PPID | | | AL450992.2 |
| | PPIE | | | SIX1 |
| | PPIG | | | ZBTB32 |
| | PPIH | | | SPTB |
| | PPIL1 | | | AL645939.2 |
| | PPIL4 | | | SLC49A4 |
| | PPL | | | LINC01215 |
| | PPM1A | | | KIF26B |
| | PPM1F | | | ID4 |
| | PPM1H | | | MTCO3P11 |
| | PPM1K | | | LPXN |
| | PPM1M | | | TMEM106A |
| | PPME1 | | | TMEM14A |
| | PPOX | | | TINAGL1 |
| | PPP1CA | | | ASH1L-AS1 |
| | PPP1R10 | | | GRM8 |
| | PPP1R11 | | | ZNF394 |
| | PPP1R12A | | | MTIF3 |

| | | | | |
|---|---|---|---|---|
| | PPP1R12B | | | PPP2R5B |
| | PPP1R12C | | | RARS2 |
| | PPP1R13B | | | SIAH1 |
| | PPP1R14A | | | MUC5AC |
| | PPP1R14B | | | HES1 |
| | PPP1R14B-AS1 | | | KCNQ3 |
| | PPP1R15A | | | AC108749.1 |
| | PPP1R16B | | | MTFR1L |
| | PPP1R18 | | | CIB2 |
| | PPP1R2 | | | C5orf63 |
| | PPP1R3C | | | VCPKMT |
| | PPP1R8 | | | TRAK1 |
| | PPP1R9A | | | BEST3 |
| | PPP1R9B | | | ZZEF1 |
| | PPP2CA | | | MTMR2 |
| | PPP2R2C | | | GNAQ |
| | PPP2R2D | | | AL163051.1 |
| | PPP2R3C | | | RAD52 |
| | PPP2R5B | | | FXYD5 |
| | PPP2R5D | | | RAD54L2 |
| | PPP3CC | | | GNG10 |
| | PPP3R1 | | | ACADSB |
| | PPP4C | | | POLR2J |
| | PPP4R1 | | | KBTBD2 |
| | PPP4R2 | | | SLC25A20 |
| | PPP4R3A | | | TRIM15 |
| | PPP4R3B | | | SETX |
| | PPP5C | | | CDK5RAP2 |
| | PPP6C | | | PIR |
| | PPP6R2 | | | SNX25 |
| | PPP6R3 | | | SMIM20 |
| | PPRC1 | | | DOCK7 |
| | PPTC7 | | | TMEM126B |
| | PQBP1 | | | MTCL1 |
| | PRADC1 | | | AC242842.3 |
| | PRAF2 | | | SDHC |
| | PRAME | | | EFL1P1 |
| | PRDM1 | | | PLXNB1 |
| | PRDM11 | | | SERPINA1 |
| | PRDM15 | | | MCM4 |
| | PRDM2 | | | PTOV1 |
| | PRDM4 | | | GEMIN8 |
| | PRDX1 | | | TRAPPC6B |
| | PRDX2 | | | KIF13B |
| | PRDX5 | | | SDHD |
| | PRDX6 | | | PWP1 |
| | PREB | | | ILF2 |
| | PRELID1 | | | CORO1B |
| | PRELID2 | | | AC005072.1 |
| | PREPL | | | SLC9A2 |
| | PRICKLE2 | | | FAM126A |
| | PRKAB1 | | | AL161421.1 |
| | PRKACB | | | AC008517.1 |
| | PRKAG2 | | | RPSA |
| | PRKAR1B | | | AGBL2 |
| | PRKAR2A | | | TBC1D2B |
| | PRKCA | | | CDCA8 |
| | PRKCE | | | NME2 |
| | PRKCZ | | | MIF |
| | PRKD1 | | | MAN2A1 |

| | | | | |
|---|---|---|---|---|
| | PRKRIP1 | | | TAOK1 |
| | PRKX | | | PDE3B |
| | PRKY | | | RRP7A |
| | PRMT1 | | | AP000944.2 |
| | PRMT2 | | | NUDCD3 |
| | PRMT5 | | | CALB2 |
| | PRMT7 | | | NUBPL |
| | PRMT9 | | | TAF1C |
| | PRPF19 | | | SCRN1 |
| | PRPF31 | | | ABHD15 |
| | PRPF38A | | | GPX4 |
| | PRPF38B | | | NUDCD2 |
| | PRPF39 | | | ACTR6 |
| | PRPF4 | | | ST7-OT4 |
| | PRPF4B | | | SSR2 |
| | PRPF6 | | | PACSIN3 |
| | PRPF8 | | | CYS1 |
| | PRPS1 | | | ATAD2 |
| | PRPS2 | | | ZBTB42 |
| | PRPSAP2 | | | AAMP |
| | PRR13 | | | AC083973.1 |
| | PRR13P5 | | | FLOT2 |
| | PRR14 | | | JAZF1-AS1 |
| | PRR14L | | | REEP1 |
| | PRR3 | | | AC090844.3 |
| | PRRC1 | | | SZRD1 |
| | PRRC2A | | | DHRS3 |
| | PRRG1 | | | MARF1 |
| | PRRG4 | | | AL138963.2 |
| | PRSS16 | | | AL590068.3 |
| | PRSS23 | | | GSTO2 |
| | PRSS8 | | | PDP1 |
| | PRTG | | | RRM2 |
| | PRX | | | AP001372.2 |
| | PRXL2B | | | ZNHIT2 |
| | PSAP | | | B4GALT6 |
| | PSAT1 | | | FAM126B |
| | PSEN1 | | | CREG1 |
| | PSEN2 | | | AC010336.8 |
| | PSENEN | | | TRAF3 |
| | PSMA4 | | | STAT5A |
| | PSMA7 | | | EIF2B5 |
| | PSMB1 | | | SCRIB |
| | PSMB10 | | | NBDY |
| | PSMB2 | | | FOXO3 |
| | PSMB3 | | | AC008870.3 |
| | PSMB4 | | | CLK2 |
| | PSMB5 | | | EVI5 |
| | PSMB7 | | | SERPINE1 |
| | PSMB8 | | | GTSF1 |
| | PSMB9 | | | UBE4A |
| | PSMC1 | | | ARHGEF39 |
| | PSMC1P1 | | | B4GALNT3 |
| | PSMC3 | | | BAP1 |
| | PSMC4 | | | KIF1A |
| | PSMC5 | | | CCNH |
| | PSMC6 | | | WASH3P |
| | PSMD1 | | | TIMM21 |
| | PSMD11 | | | SLC20A1 |
| | PSMD13 | | | CTDSP1 |

| | | | | |
|---|---|---|---|---|
| | PSMD2 | | | ZNF783 |
| | PSMD3 | | | VAMP2 |
| | PSMD4 | | | SUPT16H |
| | PSMD5 | | | PALM2AKAP2 |
| | PSMD6 | | | TOP3A |
| | PSMD7 | | | AC092120.1 |
| | PSMD8 | | | FAM102B |
| | PSME3 | | | S100A10 |
| | PSMF1 | | | CCNY |
| | PSMG2 | | | AP000845.1 |
| | PSPC1-AS2 | | | AKAP6 |
| | PSTPIP2 | | | LINC01126 |
| | PTAFR | | | C14orf132 |
| | PTBP1 | | | COQ9 |
| | PTBP2 | | | FBXL6 |
| | PTBP3 | | | BMP8B |
| | PTCD3 | | | ICA1 |
| | PTDSS2 | | | ASRGL1 |
| | PTEN | | | TBL1X |
| | PTENP1 | | | C19orf54 |
| | PTGER2 | | | ACADVL |
| | PTGIS | | | NF2 |
| | PTGR1 | | | INSL4 |
| | PTH1R | | | YY1AP1 |
| | PTK2 | | | H2AC11 |
| | PTK2B | | | USP21 |
| | PTK6 | | | CLDND1 |
| | PTK7 | | | FEZ2 |
| | PTMAP5 | | | LHFPL2 |
| | PTMS | | | CLDN12 |
| | PTOV1 | | | ZNF605 |
| | PTP4A1 | | | RABGAP1 |
| | PTP4A3 | | | CHST10 |
| | PTPA | | | TOMM5 |
| | PTPDC1 | | | TMEM53 |
| | PTPN1 | | | AL118516.1 |
| | PTPN13 | | | ANKHD1-EIF4EBP3 |
| | PTPN2 | | | MCTP1 |
| | PTPN21 | | | KDELR3 |
| | PTPN3 | | | NUP43 |
| | PTPN6 | | | TRMT2B |
| | PTPN9 | | | C19orf53 |
| | PTPRB | | | AC147651.1 |
| | PTPRC | | | ME2P1 |
| | PTPRD | | | AL359878.1 |
| | PTPRE | | | HEATR5B |
| | PTPRG | | | TRPV4 |
| | PTPRK | | | TRMT44 |
| | PTPRM | | | AL161454.1 |
| | PTPRN2 | | | NAE1 |
| | PTPRZ1 | | | DHTKD1 |
| | PTS | | | SH3YL1 |
| | PTTG1IP | | | PTCD3 |
| | PUDP | | | RNVU1-30 |
| | PUF60 | | | RNU6-33P |
| | PUM2 | | | TPRG1L |
| | PUM3 | | | MEF2A |
| | PURPL | | | AL354920.1 |
| | PUS1 | | | HNRNPUL2-BSCL2 |
| | PVR | | | SLC37A4 |

| | | | | |
|---|---|---|---|---|
| | PWP2 | | | CSNK1A1 |
| | PWWP2A | | | SNTA1 |
| | PWWP3A | | | GABPB1 |
| | PWWP3B | | | SRP9P1 |
| | PXDC1 | | | SURF1 |
| | PXDN | | | ZNF519 |
| | PXK | | | TP53RK |
| | PXN | | | ABCB1 |
| | PYCR2 | | | RPL7P9 |
| | PYGB | | | SMG5 |
| | PYGL | | | PRUNE2 |
| | PYGO2 | | | IFNA8 |
| | PYROXD1 | | | HRC |
| | QARS1 | | | CEP112 |
| | QKI | | | MIR3190 |
| | QPCT | | | AP000311.1 |
| | QPCTL | | | AC116366.2 |
| | QRICH1 | | | AC002377.1 |
| | QSER1 | | | CDKL5 |
| | QSOX1 | | | GNAI3 |
| | QSOX2 | | | RPS29P14 |
| | QTRT1 | | | AC092958.3 |
| | R3HCC1 | | | PPP1R3B |
| | R3HDM2 | | | BAD |
| | R3HDM4 | | | LEPR |
| | RAB10 | | | CSGALNACT1 |
| | RAB11B | | | PKD1L1 |
| | RAB11FIP1 | | | BRIP1 |
| | RAB11FIP3 | | | PEX13 |
| | RAB11FIP5 | | | AP002992.1 |
| | RAB13 | | | TMEM256 |
| | RAB15 | | | IL17RD |
| | RAB17 | | | ACAD9 |
| | RAB1B | | | SLC35E2B |
| | RAB20 | | | MPST |
| | RAB21 | | | SLC6A13 |
| | RAB24 | | | MRPS21 |
| | RAB25 | | | BLM |
| | RAB27A | | | SAFB |
| | RAB29 | | | NSFL1C |
| | RAB2B | | | MIA |
| | RAB30 | | | LINC00326 |
| | RAB32 | | | TRPA1 |
| | RAB33B | | | GRIK5 |
| | RAB34 | | | PDE4D |
| | RAB35 | | | SMIM19 |
| | RAB3GAP2 | | | STX16-NPEPL1 |
| | RAB40B | | | MIR5047 |
| | RAB40C | | | ZNF185 |
| | RAB4A | | | TLCD2 |
| | RAB4B | | | ACOT9 |
| | RAB5C | | | STK17A |
| | RAB6A | | | RPSAP58 |
| | RAB6C | | | CHID1 |
| | RAB7A | | | HSPE1P11 |
| | RAB7B | | | ATG12 |
| | RAB8A | | | CCDC69 |
| | RAB8B | | | MOCS3 |
| | RABAC1 | | | SNORA28 |
| | RABGAP1 | | | MAML2 |

| | | | | |
|---|---|---|---|---|
| | RABGAP1L | | | LINC01271 |
| | RABGGTA | | | AL731571.1 |
| | RABIF | | | MTOR |
| | RABL2A | | | PLEKHG3 |
| | RABL3 | | | DPH2 |
| | RABL6 | | | C1QTNF1 |
| | RAC2 | | | CRKL |
| | RACK1 | | | AL137781.1 |
| | RAD18 | | | AL160408.2 |
| | RAD23A | | | AP001062.2 |
| | RAD54L2 | | | FAM83F |
| | RAE1 | | | AP1S3 |
| | RAET1G | | | ANKMY1 |
| | RAET1L | | | AC067968.1 |
| | RAF1 | | | LGALS1 |
| | RAI2 | | | ZNF862 |
| | RALB | | | PEX2 |
| | RALBP1 | | | H2AC16 |
| | RALGAPA2 | | | LYPD3 |
| | RALGAPB | | | TRIP12 |
| | RALGDS | | | B4GALT3 |
| | RALGPS2 | | | MYL5 |
| | RALY | | | RSRP1 |
| | RAN | | | AC004264.1 |
| | RANBP1 | | | ART3 |
| | RANBP10 | | | STEAP2 |
| | RANBP3 | | | PTMS |
| | RANBP9 | | | SGMS1-AS1 |
| | RANGAP1 | | | TP53INP2 |
| | RAP1A | | | PTMAP2 |
| | RAP2C | | | LRRCC1 |
| | RAPGEF1 | | | DNAL1 |
| | RAPGEF2 | | | PPCDC |
| | RAPGEF3 | | | UCP2 |
| | RAPGEF5 | | | P4HTM |
| | RAPGEFL1 | | | EIF2B1 |
| | RASA1 | | | AC092868.3 |
| | RASA3 | | | AGAP1 |
| | RASA4 | | | WNK3 |
| | RASA4B | | | EIF2AK3 |
| | RASD2 | | | TMEM254 |
| | RASGEF1B | | | NAGA |
| | RASGRF2 | | | CEP63 |
| | RASIP1 | | | GSE1 |
| | RASL11A | | | FEZ1 |
| | RASL12 | | | ABCE1 |
| | RASSF1 | | | PNKD |
| | RASSF10 | | | PIK3C3 |
| | RASSF2 | | | YKT6 |
| | RASSF7 | | | GRK3 |
| | RBBP4 | | | NUTF2 |
| | RBBP6 | | | ANXA2R |
| | RBBP8 | | | CC2D2A |
| | RBBP9 | | | AC007780.1 |
| | RBCK1 | | | ZBTB41 |
| | RBFOX2 | | | FAM189B |
| | RBM10 | | | MPHOSPH8 |
| | RBM11 | | | NCL |
| | RBM12 | | | HAUS1 |
| | RBM14 | | | HENMT1 |

| | | | | |
|---|---|---|---|---|
| | RBM15B | | | UQCRC2 |
| | RBM17 | | | SCOC |
| | RBM19 | | | ZC3H3 |
| | RBM22 | | | PSMD11 |
| | RBM26 | | | PNRC2 |
| | RBM28 | | | RBM8A |
| | RBM3 | | | AC011448.1 |
| | RBM33 | | | LINC01285 |
| | RBM34 | | | GOLPH3 |
| | RBM39 | | | TRPC1 |
| | RBM43 | | | ENTPD3 |
| | RBM47 | | | TTI2 |
| | RBM5 | | | DUS3L |
| | RBM6 | | | PHF12 |
| | RBM7 | | | PLD3 |
| | RBMS1 | | | UQCR11 |
| | RBMX2 | | | QPCT |
| | RBMXL1 | | | COX19 |
| | RBP4 | | | C8orf31 |
| | RBPMS | | | LRIG2 |
| | RBSN | | | MCM8 |
| | RBX1 | | | CNKSR2 |
| | RC3H2 | | | NUPR1 |
| | RCAN1 | | | RFPL4A |
| | RCBTB1 | | | AC019117.3 |
| | RCBTB2 | | | AL035460.1 |
| | RCC1L | | | LDC1P |
| | RCC2 | | | AC009949.1 |
| | RCE1 | | | GMDS |
| | RCOR1 | | | NFATC2 |
| | RDH10 | | | MMP15 |
| | RDH11 | | | MZT2A |
| | RDH14 | | | SNHG25 |
| | RDM1 | | | NEK7 |
| | RDX | | | TM9SF2 |
| | REC8 | | | SPAST |
| | RECK | | | RNVU1-25 |
| | RECQL4 | | | DUT |
| | REL | | | CDK11B |
| | RENBP | | | COPRS |
| | REPS2 | | | COX4I1 |
| | RERE | | | COPS5 |
| | RESF1 | | | BLOC1S1 |
| | REST | | | MT1X |
| | RETREG1 | | | EMB |
| | RETREG3 | | | POMGNT1 |
| | RETSAT | | | NDFIP1 |
| | REV1 | | | SLC25A10 |
| | REX1BD | | | NEK10 |
| | REXO4 | | | BTBD2 |
| | RFC1 | | | IGHE |
| | RFLNB | | | SDSL |
| | RFNG | | | AP1S2 |
| | RFT1 | | | GLG1 |
| | RFTN1 | | | FOXP2 |
| | RFX1 | | | ANKHD1 |
| | RFX7 | | | ARHGAP17 |
| | RFXANK | | | PCNA |
| | RGCC | | | TJP3 |
| | RGL1 | | | SETD1B |

| | | | | |
|---|---|---|---|---|
| | RGL2 | | | PTN |
| | RGL4 | | | RBM25 |
| | RGMB | | | PRPS1 |
| | RGP1 | | | OMA1 |
| | RGPD1 | | | GPC1 |
| | RGPD2 | | | DHX29 |
| | RGPD3 | | | LINC01534 |
| | RGPD5 | | | MRPS34 |
| | RGPD6 | | | RASSF4 |
| | RGPD8 | | | LIPG |
| | RGS12 | | | RASSF6 |
| | RGS18 | | | RALGPS1 |
| | RGS2 | | | NFATC3 |
| | RGS3 | | | IGBP1 |
| | RGS4 | | | ETNK1 |
| | RGSL1 | | | CD99P1 |
| | RHBDD1 | | | RNA5S9 |
| | RHBDD2 | | | ZNF720 |
| | RHBDF1 | | | TCF7L2 |
| | RHBDF2 | | | PLEKHN1 |
| | RHOA | | | ERO1A |
| | RHOB | | | NBPF26 |
| | RHOC | | | TOR3A |
| | RHOG | | | TPR |
| | RHOH | | | CACUL1 |
| | RHOJ | | | AC105219.1 |
| | RIBC1 | | | DDX42 |
| | RIC1 | | | MORN2 |
| | RIC8B | | | AC025431.1 |
| | RICTOR | | | AC007032.1 |
| | RILPL1 | | | DOK4 |
| | RIMKLB | | | PPOX |
| | RIMS2 | | | EMSY |
| | RIMS3 | | | SPR |
| | RIN2 | | | PGF |
| | RING1 | | | MAK |
| | RIOK2 | | | TOMM70 |
| | RIOK3 | | | ECPAS |
| | RIPK1 | | | RANBP3 |
| | RIPK2 | | | RPAIN |
| | RIPOR1 | | | AP003419.1 |
| | RIPOR2 | | | TMEM38B |
| | RIPOR3 | | | UBR5 |
| | RIPPLY3 | | | NDUFB9 |
| | RLF | | | VPS39 |
| | RLIM | | | GARS1 |
| | RMDN1 | | | SKA3 |
| | RMDN3 | | | ARID1B |
| | RMND1 | | | CENPM |
| | RMND5A | | | DDX50 |
| | RMND5B | | | IL20RB |
| | RN7SL396P | | | CCT2 |
| | RN7SL689P | | | LRFN1 |
| | RNA5S9 | | | ST6GALNAC2 |
| | RNA5SP141 | | | LRRC41 |
| | RNA5SP145 | | | CHKB |
| | RNA5SP225 | | | ENTPD6 |
| | RNA5SP298 | | | ZNF827 |
| | RNA5SP481 | | | SPC24 |
| | RNASE2 | | | TCEA3 |

| | | | | |
|---|---|---|---|---|
| | RNASEH1 | | | URB2 |
| | RNASEH2B | | | ZMYM1 |
| | RNASEH2C | | | UBE2G1 |
| | RNASEL | | | SIDT1 |
| | RND1 | | | DLG1 |
| | RND3 | | | LRCH4 |
| | RNF113A | | | RAP1GDS1 |
| | RNF121 | | | SYNM |
| | RNF122 | | | DDX24 |
| | RNF13 | | | NDUFB8 |
| | RNF139 | | | LURAP1L-AS1 |
| | RNF144A | | | AC034236.1 |
| | RNF144B | | | NPM3 |
| | RNF145 | | | CHRNA7 |
| | RNF149 | | | CASR |
| | RNF152 | | | MARC1 |
| | RNF166 | | | TSPAN18 |
| | RNF167 | | | NLRP4 |
| | RNF168 | | | OR52K1 |
| | RNF170 | | | AC092168.1 |
| | RNF175 | | | AC022092.1 |
| | RNF185 | | | HSPB9 |
| | RNF19B | | | ADPGK-AS1 |
| | RNF20 | | | AC010501.2 |
| | RNF213 | | | AC011369.2 |
| | RNF216 | | | ARMC10P1 |
| | RNF216P1 | | | CAVIN4 |
| | RNF220 | | | POP5 |
| | RNF224 | | | ZNF594 |
| | RNF227 | | | ZHX3 |
| | RNF25 | | | ANAPC15 |
| | RNF34 | | | BMS1P4 |
| | RNF4 | | | AC015813.2 |
| | RNF40 | | | GRM5 |
| | RNF44 | | | ERCC6L |
| | RNF5 | | | TIMM8B |
| | RNF8 | | | LONRF3 |
| | RNGTT | | | UBR3 |
| | RNH1 | | | SNORA47 |
| | RNMT | | | HMGB1P3 |
| | RNPEP | | | CNTF |
| | RNPS1 | | | TMEM65 |
| | RNU4-1 | | | STK24-AS1 |
| | RNU4ATAC | | | DCUN1D4 |
| | RNU5A-1 | | | NANS |
| | RNU6-1 | | | LINC00240 |
| | RNU6-2 | | | SNORC |
| | RNU6-36P | | | FANCF |
| | RNU6-37P | | | HABP2 |
| | RNU6-5P | | | ALKBH5 |
| | RNU6-9 | | | AL034430.1 |
| | RNU6ATAC | | | MBOAT7 |
| | RNVU1-28 | | | IL17RC |
| | RNVU1-2A | | | PQBP1 |
| | RNVU1-31 | | | SOX13 |
| | ROBO3 | | | ZNF664 |
| | ROBO4 | | | RPS26 |
| | ROCK1 | | | ZNF777 |
| | ROCK2 | | | RPL23AP82 |
| | ROMO1 | | | CSNK1G1 |

| | | | | |
|---|---|---|---|---|
| | RORA | | | WNK1 |
| | RP11-173M1.8 | | | SNORD15B |
| | RP11-175O19.4 | | | LINC02035 |
| | RP11-206L10.3 | | | IPO9 |
| | RP11-211G3.2 | | | PHLDA2 |
| | RP11-401P9.4 | | | COPB1 |
| | RP11-435O5.2 | | | AC120024.1 |
| | RP11-474N8.8 | | | SUPT20H |
| | RP11-485G4.2 | | | AC090114.3 |
| | RP11-680G24.5 | | | TBC1D9B |
| | RP1-168P16.2 | | | CCL22 |
| | RP11-706O15.3 | | | LINC00511 |
| | RP11-706O15.5 | | | EXOC6B |
| | RP1-178F10.3 | | | AC019186.1 |
| | RP11-872J21.5 | | | CSF2RA |
| | RP11-89K10.1 | | | SLC39A1 |
| | RP1-71H24.6 | | | YES1 |
| | RP2 | | | SEC11A |
| | RP3-323A16.1 | | | RPUSD1 |
| | RP3-368A4.5 | | | MORC3 |
| | RP5-1039K5.19 | | | CDC6 |
| | RP9P | | | NDUFA3 |
| | RPA1 | | | PROM2 |
| | RPA2 | | | CINP |
| | RPA3 | | | TOX4 |
| | RPAIN | | | VPS9D1-AS1 |
| | RPAP2 | | | ENSA |
| | RPAP3 | | | AC016405.1 |
| | RPGR | | | C2-AS1 |
| | RPH3A | | | MAP2K5 |
| | RPH3AL | | | LINC00973 |
| | RPIA | | | C1orf198 |
| | RPL10 | | | FASTKD1 |
| | RPL10A | | | CDC42BPB |
| | RPL10P16 | | | EIF3H |
| | RPL10P3 | | | STN1 |
| | RPL10P6 | | | CROT |
| | RPL12 | | | PINK1 |
| | RPL12P4 | | | KIAA0408 |
| | RPL13 | | | RFC4 |
| | RPL13A | | | LRRC75B |
| | RPL13AP20 | | | ARRB1 |
| | RPL13AP25 | | | WDR46 |
| | RPL13AP5 | | | SEPTIN4 |
| | RPL13AP7 | | | CYP21A2 |
| | RPL14 | | | HSPA14 |
| | RPL15 | | | AL390719.1 |
| | RPL15P2 | | | FGF18 |
| | RPL18 | | | MTMR11 |
| | RPL18A | | | CDR2 |
| | RPL19 | | | TRIM34 |
| | RPL21 | | | SCAND2P |
| | RPL22P1 | | | PJA1 |
| | RPL23AP53 | | | TWNK |
| | RPL23AP61 | | | ZNF574 |
| | RPL24 | | | MED12 |
| | RPL26 | | | ALAS1 |
| | RPL26P19 | | | MCUR1 |
| | RPL3 | | | UBB |
| | RPL31 | | | AC005258.1 |

| | | | | |
|---|---|---|---|---|
| | RPL31P63 | | | MAT2A |
| | RPL32 | | | TBC1D4 |
| | RPL32P3 | | | DAD1 |
| | RPL34 | | | COLEC10 |
| | RPL34P18 | | | TACC2 |
| | RPL35A | | | CDS1 |
| | RPL36 | | | EPS15 |
| | RPL36AL | | | NIBAN1 |
| | RPL36AP26 | | | CDH2 |
| | RPL37P2 | | | CPSF1 |
| | RPL38 | | | THRB-IT1 |
| | RPL39 | | | CCDC124 |
| | RPL39P3 | | | UGT2A3 |
| | RPL4 | | | LINC01002 |
| | RPL41P2 | | | ZFHX4 |
| | RPL5 | | | RBM7 |
| | RPL6 | | | ZNF281 |
| | RPL6P27 | | | FBXW7 |
| | RPL7 | | | ZNF649 |
| | RPL7A | | | USP46 |
| | RPL7AP66 | | | TCF3P1 |
| | RPL7P6 | | | CIAPIN1 |
| | RPL7P9 | | | H2AC5P |
| | RPL8 | | | AL355377.1 |
| | RPL9 | | | SERGEF |
| | RPL9P29 | | | BDNF-AS |
| | RPL9P7 | | | CREB3L4 |
| | RPLP0 | | | NPC1L1 |
| | RPLP1 | | | MIIP |
| | RPLP1P6 | | | CYP20A1 |
| | RPLP2 | | | DERL1 |
| | RPN1 | | | CFAP298 |
| | RPP25 | | | ZGPAT |
| | RPRD1A | | | CALCA |
| | RPRD1B | | | CAPZA1 |
| | RPRD2 | | | ZFYVE27 |
| | RPS10 | | | ALG5 |
| | RPS11P5 | | | ZFP41 |
| | RPS13 | | | RIPOR1 |
| | RPS13P2 | | | PLSCR2 |
| | RPS14 | | | LYPLA1 |
| | RPS15A | | | LINC01089 |
| | RPS15AP1 | | | ST6GALNAC1 |
| | RPS16 | | | MRPS5 |
| | RPS18 | | | AC068338.2 |
| | RPS19 | | | APOM |
| | RPS19BP1 | | | TMED8 |
| | RPS23 | | | ZC3H12D |
| | RPS23P8 | | | AL805961.1 |
| | RPS24 | | | ANKRD36C |
| | RPS24P8 | | | AC104472.1 |
| | RPS26P11 | | | CLASRP |
| | RPS26P15 | | | FADD |
| | RPS26P31 | | | DMTF1 |
| | RPS26P47 | | | COQ10B |
| | RPS27 | | | MAP7 |
| | RPS27A | | | CD163 |
| | RPS27P29 | | | CREB3L2 |
| | RPS28 | | | TOMM7 |
| | RPS29 | | | TMSB4XP8 |

| | | | | |
|---|---|---|---|---|
| | RPS2P35 | | | ZSCAN20 |
| | RPS2P5 | | | CMTM7 |
| | RPS2P55 | | | SSH3 |
| | RPS3 | | | LAPTM4A |
| | RPS3A | | | ABLIM3 |
| | RPS5 | | | RALGAPA2 |
| | RPS6 | | | THAP1 |
| | RPS6KA2 | | | TBC1D9 |
| | RPS6KA5 | | | HTR3E |
| | RPS6KA6 | | | WDR5B |
| | RPS6KB1 | | | ADAMTSL2 |
| | RPS6KB2 | | | TAF9B |
| | RPS6KC1 | | | ZBTB33 |
| | RPS7P1 | | | TICRR |
| | RPS7P10 | | | AC067945.1 |
| | RPS7P11 | | | ANGPTL1 |
| | RPS8 | | | SYNJ2BP |
| | RPS9 | | | RN7SKP9 |
| | RPSA | | | TMEM236 |
| | RPSAP12 | | | ARL15 |
| | RPSAP4 | | | FZD1 |
| | RPSAP47 | | | RANBP6 |
| | RPSAP54 | | | EPHB4 |
| | RPTOR | | | TXN2 |
| | RPUSD2 | | | SYNRG |
| | RRAD | | | AC073508.2 |
| | RRAS | | | BBS2 |
| | RRAS2 | | | TOMM6 |
| | RRBP1 | | | LINC01467 |
| | RREB1 | | | AJ003147.1 |
| | RRM2 | | | CCL4 |
| | RRN3 | | | SGMS2 |
| | RRP1 | | | DDX47 |
| | RRP12 | | | CEP41 |
| | RRP36 | | | ZSWIM4 |
| | RRP7A | | | NDUFB11 |
| | RRP7BP | | | AF127936.1 |
| | RRP9 | | | EXD2 |
| | RSAD1 | | | BACE1-AS |
| | RSAD2 | | | CLOCK |
| | RSBN1L | | | GGT1 |
| | RSF1 | | | HYKK |
| | RSPO3 | | | TTPAL |
| | RSPRY1 | | | PHF10 |
| | RSRC2 | | | KCNS1 |
| | RTCB | | | SLC25A36 |
| | RTF1 | | | LINC01226 |
| | RTKN | | | LITAF |
| | RTKN2 | | | BICD1 |
| | RTL10 | | | EID2B |
| | RTL5 | | | FADS3 |
| | RTL8A | | | SPDYA |
| | RTN3 | | | GALNT14 |
| | RTN4 | | | DKK3 |
| | RTP4 | | | PYCR1 |
| | RUBCN | | | KIF13A |
| | RUFY2 | | | ZBED9 |
| | RUFY3 | | | LIMD2 |
| | RUNDC3A-AS1 | | | UNC13B |
| | RUSC2 | | | NUP107 |

| | | | | |
|---|---|---|---|---|
| | RUVBL2 | | | CLCN5 |
| | RWDD4 | | | CISD2 |
| | RXRA | | | EI24 |
| | RXRB | | | PRSS8 |
| | RYK | | | SLC7A5P1 |
| | S100A11 | | | SPATA6 |
| | S100A12 | | | SF3B5 |
| | S100A13 | | | INO80B-WBP1 |
| | S100A14 | | | SNORA63B |
| | S100A16 | | | LMBRD1 |
| | S100A2 | | | MLX |
| | S100A8 | | | MAPK8IP2 |
| | S100A9 | | | NSMCE1 |
| | S100P | | | VEGFB |
| | S100PBP | | | TAGLN2P1 |
| | SAA1 | | | AC037459.2 |
| | SAA2 | | | PHF20 |
| | SACS | | | SNX30 |
| | SAE1 | | | CTSZ |
| | SAFB | | | KLHL15 |
| | SAFB2 | | | SNX11 |
| | SAMD4A | | | ABCA12 |
| | SAMD9 | | | TTLL5 |
| | SAMD9L | | | ING1 |
| | SAMHD1 | | | ZNF202 |
| | SAMM50 | | | KIF22 |
| | SAP18 | | | AC092073.1 |
| | SAP30 | | | ERCC6 |
| | SAP30BP | | | CYREN |
| | SAP30L | | | UPF2 |
| | SAR1B | | | SCYL1 |
| | SARAF | | | FAM81A |
| | SARS1 | | | AC008581.2 |
| | SART1 | | | ICAM3 |
| | SASH1 | | | VAMP7 |
| | SAT1 | | | SPTAN1 |
| | SBDSP1 | | | CASZ1 |
| | SBF1 | | | EIF1 |
| | SBNO1 | | | SNORA16A |
| | SC5D | | | SNORA16A |
| | SCAF11 | | | TMED2-DT |
| | SCAF4 | | | KRT8 |
| | SCAMP1-AS1 | | | NCF2 |
| | SCAMP3 | | | SNHG26 |
| | SCAND1 | | | MANSC1 |
| | SCAP | | | MYBPC1 |
| | SCARA3 | | | AL139100.2 |
| | SCARB1 | | | SNX6 |
| | SCARB2 | | | CDC20B |
| | SCD5 | | | AC139491.2 |
| | SCDP1 | | | AC006065.4 |
| | SCGB1A1 | | | C20orf141 |
| | SCLT1 | | | AC098934.4 |
| | SCMH1 | | | ZFP30 |
| | SCML1 | | | PARM1 |
| | SCN2A | | | AC092747.2 |
| | SCN4B | | | CSPG5 |
| | SCN9A | | | ADNP2 |
| | SCNN1B | | | DCUN1D3 |
| | SCNN1G | | | TMEM33 |

| | | | | |
|---|---|---|---|---|
| | SCO1 | | | SFXN2 |
| | SCOC-AS1 | | | CPAMD8 |
| | SCRN1 | | | CEP162 |
| | SCRN2 | | | AL365181.2 |
| | SCYL1 | | | AC005899.5 |
| | SCYL2 | | | ARFGAP2 |
| | SDAD1 | | | ZYG11B |
| | SDAD1P1 | | | EPDR1 |
| | SDC3 | | | BYSL |
| | SDCBP | | | MAP1S |
| | SDCCAG8 | | | H3C4 |
| | SDE2 | | | MSN |
| | SDF2L1 | | | ABTB1 |
| | SDHA | | | RNVU1-21 |
| | SDHAF1 | | | DTX3 |
| | SDHAF2 | | | AC012513.3 |
| | SDHAF3 | | | TMEM187 |
| | SDHAP1 | | | LONRF2 |
| | SDHB | | | INTS8 |
| | SDHD | | | ZFYVE1 |
| | SEC13 | | | SERTAD4 |
| | SEC16A | | | NOS3 |
| | SEC22B | | | LINC01003 |
| | SEC22C | | | RFPL2 |
| | SEC23A | | | MXI1 |
| | SEC23IP | | | OGA |
| | SEC24C | | | THNSL1 |
| | SEC31A | | | PLXNA4 |
| | SEC31B | | | GCNT3 |
| | SEC61A1 | | | C6orf223 |
| | SEC61A2 | | | NID1 |
| | SEC62 | | | ANAPC5 |
| | SEC63 | | | TNNC1 |
| | SECISBP2L | | | RTL10 |
| | SECTM1 | | | KRT10 |
| | SEL1L | | | SDHA |
| | SELENBP1 | | | LTBR |
| | SELENOH | | | WDR35 |
| | SELENOM | | | IMMP2L |
| | SELENON | | | SIRT5 |
| | SELENOO | | | MEAF6 |
| | SELENOS | | | TTC9 |
| | SELENOW | | | WWC3 |
| | SELL | | | AC004870.2 |
| | SELPLG | | | SNRPGP2 |
| | SEM1 | | | ZNF710-AS1 |
| | SEMA3A | | | GPR35 |
| | SEMA3B | | | TANGO6 |
| | SEMA3C | | | VASH1 |
| | SEMA3F | | | C7orf50 |
| | SEMA4A | | | CREB1 |
| | SEMA4B | | | RN7SKP175 |
| | SEMA4C | | | KPNA4 |
| | SEMA6B | | | ZNF488 |
| | SEMA6D | | | PAIP1P1 |
| | SENP3 | | | HCG25 |
| | SENP5 | | | VEGFD |
| | SENP6 | | | NDUFA12 |
| | SEPHS1 | | | ATXN7L2 |
| | SEPHS2 | | | RPS19BP1 |

| | | | | |
|---|---|---|---|---|
| | SEPTIN11 | | | SLC22A18AS |
| | SEPTIN4 | | | CNOT10 |
| | SEPTIN6 | | | INHBE |
| | SEPTIN7P2 | | | VXN |
| | SEPTIN8 | | | CBX2 |
| | SEPTIN9 | | | SGO1 |
| | SERAC1 | | | TIRAP |
| | SERBP1 | | | ZNF595 |
| | SERF2 | | | HMGN4 |
| | SERGEF | | | CCDC50 |
| | SERINC1 | | | AC073046.1 |
| | SERINC5 | | | ZNF195 |
| | SERP1 | | | GLIPR1 |
| | SERPINA1 | | | GPX2 |
| | SERPINA6 | | | BRPF3 |
| | SERPINB1 | | | AP001528.1 |
| | SERPINB2 | | | KRT8P12 |
| | SERPINB6 | | | SMIM25 |
| | SERPINB8 | | | PFKL |
| | SERPINB9 | | | CD14 |
| | SERPINB9P1 | | | ZBTB46 |
| | SERPINE2 | | | FIG4 |
| | SERPING1 | | | TUT1 |
| | SERTAD1 | | | ZNF491 |
| | SERTAD2 | | | RSPH3 |
| | SERTAD3 | | | TRPS1 |
| | SESN1 | | | FOXN2 |
| | SESN2 | | | RNU7-196P |
| | SETD1A | | | AC009242.1 |
| | SETD3 | | | VDAC2 |
| | SETD5 | | | PTPN13 |
| | SETD6 | | | CELF1 |
| | SETD7 | | | GPR83 |
| | SETDB1 | | | PACS2 |
| | SEZ6L2 | | | GALT |
| | SF1 | | | NTPCR |
| | SF3A2 | | | AC145207.5 |
| | SF3A3 | | | CCDC9B |
| | SF3B1 | | | B4GALNT2 |
| | SF3B2 | | | AMN1 |
| | SF3B4 | | | PRKACA |
| | SF3B5 | | | GSS |
| | SF3B6 | | | SMARCB1 |
| | SFI1 | | | SERPINA10 |
| | SFMBT1 | | | PSMC1P2 |
| | SFN | | | CSNK1G2P1 |
| | SFPQ | | | AC090673.1 |
| | SFR1 | | | AC097636.2 |
| | SFRP4 | | | NDEL1 |
| | SFSWAP | | | PGPEP1 |
| | SFT2D1 | | | SREK1IP1 |
| | SFTPA1 | | | ADGRG6 |
| | SFTPA2 | | | KCTD9 |
| | SFTPB | | | CT83 |
| | SFTPC | | | TALDO1 |
| | SFTPD | | | FANCL |
| | SFXN3 | | | LARP4B |
| | SFXN4 | | | FGD2 |
| | SFXN5 | | | SUPV3L1 |
| | SGCA | | | TRIM6-TRIM34 |

| | | | | |
|---|---|---|---|---|
| | SGCE | | | AC027309.2 |
| | SGIP1 | | | RNF14 |
| | SGK1 | | | GDAP1 |
| | SGMS1 | | | MAOA |
| | SGPL1 | | | GLB1L2 |
| | SGPP2 | | | REEP5 |
| | SGSH | | | ZBED4 |
| | SGSM2 | | | SH3GLB1 |
| | SGSM3 | | | BICD2 |
| | SGTA | | | AP002387.2 |
| | SH2B1 | | | TNPO3 |
| | SH2B3 | | | AC007849.1 |
| | SH3BGRL2 | | | MPV17L |
| | SH3BGRL3 | | | PREPL |
| | SH3BP2 | | | SCAF11 |
| | SH3BP5 | | | FJX1 |
| | SH3BP5-AS1 | | | ZSCAN25 |
| | SH3D19 | | | KPNA5 |
| | SH3GL1 | | | UBA3 |
| | SH3PXD2A | | | LINC02331 |
| | SH3PXD2B | | | RPL39 |
| | SH3RF1 | | | PRKCE |
| | SH3RF3 | | | HHEX |
| | SH3TC1 | | | MRPL34 |
| | SH3TC2 | | | TXN |
| | SH3YL1 | | | SYT1 |
| | SHANK3 | | | TCTN3 |
| | SHB | | | DGKG |
| | SHC1 | | | RBM4 |
| | SHC2 | | | DNALI1 |
| | SHE | | | PUM1 |
| | SHFL | | | TMEM263 |
| | SHISA5 | | | ZNF43 |
| | SHISA9 | | | SLC1A4 |
| | SHKBP1 | | | ITGA9-AS1 |
| | SHOC2 | | | PSMA1 |
| | SHQ1 | | | ITFG1 |
| | SIAH2 | | | ZNF205 |
| | SIDT2 | | | SMIM14 |
| | SIGLEC16 | | | CWC15 |
| | SIK1 | | | PRPF3 |
| | SIK1B | | | KIFC3 |
| | SIK2 | | | HERC3 |
| | SIL1 | | | NEDD4L |
| | SIMC1 | | | MMS19 |
| | SIN3A | | | AC022167.2 |
| | SIN3B | | | SPRED2 |
| | SIRPA | | | NUCB2 |
| | SIRPB1 | | | ZNF737 |
| | SIRT1 | | | AL139174.1 |
| | SIRT5 | | | LEPROTL1 |
| | SIRT7 | | | FIZ1 |
| | SIVA1 | | | LRRC37B |
| | SIX1 | | | RN7SL674P |
| | SKA2 | | | FAM53B |
| | SKIV2L | | | TLE3 |
| | SKP1 | | | SETMAR |
| | SLAIN2 | | | IRF8 |
| | SLAMF7 | | | AC093155.3 |
| | SLBP | | | DEPDC7 |

| | | | | |
|---|---|---|---|---|
| | SLC10A3 | | | PCMT1 |
| | SLC12A2 | | | SLU7 |
| | SLC12A4 | | | AC092279.2 |
| | SLC12A5 | | | CEBPG |
| | SLC12A6 | | | RPS19P1 |
| | SLC12A7 | | | RIT1 |
| | SLC13A1 | | | TP53TG1 |
| | SLC15A2 | | | UCK2 |
| | SLC16A4 | | | RPRD2 |
| | SLC16A5 | | | AKR7A2 |
| | SLC16A7 | | | WDR55 |
| | SLC18B1 | | | LINC00158 |
| | SLC19A2 | | | AC078923.1 |
| | SLC22A15 | | | TMED7 |
| | SLC22A17 | | | AC087190.3 |
| | SLC22A23 | | | TMEM109 |
| | SLC24A4 | | | RHOBTB3 |
| | SLC25A1 | | | TMEM120B |
| | SLC25A11 | | | CTSL |
| | SLC25A17 | | | HAT1 |
| | SLC25A19 | | | NDUFV2-AS1 |
| | SLC25A21 | | | SMAD6 |
| | SLC25A23 | | | RAB9A |
| | SLC25A25 | | | NCKAP5 |
| | SLC25A26 | | | SNF8 |
| | SLC25A28 | | | IREB2 |
| | SLC25A29 | | | TMEM218 |
| | SLC25A3 | | | KAT2B |
| | SLC25A30 | | | FLAD1 |
| | SLC25A33 | | | SLC38A10 |
| | SLC25A36 | | | RPL11 |
| | SLC25A4 | | | MSH2 |
| | SLC25A44 | | | CLDN4 |
| | SLC25A5 | | | AC104035.1 |
| | SLC25A6 | | | PRKAA1 |
| | SLC26A4 | | | IL23A |
| | SLC26A6 | | | C1GALT1C1 |
| | SLC26A8 | | | AL117339.5 |
| | SLC27A1 | | | DUSP14 |
| | SLC27A3 | | | AP2A2 |
| | SLC28A2 | | | WDR61 |
| | SLC29A1 | | | AARS1 |
| | SLC2A1 | | | RTN4 |
| | SLC30A1 | | | ANKRD36 |
| | SLC30A4 | | | ARHGAP12 |
| | SLC31A1 | | | TXNL4B |
| | SLC34A2 | | | LINC00205 |
| | SLC35A3 | | | ELOA |
| | SLC35A4 | | | RPS2P44 |
| | SLC35B2 | | | CLDN2 |
| | SLC35C2 | | | NRBP2 |
| | SLC35E1 | | | FGFR1OP2 |
| | SLC35F1 | | | FAM193A |
| | SLC35F2 | | | FCGRT |
| | SLC35F3 | | | RPSAP9 |
| | SLC35G2 | | | DST |
| | SLC37A4 | | | DYNC1LI1 |
| | SLC38A10 | | | PRIMPOL |
| | SLC38A2 | | | IFT81 |
| | SLC38A5 | | | AC091959.3 |

| | | | | |
|---|---|---|---|---|
| | SLC39A1 | | | AC016747.1 |
| | SLC39A13 | | | FIBP |
| | SLC39A3 | | | GNB5 |
| | SLC39A6 | | | AC138969.1 |
| | SLC39A7 | | | CHKA |
| | SLC39A8 | | | SSUH2 |
| | SLC40A1 | | | NBAS |
| | SLC41A1 | | | SNORD101 |
| | SLC41A2 | | | LIX1L |
| | SLC41A3 | | | CCRL2 |
| | SLC43A1 | | | SLC43A3 |
| | SLC44A1 | | | ZMYM3 |
| | SLC44A2 | | | KDM1B |
| | SLC44A3-AS1 | | | NOC3L |
| | SLC44A4 | | | CBX1P3 |
| | SLC48A1 | | | FDPSP4 |
| | SLC4A1AP | | | MIR3173 |
| | SLC4A7 | | | H3C5P |
| | SLC51A | | | PIGL |
| | SLC52A2 | | | DTWD2 |
| | SLC5A3 | | | ULBP3 |
| | SLC5A6 | | | PAN2 |
| | SLC66A2 | | | EEA1 |
| | SLC6A4 | | | ECT2 |
| | SLC6A8 | | | AC099518.1 |
| | SLC7A2 | | | FKBP15 |
| | SLC7A5 | | | ANKS4B |
| | SLC7A7 | | | RNASET2 |
| | SLC8A1 | | | AL358472.7 |
| | SLC8B1 | | | SNRNP25 |
| | SLC9A1 | | | SOCS6 |
| | SLC9A2 | | | ATR |
| | SLC9A3R2 | | | MFSD6 |
| | SLC9A4 | | | ELOVL6 |
| | SLC9A8 | | | C17orf113 |
| | SLC9B2 | | | ENTPD4 |
| | SLCO2B1 | | | UBR1 |
| | SLCO4A1 | | | OR7E91P |
| | SLF2 | | | FAT2 |
| | SLFN11 | | | PLXND1 |
| | SLFN13 | | | TGFB2 |
| | SLFN5 | | | FUT11 |
| | SLIRP | | | PHLDB1 |
| | SLIT2 | | | PXMP2 |
| | SLIT3 | | | DZIP1 |
| | SLK | | | CCPG1 |
| | SLPI | | | ANKRD26P1 |
| | SLTM | | | PLAAT3 |
| | SLU7 | | | RBM48 |
| | SLX1A | | | AC022784.1 |
| | SLX1B | | | AL590714.1 |
| | SMAD3 | | | CSTB |
| | SMAD7 | | | ZNF101 |
| | SMAP2 | | | FAM229B |
| | SMARCA1 | | | RDH10 |
| | SMARCA4 | | | AL158196.1 |
| | SMARCAL1 | | | PIEZO2 |
| | SMARCB1 | | | AC009226.1 |
| | SMARCD1 | | | GPN1 |
| | SMARCD2 | | | SGCB |

| | | | | |
|---|---|---|---|---|
| | SMARCD3 | | | TLCD4-RWDD3 |
| | SMC1B | | | NBN |
| | SMC2-AS1 | | | ONECUT2 |
| | SMC3 | | | LINC01559 |
| | SMCHD1 | | | ERCC2 |
| | SMCO4 | | | IMMT |
| | SMDT1 | | | AP003499.3 |
| | SMG1P1 | | | WDR26 |
| | SMG1P2 | | | BIN3 |
| | SMG1P3 | | | DNAH14 |
| | SMG1P4 | | | EIF4BP6 |
| | SMG5 | | | H2BC21 |
| | SMG7 | | | PSMB1 |
| | SMG7-AS1 | | | MYRIP |
| | SMG9 | | | AC004494.1 |
| | SMIM11A | | | TGFBR3 |
| | SMIM11B | | | MLPH |
| | SMIM15 | | | TDP1 |
| | SMIM25 | | | CYP3A4 |
| | SMIM26 | | | AJ003147.3 |
| | SMIM27 | | | DYNC1I2P1 |
| | SMIM3 | | | CPED1 |
| | SMIM4 | | | TBC1D23 |
| | SMNDC1 | | | L3MBTL2 |
| | SMPD1 | | | RAB43 |
| | SMPD2 | | | DPYSL4 |
| | SMPD4 | | | AREL1 |
| | SMTN | | | PLEKHS1 |
| | SMURF1 | | | CRISPLD2 |
| | SMYD3 | | | MINDY2 |
| | SMYD5 | | | GABARAPL1 |
| | SNAI1 | | | LINS1 |
| | SNAP47 | | | CLDN7 |
| | SNAPC2 | | | SDR42E1 |
| | SNCG | | | COPS7A |
| | SNED1 | | | MLLT3 |
| | SNF8 | | | SRGAP2 |
| | SNHG1 | | | NUP133 |
| | SNHG11 | | | SMIM22 |
| | SNHG12 | | | RAC1 |
| | SNHG14 | | | NDUFC1 |
| | SNHG15 | | | SLC29A1 |
| | SNHG17 | | | UNC80 |
| | SNHG19 | | | WWP1P1 |
| | SNHG25 | | | RPL39L |
| | SNHG28 | | | ATP5MD |
| | SNHG29 | | | ZNF326 |
| | SNHG3 | | | MIR425 |
| | SNHG32 | | | FAM234A |
| | SNHG5 | | | TRAF6 |
| | SNHG7 | | | SMIM31 |
| | SNHG8 | | | KTI12 |
| | SNHG9 | | | AC016596.1 |
| | SNN | | | H3-3A |
| | SNORA11F | | | GOT1 |
| | SNORA12 | | | AK7 |
| | SNORA3B | | | AHRR |
| | SNORA73B | | | CAP2 |
| | SNORA79B | | | LONRF1 |
| | SNRNP200 | | | RIDA |

| | | | | |
|---|---|---|---|---|
| | SNRNP27 | | | SNTB2 |
| | SNRNP35 | | | EAF1 |
| | SNRNP70 | | | FAM83G |
| | SNRPA1 | | | MED7 |
| | SNRPC | | | ATAD3B |
| | SNRPD2 | | | NFS1 |
| | SNRPGP15 | | | PTPA |
| | SNRPN | | | MMS22L |
| | SNTA1 | | | MICU2 |
| | SNU13 | | | KLHDC8B |
| | SNUPN | | | PRPF4 |
| | SNW1 | | | RNVU1-24 |
| | SNX10 | | | NDRG1 |
| | SNX11 | | | HOXA-AS3 |
| | SNX14 | | | PRUNE1 |
| | SNX16 | | | HSPA9 |
| | SNX17 | | | AC009090.5 |
| | SNX19 | | | HSPB1 |
| | SNX2 | | | GINS1 |
| | SNX21 | | | NGFR |
| | SNX25 | | | ADGRL3 |
| | SNX27 | | | COL20A1 |
| | SNX3 | | | AC099489.3 |
| | SNX33 | | | KEAP1 |
| | SNX5 | | | PRELID2 |
| | SNX6 | | | ISCU |
| | SNX8 | | | CARD10 |
| | SOAT1 | | | SLC7A2 |
| | SOCS1 | | | IPCEF1 |
| | SOCS2 | | | CHGA |
| | SOCS4 | | | ACTA2-AS1 |
| | SOCS5 | | | TUBA8 |
| | SOCS7 | | | UMLILO |
| | SOD2 | | | MTND5P14 |
| | SOD3 | | | AL355312.4 |
| | SOGA1 | | | HOXB8 |
| | SON | | | CFLAR-AS1 |
| | SORBS3 | | | RPL29 |
| | SORL1 | | | SWI5 |
| | SOS1 | | | SERPINA5 |
| | SOS1-IT1 | | | MFSD14C |
| | SOWAHC | | | MYL9 |
| | SOX13 | | | SMDT1 |
| | SOX21 | | | FAR2 |
| | SOX4 | | | AC022400.5 |
| | SOX9 | | | PINK1-AS |
| | SP100 | | | RAB30-DT |
| | SP110 | | | PSMB5 |
| | SP140 | | | WBP1 |
| | SP140L | | | RTKN |
| | SP2 | | | MUS81 |
| | SP6 | | | SPRY1 |
| | SPAG1 | | | RBBP8NL |
| | SPAG7 | | | PSME2P2 |
| | SPATA13 | | | RPL5P34 |
| | SPATA18 | | | BLOC1S5-TXNDC5 |
| | SPATS2L | | | CCDC90B |
| | SPC24 | | | SELENOS |
| | SPC25 | | | GLUL |
| | SPCS1 | | | GNPTG |

| | | | | |
|---|---|---|---|---|
| | SPCS2P4 | | | BECN1 |
| | SPDEF | | | BBS10 |
| | SPECC1 | | | SRGAP2C |
| | SPECC1L | | | THAP3 |
| | SPEN | | | BORA |
| | SPG21 | | | CARD8-AS1 |
| | SPG7 | | | SULT1C2 |
| | SPHK1 | | | TMEM147 |
| | SPI1 | | | ZFHX3 |
| | SPIDR | | | AP000350.4 |
| | SPIN1 | | | STAG1 |
| | SPINDOC | | | SEMA4C |
| | SPINT2 | | | ARMCX5 |
| | SPOP | | | TMEM14C |
| | SPOPL | | | IWS1 |
| | SPOUT1 | | | H2BC3 |
| | SPPL2A | | | IRF2BPL |
| | SPRED1 | | | TFE3 |
| | SPRED2 | | | ZNF586 |
| | SPRR1A | | | RPS16 |
| | SPRR2B | | | NIP7 |
| | SPRR2D | | | CSNK1G2 |
| | SPRY1 | | | SLC25A37 |
| | SPRY2 | | | PFDN6 |
| | SPRY4 | | | HEATR5A |
| | SPRYD3 | | | TMEM147-AS1 |
| | SPRYD4 | | | CRAMP1 |
| | SPRYD7 | | | HS6ST2 |
| | SPSB1 | | | RAB30 |
| | SPTBN1 | | | TNS4 |
| | SPTLC2 | | | TTLL11-IT1 |
| | SQOR | | | ZNF773 |
| | SQSTM1 | | | DNAH8 |
| | SRA1 | | | KANK1 |
| | SRBD1 | | | RPS3AP26 |
| | SRCAP | | | TNFRSF25 |
| | SRD5A3 | | | NMT1 |
| | SREBF2 | | | ANKZF1 |
| | SREK1 | | | FAM117B |
| | SRFBP1 | | | NDUFB10 |
| | SRGAP1 | | | SENP1 |
| | SRGAP2B | | | EDC4 |
| | SRGAP2C | | | PTPRO |
| | SRGN | | | SLC35A1 |
| | SRL | | | IFNA2 |
| | SRP68 | | | AL354702.1 |
| | SRR | | | AC011586.1 |
| | SRRD | | | AC093283.1 |
| | SRRM2 | | | AC010271.2 |
| | SRRM3 | | | ADCY3 |
| | SRRT | | | GPATCH11 |
| | SRSF5 | | | RSRC1 |
| | SRSF6 | | | PAXIP1 |
| | SRSF7 | | | COX6C |
| | SRSF8 | | | ZDHHC9 |
| | SRSF9 | | | BUD23 |
| | SS18L1 | | | IKBKG |
| | SSB | | | KCNJ2 |
| | SSBP2 | | | AGTRAP |
| | SSBP4 | | | RFPL4AP6 |

| | | | | |
|---|---|---|---|---|
| | SSH1 | | | DRAM2 |
| | SSNA1 | | | HPS6 |
| | SSR1 | | | SLC35A4 |
| | SSR2 | | | AL596244.1 |
| | SSR4 | | | WNK2 |
| | SSRP1 | | | SYT11 |
| | SSTR2 | | | EIF4G3 |
| | ST20 | | | BICRAL |
| | ST3GAL1 | | | CCDC198 |
| | ST3GAL3 | | | AC013717.1 |
| | ST6GAL1 | | | SNX18P7 |
| | ST6GAL2 | | | DPH6-DT |
| | ST6GALNAC2 | | | STRIP2 |
| | ST6GALNAC4 | | | ATP5IF1 |
| | ST6GALNAC6 | | | CCNK |
| | ST7L | | | SNX19 |
| | STAB1 | | | ENKD1 |
| | STAG1 | | | NARS2 |
| | STAM | | | TAF10 |
| | STAM2 | | | Z68871.1 |
| | STAP1 | | | C17orf80 |
| | STAP2 | | | RPS14 |
| | STARD10 | | | PIGO |
| | STARD13 | | | Z97832.2 |
| | STARD3 | | | LINC01138 |
| | STARD3NL | | | CRYBB2P1 |
| | STARD7 | | | AC026403.1 |
| | STARD8 | | | FAM122B |
| | STARD9 | | | LRRC75A |
| | STAT1 | | | OSER1-DT |
| | STAT2 | | | TLR2 |
| | STAT3 | | | RUNX2 |
| | STAT4 | | | COL17A1 |
| | STAT5A | | | ZNF107 |
| | STAT5B | | | SNORA38 |
| | STAT6 | | | ZC3H10 |
| | STAU1 | | | PIGX |
| | STAU2 | | | AL137856.1 |
| | STC1 | | | AC133644.2 |
| | STEAP1 | | | TAF9 |
| | STEAP4 | | | MTHFS |
| | STIM1 | | | SGMS1 |
| | STIMATE | | | ATF6B |
| | STING1 | | | KIAA1211L |
| | STIP1 | | | METTL9 |
| | STK10 | | | AC008982.1 |
| | STK11IP | | | RPS6KA2 |
| | STK16 | | | CCDC9 |
| | STK17B | | | AC097634.4 |
| | STK19 | | | EPB41L4A |
| | STK25 | | | AP4M1 |
| | STK36 | | | VKORC1L1 |
| | STK38 | | | AC006116.9 |
| | STK38L | | | SDHB |
| | STK4 | | | RCBTB2 |
| | STK40 | | | H2AC6 |
| | STMN1 | | | NUS1 |
| | STMP1 | | | PGGHG |
| | STN1 | | | RAI14 |
| | STOM | | | RNU5A-1 |

| | | | | |
|---|---|---|---|---|
| | STOML2 | | | KREMEN1 |
| | STON1 | | | |
| | STRAP | | | |
| | STRIP1 | | | |
| | STRN4 | | | |
| | STT3B | | | |
| | STUB1 | | | |
| | STX11 | | | |
| | STX12 | | | |
| | STX17 | | | |
| | STX18 | | | |
| | STX4 | | | |
| | STX5 | | | |
| | STXBP2 | | | |
| | STXBP6 | | | |
| | STYX | | | |
| | SUCNR1 | | | |
| | SUGP1 | | | |
| | SUGP2 | | | |
| | SUGT1P4-STRA6LP | | | |
| | SULF2 | | | |
| | SULT1A1 | | | |
| | SULT1A3 | | | |
| | SULT1B1 | | | |
| | SUMF1 | | | |
| | SUMF2 | | | |
| | SUMO2 | | | |
| | SUMO2P17 | | | |
| | SUN1 | | | |
| | SUN2 | | | |
| | SUOX | | | |
| | SUPT16H | | | |
| | SUPT20H | | | |
| | SUPT3H | | | |
| | SUPT4H1 | | | |
| | SUPT5H | | | |
| | SUPT6H | | | |
| | SURF1 | | | |
| | SURF4 | | | |
| | SURF6 | | | |
| | SUSD1 | | | |
| | SUSD3 | | | |
| | SUSD6 | | | |
| | SUZ12P1 | | | |
| | SVBP | | | |
| | SWAP70 | | | |
| | SYAP1 | | | |
| | SYBU | | | |
| | SYF2 | | | |
| | SYN1 | | | |
| | SYNE1 | | | |
| | SYNE3 | | | |
| | SYNGAP1 | | | |
| | SYNGR1 | | | |
| | SYNGR2 | | | |
| | SYNM | | | |
| | SYNPO | | | |
| | SYNPO2 | | | |
| | SYPL1 | | | |
| | SYT11 | | | |

| | |
|---|---|
| | SYT13 |
| | SYVN1 |
| | SZRD1 |
| | TACC1 |
| | TACC2 |
| | TAF1 |
| | TAF10 |
| | TAF11 |
| | TAF13 |
| | TAF1C |
| | TAF4 |
| | TAF5L |
| | TAF7 |
| | TAFA2 |
| | TAGAP |
| | TAGLN |
| | TAGLN2 |
| | TALDO1 |
| | TANC1 |
| | TANC2 |
| | TANGO6 |
| | TAOK1 |
| | TAOK3 |
| | TAP1 |
| | TAP2 |
| | TAPT1-AS1 |
| | TARBP2 |
| | TARDBP |
| | TARS1 |
| | TARS2 |
| | TARS3 |
| | TAS2R14 |
| | TASOR |
| | TASOR2 |
| | TASP1 |
| | TATDN2 |
| | TAX1BP1 |
| | TAZ |
| | TBC1D10B |
| | TBC1D15 |
| | TBC1D16 |
| | TBC1D17 |
| | TBC1D2 |
| | TBC1D20 |
| | TBC1D22A |
| | TBC1D25 |
| | TBC1D2B |
| | TBC1D9 |
| | TBC1D9B |
| | TBCD |
| | TBCEL |
| | TBKBP1 |
| | TBL3 |
| | TBP |
| | TBPL1 |
| | TBRG1 |
| | TBRG4 |
| | TBX2 |
| | TBX3 |
| | TCAF1 |

| | |
|---|---|
| | TCEA3 |
| | TCEAL4 |
| | TCEAL8 |
| | TCEAL9 |
| | TCEANC2 |
| | TCERG1 |
| | TCF12 |
| | TCF21 |
| | TCF25 |
| | TCF3 |
| | TCF7L1 |
| | TCF7L2 |
| | TCFL5 |
| | TCN1 |
| | TCN2 |
| | TCP1 |
| | TCTN1 |
| | TCTN3 |
| | TDP2 |
| | TDRD1 |
| | TDRD7 |
| | TDRKH-AS1 |
| | TDRP |
| | TEAD2 |
| | TEAD3 |
| | TEAD4 |
| | TECPR1 |
| | TECRP1 |
| | TECTA |
| | TEDC2 |
| | TEF |
| | TELO2 |
| | TENT4A |
| | TENT4B |
| | TENT5B |
| | TENT5C |
| | TERF2 |
| | TERF2IP |
| | TESK1 |
| | TET2 |
| | TEX10 |
| | TEX2 |
| | TEX264 |
| | TEX41 |
| | TEX9 |
| | TFAP2A |
| | TFB2M |
| | TFE3 |
| | TFEB |
| | TFEC |
| | TFIP11 |
| | TFPI2 |
| | TG |
| | TGFA |
| | TGFB1 |
| | TGFB1I1 |
| | TGFB2 |
| | TGFBR1 |
| | TGFBR2 |
| | TGFBRAP1 |

| | |
|---|---|
| | TGIF2 |
| | TGM2 |
| | THAP1 |
| | THAP12 |
| | THAP2 |
| | THAP3 |
| | THAP4 |
| | THAP5 |
| | THAP7 |
| | THAP9-AS1 |
| | THBD |
| | THBS3 |
| | THEM4 |
| | THEMIS2 |
| | THG1L |
| | THOC1 |
| | THOC3 |
| | THOC6 |
| | THOC7 |
| | THOP1 |
| | THRA |
| | THRB |
| | THSD4 |
| | THUMPD1 |
| | THUMPD3 |
| | TIA1 |
| | TIAM1 |
| | TICAM1 |
| | TIFA |
| | TIGAR |
| | TIMELESS |
| | TIMM10 |
| | TIMM10B |
| | TIMM13 |
| | TIMM17B |
| | TIMM23 |
| | TIMM44 |
| | TIMM9 |
| | TIMP1 |
| | TIMP3 |
| | TINAGL1 |
| | TINCR |
| | TIPRL |
| | TJAP1 |
| | TJP1 |
| | TJP2 |
| | TJP3 |
| | TKFC |
| | TKT |
| | TLCD3A |
| | TLCD4 |
| | TLCD5 |
| | TLDC2 |
| | TLE1 |
| | TLE2 |
| | TLE3 |
| | TLE4 |
| | TLL1 |
| | TLN1 |
| | TLR1 |

| | |
|---|---|
| | TLR2 |
| | TLR4 |
| | TLR6 |
| | TLR7 |
| | TM4SF1 |
| | TM4SF19 |
| | TM4SF20 |
| | TM7SF3 |
| | TM9SF3 |
| | TMA7 |
| | TMBIM1 |
| | TMC6 |
| | TMCO1 |
| | TMCO3 |
| | TMCO4 |
| | TMED10 |
| | TMED4 |
| | TMED5 |
| | TMED8 |
| | TMED9 |
| | TMEM100 |
| | TMEM109 |
| | TMEM11 |
| | TMEM115 |
| | TMEM123 |
| | TMEM125 |
| | TMEM126B |
| | TMEM127 |
| | TMEM131 |
| | TMEM133 |
| | TMEM134 |
| | TMEM141 |
| | TMEM14B |
| | TMEM150A |
| | TMEM154 |
| | TMEM159 |
| | TMEM160 |
| | TMEM161A |
| | TMEM171 |
| | TMEM175 |
| | TMEM176B |
| | TMEM18 |
| | TMEM183A |
| | TMEM184B |
| | TMEM185A |
| | TMEM189 |
| | TMEM192 |
| | TMEM198B |
| | TMEM201 |
| | TMEM203 |
| | TMEM205 |
| | TMEM208 |
| | TMEM209 |
| | TMEM212 |
| | TMEM214 |
| | TMEM218 |
| | TMEM219 |
| | TMEM222 |
| | TMEM230 |
| | TMEM243 |

| | |
|---|---|
| | TMEM245 |
| | TMEM254 |
| | TMEM254-AS1 |
| | TMEM255B |
| | TMEM258 |
| | TMEM259 |
| | TMEM260 |
| | TMEM39A |
| | TMEM39B |
| | TMEM41B |
| | TMEM43 |
| | TMEM45A |
| | TMEM45B |
| | TMEM50A |
| | TMEM50B |
| | TMEM51 |
| | TMEM52B |
| | TMEM54 |
| | TMEM59 |
| | TMEM60 |
| | TMEM69 |
| | TMEM71 |
| | TMEM74B |
| | TMEM87B |
| | TMEM88 |
| | TMEM92 |
| | TMEM94 |
| | TMEM97 |
| | TMEM9B |
| | TMLHE-AS1 |
| | TMOD1 |
| | TMOD2 |
| | TMOD3 |
| | TMPRSS2 |
| | TMSB10 |
| | TMSB4XP4 |
| | TMTC1 |
| | TMUB2 |
| | TMX1 |
| | TMX2 |
| | TNF |
| | TNFAIP1 |
| | TNFAIP6 |
| | TNFAIP8 |
| | TNFAIP8L3 |
| | TNFRSF10B |
| | TNFRSF10D |
| | TNFRSF14 |
| | TNFRSF1B |
| | TNFRSF21 |
| | TNFSF10 |
| | TNFSF13B |
| | TNFSF14 |
| | TNFSF8 |
| | TNIK |
| | TNIP1 |
| | TNIP2 |
| | TNIP3 |
| | TNNC1 |
| | TNNT2 |

| | |
|---|---|
| | TNPO2 |
| | TNRC18P3 |
| | TNRC6C |
| | TNS1 |
| | TNS2 |
| | TNS4 |
| | TNXA |
| | TNXB |
| | TOB2 |
| | TOE1 |
| | TOLLIP |
| | TOM1 |
| | TOM1L2 |
| | TOMM34 |
| | TOMM40 |
| | TOMM40L |
| | TOP1MT |
| | TOP2A |
| | TOP2B |
| | TOP3A |
| | TOR1A |
| | TOR1AIP1 |
| | TOR1AIP2 |
| | TOR1B |
| | TOR4A |
| | TOX4 |
| | TP53 |
| | TP53BP1 |
| | TP53BP2 |
| | TP53I3 |
| | TP53INP2 |
| | TP53RK |
| | TP53TG3E |
| | TP73 |
| | TP73-AS1 |
| | TPBG |
| | TPCN1 |
| | TPD52L2 |
| | TPH2 |
| | TPM1 |
| | TPM2 |
| | TPM4 |
| | TPMT |
| | TPP1 |
| | TPPP |
| | TPR |
| | TPRA1 |
| | TPRG1L |
| | TPRKB |
| | TPSAB1 |
| | TPSB2 |
| | TPST1 |
| | TPT1 |
| | TPT1-AS1 |
| | TPTEP1 |
| | TRA2B |
| | TRABD |
| | TRAF1 |
| | TRAF2 |
| | TRAF3IP2 |

| | |
|---|---|
| | TRAF3IP2-AS1 |
| | TRAF4 |
| | TRAF5 |
| | TRAF6 |
| | TRAK1 |
| | TRAK2 |
| | TRAM2 |
| | TRAP1 |
| | TRAPPC1 |
| | TRAPPC10 |
| | TRAPPC12 |
| | TRAPPC2B |
| | TRAPPC3 |
| | TRAPPC4 |
| | TRAPPC9 |
| | TREM1 |
| | TREML3P |
| | TRGC2 |
| | TRIB1 |
| | TRIB3 |
| | TRIM11 |
| | TRIM14 |
| | TRIM16 |
| | TRIM16L |
| | TRIM21 |
| | TRIM22 |
| | TRIM25 |
| | TRIM26 |
| | TRIM27 |
| | TRIM29 |
| | TRIM3 |
| | TRIM38 |
| | TRIM5 |
| | TRIM52-AS1 |
| | TRIM59 |
| | TRIM68 |
| | TRIM69 |
| | TRIO |
| | TRIP10 |
| | TRIP6 |
| | TRIQK |
| | TRIR |
| | TRIT1 |
| | TRMT1 |
| | TRMT10A |
| | TRMT11 |
| | TRMT12 |
| | TRMT1L |
| | TRMT5 |
| | TRMT61A |
| | TRMU |
| | TRNAU1AP |
| | TROAP |
| | TRPA1 |
| | TRPC4AP |
| | TRPC6 |
| | TRPM2-AS |
| | TRPM4 |
| | TRPM7 |
| | TRPM8 |

| | |
|---|---|
| | TRPS1 |
| | TRRAP |
| | TRUB2 |
| | TSBP1-AS1 |
| | TSC1 |
| | TSC2 |
| | TSC22D1 |
| | TSC22D3 |
| | TSC22D4 |
| | TSEN34 |
| | TSEN54 |
| | TSFM |
| | TSHZ1 |
| | TSHZ3 |
| | TSIX |
| | TSN |
| | TSPAN14 |
| | TSPAN17 |
| | TSPAN18 |
| | TSPAN31 |
| | TSPAN4 |
| | TSPAN7 |
| | TSPAN9 |
| | TSPO |
| | TSPY26P |
| | TSPYL1 |
| | TSPYL2 |
| | TSPYL4 |
| | TSR1 |
| | TSR3 |
| | TSSC4 |
| | TSSK6 |
| | TSTA3 |
| | TTBK2 |
| | TTC14 |
| | TTC19 |
| | TTC3 |
| | TTC31 |
| | TTC38 |
| | TTC39B |
| | TTC7A |
| | TTC7B |
| | TTC9C |
| | TTF1 |
| | TTL |
| | TTLL4 |
| | TTPA |
| | TTTY15 |
| | TUBA1A |
| | TUBA1B |
| | TUBA1C |
| | TUBA3D |
| | TUBA4A |
| | TUBAP2 |
| | TUBB |
| | TUBB2A |
| | TUBB2B |
| | TUBB4B |
| | TUBB6 |
| | TUBG1 |

| | |
|---|---|
| | TUBGCP2 |
| | TUBGCP3 |
| | TUBGCP4 |
| | TUBGCP6 |
| | TUFM |
| | TUG1 |
| | TULP3 |
| | TULP4 |
| | TUSC2 |
| | TUSC3 |
| | TUT4 |
| | TVP23B |
| | TWF1P1 |
| | TWF2 |
| | TWIST2 |
| | TWNK |
| | TXLNA |
| | TXLNGY |
| | TXNDC11 |
| | TXNDC12 |
| | TXNDC15 |
| | TXNL4A |
| | TXNL4B |
| | TYMS |
| | TYRO3 |
| | TYW1B |
| | U2AF1 |
| | U2AF1L5 |
| | U2AF2 |
| | U2SURP |
| | U62317.1 |
| | UACA |
| | UAP1 |
| | UAP1L1 |
| | UBA1 |
| | UBA2 |
| | UBA52 |
| | UBA7 |
| | UBAC2 |
| | UBAC2-AS1 |
| | UBALD1 |
| | UBAP2 |
| | UBASH3B |
| | UBB |
| | UBC |
| | UBE2A |
| | UBE2B |
| | UBE2D1 |
| | UBE2D2 |
| | UBE2D3 |
| | UBE2E1 |
| | UBE2E2 |
| | UBE2E3 |
| | UBE2F |
| | UBE2G2 |
| | UBE2H |
| | UBE2I |
| | UBE2J1 |
| | UBE2J2 |
| | UBE2L6 |

| | |
|---|---|
| | UBE2M |
| | UBE2N |
| | UBE2O |
| | UBE2Q1 |
| | UBE2Z |
| | UBE3A |
| | UBE3C |
| | UBIAD1 |
| | UBL3 |
| | UBL4A |
| | UBL5 |
| | UBL7 |
| | UBN1 |
| | UBQLN1 |
| | UBQLN4 |
| | UBR4 |
| | UBR5 |
| | UBTF |
| | UBXN1 |
| | UBXN11 |
| | UBXN2B |
| | UBXN6 |
| | UBXN7 |
| | UCHL1 |
| | UCHL5 |
| | UCK1 |
| | UCK2 |
| | UCP2 |
| | UEVLD |
| | UFD1 |
| | UFSP2 |
| | UGDH-AS1 |
| | UGGT2 |
| | UGT1A7 |
| | UHRF1BP1 |
| | UHRF2 |
| | UIMC1 |
| | ULBP2 |
| | ULK1 |
| | UMPS |
| | UNC13A |
| | UNC13B |
| | UNC13D |
| | UNC45A |
| | UNC5D |
| | UNC93B1 |
| | UNK |
| | UPF1 |
| | UPF3A |
| | UPF3B |
| | UPK2 |
| | UPK3B |
| | UPP1 |
| | UQCC1 |
| | UQCC2 |
| | UQCRC1 |
| | UQCRC2 |
| | UQCRFS1 |
| | URB1 |
| | URB2 |

| | |
|---|---|
| | URGCP |
| | URM1 |
| | UROD |
| | USB1 |
| | USE1 |
| | USF1 |
| | USF3 |
| | USH1C |
| | USHBP1 |
| | USP10 |
| | USP15 |
| | USP18 |
| | USP19 |
| | USP20 |
| | USP21 |
| | USP22 |
| | USP25 |
| | USP3 |
| | USP30 |
| | USP39 |
| | USP4 |
| | USP40 |
| | USP47 |
| | USP48 |
| | USP49 |
| | USP5 |
| | USP53 |
| | USP7 |
| | UTP14C |
| | UTP20 |
| | UTP23 |
| | UTP3 |
| | UTP4 |
| | UTP6 |
| | VAC14 |
| | VAMP1 |
| | VAMP3 |
| | VAMP5 |
| | VANGL1 |
| | VANGL2 |
| | VAPB |
| | VASH1 |
| | VASN |
| | VASP |
| | VAT1 |
| | VAV1 |
| | VAV2 |
| | VAV3 |
| | VAX1 |
| | VCAN |
| | VCP |
| | VCPIP1 |
| | VCPKMT |
| | VDAC1 |
| | VDAC3 |
| | VEGFA |
| | VEGFB |
| | VEGFD |
| | VEZT |
| | VGLL3 |

| | |
|---|---|
| | VGLL4 |
| | VIM |
| | VIPAS39 |
| | VIPR1 |
| | VIRMA |
| | VKORC1 |
| | VLDLR |
| | VNN2 |
| | VNN3 |
| | VOPP1 |
| | VPS11 |
| | VPS13D |
| | VPS16 |
| | VPS18 |
| | VPS25 |
| | VPS26A |
| | VPS26C |
| | VPS28 |
| | VPS29 |
| | VPS33B |
| | VPS35 |
| | VPS37B |
| | VPS39 |
| | VPS45 |
| | VPS51 |
| | VPS53 |
| | VPS54 |
| | VPS72 |
| | VPS8 |
| | VSIG1 |
| | VSIG10 |
| | VSIG10L |
| | VSIG2 |
| | VSX1 |
| | VTI1A |
| | VTRNA1-1 |
| | VTRNA1-2 |
| | VWA7 |
| | VWF |
| | WAPL |
| | WARS2 |
| | WAS |
| | WASF2 |
| | WASF3 |
| | WASH2P |
| | WASH3P |
| | WASH4P |
| | WASH5P |
| | WASH6P |
| | WASH7P |
| | WASH8P |
| | WASH9P |
| | WASHC1 |
| | WASHC2A |
| | WASHC4 |
| | WASL |
| | WBP1L |
| | WDFY2 |
| | WDFY3 |
| | WDR11 |

| | |
|---|---|
| | WDR13 |
| | WDR18 |
| | WDR20 |
| | WDR27 |
| | WDR34 |
| | WDR35 |
| | WDR36 |
| | WDR37 |
| | WDR4 |
| | WDR43 |
| | WDR44 |
| | WDR45B |
| | WDR46 |
| | WDR5 |
| | WDR59 |
| | WDR6 |
| | WDR60 |
| | WDR61 |
| | WDR74 |
| | WDR75 |
| | WDR78 |
| | WDR82 |
| | WDR83OS |
| | WDR88 |
| | WDR89 |
| | WDR91 |
| | WDTC1 |
| | WDYHV1 |
| | WEE1 |
| | WFDC2 |
| | WFDC21P |
| | WFS1 |
| | WHAMM |
| | WIPF1 |
| | WIPI1 |
| | WIPI2 |
| | WNK4 |
| | WNT2B |
| | WNT7B |
| | WNT9A |
| | WRAP73 |
| | WRNIP1 |
| | WTAP |
| | WTIP |
| | WWC2 |
| | WWTR1 |
| | XAB2 |
| | XAF1 |
| | XAGE1A |
| | XAGE1B |
| | XKR9 |
| | XPA |
| | XPC |
| | XPNPEP1 |
| | XPO5 |
| | XRCC6 |
| | XRN1 |
| | XRN2 |
| | XYLB |
| | XYLT1 |

| | |
|---|---|
| | XYLT2 |
| | YAE1 |
| | YAP1 |
| | YARS1 |
| | YBX1 |
| | YBX1P1 |
| | YEATS2 |
| | YIPF1 |
| | YIPF5 |
| | YIPF6 |
| | YJU2 |
| | YKT6 |
| | YPEL2 |
| | YPEL3 |
| | YRDC |
| | YTHDF1 |
| | YWHAE |
| | YWHAZ |
| | YY2 |
| | ZBED2 |
| | ZBED4 |
| | ZBED5 |
| | ZBED8 |
| | ZBP1 |
| | ZBTB1 |
| | ZBTB10 |
| | ZBTB14 |
| | ZBTB17 |
| | ZBTB18 |
| | ZBTB2 |
| | ZBTB24 |
| | ZBTB33 |
| | ZBTB34 |
| | ZBTB38 |
| | ZBTB39 |
| | ZBTB4 |
| | ZBTB40 |
| | ZBTB44 |
| | ZBTB46 |
| | ZBTB5 |
| | ZBTB7A |
| | ZBTB8OS |
| | ZBTB9 |
| | ZC3H11A |
| | ZC3H12A |
| | ZC3H18 |
| | ZC3H4 |
| | ZC3H6 |
| | ZC3H7A |
| | ZC3H7B |
| | ZC3HAV1 |
| | ZC3HC1 |
| | ZCCHC14 |
| | ZCCHC24 |
| | ZCCHC3 |
| | ZCCHC7 |
| | ZCCHC8 |
| | ZCCHC9 |
| | ZDHHC11 |
| | ZDHHC11B |

| | |
|---|---|
| | ZDHHC12 |
| | ZDHHC13 |
| | ZDHHC14 |
| | ZDHHC16 |
| | ZDHHC19 |
| | ZDHHC2 |
| | ZDHHC6 |
| | ZDHHC7 |
| | ZDHHC8P1 |
| | ZEB1 |
| | ZER1 |
| | ZFAND2A |
| | ZFAND2B |
| | ZFAND5 |
| | ZFAND6 |
| | ZFAT |
| | ZFP28 |
| | ZFP3 |
| | ZFP36L1 |
| | ZFP36L2 |
| | ZFP64 |
| | ZFP90 |
| | ZFYVE1 |
| | ZFYVE19 |
| | ZFYVE27 |
| | ZHX3 |
| | ZKSCAN1 |
| | ZKSCAN5 |
| | ZMAT2 |
| | ZMIZ1 |
| | ZMIZ2 |
| | ZMPSTE24 |
| | ZMYM2 |
| | ZMYM4 |
| | ZMYND12 |
| | ZMYND19 |
| | ZNF10 |
| | ZNF100 |
| | ZNF106 |
| | ZNF12 |
| | ZNF121 |
| | ZNF131 |
| | ZNF133 |
| | ZNF134 |
| | ZNF136 |
| | ZNF140 |
| | ZNF142 |
| | ZNF154 |
| | ZNF160 |
| | ZNF175 |
| | ZNF184 |
| | ZNF185 |
| | ZNF197 |
| | ZNF211 |
| | ZNF212 |
| | ZNF22 |
| | ZNF221 |
| | ZNF222 |
| | ZNF227 |
| | ZNF24 |

| | |
|---|---|
| | ZNF248 |
| | ZNF25 |
| | ZNF26 |
| | ZNF263 |
| | ZNF266 |
| | ZNF273 |
| | ZNF276 |
| | ZNF277 |
| | ZNF282 |
| | ZNF286A |
| | ZNF3 |
| | ZNF317 |
| | ZNF330 |
| | ZNF331 |
| | ZNF333 |
| | ZNF335 |
| | ZNF343 |
| | ZNF35 |
| | ZNF354A |
| | ZNF384 |
| | ZNF385A |
| | ZNF385D |
| | ZNF394 |
| | ZNF395 |
| | ZNF407 |
| | ZNF408 |
| | ZNF431 |
| | ZNF433 |
| | ZNF440 |
| | ZNF449 |
| | ZNF45 |
| | ZNF451 |
| | ZNF461 |
| | ZNF462 |
| | ZNF468 |
| | ZNF473 |
| | ZNF480 |
| | ZNF488 |
| | ZNF496 |
| | ZNF497 |
| | ZNF502 |
| | ZNF513 |
| | ZNF514 |
| | ZNF518A |
| | ZNF558 |
| | ZNF576 |
| | ZNF577 |
| | ZNF583 |
| | ZNF585B |
| | ZNF589 |
| | ZNF592 |
| | ZNF595 |
| | ZNF597 |
| | ZNF598 |
| | ZNF599 |
| | ZNF600 |
| | ZNF610 |
| | ZNF614 |
| | ZNF621 |
| | ZNF622 |

| | |
|---|---|
| | ZNF629 |
| | ZNF638 |
| | ZNF655 |
| | ZNF664 |
| | ZNF672 |
| | ZNF687 |
| | ZNF688 |
| | ZNF689 |
| | ZNF692 |
| | ZNF697 |
| | ZNF700 |
| | ZNF710 |
| | ZNF720 |
| | ZNF732 |
| | ZNF740 |
| | ZNF76 |
| | ZNF761 |
| | ZNF766 |
| | ZNF768 |
| | ZNF776 |
| | ZNF778 |
| | ZNF783 |
| | ZNF785 |
| | ZNF800 |
| | ZNF81 |
| | ZNF816 |
| | ZNF823 |
| | ZNF829 |
| | ZNF83 |
| | ZNF836 |
| | ZNF84 |
| | ZNF841 |
| | ZNF862 |
| | ZNF876P |
| | ZNHIT1 |
| | ZNHIT3 |
| | ZNNT1 |
| | ZNRD2 |
| | ZNRF2 |
| | ZP3 |
| | ZPLD1 |
| | ZPR1 |
| | ZRANB1 |
| | ZRANB3 |
| | ZRSR2 |
| | ZSCAN16 |
| | ZSCAN18 |
| | ZSCAN22 |
| | ZSCAN29 |
| | ZSCAN31 |
| | ZSCAN9 |
| | ZSWIM7 |
| | ZSWIM8 |
| | ZUP1 |
| | ZW10 |
| | ZWINT |
| | ZYG11A |
| | ZZEF1 |
