## Supplementary file 6 for "Host transcriptional responses and SARS-CoV-2 isolates from the nasopharyngeal samples of Bangladeshi COVID-19 patients"

**Supplementary file 6: Genes and associated terms used for filtering the expression values used in Figure 6.**

| Cytokine related genes | Associated terms |
|---|---|
| CD55 | CD4-positive, alpha-beta T cell cytokine production |
| E2F8 | cell cycle comprising mitosis without cytokinesis |
| AKAP6 | cellular response to cytokine stimulus |
| ASAH2 | cellular response to cytokine stimulus |
| CASP1 | cellular response to cytokine stimulus |
| CCR7 | cellular response to cytokine stimulus |
| CRNN | cellular response to cytokine stimulus |
| CSF1R | cellular response to cytokine stimulus |
| CSF3 | cellular response to cytokine stimulus |
| CXCR4 | cellular response to cytokine stimulus |
| FLT3 | cellular response to cytokine stimulus |
| FOXF1 | cellular response to cytokine stimulus |
| HAX1 | cellular response to cytokine stimulus |
| HCLS1 | cellular response to cytokine stimulus |
| IL13 | cellular response to cytokine stimulus |
| IL18BP | cellular response to cytokine stimulus |
| IL18R1 | cellular response to cytokine stimulus |
| IL18RAP | cellular response to cytokine stimulus |
| IL1RL2 | cellular response to cytokine stimulus |
| IL37 | cellular response to cytokine stimulus |
| ITGA4 | cellular response to cytokine stimulus |
| LEF1 | cellular response to cytokine stimulus |
| PID1 | cellular response to cytokine stimulus |
| PTPN14 | cellular response to cytokine stimulus |
| PTPN2 | cellular response to cytokine stimulus |
| PTPN7 | cellular response to cytokine stimulus |
| STAT1 | cellular response to cytokine stimulus |
| TRPV1 | cellular response to cytokine stimulus |
| CD28 | cytokine biosynthetic process |
| CEBPE | cytokine biosynthetic process |
| BATF | cytokine-mediated signaling pathway |
| BCL6 | cytokine-mediated signaling pathway |
| BIRC5 | cytokine-mediated signaling pathway |
| CASP1 | cytokine-mediated signaling pathway |
| CCL11 | cytokine-mediated signaling pathway |
| CCL20 | cytokine-mediated signaling pathway |
| CCL22 | cytokine-mediated signaling pathway |
| CCL2 | cytokine-mediated signaling pathway |
| CCL4 | cytokine-mediated signaling pathway |
| CCL5 | cytokine-mediated signaling pathway |
| CCR1 | cytokine-mediated signaling pathway |
| CCR2 | cytokine-mediated signaling pathway |
| CCR5 | cytokine-mediated signaling pathway |
| CD36 | cytokine-mediated signaling pathway |
| CD4 | cytokine-mediated signaling pathway |

| | |
|---|---|
| CD80 | cytokine-mediated signaling pathway |
| CD86 | cytokine-mediated signaling pathway |
| CNOT9 | cytokine-mediated signaling pathway |
| CNTF | cytokine-mediated signaling pathway |
| CRK | cytokine-mediated signaling pathway |
| CSF1 | cytokine-mediated signaling pathway |
| CSF1R | cytokine-mediated signaling pathway |
| CSF2RB | cytokine-mediated signaling pathway |
| CSF3 | cytokine-mediated signaling pathway |
| CX3CL1 | cytokine-mediated signaling pathway |
| CXCL10 | cytokine-mediated signaling pathway |
| CXCL1 | cytokine-mediated signaling pathway |
| CXCL2 | cytokine-mediated signaling pathway |
| CXCL8 | cytokine-mediated signaling pathway |
| DUOX1 | cytokine-mediated signaling pathway |
| DUOX2 | cytokine-mediated signaling pathway |
| EPOR | cytokine-mediated signaling pathway |
| F3 | cytokine-mediated signaling pathway |
| FASLG | cytokine-mediated signaling pathway |
| FER | cytokine-mediated signaling pathway |
| FGF2 | cytokine-mediated signaling pathway |
| FLT3 | cytokine-mediated signaling pathway |
| FLT3LG | cytokine-mediated signaling pathway |
| FN1 | cytokine-mediated signaling pathway |
| FOS | cytokine-mediated signaling pathway |
| FYN | cytokine-mediated signaling pathway |
| FZD4 | cytokine-mediated signaling pathway |
| GREM2 | cytokine-mediated signaling pathway |
| HGF | cytokine-mediated signaling pathway |
| ICAM1 | cytokine-mediated signaling pathway |
| IFNL1 | cytokine-mediated signaling pathway |
| IFNL2 | cytokine-mediated signaling pathway |
| IFNL3 | cytokine-mediated signaling pathway |
| IFNLR1 | cytokine-mediated signaling pathway |
| IGHE | cytokine-mediated signaling pathway |
| IGHG1 | cytokine-mediated signaling pathway |
| IL11 | cytokine-mediated signaling pathway |
| IL11RA | cytokine-mediated signaling pathway |
| IL12A | cytokine-mediated signaling pathway |
| IL13 | cytokine-mediated signaling pathway |
| IL16 | cytokine-mediated signaling pathway |
| IL17RB | cytokine-mediated signaling pathway |
| IL1A | cytokine-mediated signaling pathway |
| IL1B | cytokine-mediated signaling pathway |
| IL1R2 | cytokine-mediated signaling pathway |
| IL1RAP | cytokine-mediated signaling pathway |
| IL1RAPL2 | cytokine-mediated signaling pathway |
| IL1RL1 | cytokine-mediated signaling pathway |
| IL1RL2 | cytokine-mediated signaling pathway |
| IL1RN | cytokine-mediated signaling pathway |

| | |
|---|---|
| IL20 | cytokine-mediated signaling pathway |
| IL20RB | cytokine-mediated signaling pathway |
| IL22RA1 | cytokine-mediated signaling pathway |
| IL23A | cytokine-mediated signaling pathway |
| IL23R | cytokine-mediated signaling pathway |
| IL24 | cytokine-mediated signaling pathway |
| IL2RA | cytokine-mediated signaling pathway |
| IL2RG | cytokine-mediated signaling pathway |
| IL31RA | cytokine-mediated signaling pathway |
| IL32 | cytokine-mediated signaling pathway |
| IL34 | cytokine-mediated signaling pathway |
| IL36RN | cytokine-mediated signaling pathway |
| IL37 | cytokine-mediated signaling pathway |
| IL3RA | cytokine-mediated signaling pathway |
| IL4 | cytokine-mediated signaling pathway |
| IL4R | cytokine-mediated signaling pathway |
| IL5RA | cytokine-mediated signaling pathway |
| IL6 | cytokine-mediated signaling pathway |
| IL6R | cytokine-mediated signaling pathway |
| IL6ST | cytokine-mediated signaling pathway |
| INPP5D | cytokine-mediated signaling pathway |
| IRF5 | cytokine-mediated signaling pathway |
| ITGAM | cytokine-mediated signaling pathway |
| ITGAX | cytokine-mediated signaling pathway |
| ITGB2 | cytokine-mediated signaling pathway |
| JAK2 | cytokine-mediated signaling pathway |
| JUNB | cytokine-mediated signaling pathway |
| KRAS | cytokine-mediated signaling pathway |
| LBP | cytokine-mediated signaling pathway |
| LRP8 | cytokine-mediated signaling pathway |
| MCL1 | cytokine-mediated signaling pathway |
| MPL | cytokine-mediated signaling pathway |
| MYC | cytokine-mediated signaling pathway |
| MYD88 | cytokine-mediated signaling pathway |
| OPRD1 | cytokine-mediated signaling pathway |
| OPRM1 | cytokine-mediated signaling pathway |
| PIK3CB | cytokine-mediated signaling pathway |
| POMC | cytokine-mediated signaling pathway |
| PTGS2 | cytokine-mediated signaling pathway |
| PTPRN | cytokine-mediated signaling pathway |
| SAA1 | cytokine-mediated signaling pathway |
| SOCS1 | cytokine-mediated signaling pathway |
| SOCS3 | cytokine-mediated signaling pathway |
| STAT1 | cytokine-mediated signaling pathway |
| STAT2 | cytokine-mediated signaling pathway |
| STAT4 | cytokine-mediated signaling pathway |
| STAT5B | cytokine-mediated signaling pathway |
| STX4 | cytokine-mediated signaling pathway |
| TNF | cytokine-mediated signaling pathway |
| TNFRSF1A | cytokine-mediated signaling pathway |

| | |
|---|---|
| TP53 | cytokine-mediated signaling pathway |
| VCAM1 | cytokine-mediated signaling pathway |
| YWHAZ | cytokine-mediated signaling pathway |
| TNFSF15 | cytokine metabolic process |
| AZI2 | cytokine production |
| BATF | cytokine production |
| CD226 | cytokine production |
| CD4 | cytokine production |
| DBH | cytokine production |
| FABP4 | cytokine production |
| FOXP3 | cytokine production |
| MAF | cytokine production |
| NFATC1 | cytokine production |
| NFATC2 | cytokine production |
| NFATC3 | cytokine production |
| PIK3CG | cytokine production |
| S1PR3 | cytokine production |
| CD96 | cytokine production involved in inflammatory response |
| IDO1 | cytokine production involved in inflammatory response |
| BTN3A1 | cytokine secretion |
| NLRP3 | cytokine secretion involved in immune response |
| CECR2 | cytoskeleton-dependent cytokinesis |
| SEPTIN12 | cytoskeleton-dependent cytokinesis |
| SEPTIN1 | cytoskeleton-dependent cytokinesis |
| SEPTIN4 | cytoskeleton-dependent cytokinesis |
| SEPTIN5 | cytoskeleton-dependent cytokinesis |
| SEPTIN6 | cytoskeleton-dependent cytokinesis |
| SEPTIN9 | cytoskeleton-dependent cytokinesis |
| ANK3 | mitotic cytokinesis |
| ANLN | mitotic cytokinesis |
| APC | mitotic cytokinesis |
| CENPA | mitotic cytokinesis |
| CEP55 | mitotic cytokinesis |
| CKAP2 | mitotic cytokinesis |
| ECT2 | mitotic cytokinesis |
| EFHC1 | mitotic cytokinesis |
| ESPL1 | mitotic cytokinesis |
| KIF20A | mitotic cytokinesis |
| KIF23 | mitotic cytokinesis |
| KIF4A | mitotic cytokinesis |
| PLK1 | mitotic cytokinesis |
| RACGAP1 | mitotic cytokinesis |
| RHOB | mitotic cytokinesis |
| ROCK1 | mitotic cytokinesis |
| ROCK2 | mitotic cytokinesis |
| SEPTIN6 | mitotic cytokinesis |
| SPTBN1 | mitotic cytokinesis |
| TRIM36 | mitotic cytokinesis |
| UNC119 | mitotic cytokinesis |
| AURKB | mitotic cytokinesis checkpoint |

| | |
|---|---|
| CNTROB | mitotic cytokinetic process |
| ASB1 | negative regulation of cytokine biosynthetic process |
| FOXP3 | negative regulation of cytokine biosynthetic process |
| TIA1 | negative regulation of cytokine biosynthetic process |
| IL36RN | negative regulation of cytokine-mediated signaling pathway |
| PTPRC | negative regulation of cytokine-mediated signaling pathway |
| PXDN | negative regulation of cytokine-mediated signaling pathway |
| AXL | negative regulation of cytokine production |
| BTK | negative regulation of cytokine production |
| CLEC4A | negative regulation of cytokine production |
| MIR155 | negative regulation of cytokine production |
| NFKB1 | negative regulation of cytokine production |
| TWSG1 | negative regulation of cytokine production |
| ABCD2 | negative regulation of cytokine production involved in inflammatory response |
| ADCY7 | negative regulation of cytokine production involved in inflammatory response |
| APOD | negative regulation of cytokine production involved in inflammatory response |
| F2 | negative regulation of cytokine production involved in inflammatory response |
| IL1R2 | negative regulation of cytokine production involved in inflammatory response |
| MEFV | negative regulation of cytokine production involved in inflammatory response |
| MIR155 | negative regulation of cytokine production involved in inflammatory response |
| ZC3H12A | negative regulation of cytokine production involved in inflammatory response |
| BTN2A2 | negative regulation of cytokine secretion |
| FCGR2B | negative regulation of cytokine secretion |
| FFAR4 | negative regulation of cytokine secretion |
| FOXP3 | negative regulation of cytokine secretion |
| PTGER4 | negative regulation of cytokine secretion |
| SRGN | negative regulation of cytokine secretion |
| ANGPT1 | negative regulation of cytokine secretion involved in immune response |
| APOA1 | negative regulation of cytokine secretion involved in immune response |
| APOA2 | negative regulation of cytokine secretion involved in immune response |
| LILRB1 | negative regulation of cytokine secretion involved in immune response |
| TNF | negative regulation of cytokine secretion involved in immune response |
| AURKB | negative regulation of cytokinesis |
| E2F7 | negative regulation of cytokinesis |
| E2F8 | negative regulation of cytokinesis |
| TGFB2 | negative regulation of macrophage cytokine production |
| TGFB3 | negative regulation of macrophage cytokine production |
| BCL6 | negative regulation of mast cell cytokine production |
| CD96 | negative regulation of natural killer cell cytokine production |
| HLA-F | negative regulation of natural killer cell cytokine production |
| BST2 | negative regulation of plasmacytoid dendritic cell cytokine production |
| APOA1 | negative regulation of response to cytokine stimulus |
| KLF4 | negative regulation of response to cytokine stimulus |
| MAPK7 | negative regulation of response to cytokine stimulus |
| FOXP3 | negative regulation of T cell cytokine production |
| HLA-F | negative regulation of T cell cytokine production |
| SMAD7 | negative regulation of T cell cytokine production |
| ARG1 | negative regulation of T-helper 2 cell cytokine production |
| NRP1 | positive regulation of cytokine activity |
| AXL | positive regulation of cytokine-mediated signaling pathway |

| | |
|---|---|
| CD74 | positive regulation of cytokine-mediated signaling pathway |
| RIPK2 | positive regulation of cytokine-mediated signaling pathway |
| IFI16 | positive regulation of cytokine production |
| IL21 | positive regulation of cytokine production |
| IL33 | positive regulation of cytokine production |
| NFAM1 | positive regulation of cytokine production |
| TLR3 | positive regulation of cytokine production |
| TNF | positive regulation of cytokine production |
| WNT5A | positive regulation of cytokine production |
| CD6 | positive regulation of cytokine production involved in inflammatory response |
| CLEC7A | positive regulation of cytokine production involved in inflammatory response |
| GBP5 | positive regulation of cytokine production involved in inflammatory response |
| KARS1 | positive regulation of cytokine production involved in inflammatory response |
| MIR21 | positive regulation of cytokine production involved in inflammatory response |
| MYD88 | positive regulation of cytokine production involved in inflammatory response |
| TICAM1 | positive regulation of cytokine production involved in inflammatory response |
| TLR4 | positive regulation of cytokine production involved in inflammatory response |
| TLR6 | positive regulation of cytokine production involved in inflammatory response |
| C1QTNF3 | positive regulation of cytokine secretion |
| CADM1 | positive regulation of cytokine secretion |
| FGR | positive regulation of cytokine secretion |
| IL1A | positive regulation of cytokine secretion |
| PTGER4 | positive regulation of cytokine secretion |
| SAA1 | positive regulation of cytokine secretion |
| TNF | positive regulation of cytokine secretion |
| KARS1 | positive regulation of cytokine secretion involved in immune response |
| TNFRSF14 | positive regulation of cytokine secretion involved in immune response |
| WNT5A | positive regulation of cytokine secretion involved in immune response |
| AURKB | positive regulation of cytokinesis |
| CDC14A | positive regulation of cytokinesis |
| CDC25B | positive regulation of cytokinesis |
| CUL3 | positive regulation of cytokinesis |
| CXCR5 | positive regulation of cytokinesis |
| DRD3 | positive regulation of cytokinesis |
| ECT2 | positive regulation of cytokinesis |
| KIF14 | positive regulation of cytokinesis |
| KIF23 | positive regulation of cytokinesis |
| OR1A2 | positive regulation of cytokinesis |
| RACGAP1 | positive regulation of cytokinesis |
| RXFP3 | positive regulation of cytokinesis |
| CD36 | positive regulation of macrophage cytokine production |
| CD74 | positive regulation of macrophage cytokine production |
| HLA-G | positive regulation of macrophage cytokine production |
| LILRB1 | positive regulation of macrophage cytokine production |
| SEMA7A | positive regulation of macrophage cytokine production |
| SPON2 | positive regulation of macrophage cytokine production |
| TLR4 | positive regulation of macrophage cytokine production |
| WNT5A | positive regulation of macrophage cytokine production |
| FCER1G | positive regulation of mast cell cytokine production |
| NR4A3 | positive regulation of mast cell cytokine production |

| | |
|---|---|
| NUP62 | positive regulation of mitotic cytokinetic process |
| DHX36 | positive regulation of myeloid dendritic cell cytokine production |
| TICAM1 | positive regulation of myeloid dendritic cell cytokine production |
| CD226 | positive regulation of natural killer cell cytokine production |
| CLNK | positive regulation of natural killer cell cytokine production |
| HLA-E | positive regulation of natural killer cell cytokine production |
| HLA-F | positive regulation of natural killer cell cytokine production |
| HLA-G | positive regulation of natural killer cell cytokine production |
| IFIH1 | positive regulation of response to cytokine stimulus |
| WNT5A | positive regulation of response to cytokine stimulus |
| B2M | positive regulation of T cell cytokine production |
| FZD5 | positive regulation of T cell cytokine production |
| SASH3 | positive regulation of T cell cytokine production |
| IL18R1 | positive regulation of T-helper 1 cell cytokine production |
| IL1B | positive regulation of T-helper 1 cell cytokine production |
| CD81 | positive regulation of T-helper 2 cell cytokine production |
| IL6 | positive regulation of T-helper 2 cell cytokine production |
| NLRP3 | positive regulation of T-helper 2 cell cytokine production |
| RSAD2 | positive regulation of T-helper 2 cell cytokine production |
| BTK | regulation of B cell cytokine production |
| GREM2 | regulation of cytokine activity |
| IGF2BP2 | regulation of cytokine biosynthetic process |
| MAP2K3 | regulation of cytokine biosynthetic process |
| ELF1 | regulation of cytokine-mediated signaling pathway |
| RUNX1 | regulation of cytokine-mediated signaling pathway |
| BTN2A1 | regulation of cytokine production |
| BTN2A2 | regulation of cytokine production |
| BTN3A1 | regulation of cytokine production |
| BTN3A2 | regulation of cytokine production |
| BTN3A3 | regulation of cytokine production |
| BTNL2 | regulation of cytokine production |
| BTNL3 | regulation of cytokine production |
| BTNL8 | regulation of cytokine production |
| BTNL9 | regulation of cytokine production |
| ELF1 | regulation of cytokine production |
| ICOSLG | regulation of cytokine production |
| JPH4 | regulation of cytokine production |
| MOG | regulation of cytokine production |
| TRIL | regulation of cytokine production involved in immune response |
| PER1 | regulation of cytokine production involved in inflammatory response |
| CCN4 | regulation of cytokine secretion |
| SOCS1 | regulation of cytokine secretion |
| TLR10 | regulation of cytokine secretion |
| TLR6 | regulation of cytokine secretion |
| TLR8 | regulation of cytokine secretion |
| MIR155 | regulation of cytokine secretion involved in immune response |
| AURKA | regulation of cytokinesis |
| AURKB | regulation of cytokinesis |
| CCP110 | regulation of cytokinesis |
| FLCN | regulation of cytokinesis |

| | |
|---|---|
| KIF13A | regulation of cytokinesis |
| KIF20A | regulation of cytokinesis |
| KLHL13 | regulation of cytokinesis |
| MYO19 | regulation of cytokinesis |
| PLK1 | regulation of cytokinesis |
| PLK2 | regulation of cytokinesis |
| PLK4 | regulation of cytokinesis |
| PRC1 | regulation of cytokinesis |
| RAB11FIP3 | regulation of cytokinesis |
| UVRAG | regulation of cytokinesis |
| ECT2 | regulation of cytokinesis, actomyosin contractile ring assembly |
| TLR3 | regulation of dendritic cell cytokine production |
| TLR4 | regulation of dendritic cell cytokine production |
| CCR2 | regulation of T cell cytokine production |
| CLC | regulation of T cell cytokine production |
| ACP5 | response to cytokine |
| AIF1 | response to cytokine |
| ALDH1A2 | response to cytokine |
| AVPR2 | response to cytokine |
| CCL5 | response to cytokine |
| FOSL1 | response to cytokine |
| FOS | response to cytokine |
| IL6R | response to cytokine |
| IL6ST | response to cytokine |
| ITIH4 | response to cytokine |
| JUNB | response to cytokine |
| JUN | response to cytokine |
| MAPKAPK3 | response to cytokine |
| MCL1 | response to cytokine |
| NFKB1 | response to cytokine |
| NFKB2 | response to cytokine |
| OXTR | response to cytokine |
| PLA2G5 | response to cytokine |
| PML | response to cytokine |
| PTGS2 | response to cytokine |
| RARA | response to cytokine |
| RELB | response to cytokine |
| REL | response to cytokine |
| SRF | response to cytokine |
| STAT1 | response to cytokine |
| SYNJ1 | response to cytokine |
| TIMP3 | response to cytokine |
| TIMP4 | response to cytokine |
| TYMS | response to cytokine |
| XCR1 | response to cytokine |
| IL12A | T-helper 1 cell cytokine production |
| IL18RAP | T-helper 1 cell cytokine production |
| DENND1B | T-helper 2 cell cytokine production |
| IL31RA | T-helper 2 cell cytokine production |
| IL4 | T-helper 2 cell cytokine production |

| Inflammation related genes | Associated terms |
|---|---|
| C4B | inflammatory response |
| F3 | activation of plasma proteins involved in acute inflammatory response |
| APOA2 | acute inflammatory response |
| NUPR1 | acute inflammatory response |
| OGG1 | acute inflammatory response |
| TREM1 | acute inflammatory response |
| CD6 | acute inflammatory response to antigenic stimulus |
| ICAM1 | acute inflammatory response to antigenic stimulus |
| IL31RA | acute inflammatory response to antigenic stimulus |
| OPRM1 | acute inflammatory response to antigenic stimulus |
| SERPINC1 | acute inflammatory response to antigenic stimulus |
| CCL11 | chronic inflammatory response |
| CXCL13 | chronic inflammatory response |
| THBS1 | chronic inflammatory response |
| TNF | chronic inflammatory response to antigenic stimulus |
| IL1A | connective tissue replacement involved in inflammatory response wound healing |
| CD96 | cytokine production involved in inflammatory response |
| IDO1 | cytokine production involved in inflammatory response |
| FASLG | inflammatory cell apoptotic process |
| ACKR2 | inflammatory response |
| ACOD1 | inflammatory response |
| ADAM8 | inflammatory response |
| ADORA2A | inflammatory response |
| ADORA3 | inflammatory response |
| AFAP1L2 | inflammatory response |
| AGER | inflammatory response |
| AGTR1 | inflammatory response |
| AIF1 | inflammatory response |
| AIM2 | inflammatory response |
| AOC3 | inflammatory response |
| APOL3 | inflammatory response |
| AXL | inflammatory response |
| BCL6 | inflammatory response |
| BMPR1B | inflammatory response |
| C3AR1 | inflammatory response |
| C3 | inflammatory response |
| C4A | inflammatory response |
| C4B | inflammatory response |
| CALCA | inflammatory response |
| CAMK1D | inflammatory response |
| CCL11 | inflammatory response |
| CCL14 | inflammatory response |

| | |
|---|---|
| CCL15-CCL14 | inflammatory response |
| CCL16 | inflammatory response |
| CCL17 | inflammatory response |
| CCL20 | inflammatory response |
| CCL22 | inflammatory response |
| CCL2 | inflammatory response |
| CCL4 | inflammatory response |
| CCL5 | inflammatory response |
| CCL8 | inflammatory response |
| CCR1 | inflammatory response |
| CCR2 | inflammatory response |
| CCR5 | inflammatory response |
| CCR7 | inflammatory response |
| CD163 | inflammatory response |
| CD180 | inflammatory response |
| CD40 | inflammatory response |
| CD44 | inflammatory response |
| CD5L | inflammatory response |
| CHI3L1 | inflammatory response |
| CHUK | inflammatory response |
| CIITA | inflammatory response |
| CLEC7A | inflammatory response |
| CMKLR1 | inflammatory response |
| CNR2 | inflammatory response |
| CRP | inflammatory response |
| CSF1 | inflammatory response |
| CSF1R | inflammatory response |
| CSRP3 | inflammatory response |
| CX3CL1 | inflammatory response |
| CXCL10 | inflammatory response |
| CXCL11 | inflammatory response |
| CXCL13 | inflammatory response |
| CXCL1 | inflammatory response |
| CXCL2 | inflammatory response |
| CXCL3 | inflammatory response |
| CXCL6 | inflammatory response |
| CXCL8 | inflammatory response |
| CXCL9 | inflammatory response |
| CXCR3 | inflammatory response |
| CXCR4 | inflammatory response |
| CXCR6 | inflammatory response |
| DAB2IP | inflammatory response |
| FCGR2B | inflammatory response |
| FOS | inflammatory response |
| FPR2 | inflammatory response |
| FPR3 | inflammatory response |
| FUT7 | inflammatory response |
| GBP5 | inflammatory response |
| GPER1 | inflammatory response |
| GPR68 | inflammatory response |

| | |
|---|---|
| HDAC4 | inflammatory response |
| HMGB2 | inflammatory response |
| HRH1 | inflammatory response |
| HRH4 | inflammatory response |
| HYAL3 | inflammatory response |
| IDO1 | inflammatory response |
| IFI16 | inflammatory response |
| IL13 | inflammatory response |
| IL15 | inflammatory response |
| IL17C | inflammatory response |
| IL18R1 | inflammatory response |
| IL18RAP | inflammatory response |
| IL1A | inflammatory response |
| IL1B | inflammatory response |
| IL1RAP | inflammatory response |
| IL1RL2 | inflammatory response |
| IL1RN | inflammatory response |
| IL23A | inflammatory response |
| IL23R | inflammatory response |
| IL2RA | inflammatory response |
| IL34 | inflammatory response |
| IL36RN | inflammatory response |
| IL37 | inflammatory response |
| IL6 | inflammatory response |
| IRGM | inflammatory response |
| ITGB2 | inflammatory response |
| KDM6B | inflammatory response |
| KNG1 | inflammatory response |
| KRT16 | inflammatory response |
| LGALS9 | inflammatory response |
| LTB4R2 | inflammatory response |
| LTB4R | inflammatory response |
| LYZ | inflammatory response |
| MAP2K3 | inflammatory response |
| MEFV | inflammatory response |
| MMP25 | inflammatory response |
| MS4A2 | inflammatory response |
| MYD88 | inflammatory response |
| NCR3 | inflammatory response |
| NDST1 | inflammatory response |
| NFAM1 | inflammatory response |
| NFATC3 | inflammatory response |
| NFKB1 | inflammatory response |
| NFKB2 | inflammatory response |
| NFKBID | inflammatory response |
| NLRP1 | inflammatory response |
| NLRP3 | inflammatory response |
| NLRP4 | inflammatory response |
| NMI | inflammatory response |
| NOD1 | inflammatory response |

| | |
|---|---|
| NOX1 | inflammatory response |
| NRROS | inflammatory response |
| ORM1 | inflammatory response |
| PIK3CG | inflammatory response |
| PLA2G2D | inflammatory response |
| PLA2G2E | inflammatory response |
| PLA2G4C | inflammatory response |
| POLB | inflammatory response |
| PPBP | inflammatory response |
| PROK2 | inflammatory response |
| PTGDR | inflammatory response |
| PTGER1 | inflammatory response |
| PTGER3 | inflammatory response |
| PTGER4 | inflammatory response |
| PTGS1 | inflammatory response |
| PTGS2 | inflammatory response |
| PTX3 | inflammatory response |
| PXK | inflammatory response |
| RARRES2 | inflammatory response |
| REG3A | inflammatory response |
| RELB | inflammatory response |
| REL | inflammatory response |
| RIPK2 | inflammatory response |
| S1PR3 | inflammatory response |
| SCG2 | inflammatory response |
| SELE | inflammatory response |
| SEMA7A | inflammatory response |
| STAB1 | inflammatory response |
| TACR1 | inflammatory response |
| TBXA2R | inflammatory response |
| THBS1 | inflammatory response |
| THEMIS2 | inflammatory response |
| TICAM1 | inflammatory response |
| TLR10 | inflammatory response |
| TLR3 | inflammatory response |
| TLR4 | inflammatory response |
| TLR6 | inflammatory response |
| TLR8 | inflammatory response |
| TMIGD3 | inflammatory response |
| TNFAIP3 | inflammatory response |
| TNF | inflammatory response |
| TNFRSF1A | inflammatory response |
| TNIP1 | inflammatory response |
| TOLLIP | inflammatory response |
| TREX1 | inflammatory response |
| TRIL | inflammatory response |
| TRPV1 | inflammatory response |
| TSPAN2 | inflammatory response |
| XCR1 | inflammatory response |
| ZC3H12A | inflammatory response |

| | |
|---|---|
| CYSLTR1 | inflammatory response to antigenic stimulus |
| HLA-DRB1 | inflammatory response to antigenic stimulus |
| HMGB2 | inflammatory response to antigenic stimulus |
| IL1A | inflammatory response to antigenic stimulus |
| IL1B | inflammatory response to antigenic stimulus |
| IL1RN | inflammatory response to antigenic stimulus |
| IL20RB | inflammatory response to antigenic stimulus |
| IL2RA | inflammatory response to antigenic stimulus |
| IL36RN | inflammatory response to antigenic stimulus |
| IL37 | inflammatory response to antigenic stimulus |
| IL5RA | inflammatory response to antigenic stimulus |
| KDM6B | inflammatory response to antigenic stimulus |
| NOTCH1 | inflammatory response to antigenic stimulus |
| RBPJ | inflammatory response to antigenic stimulus |
| TREX1 | inflammatory response to antigenic stimulus |
| CCR2 | inflammatory response to wounding |
| LBP | leukocyte chemotaxis involved in inflammatory response |
| SLAMF8 | leukocyte chemotaxis involved in inflammatory response |
| ADAM8 | leukocyte migration involved in inflammatory response |
| CCR6 | leukocyte migration involved in inflammatory response |
| CX3CL1 | leukocyte migration involved in inflammatory response |
| FUT7 | leukocyte migration involved in inflammatory response |
| ITGB2 | leukocyte migration involved in inflammatory response |
| JAM3 | leukocyte migration involved in inflammatory response |
| SELE | leukocyte migration involved in inflammatory response |
| APCS | negative regulation of acute inflammatory response |
| NLRP3 | negative regulation of acute inflammatory response |
| PPARG | negative regulation of acute inflammatory response |
| FCGR2B | negative regulation of acute inflammatory response to antigenic stimulus |
| CYP19A1 | negative regulation of chronic inflammatory response |
| FOXP3 | negative regulation of chronic inflammatory response |
| TNFAIP3 | negative regulation of chronic inflammatory response |
| AGER | negative regulation of connective tissue replacement involved in inflammatory response wound healing |
| ABCD2 | negative regulation of cytokine production involved in inflammatory response |
| ADCY7 | negative regulation of cytokine production involved in inflammatory response |
| APOD | negative regulation of cytokine production involved in inflammatory response |
| F2 | negative regulation of cytokine production involved in inflammatory response |
| IL1R2 | negative regulation of cytokine production involved in inflammatory response |
| MEFV | negative regulation of cytokine production involved in inflammatory response |
| MIR155 | negative regulation of cytokine production involved in inflammatory response |
| ZC3H12A | negative regulation of cytokine production involved in inflammatory response |
| ABR | negative regulation of inflammatory response |
| ACOD1 | negative regulation of inflammatory response |
| ACP5 | negative regulation of inflammatory response |
| ADORA2A | negative regulation of inflammatory response |
| APOA1 | negative regulation of inflammatory response |
| APOE | negative regulation of inflammatory response |
| BCR | negative regulation of inflammatory response |
| C1QTNF3 | negative regulation of inflammatory response |

| | |
|---|---|
| CNR2 | negative regulation of inflammatory response |
| CXCL17 | negative regulation of inflammatory response |
| FFAR4 | negative regulation of inflammatory response |
| FOXF1 | negative regulation of inflammatory response |
| FOXP3 | negative regulation of inflammatory response |
| FPR2 | negative regulation of inflammatory response |
| GHRL | negative regulation of inflammatory response |
| GHSR | negative regulation of inflammatory response |
| GPER1 | negative regulation of inflammatory response |
| HGF | negative regulation of inflammatory response |
| IL2RA | negative regulation of inflammatory response |
| KLF4 | negative regulation of inflammatory response |
| KRT1 | negative regulation of inflammatory response |
| MAPK7 | negative regulation of inflammatory response |
| MEFV | negative regulation of inflammatory response |
| MIR147B | negative regulation of inflammatory response |
| MIR155 | negative regulation of inflammatory response |
| MIR205 | negative regulation of inflammatory response |
| MIR223 | negative regulation of inflammatory response |
| MVK | negative regulation of inflammatory response |
| NFKB1 | negative regulation of inflammatory response |
| NLRP3 | negative regulation of inflammatory response |
| NR1D1 | negative regulation of inflammatory response |
| PBK | negative regulation of inflammatory response |
| PPARG | negative regulation of inflammatory response |
| PTGER4 | negative regulation of inflammatory response |
| PTPN2 | negative regulation of inflammatory response |
| SAA1 | negative regulation of inflammatory response |
| SOCS3 | negative regulation of inflammatory response |
| TEK | negative regulation of inflammatory response |
| TNFAIP3 | negative regulation of inflammatory response |
| TNFRSF1A | negative regulation of inflammatory response |
| ZFP36 | negative regulation of inflammatory response |
| GPR17 | negative regulation of inflammatory response to antigenic stimulus |
| HLA-DRB1 | negative regulation of inflammatory response to antigenic stimulus |
| SIGLEC10 | negative regulation of inflammatory response to wounding |
| ARG2 | negative regulation of macrophage inflammatory protein 1 alpha production |
| MEFV | negative regulation of macrophage inflammatory protein 1 alpha production |
| CD200 | negative regulation of neuroinflammatory response |
| CD200R1 | negative regulation of neuroinflammatory response |
| IGF1 | negative regulation of neuroinflammatory response |
| IL4 | negative regulation of neuroinflammatory response |
| MIR195 | negative regulation of neuroinflammatory response |
| NR1D1 | negative regulation of neuroinflammatory response |
| MEFV | negative regulation of NLRP3 inflammasome complex assembly |
| DUSP10 | negative regulation of respiratory burst involved in inflammatory response |
| SLAMF8 | negative regulation of respiratory burst involved in inflammatory response |
| ADCY1 | neuroinflammatory response |
| ADCY8 | neuroinflammatory response |
| IL4 | neuroinflammatory response |

| | |
|---|---|
| TLR4 | nitric oxide production involved in inflammatory response |
| NLRP1 | NLRP1 inflammasome complex assembly |
| NLRP3 | NLRP3 inflammasome complex assembly |
| ADAM8 | positive regulation of acute inflammatory response |
| C2CD4A | positive regulation of acute inflammatory response |
| CREB3L3 | positive regulation of acute inflammatory response |
| IL6 | positive regulation of acute inflammatory response |
| IL6ST | positive regulation of acute inflammatory response |
| PIK3CG | positive regulation of acute inflammatory response |
| IDO1 | positive regulation of chronic inflammatory response |
| TNF | positive regulation of chronic inflammatory response to antigenic stimulus |
| CD6 | positive regulation of cytokine production involved in inflammatory response |
| CLEC7A | positive regulation of cytokine production involved in inflammatory response |
| GBP5 | positive regulation of cytokine production involved in inflammatory response |
| KARS1 | positive regulation of cytokine production involved in inflammatory response |
| MIR21 | positive regulation of cytokine production involved in inflammatory response |
| MYD88 | positive regulation of cytokine production involved in inflammatory response |
| TICAM1 | positive regulation of cytokine production involved in inflammatory response |
| TLR4 | positive regulation of cytokine production involved in inflammatory response |
| TLR6 | positive regulation of cytokine production involved in inflammatory response |
| ADAM8 | positive regulation of inflammatory response |
| AGTR1 | positive regulation of inflammatory response |
| CCN4 | positive regulation of inflammatory response |
| CCR2 | positive regulation of inflammatory response |
| CD47 | positive regulation of inflammatory response |
| CX3CL1 | positive regulation of inflammatory response |
| EGFR | positive regulation of inflammatory response |
| FABP4 | positive regulation of inflammatory response |
| IL15 | positive regulation of inflammatory response |
| IL17RB | positive regulation of inflammatory response |
| IL1B | positive regulation of inflammatory response |
| IL1RL1 | positive regulation of inflammatory response |
| IL21 | positive regulation of inflammatory response |
| IL23A | positive regulation of inflammatory response |
| IL33 | positive regulation of inflammatory response |
| JAK2 | positive regulation of inflammatory response |
| MIR155 | positive regulation of inflammatory response |
| MIR21 | positive regulation of inflammatory response |
| NAPEPLD | positive regulation of inflammatory response |
| NFKBIA | positive regulation of inflammatory response |
| PLA2G2A | positive regulation of inflammatory response |
| PTGER4 | positive regulation of inflammatory response |
| SERPINE1 | positive regulation of inflammatory response |
| SNCA | positive regulation of inflammatory response |
| STAT5B | positive regulation of inflammatory response |
| TGM2 | positive regulation of inflammatory response |
| TLR10 | positive regulation of inflammatory response |
| TLR3 | positive regulation of inflammatory response |
| TLR4 | positive regulation of inflammatory response |
| TNF | positive regulation of inflammatory response |

| | |
|---|---|
| TNFRSF1A | positive regulation of inflammatory response |
| TNIP1 | positive regulation of inflammatory response |
| TRPV4 | positive regulation of inflammatory response |
| WNT5A | positive regulation of inflammatory response |
| CD28 | positive regulation of inflammatory response to antigenic stimulus |
| CD81 | positive regulation of inflammatory response to antigenic stimulus |
| MIR21 | positive regulation of inflammatory response to wounding |
| SERPINE1 | positive regulation of leukotriene production involved in inflammatory response |
| TRPV4 | positive regulation of macrophage inflammatory protein 1 alpha production |
| IL1B | positive regulation of neuroinflammatory response |
| IL33 | positive regulation of neuroinflammatory response |
| IL6 | positive regulation of neuroinflammatory response |
| NUPR1 | positive regulation of neuroinflammatory response |
| TNF | positive regulation of neuroinflammatory response |
| CD36 | positive regulation of NLRP3 inflammasome complex assembly |
| DDX3X | positive regulation of NLRP3 inflammasome complex assembly |
| GBP5 | positive regulation of NLRP3 inflammasome complex assembly |
| TLR4 | positive regulation of NLRP3 inflammasome complex assembly |
| TLR6 | positive regulation of NLRP3 inflammasome complex assembly |
| LBP | positive regulation of respiratory burst involved in inflammatory response |
| CD36 | production of molecular mediator involved in inflammatory response |
| IL4R | production of molecular mediator involved in inflammatory response |
| DNASE1 | regulation of acute inflammatory response |
| CCL5 | regulation of chronic inflammatory response |
| PER1 | regulation of cytokine production involved in inflammatory response |
| ABHD12 | regulation of inflammatory response |
| AGER | regulation of inflammatory response |
| AGTR1 | regulation of inflammatory response |
| AKNA | regulation of inflammatory response |
| BCL6B | regulation of inflammatory response |
| BCL6 | regulation of inflammatory response |
| BIRC3 | regulation of inflammatory response |
| BRD4 | regulation of inflammatory response |
| CASP1 | regulation of inflammatory response |
| CCR2 | regulation of inflammatory response |
| CYLD | regulation of inflammatory response |
| DUOXA1 | regulation of inflammatory response |
| DUOXA2 | regulation of inflammatory response |
| ESR1 | regulation of inflammatory response |
| FABP4 | regulation of inflammatory response |
| FANCA | regulation of inflammatory response |
| GGT1 | regulation of inflammatory response |
| IL1RL2 | regulation of inflammatory response |
| IL20 | regulation of inflammatory response |
| JAK2 | regulation of inflammatory response |
| MAS1 | regulation of inflammatory response |
| MCPH1 | regulation of inflammatory response |
| MYD88 | regulation of inflammatory response |
| NLRP1 | regulation of inflammatory response |
| NLRP3 | regulation of inflammatory response |

| | |
|---|---|
| PIK3AP1 | regulation of inflammatory response |
| PTGS2 | regulation of inflammatory response |
| RICTOR | regulation of inflammatory response |
| SBNO2 | regulation of inflammatory response |
| SELE | regulation of inflammatory response |
| SEMA7A | regulation of inflammatory response |
| SPATA2 | regulation of inflammatory response |
| STING1 | regulation of inflammatory response |
| TLR4 | regulation of inflammatory response |
| TNF | regulation of inflammatory response |
| TNIP1 | regulation of inflammatory response |
| TREX1 | regulation of inflammatory response |
| USP18 | regulation of inflammatory response |
| WNT5A | regulation of inflammatory response |
| CD200R1 | regulation of neuroinflammatory response |
| CD200 | regulation of neuroinflammatory response |
| IL6 | regulation of neuroinflammatory response |
| PTGS2 | regulation of neuroinflammatory response |
| C2CD4A | regulation of vascular permeability involved in acute inflammatory response |
| MYLK3 | regulation of vascular permeability involved in acute inflammatory response |

| Integrin related genes | Associated terms |
|---|---|
| CD14 | Beta-1 integrin cell surface interactions |
| CD81 | Beta-1 integrin cell surface interactions |
| COL1A1 | Beta-1 integrin cell surface interactions |
| COL1A2 | Beta-1 integrin cell surface interactions |
| COL2A1 | Beta-1 integrin cell surface interactions |
| COL3A1 | Beta-1 integrin cell surface interactions |
| COL4A1 | Beta-1 integrin cell surface interactions |
| COL4A3 | Beta-1 integrin cell surface interactions |
| COL4A4 | Beta-1 integrin cell surface interactions |
| COL4A5 | Beta-1 integrin cell surface interactions |
| COL4A6 | Beta-1 integrin cell surface interactions |
| COL5A1 | Beta-1 integrin cell surface interactions |
| COL5A2 | Beta-1 integrin cell surface interactions |
| COL6A1 | Beta-1 integrin cell surface interactions |
| COL6A2 | Beta-1 integrin cell surface interactions |
| COL6A3 | Beta-1 integrin cell surface interactions |
| COL7A1 | Beta-1 integrin cell surface interactions |
| COL11A1 | Beta-1 integrin cell surface interactions |
| COL11A2 | Beta-1 integrin cell surface interactions |
| CSPG4 | Beta-1 integrin cell surface interactions |
| F13A1 | Beta-1 integrin cell surface interactions |
| FBN1 | Beta-1 integrin cell surface interactions |

| | |
|---|---|
| FGA | Beta-1 integrin cell surface interactions |
| FGB | Beta-1 integrin cell surface interactions |
| FGG | Beta-1 integrin cell surface interactions |
| FN1 | Beta-1 integrin cell surface interactions |
| TNC | Beta-1 integrin cell surface interactions |
| ITGA6 | Beta-1 integrin cell surface interactions |
| ITGA1 | Beta-1 integrin cell surface interactions |
| ITGA2 | Beta-1 integrin cell surface interactions |
| ITGA3 | Beta-1 integrin cell surface interactions |
| ITGA4 | Beta-1 integrin cell surface interactions |
| ITGA5 | Beta-1 integrin cell surface interactions |
| ITGA7 | Beta-1 integrin cell surface interactions |
| ITGA9 | Beta-1 integrin cell surface interactions |
| ITGAV | Beta-1 integrin cell surface interactions |
| ITGB1 | Beta-1 integrin cell surface interactions |
| LAMA2 | Beta-1 integrin cell surface interactions |
| LAMA3 | Beta-1 integrin cell surface interactions |
| LAMA4 | Beta-1 integrin cell surface interactions |
| LAMA5 | Beta-1 integrin cell surface interactions |
| LAMB1 | Beta-1 integrin cell surface interactions |
| LAMB2 | Beta-1 integrin cell surface interactions |
| LAMB3 | Beta-1 integrin cell surface interactions |
| LAMC1 | Beta-1 integrin cell surface interactions |
| LAMC2 | Beta-1 integrin cell surface interactions |
| MDK | Beta-1 integrin cell surface interactions |
| NID1 | Beta-1 integrin cell surface interactions |
| PLAU | Beta-1 integrin cell surface interactions |
| PLAUR | Beta-1 integrin cell surface interactions |
| SPP1 | Beta-1 integrin cell surface interactions |
| TGFBI | Beta-1 integrin cell surface interactions |
| TGM2 | Beta-1 integrin cell surface interactions |
| THBS1 | Beta-1 integrin cell surface interactions |
| THBS2 | Beta-1 integrin cell surface interactions |
| VCAM1 | Beta-1 integrin cell surface interactions |
| VEGFA | Beta-1 integrin cell surface interactions |
| VTN | Beta-1 integrin cell surface interactions |
| ITGA10 | Beta-1 integrin cell surface interactions |
| ITGA8 | Beta-1 integrin cell surface interactions |
| ITGA11 | Beta-1 integrin cell surface interactions |
| JAM2 | Beta-1 integrin cell surface interactions |
| COL18A1 | Beta-1 integrin cell surface interactions |
| IGSF8 | Beta-1 integrin cell surface interactions |
| NPNT | Beta-1 integrin cell surface interactions |
| LAMA1 | Beta-1 integrin cell surface interactions |
| CDH1 | Alpha-E beta-7 integrin cell surface interactions |
| ITGAE | Alpha-E beta-7 integrin cell surface interactions |
| ITGB7 | Alpha-E beta-7 integrin cell surface interactions |
| CD47 | Beta-3 integrin cell surface interactions |
| COL1A1 | Beta-3 integrin cell surface interactions |
| COL1A2 | Beta-3 integrin cell surface interactions |

| | |
|---|---|
| COL4A1 | Beta-3 integrin cell surface interactions |
| COL4A3 | Beta-3 integrin cell surface interactions |
| COL4A4 | Beta-3 integrin cell surface interactions |
| COL4A5 | Beta-3 integrin cell surface interactions |
| COL4A6 | Beta-3 integrin cell surface interactions |
| FBN1 | Beta-3 integrin cell surface interactions |
| FGA | Beta-3 integrin cell surface interactions |
| FGB | Beta-3 integrin cell surface interactions |
| FGG | Beta-3 integrin cell surface interactions |
| FN1 | Beta-3 integrin cell surface interactions |
| HMGB1 | Beta-3 integrin cell surface interactions |
| TNC | Beta-3 integrin cell surface interactions |
| IBSP | Beta-3 integrin cell surface interactions |
| CYR61 | Beta-3 integrin cell surface interactions |
| ITGA2B | Beta-3 integrin cell surface interactions |
| ITGAV | Beta-3 integrin cell surface interactions |
| ITGB3 | Beta-3 integrin cell surface interactions |
| KDR | Beta-3 integrin cell surface interactions |
| L1CAM | Beta-3 integrin cell surface interactions |
| LAMA4 | Beta-3 integrin cell surface interactions |
| LAMB1 | Beta-3 integrin cell surface interactions |
| LAMC1 | Beta-3 integrin cell surface interactions |
| PDGFB | Beta-3 integrin cell surface interactions |
| PDGFRB | Beta-3 integrin cell surface interactions |
| PECAM1 | Beta-3 integrin cell surface interactions |
| PLAU | Beta-3 integrin cell surface interactions |
| PLAUR | Beta-3 integrin cell surface interactions |
| PVR | Beta-3 integrin cell surface interactions |
| SDC1 | Beta-3 integrin cell surface interactions |
| SDC4 | Beta-3 integrin cell surface interactions |
| SPP1 | Beta-3 integrin cell surface interactions |
| TGFBI | Beta-3 integrin cell surface interactions |
| TGFBR2 | Beta-3 integrin cell surface interactions |
| THBS1 | Beta-3 integrin cell surface interactions |
| THY1 | Beta-3 integrin cell surface interactions |
| VEGFA | Beta-3 integrin cell surface interactions |
| VTN | Beta-3 integrin cell surface interactions |
| SPHK1 | Beta-3 integrin cell surface interactions |
| EDIL3 | Beta-3 integrin cell surface interactions |
| F11R | Beta-3 integrin cell surface interactions |
| RHOA | Alpha-V beta-3 integrin/OPN pathway |
| CD44 | Alpha-V beta-3 integrin/OPN pathway |
| CDC42 | Alpha-V beta-3 integrin/OPN pathway |
| CHUK | Alpha-V beta-3 integrin/OPN pathway |
| PTK2B | Alpha-V beta-3 integrin/OPN pathway |
| FOS | Alpha-V beta-3 integrin/OPN pathway |
| GSN | Alpha-V beta-3 integrin/OPN pathway |
| ILK | Alpha-V beta-3 integrin/OPN pathway |
| ITGAV | Alpha-V beta-3 integrin/OPN pathway |
| ITGB3 | Alpha-V beta-3 integrin/OPN pathway |

| | |
|---|---|
| JUN | Alpha-V beta-3 integrin/OPN pathway |
| MAP3K1 | Alpha-V beta-3 integrin/OPN pathway |
| MMP2 | Alpha-V beta-3 integrin/OPN pathway |
| MMP9 | Alpha-V beta-3 integrin/OPN pathway |
| NFKB1 | Alpha-V beta-3 integrin/OPN pathway |
| NFKBIA | Alpha-V beta-3 integrin/OPN pathway |
| PIK3CA | Alpha-V beta-3 integrin/OPN pathway |
| PIK3R1 | Alpha-V beta-3 integrin/OPN pathway |
| PLAU | Alpha-V beta-3 integrin/OPN pathway |
| MAPK1 | Alpha-V beta-3 integrin/OPN pathway |
| MAPK3 | Alpha-V beta-3 integrin/OPN pathway |
| MAPK8 | Alpha-V beta-3 integrin/OPN pathway |
| RAC1 | Alpha-V beta-3 integrin/OPN pathway |
| RELA | Alpha-V beta-3 integrin/OPN pathway |
| SPP1 | Alpha-V beta-3 integrin/OPN pathway |
| SYK | Alpha-V beta-3 integrin/OPN pathway |
| PIP5K1A | Alpha-V beta-3 integrin/OPN pathway |
| MAP3K14 | Alpha-V beta-3 integrin/OPN pathway |
| ROCK2 | Alpha-V beta-3 integrin/OPN pathway |
| BCAR1 | Alpha-V beta-3 integrin/OPN pathway |
| VAV3 | Alpha-V beta-3 integrin/OPN pathway |
| ADRB2 | Arf6 integrin-mediated signaling pathway |
| AGTR1 | Arf6 integrin-mediated signaling pathway |
| BIN1 | Arf6 integrin-mediated signaling pathway |
| ARF6 | Arf6 integrin-mediated signaling pathway |
| AVPR2 | Arf6 integrin-mediated signaling pathway |
| CDH1 | Arf6 integrin-mediated signaling pathway |
| CLTC | Arf6 integrin-mediated signaling pathway |
| CPE | Arf6 integrin-mediated signaling pathway |
| CTNNA1 | Arf6 integrin-mediated signaling pathway |
| CTNNB1 | Arf6 integrin-mediated signaling pathway |
| CTNND1 | Arf6 integrin-mediated signaling pathway |
| DNM2 | Arf6 integrin-mediated signaling pathway |
| EDNRB | Arf6 integrin-mediated signaling pathway |
| IL2RA | Arf6 integrin-mediated signaling pathway |
| INS | Arf6 integrin-mediated signaling pathway |
| ITGA6 | Arf6 integrin-mediated signaling pathway |
| ITGA1 | Arf6 integrin-mediated signaling pathway |
| ITGA2 | Arf6 integrin-mediated signaling pathway |
| ITGA3 | Arf6 integrin-mediated signaling pathway |
| ITGA4 | Arf6 integrin-mediated signaling pathway |
| ITGA5 | Arf6 integrin-mediated signaling pathway |
| ITGA7 | Arf6 integrin-mediated signaling pathway |
| ITGA9 | Arf6 integrin-mediated signaling pathway |
| ITGAV | Arf6 integrin-mediated signaling pathway |
| ITGB1 | Arf6 integrin-mediated signaling pathway |
| KLC1 | Arf6 integrin-mediated signaling pathway |
| NME1 | Arf6 integrin-mediated signaling pathway |
| PLD1 | Arf6 integrin-mediated signaling pathway |
| PLD2 | Arf6 integrin-mediated signaling pathway |

| | |
|---|---|
| RALA | Arf6 integrin-mediated signaling pathway |
| SLC2A4 | Arf6 integrin-mediated signaling pathway |
| TSHR | Arf6 integrin-mediated signaling pathway |
| ITGA10 | Arf6 integrin-mediated signaling pathway |
| ITGA8 | Arf6 integrin-mediated signaling pathway |
| ASAP2 | Arf6 integrin-mediated signaling pathway |
| SPAG9 | Arf6 integrin-mediated signaling pathway |
| VAMP3 | Arf6 integrin-mediated signaling pathway |
| ACAP1 | Arf6 integrin-mediated signaling pathway |
| SCAMP2 | Arf6 integrin-mediated signaling pathway |
| EXOC5 | Arf6 integrin-mediated signaling pathway |
| EXOC3 | Arf6 integrin-mediated signaling pathway |
| ITGA11 | Arf6 integrin-mediated signaling pathway |
| MAPK8IP3 | Arf6 integrin-mediated signaling pathway |
| EXOC7 | Arf6 integrin-mediated signaling pathway |
| PIP5K1C | Arf6 integrin-mediated signaling pathway |
| EXOC6 | Arf6 integrin-mediated signaling pathway |
| EXOC1 | Arf6 integrin-mediated signaling pathway |
| EXOC2 | Arf6 integrin-mediated signaling pathway |
| EXOC4 | Arf6 integrin-mediated signaling pathway |
| ADAM8 | Alpha-9 beta-1 integrin pathway |
| CSF2 | Alpha-9 beta-1 integrin pathway |
| CSF2RA | Alpha-9 beta-1 integrin pathway |
| F13A1 | Alpha-9 beta-1 integrin pathway |
| FIGF | Alpha-9 beta-1 integrin pathway |
| FN1 | Alpha-9 beta-1 integrin pathway |
| ADAM2 | Alpha-9 beta-1 integrin pathway |
| TNC | Alpha-9 beta-1 integrin pathway |
| ITGA9 | Alpha-9 beta-1 integrin pathway |
| ITGB1 | Alpha-9 beta-1 integrin pathway |
| KCNJ15 | Alpha-9 beta-1 integrin pathway |
| NOS2 | Alpha-9 beta-1 integrin pathway |
| PXN | Alpha-9 beta-1 integrin pathway |
| RAC1 | Alpha-9 beta-1 integrin pathway |
| SAT1 | Alpha-9 beta-1 integrin pathway |
| SPP1 | Alpha-9 beta-1 integrin pathway |
| SRC | Alpha-9 beta-1 integrin pathway |
| TGM2 | Alpha-9 beta-1 integrin pathway |
| VCAM1 | Alpha-9 beta-1 integrin pathway |
| VEGFA | Alpha-9 beta-1 integrin pathway |
| VEGFC | Alpha-9 beta-1 integrin pathway |
| ADAM12 | Alpha-9 beta-1 integrin pathway |
| ADAM15 | Alpha-9 beta-1 integrin pathway |
| BCAR1 | Alpha-9 beta-1 integrin pathway |
| PAOX | Alpha-9 beta-1 integrin pathway |
| AGER | Alpha-M beta-2 integrin signaling |
| AKT1 | Alpha-M beta-2 integrin signaling |
| APOB | Alpha-M beta-2 integrin signaling |
| RHOA | Alpha-M beta-2 integrin signaling |
| BLK | Alpha-M beta-2 integrin signaling |

| | |
|---|---|
| CTGF | Alpha-M beta-2 integrin signaling |
| FGR | Alpha-M beta-2 integrin signaling |
| FYN | Alpha-M beta-2 integrin signaling |
| HCK | Alpha-M beta-2 integrin signaling |
| HMGB1 | Alpha-M beta-2 integrin signaling |
| ICAM1 | Alpha-M beta-2 integrin signaling |
| IL6 | Alpha-M beta-2 integrin signaling |
| ITGAM | Alpha-M beta-2 integrin signaling |
| ITGB2 | Alpha-M beta-2 integrin signaling |
| LCK | Alpha-M beta-2 integrin signaling |
| LPA | Alpha-M beta-2 integrin signaling |
| LRP1 | Alpha-M beta-2 integrin signaling |
| LYN | Alpha-M beta-2 integrin signaling |
| MMP2 | Alpha-M beta-2 integrin signaling |
| MMP9 | Alpha-M beta-2 integrin signaling |
| MST1 | Alpha-M beta-2 integrin signaling |
| MST1R | Alpha-M beta-2 integrin signaling |
| MYH2 | Alpha-M beta-2 integrin signaling |
| NFKB1 | Alpha-M beta-2 integrin signaling |
| PLAT | Alpha-M beta-2 integrin signaling |
| PLAU | Alpha-M beta-2 integrin signaling |
| PLAUR | Alpha-M beta-2 integrin signaling |
| PLG | Alpha-M beta-2 integrin signaling |
| PRKCZ | Alpha-M beta-2 integrin signaling |
| RAP1A | Alpha-M beta-2 integrin signaling |
| RAP1B | Alpha-M beta-2 integrin signaling |
| ROCK1 | Alpha-M beta-2 integrin signaling |
| SELP | Alpha-M beta-2 integrin signaling |
| SELPLG | Alpha-M beta-2 integrin signaling |
| SRC | Alpha-M beta-2 integrin signaling |
| THY1 | Alpha-M beta-2 integrin signaling |
| TLN1 | Alpha-M beta-2 integrin signaling |
| TNF | Alpha-M beta-2 integrin signaling |
| YES1 | Alpha-M beta-2 integrin signaling |
| JAM2 | Alpha-M beta-2 integrin signaling |
| JAM3 | Alpha-M beta-2 integrin signaling |
| RHOA | Alpha-4 beta-7 integrin signaling |
| CD44 | Alpha-4 beta-7 integrin signaling |
| ITGA4 | Alpha-4 beta-7 integrin signaling |
| ITGB1 | Alpha-4 beta-7 integrin signaling |
| ITGB7 | Alpha-4 beta-7 integrin signaling |
| PTK2 | Alpha-4 beta-7 integrin signaling |
| PXN | Alpha-4 beta-7 integrin signaling |
| VCAM1 | Alpha-4 beta-7 integrin signaling |
| MADCAM1 | Alpha-4 beta-7 integrin signaling |
| AKT1 | Alpha-6 beta-4 integrin signaling pathway |
| RHOA | Alpha-6 beta-4 integrin signaling pathway |
| MAPK14 | Alpha-6 beta-4 integrin signaling pathway |
| EIF4EBP1 | Alpha-6 beta-4 integrin signaling pathway |
| MTOR | Alpha-6 beta-4 integrin signaling pathway |

| | |
|---|---|
| GAB1 | Alpha-6 beta-4 integrin signaling pathway |
| HRAS | Alpha-6 beta-4 integrin signaling pathway |
| ITGA6 | Alpha-6 beta-4 integrin signaling pathway |
| IRS1 | Alpha-6 beta-4 integrin signaling pathway |
| ITGB4 | Alpha-6 beta-4 integrin signaling pathway |
| LAMA2 | Alpha-6 beta-4 integrin signaling pathway |
| LAMA3 | Alpha-6 beta-4 integrin signaling pathway |
| LAMA5 | Alpha-6 beta-4 integrin signaling pathway |
| LAMB1 | Alpha-6 beta-4 integrin signaling pathway |
| LAMB2 | Alpha-6 beta-4 integrin signaling pathway |
| LAMB3 | Alpha-6 beta-4 integrin signaling pathway |
| LAMC1 | Alpha-6 beta-4 integrin signaling pathway |
| LAMC2 | Alpha-6 beta-4 integrin signaling pathway |
| PIK3R1 | Alpha-6 beta-4 integrin signaling pathway |
| PIK3R2 | Alpha-6 beta-4 integrin signaling pathway |
| PRKCA | Alpha-6 beta-4 integrin signaling pathway |
| PRKCD | Alpha-6 beta-4 integrin signaling pathway |
| MAPK1 | Alpha-6 beta-4 integrin signaling pathway |
| MAPK3 | Alpha-6 beta-4 integrin signaling pathway |
| PTK2 | Alpha-6 beta-4 integrin signaling pathway |
| PTPN11 | Alpha-6 beta-4 integrin signaling pathway |
| RAC1 | Alpha-6 beta-4 integrin signaling pathway |
| SHC1 | Alpha-6 beta-4 integrin signaling pathway |
| SRC | Alpha-6 beta-4 integrin signaling pathway |
| IRS2 | Alpha-6 beta-4 integrin signaling pathway |
| LAMA1 | Alpha-6 beta-4 integrin signaling pathway |
| AKT1 | Alpha-6 beta-1 and alpha-6 beta-4 integrin signaling |
| CASP7 | Alpha-6 beta-1 and alpha-6 beta-4 integrin signaling |
| CD9 | Alpha-6 beta-1 and alpha-6 beta-4 integrin signaling |
| CDH1 | Alpha-6 beta-1 and alpha-6 beta-4 integrin signaling |
| COL17A1 | Alpha-6 beta-1 and alpha-6 beta-4 integrin signaling |
| EGF | Alpha-6 beta-1 and alpha-6 beta-4 integrin signaling |
| EGFR | Alpha-6 beta-1 and alpha-6 beta-4 integrin signaling |
| ERBB2 | Alpha-6 beta-1 and alpha-6 beta-4 integrin signaling |
| ERBB3 | Alpha-6 beta-1 and alpha-6 beta-4 integrin signaling |
| SFN | Alpha-6 beta-1 and alpha-6 beta-4 integrin signaling |
| GRB2 | Alpha-6 beta-1 and alpha-6 beta-4 integrin signaling |
| HRAS | Alpha-6 beta-1 and alpha-6 beta-4 integrin signaling |
| IL1A | Alpha-6 beta-1 and alpha-6 beta-4 integrin signaling |
| ITGA6 | Alpha-6 beta-1 and alpha-6 beta-4 integrin signaling |
| ITGB1 | Alpha-6 beta-1 and alpha-6 beta-4 integrin signaling |
| ITGB4 | Alpha-6 beta-1 and alpha-6 beta-4 integrin signaling |
| LAMA2 | Alpha-6 beta-1 and alpha-6 beta-4 integrin signaling |
| LAMA3 | Alpha-6 beta-1 and alpha-6 beta-4 integrin signaling |
| LAMA4 | Alpha-6 beta-1 and alpha-6 beta-4 integrin signaling |
| LAMA5 | Alpha-6 beta-1 and alpha-6 beta-4 integrin signaling |
| LAMB1 | Alpha-6 beta-1 and alpha-6 beta-4 integrin signaling |
| LAMB2 | Alpha-6 beta-1 and alpha-6 beta-4 integrin signaling |
| LAMB3 | Alpha-6 beta-1 and alpha-6 beta-4 integrin signaling |
| LAMC1 | Alpha-6 beta-1 and alpha-6 beta-4 integrin signaling |

| | |
|---|---|
| LAMC2 | Alpha-6 beta-1 and alpha-6 beta-4 integrin signaling |
| MET | Alpha-6 beta-1 and alpha-6 beta-4 integrin signaling |
| MST1 | Alpha-6 beta-1 and alpha-6 beta-4 integrin signaling |
| MST1R | Alpha-6 beta-1 and alpha-6 beta-4 integrin signaling |
| PIK3CA | Alpha-6 beta-1 and alpha-6 beta-4 integrin signaling |
| PIK3R1 | Alpha-6 beta-1 and alpha-6 beta-4 integrin signaling |
| PMP22 | Alpha-6 beta-1 and alpha-6 beta-4 integrin signaling |
| PRKCA | Alpha-6 beta-1 and alpha-6 beta-4 integrin signaling |
| RAC1 | Alpha-6 beta-1 and alpha-6 beta-4 integrin signaling |
| RPS6KB1 | Alpha-6 beta-1 and alpha-6 beta-4 integrin signaling |
| RXRA | Alpha-6 beta-1 and alpha-6 beta-4 integrin signaling |
| RXRB | Alpha-6 beta-1 and alpha-6 beta-4 integrin signaling |
| RXRG | Alpha-6 beta-1 and alpha-6 beta-4 integrin signaling |
| SHC1 | Alpha-6 beta-1 and alpha-6 beta-4 integrin signaling |
| YWHAB | Alpha-6 beta-1 and alpha-6 beta-4 integrin signaling |
| YWHAE | Alpha-6 beta-1 and alpha-6 beta-4 integrin signaling |
| YWHAG | Alpha-6 beta-1 and alpha-6 beta-4 integrin signaling |
| YWHAH | Alpha-6 beta-1 and alpha-6 beta-4 integrin signaling |
| YWHAZ | Alpha-6 beta-1 and alpha-6 beta-4 integrin signaling |
| LAMC3 | Alpha-6 beta-1 and alpha-6 beta-4 integrin signaling |
| YWHAQ | Alpha-6 beta-1 and alpha-6 beta-4 integrin signaling |
| LAMA1 | Alpha-6 beta-1 and alpha-6 beta-4 integrin signaling |
| CRK | p130Cas linkage to MAPK signaling for integrins |
| PTK2 | p130Cas linkage to MAPK signaling for integrins |
| BCAR1 | p130Cas linkage to MAPK signaling for integrins |
| FES | Sema3A-plexin repulsion signaling by inhibiting integrin adhesion |
| FYN | Sema3A-plexin repulsion signaling by inhibiting integrin adhesion |
| PLXNA1 | Sema3A-plexin repulsion signaling by inhibiting integrin adhesion |
| PLXNA2 | Sema3A-plexin repulsion signaling by inhibiting integrin adhesion |
| RAC1 | Sema3A-plexin repulsion signaling by inhibiting integrin adhesion |
| RRAS | Sema3A-plexin repulsion signaling by inhibiting integrin adhesion |
| TLN1 | Sema3A-plexin repulsion signaling by inhibiting integrin adhesion |
| NRP1 | Sema3A-plexin repulsion signaling by inhibiting integrin adhesion |
| FARP2 | Sema3A-plexin repulsion signaling by inhibiting integrin adhesion |
| SEMA3A | Sema3A-plexin repulsion signaling by inhibiting integrin adhesion |
| PIP5K1C | Sema3A-plexin repulsion signaling by inhibiting integrin adhesion |
| RND1 | Sema3A-plexin repulsion signaling by inhibiting integrin adhesion |
| PLXNA3 | Sema3A-plexin repulsion signaling by inhibiting integrin adhesion |
| PLXNA4 | Sema3A-plexin repulsion signaling by inhibiting integrin adhesion |
| CRK | GRB2-SOS provides linkage to MAPK signaling for integrins |
| FGA | GRB2-SOS provides linkage to MAPK signaling for integrins |
| FGB | GRB2-SOS provides linkage to MAPK signaling for integrins |
| FGG | GRB2-SOS provides linkage to MAPK signaling for integrins |
| FN1 | GRB2-SOS provides linkage to MAPK signaling for integrins |
| GRB2 | GRB2-SOS provides linkage to MAPK signaling for integrins |
| ITGA2B | GRB2-SOS provides linkage to MAPK signaling for integrins |
| ITGB3 | GRB2-SOS provides linkage to MAPK signaling for integrins |
| PTK2 | GRB2-SOS provides linkage to MAPK signaling for integrins |
| RAP1A | GRB2-SOS provides linkage to MAPK signaling for integrins |
| RAP1B | GRB2-SOS provides linkage to MAPK signaling for integrins |

| | |
|---|---|
| SOS1 | GRB2-SOS provides linkage to MAPK signaling for integrins |
| SRC | GRB2-SOS provides linkage to MAPK signaling for integrins |
| TLN1 | GRB2-SOS provides linkage to MAPK signaling for integrins |
| VWF | GRB2-SOS provides linkage to MAPK signaling for integrins |
| BCAR1 | GRB2-SOS provides linkage to MAPK signaling for integrins |
| APBB1IP | GRB2-SOS provides linkage to MAPK signaling for integrins |
