## Supplementary file 7 for "Host transcriptional responses and SARS-CoV-2 isolates from the nasopharyngeal samples of Bangladeshi COVID-19 patients"

**Supplementary file 7: Differentially expressed genes in SARS-CoV-2 infected Lungs compared to the Bangladeshi Nasal samples used in this study.**

| Up-regulated in Lungs compared to Nasal samples (Bangladeshi) | Down-regulated in Lungs compared to Nasal samples (Bangladeshi) |
|---|---|
| CLCA4 | AC027290.3 |
| S100A2 | IL1RN |
| FGFBP1 | RN7SKP255 |
| AC091429.1 | CD53 |
| LYPD3 | SPI1 |
| RPL17P36 | PNRC1 |
| MTRNR2L1 | IFI44L |
| FOLH1B | AC073571.1 |
| RPL21P134 | MX2 |
| RPL41P2 | FBXO48 |
| EEF1A1P12 | ZFP36 |
| KRT6C | AL354919.1 |
| TPT1P9 | IFIT1 |
| KRT8P48 | AC016590.1 |
| AC209007.1 | H2BC4 |
| AC090543.3 | PTCHD4 |
| AC004453.1 | IFIT2 |
| EEF1A1P14 | DUSP5 |
| UNC93B2 | MUC5AC |
| SLC44A5 | MUC5B |
| H3C9P | AEN |
| YWHAZP4 | AC099489.1 |
| BX248409.1 | NR4A1 |
| FTH1P5 | SKAP2 |
| MT-TA | AC006435.4 |
| RPL10P4 | CROCC2 |
| AL133260.1 | H2BC8 |
| KRT8P5 | EGR1 |
| RPL31P63 | HOPX |
| AC091685.2 | EVI2A |
| PPIAP13 | SCARNA7 |
| EEF1A1P16 | C3AR1 |
| AC073072.1 | AC092299.7 |
| EEF1A1P22 | RN7SKP80 |
| LDHBP2 | AC015967.2 |
| EEF1A1P8 | FP236383.8 |
| FTH1P12 | H4C8 |
| PTMAP5 | H2BC7 |
| HSP90AA2P | MEG3 |
| RPS2P7 | MOCS1 |
| KRT8P3 | HTN3 |
| RPS27AP16 | RN7SKP203 |
| PPIAP16 | FP236383.7 |
| RPL17P22 | FAM27E3 |
| RPS18P12 | H2AC12 |
| HSPA8P1 | SNORA54 |
| PDIA3P1 | SLC38A5 |

| | |
|---|---|
| RPL10P6 | H19 |
| AC104563.1 | SIGLEC14 |
| AC022210.1 | MT-RNR1 |
| GAPDHP63 | JSRP1 |
| MT-TE | SNORA79B |
| RPL7P23 | VSIG4 |
| FTH1P16 | RN7SL396P |
| APOD | RNU5B-1 |
| PPIAP6 | SSTR5-AS1 |
| RPL24P4 | IGKC |
| GAPDHP1 | LINC01783 |
| SETP20 | AC011595.1 |
| AC104339.1 | RNU6ATAC |
| KRT8P32 | COL5A1 |
| MFSD3 | RN7SL471P |
| AC012005.1 | SNORD94 |
| BX679664.1 | HIST1H3B |
| AC025518.1 | SNORA74D |
| HLA-J | CCL19 |
| RPL7P15 | SNORA71D |
| EEF1A1P7 | H1-5 |
| RPL12P38 | SNORD17 |
| CROT | FRZB |
| RPS3AP5 | ADAMTS8 |
| AC068522.1 | RN7SKP71 |
| MTND6P3 | PLA2G5 |
| AL162430.2 | SNORA5A |
| RPS26P15 | SLC5A2 |
| AC090686.1 | AC010768.1 |
| AC092670.1 | AL121758.1 |
| PA2G4P6 | RNA5SP481 |
| RPL10AP6 | LRRC8C-DT |
| AC097658.2 | AC005515.1 |
| HNRNPA1P2 | IGF2 |
| RPL3P2 | RNA5-8SP6 |
| ANXA8 | AL161626.1 |
| HSPA8P5 | H2BC9 |
| AC006386.2 | SNORD89 |
| HNRNPKP4 | H3C11 |
| RPL13AP25 | RNA5SP74 |
| PPIAL4C | PRH2 |
| RPS13P2 | H2AC4 |
| PPIAP43 | NDUFA4L2 |
| AL049873.1 | COL5A3 |
| HSP90AA6P | AC024267.4 |
| ANXA8L1 | SNORA49 |
| SUMO2P1 | RN7SL778P |
| TPT1P5 | RN7SL151P |
| RPL7P19 | C19orf38 |
| PPIAP29 | H1-3 |
| AC005000.1 | AC024051.8 |
| RPS7P14 | AC024051.10 |
| RPL12P8 | AC024051.3 |
| RPS27P29 | AC024051.7 |
| RPS3AP25 | SNORA74A |
| AP002784.2 | RNA5SP162 |
| MTND5P11 | H4C4 |

| | |
|---|---|
| AL627402.1 | AC024051.1 |
| AL121871.1 | AL162581.1 |
| TAPT1-AS1 | RNU2-1 |
| RPS2P4 | AC105036.3 |
| RPS26P39 | KIAA1614-AS1 |
| MT-TL1 | RP11-180P8.3 |
| RPSAP19 | PTGDS |
| AC078819.1 | H4C3 |
| RPS3AP47 | AC027281.2 |
| FTH1P3 | ARHGAP26-AS1 |
| RPL15P18 | H1-4 |
| RPS20P14 | H4C6 |
| AL009174.1 | SCARNA5 |
| RPL23AP65 | KCNK12 |
| AC083873.1 | AC026369.2 |
| ACTA1 | SNORA80A |
| Z97353.1 | ADGRL3 |
| KRT8P45 | CTC-251I16.1 |
| RPL4P3 | FAM27B |
| RPL15P20 | RNA5SP225 |
| EEF1A1P29 | RPA4 |
| AC092865.1 | RNA5SP149 |
| EIF4A1P10 | AC087521.3 |
| AC100757.1 | HMGCLL1 |
| FTH1P20 | RNU1-67P |
| TMX1 | NCAM2 |
| IMP3 | PAPPA |
| AC104619.3 | KL |
| ST13P3 | RNA5SP141 |
| AC104257.1 | FP236383.5 |
| AC136632.1 | RNU4ATAC |
| AC020898.1 | MIR5188 |
| AL139095.2 | MAGI1-AS1 |
| EPHA1 | RNA5SP506 |
| HLA-G | FZD10 |
| TMSB4XP2 | HES5 |
| AC113404.3 | AC027514.1 |
| HSP90AB3P | MIR23A |
| EEF1A1P38 | AC024051.9 |
| COX6A1P2 | H4C12 |
| DTYMK | RNA5SP429 |
| ALG5 | RNA28S5 |
| AL162151.2 | B4GALNT1 |
| GAPDHP61 | AFF2 |
| RPL7AP11 | H2BC14 |
| DPYD | RNA5SP145 |
| KRT8P33 | AC005476.2 |
| YWHAZP3 | AL157895.2 |
| SCAMP3 | AC006449.2 |
| RPS26P28 | MIR3648-2 |
| AC073861.1 | AC025423.1 |
| RPL23P8 | AL355388.1 |
| MORF4L1P1 | SNORD15B |
| NACA3P | ALX4 |
| DENND10P1 | CCDC33 |
| PTN | REXO1L2P |
| AC010468.1 | RAG2 |

| | |
|---|---|
| UQCRFS1P1 | AC092612.1 |
| RPSAP61 | PCA3 |
| ALKBH3 | RNVU1-27 |
| RPS26P3 | DIRC3 |
| AC244034.1 | H4C11 |
| CRYZ | ROBO3 |
| AL355802.1 | MIR3648-1 |
| FTH1P11 | RN7SL752P |
| PTGES3P3 | RN7SL801P |
| RPSAP5 | RN7SL4P |
| YWHAZP5 | FP236383.10 |
| AC112187.1 | AL109920.1 |
| AL158206.1 | RN7SL5P |
| CDC42P6 | RNA5SP226 |
| RPL21P93 | FP236383.12 |
| RBM8B | FP236383.4 |
| XRCC6P2 | FP671120.7 |
| GAPDHP65 | PDZD4 |
| DBP | IDSP1 |
| TPI1P1 | AC010768.2 |
| EZH2 | OLFM5P |
| COQ5 | SNORA53 |
| AC092597.1 | PRB1 |
| CHCHD1 | RNU5A-1 |
| PRCP | H4C5 |
| PGDP1 | ERICH6B |
| FOLR1 | TMIGD3 |
| TOP2A | AC090970.1 |
| PIGO | SNORA74B |
| RPL7AP31 | RNA5SP161 |
| SIL1 | RNY4P6 |
| AC000089.1 | OR10A3 |
| TAGLN2P1 | LY86-AS1 |
| C1GALT1C1 | AC074135.1 |
| RPL10P12 | PRB2 |
| PFN1P1 | SCAT2 |
| RTN3P1 | AL135938.1 |
| PARP2 | RNA5-8SN1 |
| AC107032.1 | FP671120.4 |
| CHST12 | RNA5-8SN2 |
| HMGN2P46 | RNA5-8SN3 |
| GPR89A | PRB4 |
| GLULP4 | LINC00901 |
| HLA-DRB6 | AL031716.1 |
| TMEM179B | AL160408.1 |
| AC026271.1 | RNU4-1 |
| TMSB4XP4 | AC079601.1 |
| EIF4BP7 | RNY3 |
| ABHD3 | HCN2 |
| RPS23P8 | BAIAP2L2 |
| EIF4HP1 | AC024051.4 |
| MCM2 | AC004223.2 |
| DDX50 | RNVU1-31 |
| RPSAP18 | AC024051.6 |
| RPL26P19 | CNTN3 |
| SLC27A4 | AC004817.2 |
| TTC3P1 | CHRNA4 |

| | |
|---|---|
| B3GALT6 | RNA5SP502 |
| EIF4BP6 | AC024051.2 |
| SETP14 | MYH1 |
| AL080243.4 | RNA5SP387 |
| AP001324.1 | SNORA73B |
| RFNG | RNY3P1 |
| HYAL2 | RNA5SP298 |
| HMGB1P1 | DNASE2B |
| AGAP14P | DLK1 |
| AC022968.1 | AC024051.5 |
| RPL12P6 | AC024051.11 |
| PWP2 | FP671120.2 |
| EEF1B2P3 | CR392039.1 |
| FAM171A1 | FP236383.9 |
| HNRNPCP2 | RP11-216P16.8 |
| AC092115.2 | HNRNPA1P9 |
| EIF4BP3 | AC019070.2 |
| FKBP9P1 | RNA5SP389 |
| FP565260.1 | RN7SL753P |
| MMP15 | AC245008.1 |
| EIF4A1P2 | AL133368.1 |
| UNC50 | DCAF13P2 |
| ST13P5 | AL365209.1 |
| MMAB | AC009495.3 |
| HNRNPA3P6 | RNA5SP335 |
| EIF4A1P4 | AC024051.12 |
| ACTBP2 | MIR2278 |
| ZNF226 | SMIM9 |
| DPAGT1 | AC013403.2 |
| THNSL2 | PHOX2A |
| RDH14 | FAM27E2 |
| POLR2I | LINC01391 |
| AL391244.2 | IL24 |
| NPM1P39 | AC239859.1 |
| TMED1 | RP5-1011O1.2 |
| TMEM212 | RNY4 |
| RPL12P4 | SERTM2 |
| SERBP1P5 | RNVU1-2 |
| RPSAP4 | AC087239.1 |
| RNF26 | AC115989.1 |
| RAC1P2 | AC125793.1 |
| MIR22HG | CTD-2306M10.1 |
| TMEM147 | RNU4-2 |
| RPL7AP50 | NPTX1 |
| ABCB7 | FP236383.6 |
| MSH2 | SLC45A2 |
| MCM5 | VTRNA1-1 |
| ADH1A | RNA5S9 |
| LRRCC1 | RNU1-11P |
| ERO1B | RNA5SP202 |
| AP000936.3 | RNU1-3 |
| AL354702.1 | RNU1-1 |
| PTK7 | RNA5SP370 |
| SIRT3 | RNVU1-28 |
| RPL23AP7 | CDR1 |
| ALG6 | RNVU1-18 |
| EIF3FP3 | RNU1-2 |

| | |
|---|---|
| CFB | RNU1-4 |
| MSTO1 | RNA5S4 |
| MCM6 | RNA5S5 |
| STAG3L1 | RNA5S6 |
| SMG1P1 | RNA5S7 |
| TOPORS | RNA5S3 |
| TMTC4 | RNA5S2 |
| TAP2 | RNU1-88P |
| DNAJB9 | RNA5S1 |
| EIF2S2P4 | RNY1 |
| PMS1 | RNA5S12 |
| EXOSC8 | RNA5S17 |
| RAMAC | RNA5S16 |
| G6PC3 | RNA5S10 |
| SETSIP | RNA5S15 |
| NSMCE3 | RNA5S13 |
| TMEM43 | RNA5S8 |
| GPR87 | RNA5S11 |
| SRD5A3 | RNA5S14 |
| AC105250.1 | RNVU1-29 |
| H3-5 | RNVU1-7 |
| SMIM4 | RNU1-27P |
| LSAMP | RNU1-28P |
| FSCN1 | |
| COL6A3 | |
| CYP3A5 | |
| NR2C1 | |
| TMEM106C | |
| FRG1HP | |
| ANAPC4 | |
| BMI1 | |
| ATP5ME | |
| NAMPTP1 | |
| ABHD6 | |
| ILKAP | |
| WFS1 | |
| AC092490.1 | |
| OCLNP1 | |
| ACAD10 | |
| ARAP3 | |
| FBLN1 | |
| MITD1 | |
| CPQ | |
| BORCS5 | |
| AC115223.1 | |
| LTV1 | |
| SESN1 | |
| DSE | |
| CARM1 | |
| FKSG70 | |
| TCN1 | |
| PGM5P2 | |
| KLHL42 | |
| PROS1 | |
| FKSG61 | |
| ERVK3-1 | |
| EEF1A1P13 | |

| |
|---|
| B3GNT3 |
| RPS26P31 |
| AL663070.2 |
| ANXA2P2 |
| PPIAP31 |
| RPS26P47 |
| RPS7P1 |
| PPIAP22 |
| FAM3D |
| TP63 |
| RPL9P7 |
| GPX1P1 |
| EEF1A1P11 |
| MT-TY |
| PPIC |
| AC004057.1 |
| ATP1B3 |
| RPL3P4 |
| KRT5 |
| ANKRD66 |
| RPS3AP6 |
| RPP25L |
| AC024293.1 |
| EEF1A1P9 |
| RPS15AP1 |
| FTH1P7 |
| RPS26P8 |
| ADH1B |
| UBE2I |
| RPS24P8 |
| AC004552.1 |
| KRT6B |
| GALNT14 |
| KIT |
| ITGA6 |
| AC092683.1 |
| HMGB1P5 |
| FGFR3 |
| AL049597.1 |
| TMEM183B |
| SLC52A2 |
| NDUFA12 |
| H3P47 |
| MTRNR2L9 |
| DCAF13 |
| EEF1A1P25 |
| H3P6 |
| HMGB1P6 |
| RARRES1 |
| MST1R |
| MIR205HG |
| ACOT1 |
| MFSD5 |
| AC012085.1 |
| CLCA2 |
| SERPINB4 |
| ARSD |

| |
|---|
| MAD2L1BP |
| DPY30 |
| KRT6A |
| RHBDL2 |
| S100A4 |
| EMC7 |
| LPCAT3 |
| RPL13AP20 |
| DSG3 |
| APLP2 |
| ELMO3 |
| RPL7P1 |
| AC009245.1 |
| H3P16 |
| SVBP |
| THYN1 |
| IFNGR1 |
| MRPS21 |
| RPS7P11 |
| SRSF2 |
| AQP3 |
| ERAL1 |
| TEX264 |
| SLC44A3 |
| FCGRT |
| AC106795.1 |
| PHF14 |
| CYP2J2 |
| RPS2P55 |
| EDEM2 |
| ITGB6 |
| CD14 |
| JPT1 |
| FTLP3 |
| ATP10B |
| HNRNPA1P7 |
| SELENOS |
| BZW1P2 |
| UQCC2 |
| RPL7P9 |
| RPS26P6 |
| MINPP1 |
| PLLP |
| TBL2 |
| PDIA3 |
| DDAH2 |
| ITM2C |
| RPN2 |
| PSPC1 |
| TUBAP2 |
| NPC1 |
| EEF1A1P19 |
| ECI1 |
| LAMC1 |
| SLC35B2 |
| ARMT1 |
| EXOSC9 |

| |
|---|
| CLDN1 |
| HIBCH |
| FMO5 |
| RPL7P32 |
| SNHG6 |
| PRDX4 |
| RPS2P46 |
| PRPF39 |
| PSME1 |
| SCYL3 |
| AC064799.1 |
| UBBP4 |
| MANF |
| UXS1 |
| PAPSS2 |
| SPINT2 |
| RPL7AP6 |
| RPL21P28 |
| TST |
| RMDN3 |
| TUSC3 |
| METTL17 |
| SCPEP1 |
| NOMO2 |
| PBXIP1 |
| PYCARD |
| CTSC |
| CD9 |
| HLA-A |
| PRMT7 |
| CDS2 |
| CCDC59 |
| AC113935.1 |
| SMC4 |
| LMAN2 |
| FANCI |
| HNRNPA1P48 |
| RPL14P1 |
| AL135745.1 |
| GAS6 |
| GPN1 |
| PGRMC1 |
| ATRAID |
| TSPAN6 |
| MTCO2P12 |
| CCT6A |
| HEXB |
| FKBP9 |
| TMCO1 |
| CTSH |
| MFSD11 |
| HLA-F |
| ETHE1 |
| SNX14 |
| DNAJB11 |
| PFKM |
| ASB3 |

| |
|---|
| HLA-H |
| EEF1A1P4 |
| LAMB3 |
| BCAP31 |
| UPK1B |
| SLC39A6 |
| GUSB |
| ITM2B |
| NDUFAB1 |
| PPIAL4G |
| C1QBP |
| PTDSS1 |
| RUFY2 |
| KRT8 |
| KTN1 |
| AC090498.1 |
| SDHA |
| DSC3 |
| CXADR |
| SNRPD2 |
| KDSR |
| HLA-C |
| UFD1 |
| THOC3 |
| HADH |
| HSP90B1 |
| ATP6AP1 |
| HSP90AA1 |
| RRAGA |
| LRRC8D |
| ATP6AP2 |
| PSMD1 |
| EBPL |
| ARMC1 |
| RPL22P1 |
| TMEM9 |
| AGR2 |
| GBA |
| NPTN |
| UQCRC1 |
| PSENEN |
| CUEDC1 |
| CHPF |
| MAPKAPK3 |
| BX679664.3 |
| PLS1 |
| CALR |
| AC026403.1 |
| CLK1 |
| PRSS8 |
| RCN2 |
| SNRPA1 |
| RRM1 |
| VARS1 |
| ADAM28 |
| AC233968.1 |
| CDH1 |

| |
|---|
| CTNNAL1 |
| RPL13P12 |
| BCAT2 |
| SMARCAD1 |
| IL10RB |
| ACSF2 |
| CAMK2G |
| CHKA |
| PLTP |
| GFM2 |
| ARL3 |
| RPAP2 |
| SNHG1 |
| ATP5F1B |
| AC233699.1 |
| RPN1 |
| RPL10P16 |
| ACOT2 |
| GTF2H2B |
| GAA |
| CD81 |
| NOMO1 |
| TWF2 |
| ABCE1 |
| GTF2H2C |
| CHPT1 |
| RPL13AP5 |
| C5orf15 |
| LUC7L3 |
| RARS1 |
| NT5DC1 |
| SURF1 |
| PRDX1 |
| LRG1 |
| ERMP1 |
| CKMT1A |
| RPS2P5 |
| ITFG1 |
| CRYM |
| AL592114.1 |
| CEACAM5 |
| LRP5 |
| TAP1 |
| F11R |
| SRSF11 |
| AKAP1 |
| HSD17B13 |
| FAT1 |
| ANXA4 |
| HSD17B12 |
| TXNDC15 |
| RTN3 |
| KIFAP3 |
| CAPG |
| DNAJC1 |
| ARF5 |
| GCLC |

| |
|---|
| SDF4 |
| TMEM59 |
| KRT19 |
| CDH3 |
| HLA-B |
| GSTK1 |
| TM9SF2 |
| ADRM1 |
| PCNA |
| MAN1B1 |
| ANKRD36 |
| WDR61 |
| RCN1 |
| MPP7 |
| PDIA6 |
| KRT15 |
| FHL2 |
| DDOST |
| TSPAN13 |
| AGL |
| ITGAV |
| SLC35A2 |
| ASCC3 |
| TRAPPC3 |
| GSS |
| GRN |
| RASA1 |
| STAM2 |
| PRELID1 |
| SMG1P4 |
| PRKAR2B |
| NBEAL1 |
| ERLEC1 |
| RPL7AP66 |
| GNS |
| CFH |
| P4HB |
| CD47 |
| CTSB |
| GLRX5 |
| RPS21 |
| AMFR |
| MGST1 |
| B2M |
| HACD3 |
| CLDN4 |
| BCAP29 |
| WASHC2C |
| APP |
| FDFT1 |
| VPS25 |
| ATP1A1 |
| ALDH7A1 |
| PRXL2A |
| ADAM15 |
| SREK1 |
| CD151 |

| |
|---|
| TACSTD2 |
| ARL6IP1 |
| TOR3A |
| MDH1 |
| MORN2 |
| ATP2C1 |
| OS9 |
| MIPEP |
| AKR1A1 |
| DSG2 |
| SELENBP1 |
| ATP5MG |
| TSPAN1 |
| PPIB |
| OSBPL3 |
| TMEM50B |
| MBOAT2 |
| SYNGR2 |
| S100A6 |
| WDR33 |
| SLC44A2 |
| SOD1 |
| HNRNPA1P10 |
| ZMPSTE24 |
| TK2 |
| ST14 |
| TMEM9B |
| PTTG1IP |
| AC087473.1 |
| EPCAM |
| ANXA1 |
| MAT2B |
| KARS1 |
| EFTUD2 |
| TMEM87A |
| GPD2 |
| RPL21P16 |
| SLC38A10 |
| PSMB3 |
| RBM5 |
| EXOC1 |
| ARPC3 |
| MTDH |
| RETREG2 |
| REEP5 |
| ERBB3 |
| RBM6 |
| ATP6V0D1 |
| MAP3K5 |
| ALDH2 |
| TUFM |
| RPS26 |
| PSAP |
| ATP5MC3 |
| CD63 |
| POR |
| UBXN1 |

| |
|---|
| PGD |
| ALDH1A1 |
| DSP |
| LRPAP1 |
| ENO1 |
| PTPRK |
| GALNT7 |
| TOP2B |
| SARAF |
| ATXN10 |
| CCNDBP1 |
| YWHAB |
| HSPA5 |
| QSOX1 |
| EEF1A1P5 |
| SRI |
| OCIAD1 |
| EXOC3 |
| DDX19A |
| EMC4 |
| SDC1 |
| CCT5 |
| HTATIP2 |
| MMP14 |
| PKM |
| ANXA3 |
| SLC39A7 |
| TUBA1A |
| ARPC1B |
| TAGLN2 |
| ATP5F1A |
| USP48 |
| PCYOX1 |
| HMGB1 |
| PCMTD2 |
| SRSF3 |
| CLSTN1 |
| GSN |
| PPP1CA |
| DDB1 |
| NUCB2 |
| LRIG1 |
| HNRNPAB |
| CLPTM1L |
| SMARCA2 |
| LGALS3BP |
| ARPC5 |
| PYGL |
| ANKRD36B |
| RTN4 |
| PSMD4 |
| NDUFV1 |
| HNRNPA2B1 |
| ANPEP |
| PLD3 |
| NRBP1 |
| IFT57 |

| |
|---|
| CTNNB1 |
| COPB1 |
| ACSL5 |
| DDX5 |
| ACTG1 |
| COX4I1 |
| SRPRA |
| GDE1 |
| SEPHS2 |
| CAST |
| PRKCSH |
| LDHB |
| HNRNPL |
| TSPYL1 |
| NFE2L1 |
| EIF3C |
| EIF3CL |
| SFPQ |
| DHCR24 |
